# Heightened lateral habenula activity during stress produces brainwide and behavioral substrates of susceptibility

**DOI:** 10.1101/2023.11.06.565681

**Authors:** Anna Zhukovskaya, Christopher A. Zimmerman, Lindsay Willmore, Alejandro Pan-Vazquez, Sanjeev R. Janarthanan, Annegret L. Falkner, Ilana B. Witten

**Affiliations:** Princeton Neuroscience Institute, Princeton University, Princeton NJ 08544 USA

## Abstract

Some individuals are susceptible to the experience of chronic stress and others are more resilient. While many brain regions implicated in learning are dysregulated after stress, little is known about whether and how neural teaching signals during stress differ between susceptible and resilient individuals. Here, we seek to determine if activity in the lateral habenula (LHb), which encodes a negative teaching signal, differs between susceptible and resilient mice during stress to produce different outcomes. After, but not before, chronic social defeat stress (CSDS), the LHb is active when susceptible mice are in the proximity of the aggressor strain. During stress itself, LHb activity is higher in susceptible mice during aggressor proximity, and activation of the LHb during stress biases mice towards susceptibility. This manipulation generates a persistent and widespread increase in the balance of subcortical versus cortical activity in susceptible mice. Taken together, our results indicate that heightened activity in the LHb during stress produces lasting brainwide and behavioral substrates of susceptibility.

## Introduction

Chronic stress increases the risk of developing mental illness^1–3^. However, some individuals are more susceptible to the adverse effects of chronic stress, whereas others are more resilient. Pioneering work has used chronic social defeat stress (CSDS) to model this variability in rodents^4–14^. This prior work has largely focused on identifying the factors that predispose to susceptibility to stress^15–20^, or on identifying the changes in the brain after stress^4,5,10,11,21–26^. However, the question of how activity differs between susceptible and resilient individuals during stress itself in order to lead to different stress outcomes remains largely unaddressed.

To begin to address this, we recently developed approaches to automatically identify relevant behaviors during social defeat from video recordings (e.g. being attacked, fighting back)^27^. Using these tools, we observed distinct neural correlates in the midbrain dopamine system (a key component of the brain’s positive reinforcement system^28–32^) across resilient and susceptible mice: more dopamine neuron activity during an “active coping” strategy (i.e. fighting back behavior) in resilient mice, and more dopamine during escape in susceptible animals. Together, this may help explain how individuals develop resilient versus susceptible strategies. However, these differences in the dopamine system between resilient and susceptible mice emerge gradually across the ten days of defeat, leaving open the question of if there are important differences in other populations that emerge earlier.

Given that stressors have negative valence, we hypothesize that differences in the aversive learning system may be present earlier during stress and be critical to the formation of the susceptible state. The lateral habenula (LHb) is a key region for aversive learning. It encodes a negative reward prediction error^33–35^, drives aversive learning^36,37^, and is dysregulated by stress^21,38–46^. However, if and when the LHb first responds differentially to stressors in susceptible versus resilient individuals, and whether such differences are causal to the development of the susceptible state, remains unknown.

To address this, we recorded longitudinally from the LHb before, during, and after CSDS, and found elevated activity in the LHb in susceptible mice starting on the first day of stress. To determine if these differences in the LHb between susceptible and resilient mice are causal to susceptibility, we activated the LHb during defeat stress, which biased animals towards the susceptible phenotype. Finally, we examined the effects of LHb stimulation during defeat on brainwide activity, and found that LHb activation generated a sustained increase in the balance of subcortical versus cortical activity in susceptible mice.

## Results

### CSDS produces strain-specific social aversion and anxiety-like behaviors

Male mice underwent 10 days of chronic social defeat stress (CSDS), where they were defeated by a new aggressor for 5 minutes a day, and housed with the aggressor (that was separated by a barrier) for the remainder of each day (Figure 1A). CSDS was preceded by assays of sociability and followed both by assays of sociability and of anxiety-like behavior (Figure 1B-N). Consistent with previous studies^4,5,9,47,48^, a subset of mice showed decreased social interaction time with the aggressor strain after (but not before) CSDS in a “social interaction” (SI) test when the social target was behind a barrier (Figure 1B-C). Mice were defined as susceptible if their social interaction time in the SI test was less than one standard deviation below that of unstressed controls^27^; otherwise, they were considered resilient (Figure 1C).

**Figure 1.**
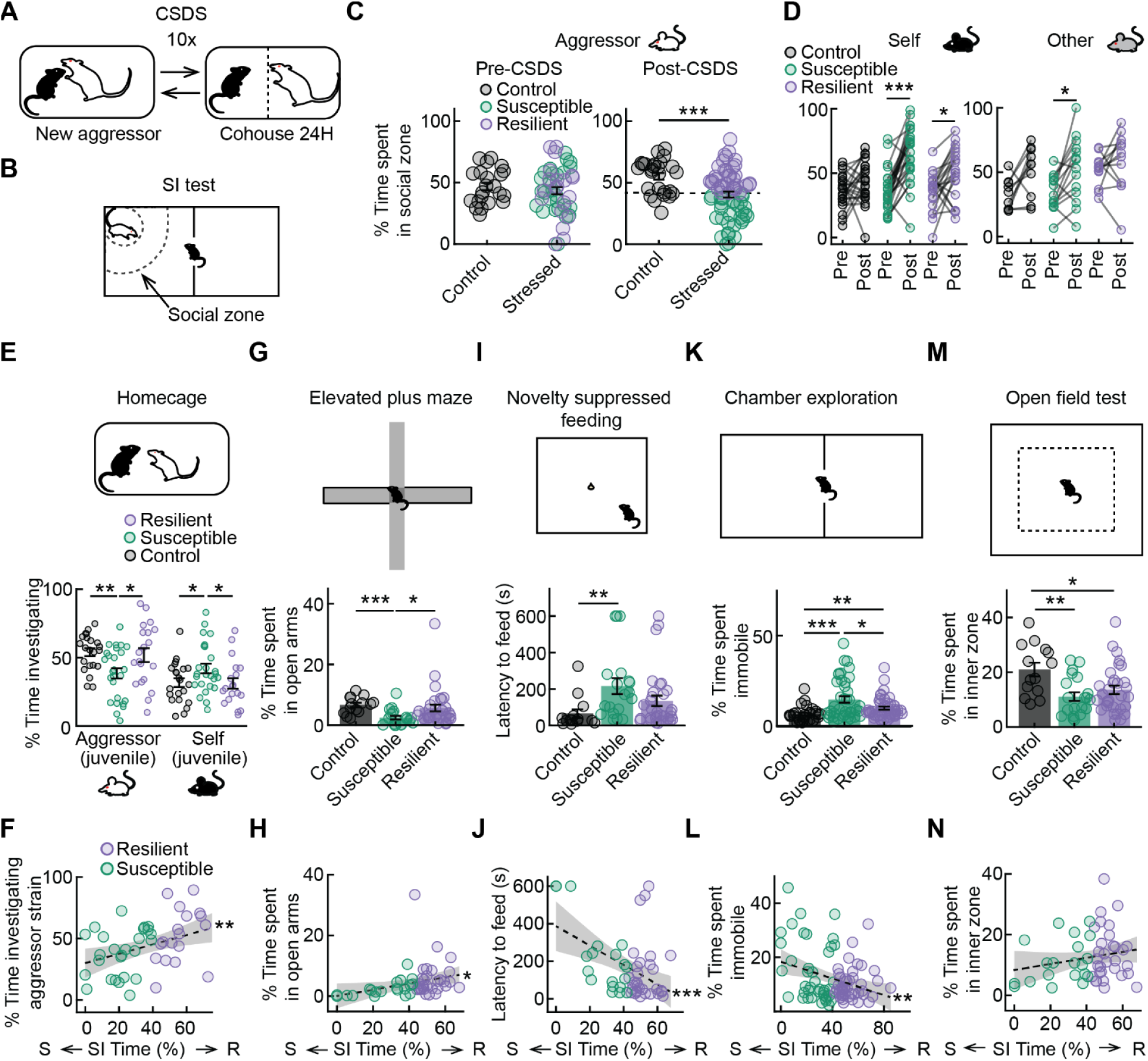
In susceptible mice, CSDS produced strain-specific aversion and changes in anxiety-like behavior and mobility. **A**. Schematic of Chronic Social Defeat Stress (CSDS) behavioral paradigm. **B**. Schematic of Social Interaction (SI) test. Social zone: 8 cm additional radius from the perimeter of the cup containing the social target. **C**. Left: time spent near the aggressor strain in SI test pre-CSDS. Right: time spent near the aggressor strain in SI test post-CSDS. Dashed line indicates the cutoff for binary categorization of Susceptible/Resilient (based on one standard deviation below the control mean). Control mice vs stressed mice post-CSDS: *t* = 3.5505, *p* = 6.1699e-04. **D**. Left: Time spent in the social zone (SI time) before vs after CSDS when the social target was of the self (BL6) strain. Right: time spent in the social zone (SI time) before vs after CSDS when the social target was of the other (AKR) strain. Self strain susceptible pre-CSDS vs self strain susceptible post-CSDS: *t* = −6.1047, *p* = 1.33e-05. Self strain resilient pre-CSDS vs self strain resilient post-CSDS: *t* = −2.9147, *p* =0.0393. Other strain susceptible pre-CSDS vs other strain susceptible post-CSDS: *t* = −3.1374, *p* =0.0393. **E**. Top: schematic of homecage assay. Bottom: percent of time spent investigating (sniffing and pursuing) social target in freely-moving assay when the social target was a juvenile of the aggressor strain or a juvenile of the self strain (control *N* = 22, susceptible *N* = 26, resilient *N* = 20). Control vs susceptible for aggressor social target: *t* = 3.2324, *p* = 0.0023. Susceptible vs resilient vs for aggressor social target: *t* = −2.1898, *p* = 0.0339. Control vs susceptible vs for self-strain social target: *t* = −2.1285, *p* = 0.0387. Susceptible vs resilient for self-strain social target: *t* = 2.0549, *p* = 0.0460. **F.** Relationship between time spent investigating a juvenile from the aggressor strain in the homecage assay and SI time after CSDS: *R* = 0.3965, *p* = 0.0064. **G**. Top: Schematic of elevated plus maze (EPM). Bottom: percent of time spent in open arms of EPM (control *N* = 14, susceptible *N* = 20, resilient *N* = 31). Control vs susceptible: *t* = 4.5352, *p* = 7.6295e-05. Susceptible vs resilient: *t* = −2.1630, *p* = 0.0354. **H**. Relationship between time spent in open arms of the EPM and SI time: *R* = 0.3100, *p* = 0.0269. **I**. Top: schematic of novelty suppressed feeding assay (NSF). Bottom: Latency to feed during NSF (control *N* = 14, susceptible *N* = 19, resilient *N* = 31). Control vs susceptible: *t* = −2.7623, *p* = 0.0096. **J**. Relationship between latency to feed in NSF and SI time: *R* = −0.4584, *p* = 0.0008. **K**. Top: Schematic of chamber exploration assay. Bottom: Percent of time immobile (speed <1cm/s) during chamber exploration (control *N* = 30, susceptible *N* = 40, resilient *N* = 46). Control vs susceptible: *t* = −3.7733, *p* = 3.4041e-04. Susceptible vs resilient: *t* = −2.5642, *p* = 0.0102. Control vs resilient: *t* = −2.7313, *p* = 0.0079. **L**. Relationship between time spent immobile during chamber exploration and SI time: *R* = −0.3119, *p* =0.0035. **M**. Top: Schematic of open field test (OFT). Bottom: percent of time spent in inner zone of OFT (control *N* = 14, susceptible *N* = 20, resilient *N* = 31). Control vs susceptible: *t* = 3.5396, *p* = 0.0013. Control vs resilient: *t* = 2.6244, *p* = 0.0102. **N**. Relationship between time spent in inner zone of OFT and SI time. *p*-value in **C** is from an unpaired 2-sided *t*-test. *p*-values in **D** are from paired 2-sided *t*-tests (with Bonferroni correction for three groups and two strains). *p*-values in **E**, **G**, **I**, **K**, **M** are from unpaired 2-sided *t*-tests following 1-way ANOVA. *p*-values in **F**, **H**, **J**, **L** are from Pearson’s correlations. Shaded areas in **F**, **H**, **J, L**, **N** represent 95% confidence interval for linear fit. \**p* ≤ 0.05, \*\**p* ≤ 0.01, \*\*\**p* ≤ 0.001. See Supplementary Table 1 for statistics summary (including ANOVA results).

As expected, susceptibility by this measure correlated with social avoidance of a juvenile of the aggressor strain in a freely moving assay (Figure 1E-F), as well as with higher anxiety-like behavior in non-social settings (elevated plus maze: Figure 1G-H; novelty-suppressed feeding: Figure 1I-J; immobility in a neutral context: Figure K-L, for pre-CSDS data see Figure S1A; open field test: Figure 1M-N)^7,8,27,39,49,50^,. These correlates were not apparent in unstressed controls (Figure S1B-F). Susceptibility did not generalize to social avoidance of the self strain or a control strain; similar to resilient mice, susceptible mice spent significantly more time with their own strain and another third strain after stress (Figure 1D for SI test; Figure 1E-F for freely moving assay).

Thus, in susceptible mice, CSDS produced a generalized anxiety-like phenotype in non-social contexts, while also generating strain-specific social avoidance. The observation of strain-specific avoidance learning as a result of CSDS is consistent with recent work^51,52^ (but see ^53^ which used longer defeat sessions and instead observed generalization of avoidance across strains).

### CSDS produces neural correlates of strain-specific aversion in the LHb in susceptible mice

To determine whether neural activity in the lateral habenula (LHb) before and after CSDS relates to the observed strain-specific avoidance learning (Figure 1B-F), we used fiber photometry (Figure 2A-J, histology summary: Figure S2A-B) and cellular resolution calcium imaging (Figure 2K-Z, histology summary: Figure S2C) to record responses to the aggressor strain, the defeated mouse’s own strain, and a control strain in the pre- and post-CSDS SI tests. In our fiber photometry experiments, we used wild type mice and a pan-neuronal GCaMP virus. In our cellular resolution experiments, we used vGlut2-Cre mice and a Cre-dependent GCaMP virus in order to focus on glutamatergic cells, as they are the predominant cell type in LHb^54^ and previous work has shown that they respond to aggressive interactions^39^.

**Figure 2.**
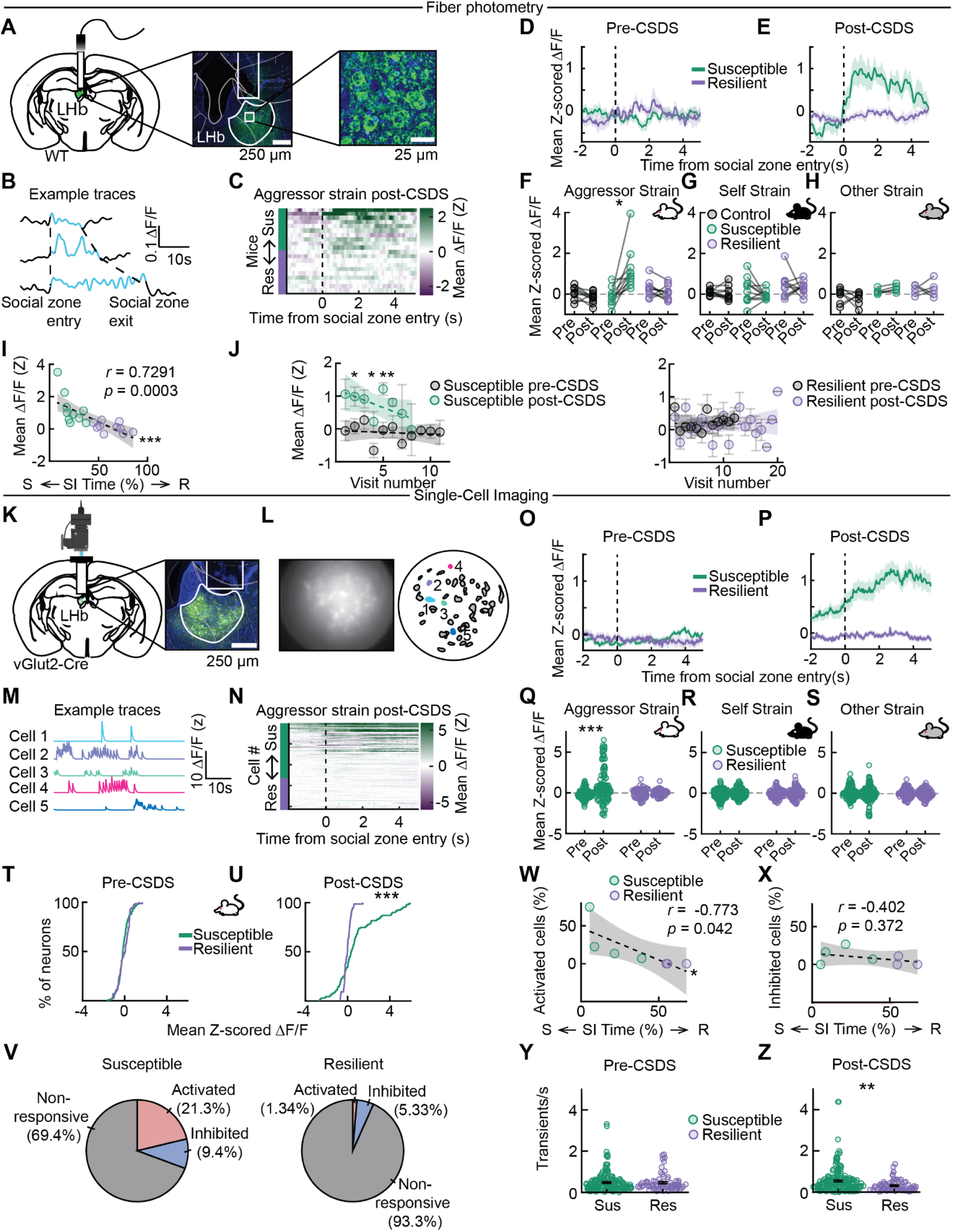
After but not before CSDS, aggressor strain-specific responses in the LHb of susceptible mice in the SI test. **A**. Left: Location of fiber photometry recordings from cell bodies in the Lateral Habenula (LHb) Middle: GCaMP (AAV5-CaMKII-GCaMP6f or AAV5-syn-jGCaMP7f) expression in LHb cell bodies in green and DAPI in blue. Right: A confocal image of LHb neurons showing nuclear exclusion of GCaMP. **B**. Responses in an example mouse to individual visits to the social zone of the aggressor (Figure 1B). Trials are organized in ascending order of time spent in the social zone. **C**. LHb signal aligned to entry of aggressor (SW) social zone during SI test. Each row is mean response in one mouse, with mice sorted by SI time, from susceptible (green, *N* = 10) to resilient (purple, *N* = 10). **D**. LHb signal aligned to entry of aggressor social zone during pre-CSDS SI test, averaged across individuals in resilient and susceptible groups (mean ± s.e.m. plotted). **E**. Same as **D** for LHb signal aligned to entry of aggressor social zone during post-CSDS SI test. **F**. Average from 1 s to 2 s post onset of entry to the social zone in **D** and **E** for susceptible (*N* = 10), resilient (*N* = 10), and control mice (*N* = 10). Susceptible pre-CSDS vs susceptible post-CSDS: *t* = −3.7842, *p* = 0.0389. **G**. Same as **F** for self-strain (BL6) social zone entry. **H**. Same as **F** for other strain (AKR) social zone entry. **I**. Correlation between the magnitude of the fluorescence response during the SI test with the aggressor strain post-CSDS and avoidance level in mice from fiber photometry experiments. *r* = −0.7291, *p* = 0.0003. **J**. Neural response as a function of visit number across mice for susceptible (left) and resilient (right) mice pre-CSDS vs post-CSDS. **K**. Left: Cellular resolution calcium imaging schematic. Right: Example histology with GRIN lens placement above LHb and AAV9-syn-FLEX-GCaMP7f expression in green and DAPI in blue. **L**. Left: Example FOV from microendoscope. Right: Same example FOV, with identified neurons outlined. **M**. Example traces of neurons with filled-in colors in **L**. **N**. LHb signal aligned to entry of aggressor (SW) social zone during SI test. Each row is a neuron, sorted from susceptible (green, n = 131 neurons, *N* = 4 mice) to resilient (purple, n = 75 neurons, *N* = 5 mice). **O**. LHb signal aligned to entry into aggressor (SW) social zone during pre-CSDS SI test, averaged across neurons in resilient and susceptible groups (mean ± s.e.m. plotted). **P**. Same as **O**, for LHb signal aligned to entry into aggressor (SW) social zone during post-CSDS SI test. **Q**. Average from 1 s to 2 s post entry into aggressor (SW) social zone in **O** and **P** plotted for susceptible and resilient groups. Susceptible pre-CSDS vs susceptible post-CSDS: *t* = −5.8304, *p* <.0001. **R**. Same as **Q** for self-strain (BL6) social zone entry. **S**. Same as **Q** for other strain (AKR) social zone entry. **T**. Distribution of responses in resilient and susceptible mice during aggressor strain proximity in the SI test before CSDS. **U**. Distribution of responses in resilient and susceptible mice during aggressor strain proximity in the SI test after CSDS. Susceptible vs resilient post-CSDS: *k =* 0.4123, *p* = 9.5142e-08. **V**. Proportion of cells that were significantly responding during aggressor proximity during the SI test after defeat in susceptible (left) and resilient (right) mice (see Figure S2**D**-**G** and Methods). **W**. Correlation between the magnitude of the fluorescence response during aggressor strain proximity in the SI test post-CSDS and avoidance level in significantly activated cells: *r* = −0.7732, *p* = 0.0414. **X**. Correlation between the magnitude of the fluorescence response during aggressor strain proximity in the SI test post-CSDS and avoidance level in significantly inhibited cells. **Y**. Spontaneous event rates in susceptible (green, n = 165 neurons, *N* = 6 mice) and resilient (purple, n = 71 neurons, *N* = 5 mice) mice during a 5 min test in a neutral chamber pre-CSDS. **Z**. Spontaneous event rates in susceptible (green, n = 182 neurons, N = 6 mice) and resilient (purple, n = 75 neurons, *N* = 5 mice) mice during a 5 min test in a neutral chamber post-CSDS. Susceptible vs resilient *t* = −3.0187, *p* = 0.0028. *p*-value in **F** is from a paired 2-sided *t*-test (with Bonferroni correction for three groups and three strains). *p*-value in **Q** is from an unpaired 2-sided *t*-test (with Bonferroni correction for two groups and three strains). *p*-value in **U** is from a Kolmogorov-Smirnov test. *p*-values in **I**, **W**-**X** are from Pearson’s correlations. *p*-values in **J**, **Z** are from 2-sided *t*-tests. Shaded areas in **I**-**J**, **W**-**X** represent 95% confidence interval for linear fit. \**p* ≤ 0.05, \*\**p* ≤ 0.01, \*\*\**p* ≤ 0.001. See Supplementary Table 1 for detailed statistics summary.

The fiber photometry recordings during the SI test revealed no modulation of the LHb to any strain in either susceptible or resilient mice before CSDS (Figure 2D). After CSDS, there was elevated activity in susceptible but not resilient mice, specifically to the aggressor strain (Figure 2E-J). Consistent with this, responses in the social zone after CSDS were inversely correlated with SI time (Figure 2I).

Given that resilient mice visit the aggressor more (Figure 1C), we sought to determine if elevated activity during aggressor visits in susceptible mice could be a consequence of attenuation of neural activity as a function of visits to the aggressor in resilient mice (i.e. adaptation). Contradicting this idea, responses did not significantly attenuate with visit number for resilient mice (Figure 2J). In susceptible mice there was response attenuation with visit number after (but not before) CSDS (Figure 2J).

Similar to the fiber photometry data (Figure 2D, F-J), the cellular resolution imaging during the SI test revealed that, on average, LHb neurons were not modulated by any strain prior to CSDS (Figure 2O, Q-S). Following CSDS, on average, LHb neurons of susceptible mice were activated by the aggressor strain and not other strains (Figure 2P-U), also similar to the fiber photometry data (Figure 2E-J).

We next examined the heterogeneity of cellular responses by identifying cells that were significantly activated or inhibited by the aggressor (Figure S2D-G; see Methods). Susceptible mice had many more activated cells, and slightly more inhibited cells (Figure 2V). Furthermore, the magnitude of the fluorescence response during the SI test with the aggressor strain post-CSDS was inversely correlated with avoidance level in activated (but not inhibited) cells (Figure 2W-X).

Though neither susceptible nor resilient mice had increased LHb activity to the social stimuli prior to CSDS (Figure 2O, Q-S), we considered whether spontaneous transient rates may differ prior to stress. Similar to previous work, we found that spontaneous transient rates were similar in susceptible and resilient mice preceding CSDS, but higher in susceptible mice following CSDS, when mice explored a neutral chamber for 5 minutes (Figure 2Y-Z, Figure S2H)^55^.

Thus, while social responses before CSDS were not apparent in the LHb in susceptible or resilient mice, stress produced neural correlates of strain-specific aversion in susceptible mice. We next sought to determine how LHb activity relates to behavior during stress itself, and if and when stress-related activity first differed between susceptible and resilient mice.

### LHb activity during defeat is elevated during attacks and other proximal behaviors

We performed high-speed, multiview videography during each defeat session, followed by automated behavioral quantification (Figure 3A). Key points in both mice were tracked (Figure 3A) and used to define 12 features - such as relative orientation and distance between the mice - to capture the postures, positions and movements of the animals (Figure 3B; see Methods for explanation of feature calculation). These features from each frame were then embedded into a 2D *t*-distributed stochastic neighbor embedding (*t-*SNE) manifold (Figure 3C, Figure S3A), which was followed by density-based clustering to define distinct social and non-social behaviors^27^. Clusters, which had similar occupancy between susceptible and resilient mice (Figure S3B), were numbered by proximity between the mice.

**Figure 3.**
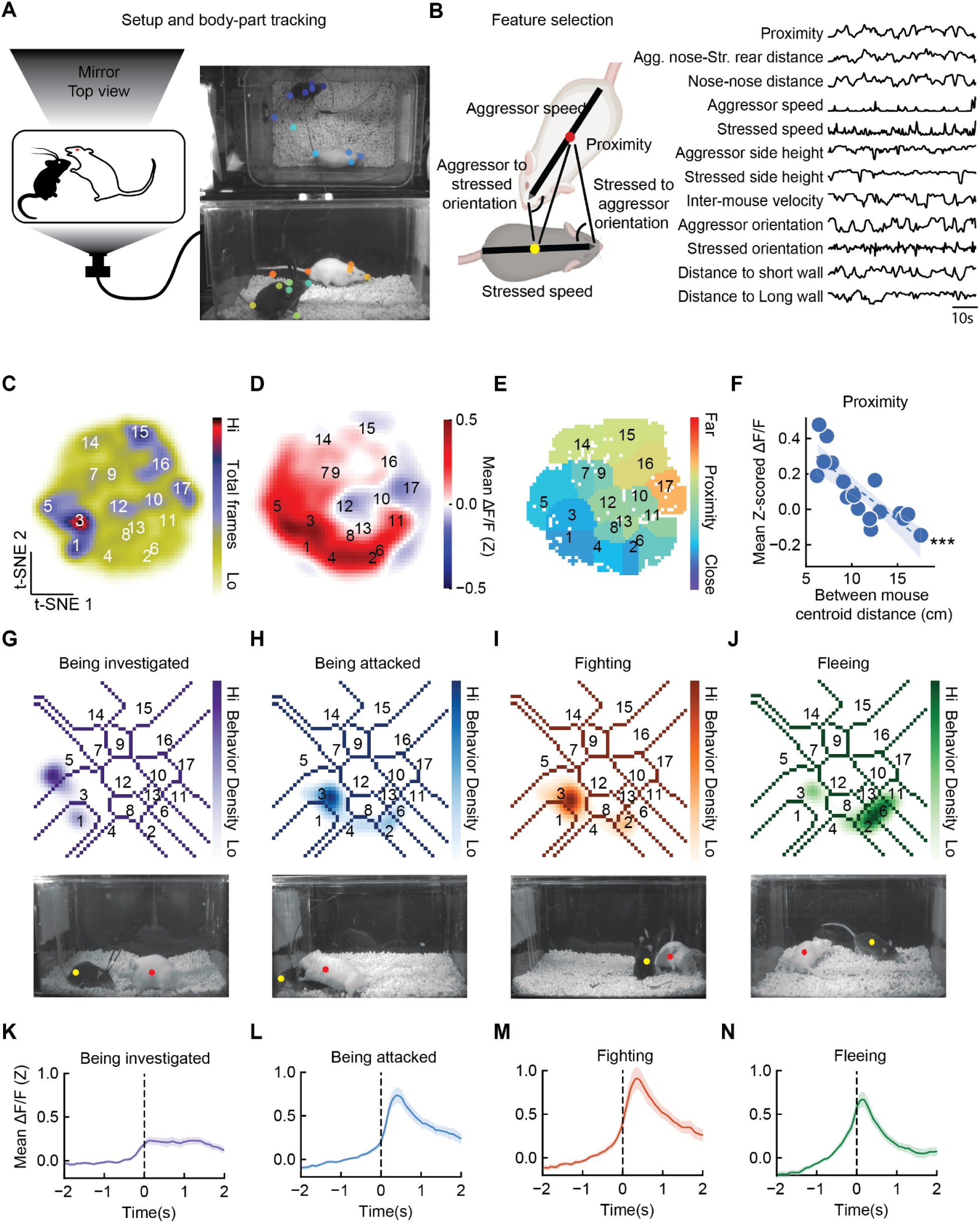
During CSDS, elevated LHb activity during attack and other proximal behaviors. **A**. Left: Behavioral setup. Right: Example video frame with tracked key points. **B**. Left: Features calculated from key points. Right: Time series of all features used in behavior quantification. **C**. Smoothed histogram of *t*-SNE embeddings from features, with clusters numbered by increasing distance between mice (*N* = 35). **D**. Mean LHb GCaMP signal across *t*-SNE behavior space in fiber photometry mice (*N* = 21). **E**. Average proximity within each *t*-SNE cluster. **F**. For each cluster, mean LHb GCaMP signal plotted against mean centroid distance between mice (*R* = −0.8215, *p* = 5.3E-5, *N* = 17 clusters). **G**. Top: Density of random forest classified investigation within *t*-SNE space. Bottom: Example frame of being investigated. Stressed mouse: yellow dot; aggressor mouse: red dot. **H**. Top: Density of random forest classified attack within *t*-SNE space. Bottom: Example frame of attack. Stressed mouse: yellow dot; aggressor mouse: red dot. **I**. Density of random forest classified fighting within *t*-SNE space. Bottom: Example frame of fighting. Stressed mouse: yellow dot; aggressor mouse: red dot. **J**. Density of random forest classified fleeing within *t*-SNE space. Bottom: Example frame of fleeing. Stressed mouse: yellow dot; aggressor mouse: red dot. **K**. Neural activity in LHb time-locked to being investigated (mean ± s.e.m. plotted). **L**. Neural activity in LHb time-locked to being attacked (mean ± s.e.m. plotted). **M**. Neural activity in LHb time-locked to fighting (mean ± s.e.m. plotted). **N**. Neural activity in LHb time-locked to fleeing (mean ± s.e.m. plotted). *p*-value in **F** is from a Pearson’s correlation. Shaded area in **F** represents 95% confidence interval for linear fit. \**p* ≤ 0.05, \*\**p* ≤ 0.01, \*\*\**p* ≤ 0.001. See Table 1 for detailed statistics summary. See Supplementary Table 1 for detailed statistics summary.

Average neural activity from the fiber photometry recordings in LHb was plotted in the *t*-SNE space (Figure 3D, results split across cohorts in: Figure S3C; see Methods). Activity was greatest for clusters that corresponded to high proximity between the mice (Figure 3E-F), consistent with the aversive nature of being near the aggressor.

To better interpret these proximal clusters, we trained random forest classifiers^27^ to identify four behaviors - being investigated, being attacked, fighting back, and fleeing - across all video frames using the same 12 features as were embedded into *t*-SNE space (Figure 3B; Figure S3D-F). These four behaviors together spanned the portion of the *t*-SNE map where LHb activity was the highest (compare Figure 3G-J for random forest densities with Figure 3D for neural data). The onset of these four behaviors had similar responses (Figure 3K-N).

Taken together, this implies that LHb activity is elevated across proximal behaviors during defeat, with little differentiation across such behaviors. This lack of behavioral differentiation contrasts with our observations in VTA dopamine neurons, as in that case we saw different response patterns in relation to different behaviors (e.g. flee versus fight back)^27^.

### From the first defeat session, LHb activity is higher in susceptible mice when attacked

Thus far, we observed differences in LHb activity in susceptible and resilient mice following but not preceding CSDS (Figure 2), as well as elevated activity during proximal behaviors during defeat, when considering all mice (susceptible or resilient; Figure 3). We next asked whether neural activity in LHb is different between susceptible and resilient mice during defeat, and if so, when differences first emerged.

Susceptible mice had higher activity than resilient mice in the portion of the *t*-SNE space corresponding to proximal behaviors such as being attacked, fighting, or fleeing (Figure 4A-C; compare to random forest densities in Figure 3G-J; consistent pattern across 2 cohorts: Figure S4A; activity timelocked to each t-SNE cluster: Figure S5-6). In contrast, resilient mice had higher activity in the portion of the *t*-SNE map corresponding to a vigilance-like posture (Figure 4C; close to a wall, low body posture, and oriented toward the aggressor: Figure S4B-G). These conclusions were also evident from directly time-locking activity to the random-forest identified behaviors (Figure 4D-G).

**Figure 4.**
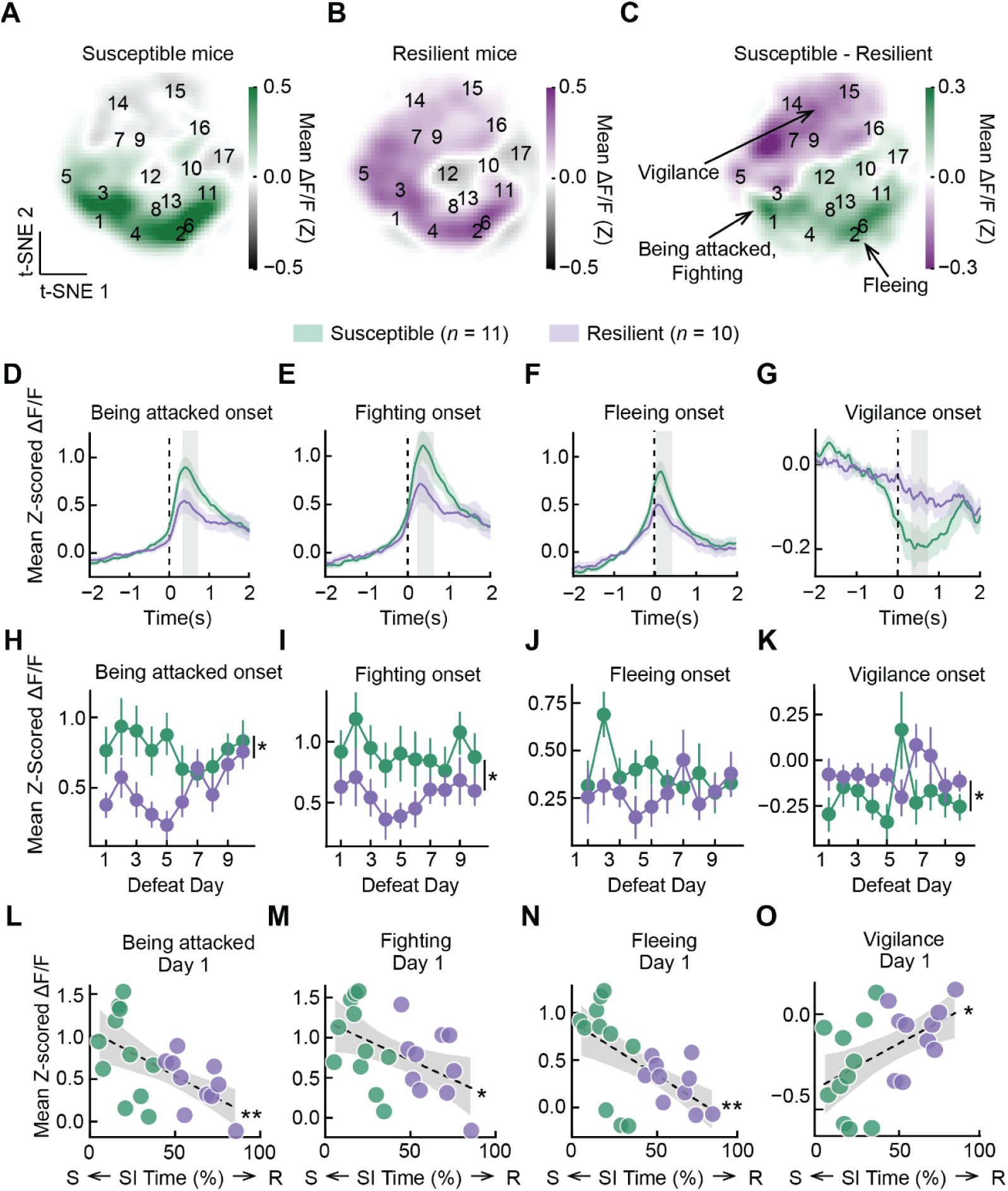
From the 1st day of defeat, higher LHb activity in susceptible mice during proximal behaviors and in resilient mice when vigilant. **A**. Mean LHb GCaMP dF/F across *t*-SNE behavior space in susceptible mice (across all 10 days of defeat; *N* = 11). **B**. Mean LHb GCaMP dF/F across *t*-SNE behavior space in resilient mice (across all 10 days of defeat; *N* = 10). **C**. Difference between susceptible and resilient LHb GCaMP dF/F (difference between panel **A** and **B**). **D**. Being attacked onset-aligned LHb responses during defeat averaged across individuals in resilient and susceptible groups (mean ± s.e.m. plotted). Grey region indicates +/-0.25 s surrounding the peak response. **E**. Fighting back onset-aligned LHb dF/F during defeat. Grey region indicates +/-0.25 s surrounding the peak response. **F**. Fleeing onset-aligned LHb dF/F during defeat. Grey region indicates +/-0.25 s surrounding the peak response. **G**. Vigilance onset-aligned LHb DF/F during defeat. Grey region indicates +/-0.25 s surrounding the minimum response. **H**. Average LHb GCaMP dF/F to attack onset from susceptible and resilient groups across defeat (mean ± s.e.m across mice; averaging across labeled gray region (+/-0.25 s peak response) in **D**). Onset activity by SI time, day, and their interaction: main effect of SI time, *Z* = −2.428, *p* = 0.015; main effect of day, *Z* = 0.537, *p* = 0.591. Interaction, *Z* = 2.088, *p* = 0.037. **I**. Average LHb GCaMP dF/F to fighting onset from susceptible and resilient groups across defeat (mean ± s.e.m across mice; averaging across labeled gray region (+/-0.25 s peak response) in **E**). Onset activity by SI time, day, and their interaction: main effect of SI time, *Z* = −2.089, *p* = 0.037; main effect of day, *Z* = −0.431, *p* = 0.666. Interaction, *Z* = 0.808, *p* = 0.491. **J**. Average LHb GCaMP dF/F to fleeing onset from susceptible and resilient groups across defeat (mean ± s.e.m across mice; averaging across labeled gray region (+/-0.25 s peak response) in **F**). Onset activity by SI time, day, and their interaction: main effect of SI time, *Z* = −1.607, *p* = 0.108; main effect of day, *Z* = −0.539, *p* = 0.590. Interaction, *Z* = 2.520, *p* = 0.012. **K.** Average LHb GCaMP dF/F to vigilance onset from susceptible and resilient groups across defeat (mean ± s.e.m across mice; averaging across labeled gray region (+/-0.2s minimum response) in **G**). Onset activity by SI time, day, and their interaction: main effect of SI time, *Z* = 2.185, *p* = 0.019; main effect of day, *Z* = 0.188, *p* = 0.851. Interaction, *Z* = 0.207, *p* = 0.836. **L**. Average LHb GCaMP responses to attack onset (labeled gray region in **D**) on day 1 plotted against SI time for each mouse (*N* = 21 mice): *R* = −0.5783, *p* = 0.0060. **M**. Average LHb GCaMP responses to fighting onset (labeled gray region in **E**) on day 1 plotted against SI time for each mouse (*N* = 21 mice): *R* = −0.4806, *p* = 0.0274. **N**. Average LHb GCaMP responses to fleeing onset (labeled gray region in **F**) on day 1 plotted against SI time for each mouse (*N* = 21 mice): *R* = −0.5807, *p* = 0.0057. **O**. Average LHb GCaMP responses to vigilance onset (labeled gray region in **G**) on day 1 plotted against SI time for each mouse (*N* = 21 mice): *R* = 0.4954, *p* = 0.0224. *p*-values in **H**-**K** are from two-sided GEE. *p*-values in **L**-**O** are from Pearson’s correlations. Shaded areas in **L**-**O** represent 95% confidence interval for linear fit. \**p* ≤ 0.05, \*\**p* ≤ 0.01, \*\*\**p* ≤ 0.001. See Supplementary Table 1 for detailed statistics summary. See Supplementary Tables 10-17 for more information on GEE statistics.

Rather than emerging gradually, these differences between susceptible and resilient mice were present from the first day of CSDS (Figure 4H-O; see Figure S4H for summary of response to each attack across day 1; analogous results from *t*-SNE in Figure S4I). These differences imply that heightened initial LHb responses to the stressor might produce susceptibility. This also provides a clear contrast to our prior observations in the VTA dopamine system, where we observed differences in neural correlates in susceptible and resilient mice emerge gradually over the course of CSDS^27^.

### Closed-loop activation of the LHb during defeat biases towards susceptibility

To determine if the elevated activity in the LHb observed in susceptible mice during defeat causes susceptibility, we performed closed-loop optogenetic activation during defeat. vGlut2-Cre males were bilaterally injected in the LHb with either a Cre-dependent excitatory opsin (ChR2 or ChRmine) or control virus (YFP), and optical fibers were implanted in the LHb (Figure 5A, validation of stimulation parameters: Figure S7A-D; histology: Figure S7E).

**Figure 5.**
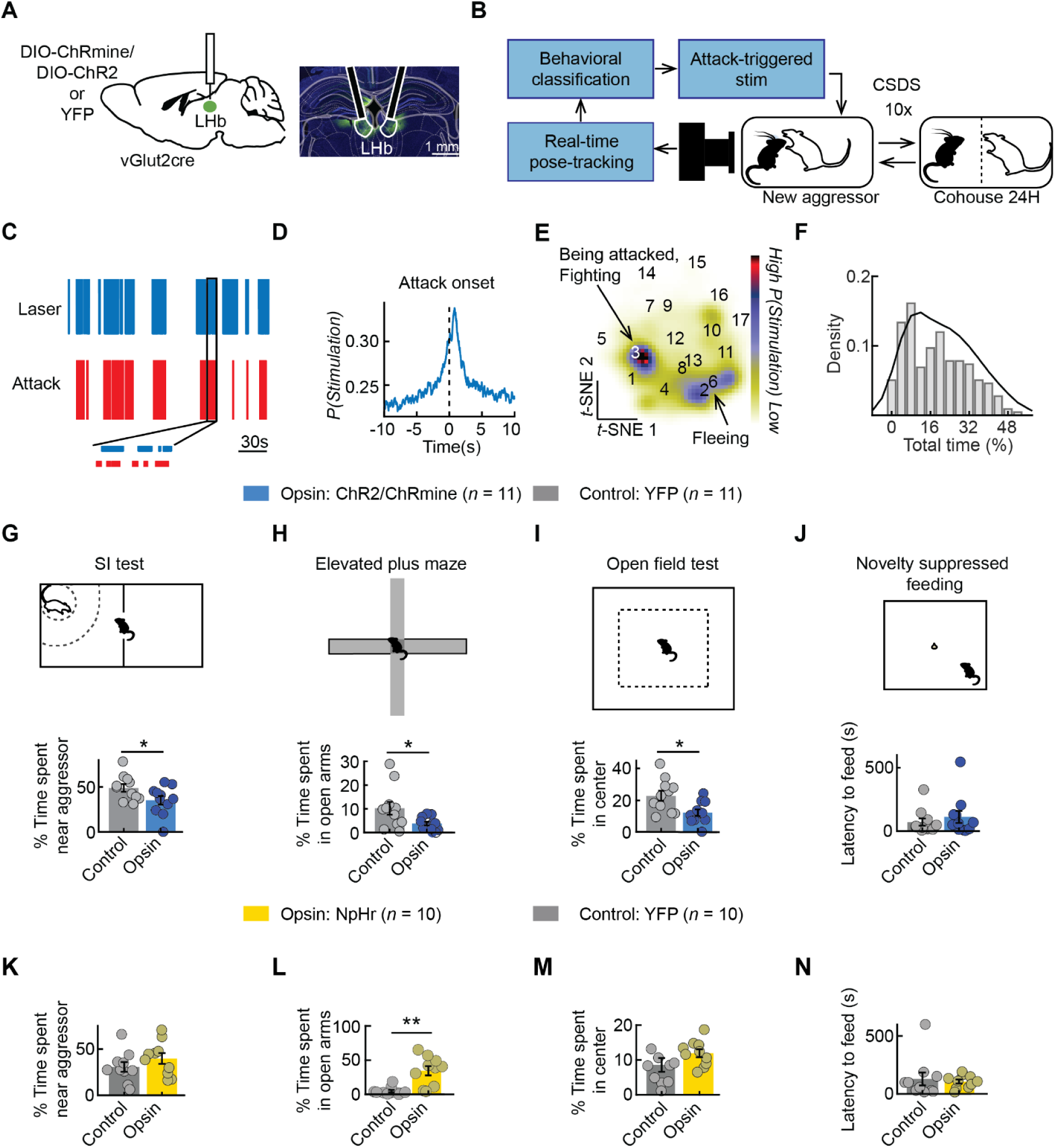
Closed-loop activation of LHb during CSDS produces susceptibility. **A**. Left: Location of virus injections and fiber targeting of cell bodies in the Lateral Habenula (LHb). Right: Example histology of virus expression. **B**. Schematic of attack-triggered stimulation. Each detected attack frame during defeat triggered 5 pulses of 20 Hz activation. **C**. Example of a defeat session with laser light delivery triggered on attack of the closed-loop mouse. Bottom inset is a 10s segment. **D**. Probability of a laser train, as a function of time relative to attack onset. **E**. Density of activation in *t*-SNE space. **F**. Distribution across sessions of percent of defeat session that mice received laser (mean is 21.34% of the defeat session). **G**. Difference in SI time between opsin (ChR2 or ChRmine) and control group. Opsin (ChR2 or ChRmine) vs control: *t* = −2.1263, *p* = 0.0461. **H**. Difference in open-arm time in the elevated plus maze between opsin (ChR2 or ChRmine) and control group. Opsin (ChR2 or ChRmine) vs control (YFP): *t* = −2.2934, *p* = 0.0328. **I**. Difference in center time in the open field between opsin (ChR2 or ChRmine) and control (YFP) group. Opsin (ChR2 or ChRmine) vs control (YFP): *t* = −2.8231, *p* = 0.0105. **J**. Difference in latency to feed in novelty suppressed feeding assay between opsin (ChR2 or ChRmine) and control (YFP) group. Opsin (ChR2 or ChRmine) vs control (YFP): *t* = −0.7336, *p* = 0.4717. **K**. Difference in SI time between opsin (NpHr) and control group. Opsin (NpHr) vs control: *t* = −1.1623, *p* = 0.2603. **L**. Difference in open-arm time in the elevated plus maze between opsin (NpHr) and control (YFP) group. Opsin (NpHr) vs control (YFP): *t* = −3.9140, *p* = 0.0010. **M**. Difference in center time in the open field between opsin (NpHr) and control (YFP) group. Opsin (NpHr) vs control (YFP): *t* = −1.4660, *p* = 0.1599. **N**. Difference in latency to feed in novelty suppressed feeding assay between opsin (NpHr) and control group. Opsin (NpHr) vs control *t* = −0.3913, *p* = 0.7002. *p*-values in **G**-**N** are from unpaired 2-sided *t*-tests. \**p* ≤ 0.05, \*\**p* ≤ 0.01, \*\*\**p* ≤ 0.001. See Supplementary Table 1 for detailed statistics summary.

In order to recapitulate the heightened LHb activity during attack, fighting, and fleeing observed in susceptible mice (Figure 4D-F), we streamed video frames to our pose-estimation network, calculated the 12 features as previously described (Figure 3B), and inputted them into the random forest classifier to identify attack and trigger laser activation^27^ (Figure 5B). Post-hoc analyses confirmed that this resulted in the greatest activation during attack (Figure 5C-E), as well as activation during fighting and fleeing which closely follow attack (Figure 5E, compare to susceptible mice LHb activity map in Figure 4A). Across the 10 days of CSDS, the average duration of this activation was ∼1.07 min/day (21.4% of session; Figure 5F).

Activation of the LHb increased freezing behavior during defeat (Figure S7F), and biased mice towards a susceptible phenotype. Specifically, mice that received activation were less social on the SI test (Figure 5G), spent less time in the open arms in the elevated plus maze (Figure 5H), and less time in the center of the open field (Figure 5I), although they did not have a significantly decreased latency to feed in a novel context (Figure 5J). These differences after chronic stress were not due to significant differences in being attacked across the groups (Figure S7F).

We next performed an analogous closed-loop optogenetic inhibition experiment during defeat to determine if inhibition of LHb during attack causes resilience. We injected vGlut2-Cre males bilaterally in the LHb with either a Cre-dependent inhibitory opsin (NpHr) or control virus (YFP), and implanted optical fibers in the LHb (Figure S7E). Inhibiting LHb during attack was not sufficient to create a robust resilient phenotype in most behavioral measures, but there was a trend towards resilience in some assays, and the effect in the elevated plus maze was highly significant (Figure 5K-N). Similar to our stimulation experiment, we did not observe a significant difference in time attacked across the groups (Figure S7G).

### Activation of the LHb during defeat produces durable, brainwide changes

Stimulation of the LHb during defeat increased susceptibility and anxiety-like behavior, a change that persisted for days after CSDS ended (Figure 5). This result raises the question of how this manipulation may alter the brain’s response to later encounters with an aggressor. To address this, we performed high resolution and high signal-to-noise measurements of brainwide activity in mice that had received attack-triggered LHb activation (*N* = 10, Figure 6A) during defeat, or mice who also underwent defeat but did not receive LHb stimulation (*N* = 44).

**Figure 6.**
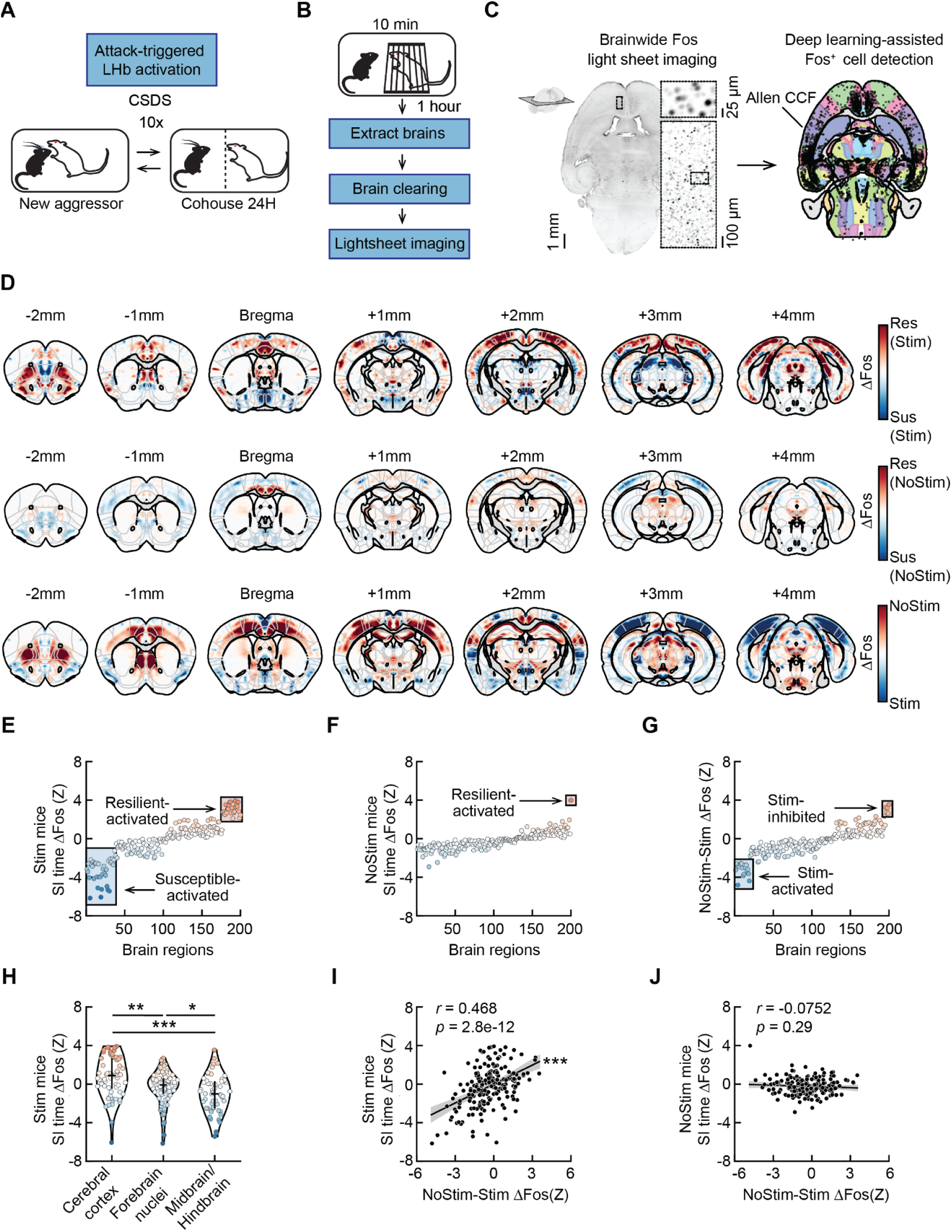
Activation of the LHb during defeat produces durable, brainwide changes. **A**. Schematic of closed-loop attack-triggered LHb stimulation during CSDS. The Fos dataset includes 10 mice that received LHb stimulation during CSDS and 44 unstimulated control mice. **B**. Approximately one week after the conclusion of CSDS, mice were placed for 10 min into the cage of a novel Swiss Webster aggressor that was restrained under a wire cup. There was no LHb stimulation during this assay. The mice were then sacrificed one hour later for Fos analysis. **C**. Left: example brainwide Fos imaging data. A 50-plane (100-µm) maximum intensity projection is shown. The insets are shown after background subtraction and filtering. Right: All detected cells overlaid on the Allen CCF for the example section on the left. **D**. The difference in Fos^+^ cell density across LHb-stimulated resilient (*N* = 4 mice) and susceptible (*N* = 6 mice) mice (top row), across unstimulated resilient (*N* = 30 mice) and susceptible (*N* = 14 mice) mice (middle row), and across all LHb-stimulated (*N* = 10 mice) and all unstimulated (*N* = 44 mice) mice (bottom row). **E**. Individual brain regions sorted by the estimated contribution of SI time to Fos^+^ cell counts based on GLM coefficients for mice that received LHb stimulation (*N* = 10 mice). Regions that were significantly different are highlighted with red (resilient-activated) and blue (susceptible-activated) boxes. **F**. Individual brain regions sorted by the estimated contribution of SI time to Fos^+^ cell counts based on GLMM coefficients for unstimulated control mice (*N* = 44 mice). **G**. Individual Allen CCF brain regions sorted by the estimated contribution of LHb stimulation to Fos^+^ cell counts based on GLMM coefficients across all mice (*N* = 54 mice). **H**. Comparison of distributions of LHb stimulation coefficients (from **G**) across all brain regions in cerebral cortex (*n* = 61 regions), forebrain nuclei (*n* = 83 regions), and midbrain/hindbrain (*n* = 56 regions). **I**. Correlation between the estimated contribution to Fos^+^ cell counts of SI time in LHb-stimulated mice (from **E**; *y*-axis) vs. of LHb stimulation across all mice (from **G**; *x*-axis) (*n* = 200 regions). **J**. Correlation between the estimated contribution to Fos^+^ cell counts of SI time in unstimulated control mice (from **F**; *y*-axis) vs. of LHb stimulation across all mice (from **G**; *x*-axis) (*n* = 200 regions). Significance in **E**-**G** is based on GLM or GLMM coefficient estimate *z*-tests corrected for 10% false discovery rate. Error bars in **H** represent median ± interquartile range. Shaded areas in **I**-**J** represent 95% confidence interval for linear fit. *p*-values in **H** are from Kolmogorov-Smirnov tests with Hochberg-Bonferroni correction for multiple comparisons. *p*-values in **I**-**J** are from Pearson correlations. \**p* ≤ 0.05, \*\**p* ≤ 0.01, \*\*\**p* ≤ 0.001. See Supplementary Tables 1 and 60–62 for detailed statistics summary.

Approximately one week after the last day of CSDS (after all the post-CSDS tests), each mouse was introduced for 10 minutes to the cage of a novel aggressor restrained under a mesh cup and was sacrificed one hour later (Figure 6B). We next cleared the brains with iDISCO+, stained for the immediate early gene Fos as a marker of neural activation, and imaged with a SPIM light sheet fluorescence microscope (Figure 6C)^56,57^. We then used an automated deep learning-assisted cell detection pipeline^58^ to generate cellular resolution maps of brainwide neural activation registered to the Allen common coordinate framework (CCF)^59^ for each animal (Figure 6C-D; see Methods). We detected a total of 19,337,269 Fos^+^ cells across all mice.

We first analyzed how the brainwide response to the aggressor differed across resilient and susceptible mice that had received LHb stimulation during CSDS. We used a generalized linear mixed model (GLMM) to estimate the contribution of post-CSDS SI time, as a proxy of susceptibility versus resilience, to neural activation (Fos^+^ cell counts) for each brain region (see Methods). This revealed that activation of a surprisingly large fraction of regions was significantly modulated by susceptibility versus resilience among LHb-stimulated mice (approximately 30%, Figure 6E; Supplementary Data Table 60; example resilient-activated regions: anterior cingulate cortex, medial entorhinal area, and piriform area, *p* < 0.01 for all; example susceptible-activated regions: subiculum, lateral amygdala, and pontine central gray, *p* < 0.05 for all). Conversely, mice that did not receive LHb stimulation were very weakly modulated by susceptibility versus resilience (Figure 6F, Supplementary Data Table 61; only the medial habenula was significantly activated in resilient mice, *p* < 0.0001).

Next, we tested how the brainwide response to the aggressor differed across mice that received LHb stimulation during defeat versus those that did not receive stimulation. We first fit a GLMM that estimated the contribution of LHb stimulation alone to neural activation. In this analysis, LHb stimulation significantly impacted the aggressor response of several regions (12%, Figure 6G; Supplementary Data Table 62; example LHb stimulation-activated regions: the parabrachial nucleus, central amygdala, and medial geniculate nucleus, *p* < 0.01 for all; example LHb stimulation-inhibited regions: primary and secondary motor areas and primary somatosensory area, *p* < 0.01 for all). When we considered the combined effects of LHb stimulation during CSDS and of post-CSDS SI time (again, as a proxy for susceptibility) using a GLMM that included both terms and their interaction, we found that many regions encoded the *interaction* between LHb stimulation and SI time, but not the main effects of SI time (i.e., in the unstimulated mice) or of LHb stimulation (Figure S8A-C, Supplementary Data Table 63-65). This is consistent with our finding above that susceptibility in LHb-stimulated mice involves strong brainwide responses upon subsequent exposure to the aggressor (Figure 6E), whereas susceptibility in unstimulated control mice involves weaker brainwide responses (Figure 6F).

Interestingly, we found that the susceptible mice (i.e., low SI time) that received LHb stimulation had strong engagement of a broad subcortical network (Figure 6H, Figure S8D). For example, in resilient LHb-stimulated mice there was more activity in the anterior cingulate cortex (dorsal and ventral; *p* < 0.001), motor cortex (secondary motor area; *p* < 0.01) and sensory cortices (anteromedial visual area, primary somatosensory area barrel field, piriform cortex, among others; *p* < 0.05, see Supplementary Data Table 60 for details). In contrast, in susceptible LHb-stimulated mice there was more activity in the parabrachial nucleus, pedunculopontine nucleus, substantia nigra pars reticulata, medial habenula, and central and lateral amygdala (*p* < 0.01 for all; see Supplementary Data Table 60 for details).

To determine if there was a relationship between susceptibility following LHb stimulation and the overall effect of LHb stimulation (versus controls), we examined the pairwise correlation of SI time coding in the LHb-stimulated mice (from Figure 6E) to the coding of LHb stimulation versus control across all mice (from Figure 6G). This revealed a strong correlation (Figure 6I), which was absent when we performed an analogous analysis for the unstimulated mice (Figure 6J). The significant correlation between susceptibility (in LHb-stimulated mice) and LHb stimulation (vs. control) suggests that the brainwide encoding of susceptibility is similar to the encoding of stimulation.

To complement the GLMM analyses above, we also performed a clustering analysis of the animal-by-animal pairwise correlation in Fos counts across all brain regions in the LHb-stimulated mice (Figure S8E). We found that activation of the the LHb cluster (red) tended to be anti-correlated with the cluster containing the dorsal raphe nucleus (DRN; yellow) and positively correlated with the cluster containing the VTA (dark blue), potentially consistent with a recent study that showed that the strength of the LHb projection to VTA but not DRN increases after stress^21^. In addition, we find some shared network structure between the LHb stimulated and unstimulated mice, particularly in the LHb cluster.

Taken together, these analyses suggest that heightened LHb activity during defeat leads to a strong difference in how the brains of resilient and susceptible animals respond to subsequent encounters with an aggressor. This differential response is characterized by the recruitment of broad subcortical vs cortical networks in susceptible vs resilient animals, that persists for many days following the end of LHb stimulation.

## Discussion

While prior work has uncovered differences between susceptible and resilient mice after CSDS, much less is known about the role of neural teaching signals during stress in driving differences in stress outcomes. Here, we focus on the LHb, which provides a negative teaching signal^33,60–62^ and is implicated in aversive learning^34,35^ and depression-related behavior^21,35,38–40,55,63,64^, to ask: 1) when and how does activity in the LHb first differ between susceptible and resilient individuals? and 2) do these differences produce behavioral and brainwide correlates of susceptibility?

We found heightened LHb activity during proximal behaviors during and after social defeat (but not before), with little dependence on the specific proximal behavior (Figure 2-3). From the first day of defeat, this elevated activity is stronger in susceptible than resilient mice (Figure 4). LHb stimulation during defeat is sufficient to produce a susceptible phenotype (Figure 5), as well as to generate a persistent shift in the balance of subcortical versus cortical activity in susceptible mice (Figure 6).

### Learning as a result of the CSDS paradigm

Though CSDS is a widely used model of chronic stress^4–14^, what animals learn as a result of the paradigm, and what signals drive this learning, remain open questions. We think our data provides evidence of specificity, as well as generalization, in terms of what mice learn. The evidence for learning specificity comes from the social avoidance tests, where mice learn to avoid the aggressor strain (although note that they do generalize across mice of that strain), but do not avoid other strains (displaying learning specificity; Figure 1). There is also evidence of a susceptible phenotype that generalizes beyond the social context, as we (and others^63,65–68^) observe a correlation between susceptibility based on social interaction (SI) time and anxiety-like behavior in the elevated plus maze, open field test, novelty suppressed feeding, as well as immobility in a neutral context (Figure 1). Note that another recent paper that used longer defeat sessions found that social aversion generalized across strains^53^, suggesting that more intense stress results in more generalization.

Regarding what signals drive this learning, our data is consistent with a model that LHb serves as an aversive teaching signal during CSDS, similar to what has been shown in other settings^33–37,60–62^. In particular, we found that: 1) LHb activity was stronger from the first social interactions in the mice that learned more social aversion (i.e. susceptible mice); 2) this difference significantly decreased across days, once the social stress experience became less unexpected; 3) closed-loop LHb activation during stress caused mice to be more susceptible to the stress.

### LHb activation and susceptibility recruits a subcortical network

Activation of the LHb during defeat produces sustained, brainwide differences in response to being near the aggressor in susceptible versus resilient mice. In particular, susceptible mice that received stimulation had greater activation of subcortical regions, while mice that did not receive stimulation had greater activation in cortical regions (Figure 6). Surprisingly, this organization of cortical activation in resilience and subcortical activation in susceptibility was not evident in mice that did not receive stimulation, suggesting that LHb stimulation increases the brainwide encoding of susceptibility vs resilience. Note that this brainwide pattern is consistent with the fact that several cortical regions have been implicated in resilience^69–74^ and several subcortical regions have been implicated in susceptibility^24,75–81^. An interesting open question is why the encoding of susceptibility was so much stronger in mice that received stimulation.

### Separate roles for LHb and VTA dopamine during defeat in the progression to susceptibility versus resilience

LHb neurons inhibit VTA dopamine neurons^36,82–87^, and the two populations are thought to have roughly opposite response profiles and functions^33,36^. Consistent with these opposing roles, activation of NAc-projecting DA neurons during defeat biases towards resilience^27^, whereas here we show activation of LHb neurons during defeat biases towards susceptibility (Figure 5). Given this, LHb activity during defeat may contribute to susceptibility at least in part by inhibiting pro-resilient VTA DA activity^44^.

However, a comparison of the current findings and our previous work^27^ points to important differences in the correlates and consequences of activity in LHb versus VTA dopamine neurons during defeat. First, the LHb shows little action selectivity during proximal behaviors during defeat (Figure 3), while NAc-projecting dopamine neurons display clear selectivity to specific actions (e.g. fighting back versus being attacked). Second, differences in activity between susceptible and resilient mice are present from the first day of defeat in LHb (Figure 4), while differences only emerge slowly during defeat in dopamine neurons. Finally, attack-triggered activation of LHb produces susceptibility (Figure 5), while attack-triggered dopamine inhibition does not.

These differences between LHb versus VTA dopamine neurons imply that dopamine neurons are not simply a reflection of LHb activity during defeat, and that LHb-mediated and dopamine-mediated mechanisms of susceptibility versus resilience are at least partially distinct. Since the LHb also sends a major projection to the raphe^88–94^, that projection is a good candidate to mediate the effects of LHb activity on susceptibility^55,89,91,95^.

One possibility is that LHb activity during defeat primarily controls the progression towards susceptibility (with less control of resilience)^38,40,96^, while dopamine activity during defeat primarily controls the progression towards resilience^27,97^ (with less control of susceptibility). This is consistent with our inability to produce susceptibility with manipulations of DA neurons during defeat^27^ as well as our inability to produce a strong resilient phenotype by inhibiting LHb (Figure 5), and the broader idea that resilience and susceptibility are distinct and actively learned processes^5,27,98^.

### Relationship to recent work on the LHb and stress

Our results align with previous work investigating the role of LHb activity both during and after stress. Similar to previous studies, we have also shown LHb is dysregulated after stress in susceptible individuals^38,41–46^. Our results complement recent work by Fan et al., 2023 which showed that LHb activity is heightened in mice at the top of the dominance hierarchy during unexpected forced loss in the tube test, and that those mice show greater depression-like behavior after the loss (based on measures of anhedonia and immobility). Our work adds to this work as we 1) leveraged a different stress paradigm and a different panel of post-stress assays, demonstrating generalizability of the importance of elevated LHb activity to the case of susceptibility to chronic social stress, 2) performed automatic social behavioral quantification to demonstrate that social proximity, rather than the specific behavior (e.g. fighting, fleeing), was most important for elevation of LHb activity during social stress, 3) performed social-behavior triggered optogenetic manipulations of LHb activity, 4) identified alterations in brainwide activation in susceptible mice days after stimulation of the LHb during stress, 5) demonstrated that stress-activated (but not stress-inhibited) LHb neurons change their activity in response to stress.

## STAR★Methods

### Key resources table

| REAGENT or RESOURCE | SOURCE | IDENTIFIER |
| --- | --- | --- |
| Antibodies |  |  |
| Rabbit anti-GFP | Novus Biologicals | CAT# NB600-308; RRID: AB_341929 |
| Rabbit anti-Fos | Synaptic Systems | CAT# 226008; RRID: AB_2891278 |
| Chicken anti-GFP | Aves | CAT# GFP-1020; RRID: AB_10000240 |
| Donkey anti-Rabbit Alexa 647 | ThermoFisher Scientific | CAT# A-31573; RRID: AB_2536183 |
| Donkey anti-Rabbit Alexa 647 | Abcam | CAT# ab150075; RRID:AB_2752244 |
| Donkey anti-Chicken Alexa 594 | Jackson Immuno | CAT# 703-585-155; RRID:AB_2340377 |
| Bacterial and virus strains |  |  |
| AAV5-CaMKII-GCaMP6f-WPRE-SV40 | Addgene | Cat# 100834-AAV5; RRID:Addgene_100834 |
| AAV5-syn-jGCaMP7f-WPRE-SV40pA | PNI vector core | N/A |
| AAV9-syn-FLEX-GCamp6s-WPRE | Addgene | CAT# 100845-AAV9; RRID:Addgene_100845 |
| AAV5-EF1a-DIO-ChRmine-EYFP/mScarlet-WPRE | PNI Virus core | N/A |
| AAV5-EF1a-DIO-hChR2(H134R)-EYFP-WPRE-HGHpA | PNI vector core | N/A |
| AAV5-EF1a-DIO-EYFP-WPRE-hGHpA | PNI vector core | N/A |
| AAV9-EF1a-DIO-eNpHR3.0-EYFP-WPRE-hGH | Addgene | CAT# 26966-AAV9; RRID:Addgene_26966 |
| Experimental models: Organisms/strains |  |  |
| Mouse: wild type C57BL/6J | The Jackson Laboratory | Stock# 000664;<br>RRID:IMSR_JAX:000664 |
| Mouse: wild type AKR/J | The Jackson Laboratory | Stock# 000648;<br>RRID:IMSR_JAX:000648 |
| Mouse: Slc17a6tm2(cre)Lowl/J | The Jackson Laboratory | Stock# 016963;<br>RRID:IMSR_JAX:016963 |
| Mouse: wild type SW | Taconic Biosciences | Stock# Tac:SW;<br>RRID:IMSR_TAC:SW |
| Software and algorithms |  |  |
| BORIS | Friard and Gamba, 2016 | http://www.boris.unito.it |
| CSDS analysis code | Willmore et al., 2022 | Database:<br>https://github.com/lwillmore/QuantifyingDefeat |
| Custom analysis code | This paper | Database:<br>https://github.com/annazhuk/CSDS_LHb |
| DeepLabCut | Mathis et al., 2018 | Database:<br>https://github.com/DeepLabCut/DeepLabCut |
| EthoVision | Noldus | https://www.noldus.com/ethovision-xt |
| Fiji<sup>99</sup> | Schindelin et al., 2012 | https://imagej.net/Fiji |
| IDPS | Inscopix | https://inscopix.com/ |
| MATLAB | The MathWorks, Inc | https://www.mathworks.com/products/matlab.html |
| Motif | LoopBio | http://loopbio.com/recording/ |
| Python | Python Software Foundation | http://www.python.org/ |
| Other |  |  |
| Patch cables for fiber photometry | Doric Lenses | MFP_400/430/1100-0.48_2m_FCM-MF2.5 |
| 0.5 mm diameter, ~6.1 mm length GRIN lens | Inscopix | CAT# 1050-004610 |
| Imaging Baseplate | Inscopix | CAT# 1050-004638 |
| Baseplate cover | Inscopix | CAT# 1050-004639 |
| Cable sheath | Inscopix | CAT# 1050-003523 |

### Resource availability

#### Lead contact

Further information and requests for resources should be directed to the lead contact, Ilana Witten.

#### Materials availability

Plasmids and viruses generated in this study are available by contacting the lead contact.

#### Data and code availability

Code used in this paper is available at: https://github.com/annazhuk/CSDS_LHb/. Data reported in this paper is available at: https://figshare.com/articles/dataset/Heightened_lateral_habenula_activity_during_stress_produ ces_brainwide_and_behavioral_substrates_of_susceptibility/26072956.

### Experimental model and study participant details

#### Mice

All experiments were approved by the Princeton University Institutional Animal Care and Use Committee and were in accordance with National Institutes of Health standards. Prior to and throughout experimental assays, experimental and stimulus animals were housed under a 12 H light-dark cycle with experiments exclusively taking place during the dark phase. Mice used in this study were C57BL/6J males (RRID:IMSR_JAX:000664) between the ages of 8 and 24 weeks old, Swiss Webster males (RRID:IMSR_TAC:SW) between the ages of 8 and 57 weeks, and AKR/J males (RRID:IMSR_JAX:000648) between the ages of 8 and 16 weeks old. A total of 150 mice were used for recordings and manipulation: 57 for fiber photometry, 11 for cellular resolution calcium imaging, 40 for fos experiments, and 42 for optogenetics experiments. An additional 161 mice were used as stimulus mice (social targets or aggressors): 110 Swiss Websters, 33 C57BL6/J, and 18 AKR/J. Mice undergoing fiber photometry experiments were wild-type. Mice undergoing cellular resolution calcium imaging experiments and mice undergoing optogenetic experiments were heterozygous vGlut2-cre (RRID:IMSR_JAX:016963). Food and water were given *ad libitum*.

### Method details

#### Surgery

At 4-12 weeks of age, animals were anesthetized (isoflurane at 5% for induction and 1-2% for maintenance) and leveled with a stereotaxic frame before injections and implants were done.

For fiber photometry recordings, mice were injected with 80 nL of viral vector expressing a GCaMP (cohort 1: AAV5-CaMKII-GCaMP6f-WPRE-SV40 at a titre of 3.13E13 genome copies/mL produced by addgene or cohort 2: AAV5-syn-jGCaMP7f-WPRE-SV40pA at a titre of 2.5E14 genome copies/mL produced by PNI viral core) in the LHb (AP −1.6 mm, ML +/− 0.46 mm, DV −3 mm relative to the skull surface at Bregma) and implanted with 400 μm core diameter optical fibers (MFC_400/430-0.48_4mm_MF2.5_FLT from Doric Lenses Inc.) in the LHb (AP −1.6 mm, ML +/− 0.46 mm, DV −2.4 mm relative to the skull surface at Bregma), with hemisphere selection counterbalanced between animals. Metabond (Parkell) was used to fix fibers to the skull. Ortho-Jet Crystal mixed with carbon glassy, spherical powder (Sigma-Aldrich) was then used to further secure the implants to the metabond. Mice were allowed to recover for at least a week before starting CSDS. Mice used in fiber photometry experiments were given 4 weeks of recovery time following surgery before experiments began.

For cellular resolution calcium imaging experiments, mice were injected with 80 nL of viral vector expressing a GCaMP (AAV9-syn-FLEX-GCamp6s-WPRE at a titre of 2.13E13 genome copies/mL produced by addgene). At least 5 days later, animals were implanted with a 0.5 mm diameter GRIN lens (1050-004610, Inscopix) in the LHb (AP −1.6 mm, ML +/− 0.47 mm, DV −2.45 mm relative to the skull surface at Bregma). At least 4 weeks later, a baseplate (1050-004638, Inscopix), attached to the miniature microscope (nVISTA 3.0, Inscopix), was positioned over the GRIN lens such that the neurons were in focus. The baseplate along with a titanium headplate^100^ were then secured to the skull using Ortho-Jet Crystal mixed with carbon glassy, spherical powder (Sigma-Aldrich), and a baseplate cover (1050-004639, Inscopix) was used to protect the GRIN lens. Mice used in cellular resolution experiments were given 2-4 weeks of recovery time following base plate implants before experiments began.

For optogenetic experiments, vGlut2-cre animals were injected with 60 nL of viral vector expressing ChR2 (AAV5-EF1a-DIO-ChR2-eYFP) at a titre of 2.40E+13 or 80 nL of viral vector expressing ChRmine (AAV5-EF1a-DIO-ChRmine-EYFP/mScarlet-WPRE-HGHpA at a titre of 1.8E13 genome copies/mLproduced by the PNI virus core) or 60 nL of viral vector expressing NpHr (AAV9-EF1a-DIO-eNpHR3.0-EYFP-WPRE-hGH at a titre of 5.6E+13 genome copies/mL) or YFP (AAV5-EF1a-DIO-EYFP-WPRE-hGHpA at a titre of 1.5-2E14 genome copies/mL produced by the PNI virus core) in each LHb (AP −1.6 mm, ML +/− 0.46 mm, DV −3 mm relative to the skull surface at Bregma) and implanted bilaterally with a 200 μm core diameter optical fibers above the LHb (AP −1.6 mm, ML +/− 0.87 mm, DV −2.28 mm relative to the skull surface at Bregma, inserted at a 10° angle). Mice were allowed to express virus for a minimum of 4 weeks before behavioral experiments were initiated. Mice used in optogenetic experiments were given 4 weeks of recovery time following surgery before experiments began.

#### Fiber photometry data acquisition

GCaMP fiber photometry recordings were collected through two different systems. Data from cohort 1 was collected using a fiber photometry set-up similar to that described in Gunaydin et al., 2014. A 488 nm laser light (Micron Technology) was filtered (FL488, Thor Labs), passed through a dichroic mirror (MD498, Thor Labs), and delivered through a patch cable (MFP_400/430/1100-0.48_2m_FCM-MF2.5, Doric Lenses) which was coupled to the fiber attached to the mouse via a ceramic split sleeve (2.5 mm diameter, Precision Fiber Products). The laser, which was modulated at 210.999 Hz, was controlled by a lock-in amplifier (Ametek, 7265 Dual Phase DSP Lock-in Amplifier). Fluorescent emission from GCaMP6f at 500-550 nm then passed through the same patch cable and dichroic mirror into a photodetector (Model 2151, New Focus), and the signal was filtered at the same 210.999 Hz using the same lock-in amplifier, and a time constant of 20 ms. AC gain on the lock-in amplifier was set to 0 dB. The signal was digitized at 1000 Hz.

Data from cohort 2 was collected using a set-up similar to that described in Willmore et al., 2022. We used a Doric Lenses photometry system (4-channel driver LEDD_r, LEDs at 465 (and 405 nm in a subset of animals), fluorescence mini cube FMC5_E1(465-480)_F1(500-540)_E2(555-570)_F2(580-680)_S, and Newport Visible Femtowatt Photoreceiver Module NPM_2151_FOA_FC). The system was driven by and recorded from using custom code written for a real-time processor (RZ5P, Tucker Davis Technologies) in OpenWorkBench (v.2.28.0). GCaMP was excited by driving a 465 nm light-emitting diode (LED) light (about 400 Hz sinusoidal modulation, at an intensity of around 10 μW, filtered between 465 and 480 nm) delivered to the brain through a fiber optic patch cord (MFP_400/430/1100-0.48_2m_FCM-MF2.5). The emission fluorescence passed from the brain through the same patch cords and was filtered (500–520 nm), amplified, detected, and demodulated in real-time by the system. Demodulated fluorescence signals were saved at a rate of about 1 kHz. Modulation at the 405 nm wavelength was not used for processing GCaMP signals.

#### Inscopix data acquisition

Data were acquired with nVista 3.0 using Inscopix Data Acquisition Software v1.7.1 (Inscopix) at 25 FPS, LED power at 0.3 mW/mm^2^. To synchronize imaging data with behavior, we recorded TTL sync pulses from the microscope and TTLs from the waveform generator (pulse pal) used to control video frame acquisitions.

#### Histology in brain slices

Mice were injected with euthasol and perfused with 4% PFA dissolved in 1x PBS. Brains were extracted and post-fixed in 4% PFA for 12–24 H, after which they were cryoprotected in 30% sucrose. Cryosections of the frozen tissue (40 µm slices) were made and stamped directly onto glass microscope slides. Slices were washed with PBS or, for immunohistochemistry, PBS+0.4% Triton (PBST). Then, for immunostaining, a blocking buffer (PBST with 2% normal donkey serum and 1% BSA) was applied for 30 min, followed by incubation by a primary antibody at 4 °C for 12–24 H. Following primary antibody incubation, slides were washed with PBST (5 rounds of 10 min each) followed by incubation at room temperature in a secondary antibody for 2 H, and a final set of washes in PBS (5 rounds of 10 min each). Stained or unstained slides were then dried and coverslipped with a mounting medium (EMS Immuno Mount DAPI and DABSCO, Electron Microscopy Sciences, 17989-98, lot 180418). After at least 12 H of drying, slides were imaged with a digital robotic slide scanner (NanoZoomer S60, C13210-01, Hamamatsu). The following antibodies were used: rabbit anti-GFP (Novus Biologicals CAT# NB600-308) 1:500, Donkey anti-Rabbit Alexa 647 (ThermoFisher Scientific CAT# A-31573), 1:1000.

#### Video recordings

For the chronic social defeat stress and homecage assays (described below), we used a BlackFly S camera (FLIR, BFS-U3-32S4M-C: 3.2 MP, 118 FPS, Sony IMX252, Mono) and recorded videos with Motif software (Loopbio). The camera was triggered by a Pulse Pal v2 (Sanworks, #1102) at a rate of 100 frames per second (FPS). The camera was oriented at 90° towards the side of the preparation and also captured the top-down view of the preparation with a mirror mounted at a 40° angle above the horizontal.

For all other behavioral assays, recordings were performed using an analog camera and Ethovision (Noldus) software which was used to track the mice.

#### Chronic social defeat stress (CSDS)

Mice were placed in the cage of a novel aggressor for 5 min of free interaction. Mice that sustained more than pinpoint wounds were euthanized. Afterward a perforated acrylic barrier (Tap Plastics) was placed between the mice. 24 H later, mice were placed in the cage of a new aggressor. This continued for a total of 10 days. Unstressed controls were pair-housed with a perforated barrier separating the two mice. They were handled and their cages rotated each day for 10 days. Following the defeat on day 10, aggressors were removed and all mice were singly housed in the shoebox cages through the remaining stages of the experiment.

Defeated mice were housed in the shoebox home cages (#5 Expanded Mouse Cage 22.2 x 30.8 x 16.2 cm, Thoren Caging Systems, Inc.). For recordings, food was removed from the cage and the typical stainless steel lids were replaced with a custom-cut sheet of clear acrylic (Tap Plastics) with a hole for patch cables to run through. Shoebox cages were placed underneath the angled mirror and video recordings were made as described above.

#### Social interaction test

Mice were placed in a two-chamber arena (56 cm x 24 cm) for 5 minutes with two empty mesh pencil cups in the far left and right corners. The mouse was then removed from the chamber, and a novel social target was placed beneath one cup. The mouse was then returned to the recording chamber for an additional 5 minutes. Mouse location was tracked via Ethovision (Noldus). We quantified the time spent within the social zone (up to 8 cm from the perimeter of the enclosure). Following day 10 of defeat, the time spent in the social interaction zone when the social target was a novel swiss webster aggressor was used to delineate resilient and susceptible mice. We defined susceptible as 1 standard deviation below the mean social interaction time of the unstressed control group^27^.

#### Elevated plus maze

Following CSDS, mice were placed in the center of an elevated plus maze (2 enclosed arms and 2 open arms; each arm 76 cm long and 6.5 cm wide). The mouse explored the maze for 7 minutes, while its centroid location was tracked via Ethovision (Noldus). The time spent in the open arms and center of the maze was measured.

#### Open field test

Mice were placed into the center of an empty area (50 cm x 50 cm). Lamps were used to illuminate the arena on the left and right so that there was a shaded area along the left and right walls. The center was 42 cm x 42 cm centered at the center of the arena. The animal explored for 10 minutes, while its centroid location was tracked via Ethovision (Noldus). The time spent in the center was measured.

#### Chamber exploration (immobility)

Animals were placed in a neutral two-chamber arena (56 cm x 24 cm) for 5 minutes. Mouse location and speed were continuously tracked via Ethovision (Noldus). We quantified the amount of time spent immobile (speed < 1 cm/s).

#### Novelty-suppressed feeding

Mice were placed in the corner of a brightly lit, neutral chamber (25 x 25 cm) with a single yogurt chip placed in the center of the camber on a plastic platform. The latency to initiate consumption of the treat was scored. After the first consumption bout, mice were placed back in their home cages with *ad libitum* food access. Mice were food deprived for 18 H.

#### Homecage assay

After defeat, video and neural recordings were taken as mice freely interacted with novel juvenile male C57BL6/J or Swiss Webster social targets. Recordings took place on the same setup described above for recording defeat. Behavior occurred in clean shoebox cages of the same type that was used for defeat. After at least 1 minute of baseline recording mice were presented with the novel mouse. Video (100 FPS) and neural recordings were taken for an additional 9 minutes. Sniffing and pursuit of the social target were then hand-scored.

#### Behavioral schedule for each cohort

Animals undergoing fiber photometry recordings were subject to the following assays in this order: social interaction assay (with a Swiss Webster, C57BL6/J, and then AKR/J social target in that order on different days), chronic social defeat stress (10 days), social interaction assay (with a Swiss Webster, C57BL6/J and then AKR/J social target in that order on different days), and homecage assay (C57BL6/J and then Swiss Webster social target, or vice versa, randomly assigned on different days). Altogether there were two cohorts of fiber photometry recordings. Animals undergoing cellular resolution calcium imaging were subject to the following assays in this order: social interaction assay (with a Swiss Webster, C57BL6/J, and then AKR/J social target in that order on different days), chronic social defeat stress, social interaction assay (with a Swiss Webster, C57BL6/J and then AKR/J social target in that order on different days), elevated plus maze assay, novelty suppressed feeding assay, open field test, and homecage assay (C57BL6/J and then Swiss Webster social target, or vice versa, randomly assigned on different days). Animals undergoing optogenetic and fos experiments were subject to the following assays in order: chronic social defeat stress, social interaction assay with a Swiss Webster, elevated plus maze assay, open field test, and novelty suppressed feeding assay. Mice undergoing optogenetic and Fos experiments were also placed for 10 min into the cage of a novel Swiss Webster aggressor that was restrained under a wire cup prior to being sacrificed an hour later for Fos analysis.

#### Closed-loop, behavior-triggered stimulation during defeat

To deliver closed-loop attack-triggered optogenetic stimulation (Figure 5), we used a pre-trained random forest (described above) for inference on video frames streamed in real-time.

Images were acquired using a FLIR BlackFlyS camera connected directly to our behavior inference computer (Ubuntu 18.04.06, equipped with a Nvidia GeForce GTX 1070 Ti graphics card). Using publicly available custom code, each video frame was captured by Motif (Loopbio) software and sent as an input to our pre-trained DeepLabCut network for estimating the positions of the interacting mice. The 12 features we defined above were calculated with minor modifications (no smoothing, using adjacent frames for instantaneous speed and velocity features). We trained a separate binary random forest classifier to detect attack behavior from the unsmoothed features using the same training set as mentioned above for the offline analysis. After detection of an attack video frame, a serial signal was passed through the USB to an Arduino, which translated the signal into a TTL for triggering the laser light delivery protocol. The frame capture, behavior inference, and trigger delivery code were run in an open loop and could achieve a speed of about 20 FPS. A list of time stamps from each frame and its probability of behavior detection and whether a trigger was delivered were saved for synchronization.

Blue (447 nm, 5 mW for ChR2 animals),green (532 nm, 0.8 mW for ChRmine animals), or yellow (593 nm, 3mW for NpHr animals) lasers were connected to a commutator (Doric Lenses, FRJ_1x2i_FC-2FC_0.22), which led to 200-μm diameter patch cords that were fastened to the implants of mice through plastic sleeves surrounded by black electric tape. For activation experiments, phasic stimulation was delivered (5 pulses of 5 ms in duration at 20 Hz) when an attack frame was detected. Once an attack frame was detected, laser stimulation could not be triggered until all 5 pulses had occurred. Pulses were continuous so long as attack behavior was ongoing. For inhibition experiments, continuous yellow light was delivered for 1s when an attack frame was detected. Similar to activation experiments, laser stimulation could not be triggered until after the 1s of light had occurred and light was continuous so long as attack behavior was ongoing. Laser stimulation pulses were recorded for synchronization. Stimulation was performed across all 10 days of defeat, but not during post hoc testing.

#### Slice electrophysiology

For the slice physiology data in Figure S7A-D, Vglut2::Cre adult mice 10-12 weeks old were injected with 80 nL of AAV5-EF1α-DIO-hChR2(H134R)-eYFP (titer: 1.2e13 GC/ml; manufacturer: PNI Viral Core Facility) bilaterally into the LHb (10+ days before the experiment; Figure S7A-D). On the day of the experiment, mice were anesthetized with isoflurane and decapitated to remove the brain. After extraction, the brain was immersed in ice-cold NMDG ACSF (92 mM NMDG, 2.5 mM KCl, 1.25 mM NaH2PO4, 30 mM NaHCO3, 20 mM HEPES, 25 mM glucose, 2 mM thiourea, 5 mM Na-ascorbate, 3 mM Na-pyruvate, 0.5 mM CaCl2 ·4H2O, 10 mM MgSO4 ·7H2O, and 12 mM N-Acetyl-L-cysteine; pH adjusted to 7.3–7.4) for approximately 2 min. Afterwards, coronal slices (300 μm) were sectioned using a vibratome (VT1200s, Leica, Germany) and then incubated in NMDG ACSF at 34 °C for approximately 15 min. Slices were then transferred into a holding solution of HEPES ACSF (92 mM NaCl, 2.5 mM KCl, 1.25 mM NaH2PO4, 30 mM NaHCO3, 20 mM HEPES, 25 mM glucose, 2 mM thiourea, 5 mM Na-ascorbate, 3 mM Na-pyruvate, 2 mM CaCl2 ·4H2O, 2 mM MgSO4 ·7H2O and 12 mM N-Acetyl-L-cysteine, bubbled at room temperature with 95% O2 /5% CO2) for at least 60 min until recordings were performed. Whole-cell recordings were performed using a Multiclamp 700B (Molecular Devices) using pipettes with a resistance of 7-8 MΩ filled with an internal solution containing 120 mM potassium gluconate, 0.2 mM EGTA, 10 mM HEPES, 5 mM NaCl, 1 mM MgCl2, 2 mM Mg-ATP and 0.3 mM NA-GTP, with the pH adjusted to 7.2 with KOH and the osmolarity adjusted to around 289 mmol kg−1 with sucrose. During recordings, slices were perfused with a recording ACSF solution (120 mM NaCl, 3.5 mM KCl, 1.25 mM NaH2PO4, 26 mM NaHCO3, 2 mM MgSO4, 2 mM CaCl2, and 11 mM D-(+)-glucose) containing the AMPA receptor blocker NBQX (10 mM), and the NMDA receptor blocker AP5 (25 mM) to avoid secondary activation of the patched cells. Infrared differential interference contrast-enhanced visual guidance was used to select neurons that were 3–4 cell layers below the surface of the slices. We targeted neurons in lHb by using the Paxinos atlas as reference. The recording solution was delivered to slices via superfusion driven by a peristaltic pump (flow rate of 4–5 ml/min) and was held at room temperature. The neurons were held at −65 mV (voltage clamp), and the pipette series resistance was monitored throughout recordings. If the series resistance was >30MΩ, the cell was highly depolarized (>-40 mV RMP) or the leak current was >250 pA the data were discarded. Pipette offsets were nulled before seal formation and pipette capacitance was compensated in the cell-attached configuration once a giga-seal was obtained. Whole-cell currents were filtered at 4 kHz online and digitized and stored at 10 KHz (Clampex 10; MDS Analytical Technologies). Bridge balance was used to compensate for series resistance in the current clamp experiments. All voltage clamp experiments were recorded after series resistance compensation for 7 MΩ. Membrane potentials were not adjusted for the liquid junction potential. All optical stimulation was delivered with a 473 nm LED (Lumincor).

Photocurrents were measured in voltage clamp configuration where we recorded 10 sweeps (2.8 s/ sweep) of light-evoked oEPSCs from 5 ms light stimulation with a power density of 8 mW/mm^2^. This power approximately matches the estimated in vivo stimulation parameters 100 um below the fiber tip. Reported photocurrents are the mean peak current over 10 sweeps of optical stimulation.

Spike fidelity at different stimulation frequencies was measured in current-clamp by applying 5 sweeps containing 250 ms trains of 5, 10, 20 and 40 Hz of 5 ms wide 8 mW/mm^2^ 473 nm pulses. Reported spike probabilities are calculated from the mean spike fidelity of each cell across the 5 sweeps.

#### Behavior for brainwide fos analysis (Figure 6)

Eight or nine days after CSDS with closed-loop attack-triggered LHb stimulation (i.e., after the completion of all post-CSDS tests), mice were placed for 10 min into the cage of a novel Swiss Webster aggressor that was restrained under a wire cup and were then sacrificed one hour later for Fos analysis. There was no LHb stimulation on the Fos day. Altogether we had three cohorts of mice: a cohort that received LHb stimulation (which had *N* = 10 mice expressing ChR2 and *N* = 5 mice expressing YFP), and two cohorts (of *N* = 19 and *N* = 20 mice each) that did not receive stimulation. Mice expressing YFP and mice from the two unstimulated cohorts were combined into “unstimulated mice” (*N* = 44).

For analysis of susceptible vs. resilient groups, mice were defined as susceptible if their social interaction time in the SI test was less than one standard deviation below that of unstressed controls (from our fiber photometry dataset, Figure 1C), otherwise they were considered resilient.

#### Tissue clearing and immunolabeling

Mice were deeply anesthetized (2 mg/kg Euthasol ip.) and then transcardially perfused with ice-cold PBS + heparin (20 U/mL; Sigma H3149) followed by ice-cold 4% PFA in PBS. Brains were then extracted and post-fixed overnight in 4% PFA at 4°C.

Brains were cleared and immunolabeled using an iDISCO+ protocol as previously described^56,57^. All incubations were performed at room temperature unless otherwise noted.

Clearing: Brains were serially dehydrated in increasing concentrations of methanol (Carolina Biological Supply 874195; 20%, 40%, 60%, 80%, 100% in doubly distilled water (ddH2O); 45 min–1 h each), bleached in 5% hydrogen peroxide (Sigma H1009) in methanol overnight, and then serially rehydrated in decreasing concentrations of methanol (100%, 80%, 60%, 40%, 20% in ddH2O; 45 min–1 h each).

Immunolabeling: Brains were washed in 0.2% Triton X-100 (Sigma T8787) in PBS, followed by 20% DMSO (Fisher Scientific D128) + 0.3 M glycine (Sigma 410225) + 0.2% Triton X-100 in PBS at 37°C for 2 d. Brains were then washed in 10% DMSO + 6% normal donkey serum (NDS; EMD Millipore S30) + 0.2% Triton X-100 in PBS at 37°C for 2–3 d to block non-specific antibody binding. Brains were then twice washed for 1 h at 37°C in 0.2% Tween-20 (Sigma P9416) + 10 mg/mL heparin in PBS (PTwH solution) followed by incubation with primary antibody solution (rabbit anti-Fos, 1:1000; Synaptic Systems CAT#226008; chicken anti-GFP, 1:500; Aves CAT#GFP-1020) in 5% DMSO + 3% NDS + PTwH at 37°C for 7 d. Brains were then washed in PTwH 6× for increasing durations (10 min, 15 min, 30 min, 1 H, 2 H, overnight) followed by incubation with secondary antibody solution (Alexa Fluor 647 donkey anti-rabbit, 1:200; Abcam CAT#ab150075; Alexa Fluor 594 donkey anti-chicken, 1:500; Jackson Immuno CAT#703-585-155) in 3% NDS + PTwH at 37°C for 7 d. Brains were then washed in PTwH 6× for increasing durations again (10 min, 15 min, 30 min, 1 H, 2 H, overnight).

Final storage and imaging: Brains were serially dehydrated in increasing concentrations of methanol (20%, 40%, 60%, 80%, 100% in ddH2O; 45 min–1 h each), then incubated in a 2:1 solution of dichloromethane (DCM; Sigma 270997) and methanol for 3 H followed by 2× 15-min washes 100% DCM. Before imaging, brains were stored in the refractive index-matching solution dibenzyl ether (DBE; Sigma 108014).

### Quantification and statistical analysis

#### Behavioral Annotation

Ground truth for supervised classification of behaviors during defeat (Figure 3-5) was determined by hand annotations of videos scored with BORIS^101^. The following behaviors were annotated: mouse being attacked, mouse being sniffed, mouse fighting back, stressed mouse running away, and mice being vigilant.

#### Marklerless pose tracking

For fiber photometry and optogenetics experiments, DeepLabCut^102^ was used for tracking the

positions of the stressed and aggressor mice during defeat. The training set included 1603 frames from 350 videos across 35 mice from 2 separate defeat cohorts). The following points were tracked:

● TopStressNose
● TopStressRightEar
● TopStressLeftEar
● TopStressFiberBase
● TopStressTTI
● TopStressTTip
● TopAggNose
● TopAggRightEar
● TopAggLeftEar
● TopAggTTI
● TopAggTTip
● BottomStressNose
● BottomStressRightEar
● BottomStressLeftEar
● BottomStressFiberBase
● BottomStressRightForePaw
● BottomStressLeftForePaw
● BottomStressTTI
● BottomStressTTip
● BottomAggNose
● BottomAggRightEar
● BottomAggLeftEar
● BottomAggTTI
● BottomAggTTip
● TopDividerRight
● TopDividerLeft
● BottomDividerTopRight
● BottomDividerTopLeft

(TTip: Tail tip, TTI: Tail-torso interface, Stress: stressed mouse, Agg: aggressor) DLC training was run for 1.03 million iterations with default parameters: training frames selected by kmeans clustering of each video session in the training set, trained on 95% of labeled frames, initialized with ResNet-50, batch size of 4.

#### Feature definition

To define the defeated mouse’s posture with respect to his environment and the aggressor, we converted pose data to the following behavioral features:

1. Between centroid distance: Euclidean distance between the midpoint between each mouse’s tail-body interface and nose, defined by the top-down view.
2. Distance between aggressor nose and stressed mouse rear
3. Distance between aggressor nose and stressed mouse nose
4. Between centroid velocity: instantaneous (every 8 frames or 0.08 s) change in between centroid distance, median smoothed with a window of 0.17 s.
5. Aggressor speed: instantaneous distance between centroid position every 10 frames, smoothed as above
6. Stressed mouse speed: same as above
7. Orientation of aggressor with respect to stressed mouse
8. Orientation of stressed mouse with respect to the aggressor
9. Height of the aggressor: side view nose Y position
10. Height of stressed mouse: same as above
11. Distance of stressed mouse from the closest short wall of the cage: based on top-down view
12. Distance of the stressed mouse from the closest long wall of the cage

#### Feature preprocessing

Before using features for random forest classification or unsupervised behavior classification, features were preprocessed. Features were truncated to fall within the 1st and 99th percentile of all recorded data for each feature (to remove extreme outliers), smoothed across time with a Gaussian filter of 0.20 s, and rescaled from −1 to 1 (sklearn.preprocessing.MixMaxScaler) within each session to account for variability in mouse size and slightly varying camera angle or height. We chose to rescale features so that no single feature dominates owing to higher magnitude while maintaining the original feature distributions and their covariances, properties that would not be maintained if each feature were normalized independently to unit variance, for example.

#### Random Forest classification

For automated identification of behaviors across our entire video dataset (Figure 3-5, Figure S3-4,7), we trained supervised random forest classifiers using manually annotated data. Behaviors of interest during defeat included being attacked, being investigated, fighting back, fleeing, and being vigilant. These each were classified by a separate binary random forest classifier (Scikit-learn). The training and testing set consisted of twenty videos each. Ground truth was determined by manual annotation (BORIS) for frames in which the behavior was occurring (see above).

For each classified behavior, the feature matrix included the 12 features described above for each video frame. The objective matrix was a binary indicator if the behavior was manually annotated in that frame. The training set was composed of all the frames in which the behavior was present and a randomly selected equal number of frames in which the behavior was absent. The classifier was trained with a maximum depth of 2 and 100 estimators.

The probability threshold for detecting behaviors was set to the most permissive possible without exceeding a false positive rate of 3% on the training set. Evaluation was conducted by plotting the receiver operator curve on the held-out testing set (Figure S3D-E).

#### Unsupervised behavior classification

To characterize behaviors as stereotyped features repeated throughout time, we followed previous work^103^ in using a low-dimensional embedding of the original features and defining behaviors as high density clusters in that low-dimensional embedding to create Figure 3C. To achieve dense clusters, we embedded our behavior features using *t*-SNE, which preserves small pairwise distances and thereby retains clustering of nearby points.

Generating this manifold involved a technique known as importance sampling, which enabled us to create a final embedding that included behaviors that might be rare or nuanced, and therefore under-represented in a uniform sampling over time. Importance sampling includes two rounds of *t*-SNE embedding. First, around 12,000 frames of behavior were uniformly sampled in time across all videos (*N* =350) analyzed. Those features were embedded into a two-component *t*-SNE manifold (sklearn.manifold.TSNE with perplexity = 100). The embedded space was binned into a 50 × 50 histogram, smoothed with a 2D Gaussian kernel (with a standard deviation of 2.5), and parcellated into 17 clusters with watershed (skimage.morphology.watershed) over the smoothed histogram. When then used a multilayer perceptron (sklearn.neural_network.MLPRegressor, hidden layer size of 400 × 200 × 50 units) to represent data from every video frame in 2D *t*-SNE space, and thus to fall into 1 of the 17 clusters defined in this space.

Because we are interested in attack behaviors, we repeated these steps but with a subset of frames that were biased to have more attack frames. To sample from aggressive behavior, we characterized the overlap between random forest-classified attack frames and the clusters in *t*-SNE space. From the cluster that most overlapped with attack, we sampled five random frames from every defeat session. From the 16 remaining non-attack clusters, we sampled 2 random frames from every defeat session. Thus, from 35 males undergoing defeat for 10 days, we sampled (2 × 16) frames from non-attack clusters and 5 frames from the attack cluster on each day for each mouse for a total of 35 × 10 × (16 × 2 + 5) = 12,950 frames. From these sampled frames, we again embedded the 12-dimensional behavior features into two-component *t*-SNE space. The full set of video frames was then mapped into this final *t*-SNE manifold using another multilayer perceptron. Then a 2D histogram of that perceptron-mapped 2D data was smoothed with a Gaussian kernel (with a standard deviation of 1.5) and divided into 17 clustered again with watershed. Gaussian kernels in both *t*-SNE steps were chosen by rounding to the nearest 0.5 and to yield 10-20 clusters from watershed clustering.

To plot behavior data from our optogenetics experiments in the same *t*-SNE space (Figure 5E), the perceptron (sklearn.neural_network.MLPRegressor) used to learn the 12 features from the fiber photometry experiments to create the *t*-SNE mapping (Figure 3C) was applied to the 12 features from the optogenetics experiments in the same way.

#### Processing of fiber photometry data

For Figure 2, raw fluorescence data in each session was converted into dF/F using a moving average (window of 30 s) to calculate F_0_. The data was then z-scored by diving by the standard deviation of the dF/F signal across the entire session. For Figures 3-4, defeat recordings for each mouse were converted into dF/F using the average of each session rather than a moving window to calculate F_0_. Sessions were then appended and z-scoring was performed by dividing by the standard deviation of all 10 days of defeat.

#### Pre-processing of cellular resolution calcium imaging data

Initial pre-processing was done in IDPS 1.8.0 (Inscopix Data Processing Software). Videos were spatially downsampled by a factor of 4 and motion-corrected with a translational correction algorithm based on cross-correlations computed on consecutive frames. Videos were subsequently exported as .tiff files and further motion-corrected using NoRMCorre^104^. After motion correction, the CNMFe algorithm^105,106^ was used to identify neurons and obtain their fluorescence traces. The fluorescence traces were then z-scored using the same method as for the fiber photometry data described above. To calculate Ca^2+^ transient rates (Figure 2Y-Z), we identified events based on the deconvolved events identified by CNMFE^105^.

#### Inter-cell activity synchrony

To determine how synchronous the neurons of susceptible and resilient mice were before and after CSDS, we calculated the fraction of the recorded population that had at least one transient in each 500 ms timebin in the neutral chamber. We then plotted the cumulative distribution of this data for each recording, and then averaged across recordings, before and after defeat (Figure S2H).

#### Determining significant neurons in calcium imaging data

To determine which neurons were significantly activated during the SI test with the aggressor strain (Figure 2V-X), we created a null distribution that maintains the autocorrelations of the real neural data, but does not preserve the temporal relationship to behavioral events^107^ by shifting the dF/F trace of each cell 5 s for each shifted sample, and repeat 1000 times to create 1000 traces (Figure S2D). We then time-locked each of the shifted 1000 traces and the real trace to entry into the social zone (Figure S2E). We then averaged over the time window of interest (0.5 s to 2.5 s post social zone entry) for the real and 1000 shifted traces and calculated which neurons were in the 2.5th percentile (inhibited) or 97.5th percentile (activated) to determine significance of *p*<0.05 for a 2-sided test (Figure S2F-G).

#### Plotting neural data in behavioral *t*-SNE space

We wanted to see the corresponding neural activity within the behavioral clusters identified from the *t*-SNE map in Figure 3D, 4A-C). Because peak neural activity to proximity-related behaviors occurred 0.5 s after the start of attack (Figure 3K-N), we shifted the fluorescent data of each video forward 0.5 s. We then identified the neural activity corresponding with each video frame and also where that video frame is located in the 50 x 50 *t*-SNE embedding. Then we smoothed the neural data plotted in *t*-SNE space with a 2D Gaussian kernel (with a standard deviation of 1.5).

#### Fos light sheet imaging

Cleared and immunolabeled brains were glued (Loctite 234796) ventral side-down to a 3D-printed holder and imaged in DBE using a dynamic axial sweeping light sheet fluorescence microscope (Life Canvas SmartSPIM). Images were acquired using a 3.6×/0.2 NA objective with a 3,650 µm×3,650 µm field-of-view onto a 2,048 px×2,048 px sCMOS camera (pixel size: 1.78 µm×1.78 µm) with a spacing of 2 µm between horizontal planes (nominal z-dimension point spread function: 3.2–4.0 µm). Imaging the entire brain required 4×6 tiling across the horizontal plane and 3,300–3,900 total horizontal planes. Autofluorescence channel images were acquired using 488 nm excitation light at 20% power and 2ms exposure time, Fos channel images were acquired using 639 nm excitation light at 90% power and 2ms exposure time, and YFP channel images were acquired using 561 nm excitation light at 20% power and 2-ms exposure time to confirm ChR2-YFP expression.

After acquisition, tiled images for the Fos channel were first stitched into a single imaging volume using the TeraStitcher C++ package (https://github.com/abria/TeraStitcher). These stitching parameters were then directly applied to the tiled autofluorescence channel images, yielding two aligned 3D imaging volumes with the same final dimensions. After tile stitching, striping artifacts were removed from each channel using the Pystripe Python package (https://github.com/chunglabmit/pystripe).

We registered the final Fos imaging volume to the Allen CCF using the autofluorescence imaging volume as an intermediary. We first downsampled both imaging volumes by a factor of 5 for computational efficiency. Autofluorescence→atlas alignment was done by applying an affine transformation to obtain general alignment using only translation, rotation, shearing, and scaling, followed by applying a b-spline transformation to account for local nonlinear variability among individual brains. Fos→autofluorescence alignment was done by applying only affine transformations to account for brain movement during imaging and wavelength-dependent aberrations. Alignment transformations were computed using the Elastix C++ package (https://github.com/SuperElastix/elastix). These transformations allowed us to transform Fos^+^ cell coordinates first from their native space to the autofluorescence space and then to Allen CCF space.

Deep learning-assisted cell detection pipeline. We first use standard machine vision approaches to identify candidate Fos^+^ cells based on peak intensity and then use a convolutional neural network to remove artifacts. Our pipeline builds upon the ClearMap Python package^56,57^ (https://github.com/ChristophKirst/ClearMap2) for identifying candidate cells and the Cellfinder Python package^108^ (https://github.com/brainglobe/cellfinder) for artifact removal.

Cell detection: ClearMap operates through a series of simple image processing steps. First, the Fos imaging volume is background-subtracted using a morphological opening (disk size: 21 px). Second, potential cell centers are found as local maxima in the background-subtracted imaging volume (structural element shape: 11 px). Third, cell size is determined for each potential cell center using a watershed algorithm (see below for details on watershed detection threshold). Fourth, a final list of candidate cells is generated by removing all potential cells that are smaller than a preset size (size threshold: 350 px). We confirmed that our findings were consistent across a wide range of potential size thresholds.

We implemented three changes to the standard ClearMap algorithm. First, we de-noised the Fos imaging volume using a median filter (function: scipy.ndimage.median_filter; size: 3 px) before the background subtraction step. Second, we dynamically adjusted the watershed detection threshold for each sample based on its fluorescence intensity. This step was important for achieving consistent cell detection performance despite changes in background and signal intensity across cohorts and samples due to technical variation in clearing, immunolabeling, and imaging. Briefly, we selected a 1,000 px×1,000 px×200 px subvolume at the center of each sample’s Fos imaging volume. We then median filtered and background subtracted this subvolume as described above. We then used sigma clipping (function: astropy.stats.sigma_clipped_stats; sigma=3.0, maxiters=10, cenfunc=’median’, stdfunc=’mad_std’) to estimate the mean background (non-cell) signal level for this subvolume, µ_bg_, and set each sample’s watershed detection threshold to 5*µ_bg_ (low-signal cohorts) or 10*µ_bg_ (high-signal cohorts). Third, we removed from further analysis all cell candidates that were located outside the brain, in the anterior olfactory areas or cerebellum (which were often damaged during dissection), or in the ventricles, fiber tracts, and grooves following registration to the Allen CCF.

Cell classification: One limitation of the watershed algorithm implemented by ClearMap is that it identifies any high-contrast feature as a candidate cell, including exterior and ventricle brain edges, tissue tears, bubbles, and other aberrations. To overcome this limitation, we re-trained the 50-layer ResNet implemented in Keras (https://keras.io) for TensorFlow (https://www.tensorflow.org) from the Cellfinder Python package to classify candidate Fos^+^ cells in our high-resolution light sheet imaging dataset as true Fos^+^ cells or artifacts. This network uses both the autofluorescence and Fos channels during classification because the autofluorescence channel has significant information about high-contrast anatomical features and imaging aberrations. We first manually annotated 2,000 true Fos^+^ cells and 1,000 artifacts from each of four brains across two technical cohorts using the Cellfinder Napari plugin, for a total training dataset of 12,000 examples. We then re-trained the Cellfinder network (which had already been trained on 2p images of GFP^+^ cells) using TensorFlow over 100 epochs with a learning rate of 0.0001 and 1,200 examples (10% of the training dataset) held out for validation. Re-training took 4 d 16 min 41 sec on a high performance computing cluster using 1 GPU and 12 CPU threads. We achieved a final validation accuracy of 98.33%. Our trained convolutional neural network removed 15.99% ± 0.58% (mean ± s.e.m.; range: 2.96% – 32.71%; n = 99 brain^58^) of cell candidates from ClearMap as artifacts.

Atlas registration: We used the ClearMap interface with Elastix to transform the coordinates of each true Fos^+^ cell to Allen CCF space using the transformations described above. We then used these coordinates to assign each Fos^+^ cell to an Allen CCF brain region. For each sample, we generated a final data structure containing the Allen CCF coordinates (*x*,*y*,*z*), size, and brain region for each true Fos^+^ cell.

#### Fos density maps

We generated 3D maps of Fos^+^ cell density by applying a gaussian kernel-density estimate (KDE) (function: *scipy.stats.gaussian_kde*) in Python to all Fos^+^ cells across all animals within a given experimental condition (for example, susceptible mice). These maps are visualized in Figure 6D (interactive visualization: https://www.brainsharer.org/ng/?id=876).

We first generated a table containing the Allen CCF coordinates (*x*,*y*,*z*) for every Fos^+^ cell in every animal within an experimental condition. At this stage, we listed each cell twice (once with its original coordinates and once with its ML (*z*) coordinate flipped to the opposite hemisphere) in order to pool data from both hemispheres. We used a modified symmetrical version of the Allen CCF to facilitate this. We then assigned each cell a weight equal to the inverse of the total number of Fos^+^ cells in that animal to ensure that each animal within an experimental condition would be weighted equally. We then fit a 3D gaussian KDE for each experimental condition using the scipy.stats.gaussian_kde function, and manually set the kernel bandwidth for every experimental condition to be equal at 0.04. We then evaluated this KDE at every voxel in the Allen CCF (excluding voxels outside the brain or in anterior olfactory areas, cerebellum, ventricles, fiber tracts, grooves) to obtain a 3D map of Fos^+^ density for each condition. Lastly, we normalized the KDE for each experimental condition by dividing by its sum as well as the voxel size of the atlas, (0.025 mm)^3^, to generate a final 3D map with units of “% Fos^+^ cells per mm^3^”. To examine the difference in Fos^+^ cell density across conditions, we simply subtracted the 3D KDE volumes for the two conditions, e.g. Resilient(Stim) – Susceptible(Stim), and then plotted coronal sections through this subtracted volume with Allen CCF boundaries overlaid. The colorbar limits for all KDE figures are ±0.5% Fos^+^ cells per mm^3^.

#### Fos GLMMs

We adopted a GLMM approach to analyze the Fos data (Figure 6E-J and Figure S8A-D). This allowed us to model the contribution of SI time (time near aggressor; z-scored across all 54 mice) and/or Stim/NoStim to neural activation in each brain region, while also accounting for the overdispersed, discrete nature of the data by employing a negative binomial link function.

We first fit a GLM for each brain region using the glmmTMB R package (https://github.com/glmmTMB/glmmTMB) with a *nbinom2* link function and the formula, ***Counts* ∼ *SI Time* + ln(*Total Counts*)**, where ***Counts*** is the number of Fos^+^ cells in a brain region, ***SI Time*** is an animal’s Social Interaction Test score, and **ln(*TotalCounts*)** is an offset term for the total number of Fos^+^ cells in each sample. We used the coefficient estimate and standard error (*Z*-value = estimate/standard error) as a proxy for modulation by susceptibility and resilience.

We calculated a *p*-value for each brain region using this statistic, and corrected for a 10% false discovery rate across all brain regions using the Benjamini-Krieger-Yekutieli two-step procedure. We performed this analysis separately for LHb-stimulated mice (Figure 6E) and for unstimulated control mice (Figure 6F). For the unstimulated control mice, our regressions also included a random effect of **(1+*SI Time*|*Cohort*)** to account for differences in behavior and Fos labeling across cohorts. Mice in the “unstimulated” group were made up of three cohorts: mice from the LHb activation experiments that were injected with a virus expressing YFP, and two cohorts of mice that underwent CSDS but were not injected with any viruses.

We then fit similar regressions using ***NoStim–Stim*** (a categorical variable, 1 for NoStim and 0 for Stim) as a regressor instead of ***SI Time*** and including both the LHb-stimulated and the unstimulated control mice, and then calculated *Z*-values (coefficient estimate/standard error) for each of these variables (Figure 6G). These regressions also included a random effect of **(1|*Cohort*)** to account for differences in behavior and Fos labeling across cohorts.

We also fit GLMMs with both SI Time and LHb stimulation, and their interaction, as regressors and including both the LHb-stimulated and the unstimulated control mice (Figure S8A-D): ***Counts* ∼ *SI Time* + *Stim* + *SI Time:Stim* + ln(*Total Counts*) + (1+*SI Time*|*Cohort*)**. Here, we treated ***Stim*** as a categorical variable where Stim=1 and NoStim=0. We then calculated *p*-values and significance for each regressor and each region as described above.

In Figure 6H and S8D, we used the Violinplot MATLAB package (https://github.com/bastibe/Violinplot-Matlab) to plot the distribution of Z-values described above for all brain regions in cortex (Cerebral Cortex, by Allen CCF designation), forebrain nuclei (Cerebral Nuclei, Thalamus, Hypothalamus, by Allen CCF designation), and midbrain/hindbrain (Midbrain, Pons, Medulla, by Allen CCF designation). We used Kolmogorov-Smirnov tests to assess whether these distributions were significantly different from each other across these thre subdivisions.

In Figure 6I, we took the pairwise correlation across brain regions for the ***SI Time*** regressor *Z*-scores for the LHb-stimulated mice and the ***NoStim–Stim*** regressor *Z*-scores across all mice. In Figure 6J, we took the pairwise correlation across brain regions for the ***SI Time*** regressor *Z*-scores for the unstimulated control mice and the ***NoStim–Stim*** regressor *Z*-scores across all mice.

#### Fos correlation analysis

To quantify Fos correlations across individual mice (Figure S8E), we considered the LHb-stimulated mice and the unstimulated control mice separately. We first assembled the relative Fos^+^ cell counts (% per mm^3^) for every brain region for each group of mice, then used the built-in MATLAB *corr* function to calculate and visualize pairwise correlations among all brain regions. We then used the used the built-in MATLAB *linkage* function (*method*=’ward’, *metric*=’chebychev’) to create a hierarchical tree using the correlation matrix for the LHb-stimulated mice, and then sorted both correlation matrices using this hierarchical tree.

## Acknowledgments

We thank Esteban Engel, Oliver Huang, Angela Chan, and the PNI Viral Core Facility for AAV production; Jeffrey Stirman and Life Canvas Technologies for light sheet imaging support; the Kleinfeld and Wang labs for collaboration on the Brainsharer web portal; Adrian Sirko and the Princeton Laboratory Animal Resources staff for help with animal husbandry; and C. Peña, M. Murthy, Rob Fetcho, Adelaide Minerva, and other members of the Witten lab for feedback on this work. This work was supported by NIH grants DP1-MH136573 (I.B.W.) and K99-DA059957 (C.A.Z.), the Simons Collaboration on the Global Brain (I.B.W., A.L.F.), the Helen Hay Whitney Foundation (C.A.Z.), the Princeton Innovation Fund (I.B.W., A.L.F.), and the New York Stem Cell Foundation (I.B.W., A.L.F.).

## Author contributions

A.Z. and I.B.W. conceived the project, designed the experiments, and interpreted the data, with input from L.W. and A.L.F. A.Z. collected and analyzed the majority of the data. S.R.J. collected the data for Figure 6 and Figure S8. A.P.V. collected the data for Figure S7A-D. L.W. provided code and advised on analysis for Figures 3–5. C.A.Z. analyzed and interpreted the data for Figure 6 and Figure S8. I.B.W. advised on the data analysis. A.Z., C.A.Z. and I.B.W. wrote the paper.

## References

1. Breslau, N., and Davis, G.C. (1986). Chronic stress and major depression. Arch. Gen. Psychiatry 43, 309–314.

2. McEwen, B.S., and Akil, H. (2020). Revisiting the Stress Concept: Implications for Affective Disorders. J. Neurosci. 40, 12–21.

3. Daviu, N., Bruchas, M.R., Moghaddam, B., Sandi, C., and Beyeler, A. (2019). Neurobiological links between stress and anxiety. Neurobiol Stress 11, 100191.

4. Berton, O., McClung, C.A., Dileone, R.J., Krishnan, V., Renthal, W., Russo, S.J., Graham, D., Tsankova, N.M., Bolanos, C.A., Rios, M., et al. (2006). Essential role of BDNF in the mesolimbic dopamine pathway in social defeat stress. Science 311, 864–868.

5. Krishnan, V., Han, M.-H., Graham, D.L., Berton, O., Renthal, W., Russo, S.J., Laplant, Q., Graham, A., Lutter, M., Lagace, D.C., et al. (2007). Molecular adaptations underlying susceptibility and resistance to social defeat in brain reward regions. Cell 131, 391–404.

6. Yohn, C.N., Dieterich, A., Bazer, A.S., Maita, I., Giedraitis, M., and Samuels, B.A. (2019). Chronic non-discriminatory social defeat is an effective chronic stress paradigm for both male and female mice. Neuropsychopharmacology 44, 2220–2229.

7. Kudryavtseva, N.N., Bakshtanovskaya, I.V., and Koryakina, L.A. (1991). Social model of depression in mice of C57BL/6J strain. Pharmacol. Biochem. Behav. 38, 315–320.

8. Rygula, R., Abumaria, N., Flügge, G., Hiemke, C., Fuchs, E., Rüther, E., and Havemann-Reinecke, U. (2006). Citalopram counteracts depressive-like symptoms evoked by chronic social stress in rats. Behav. Pharmacol. 17, 19–29.

9. Avgustinovich, D.F., Kovalenko, I.L., and Kudryavtseva, N.N. (2005). A model of anxious depression: persistence of behavioral pathology. Neurosci. Behav. Physiol. 35, 917–924.

10. Chaudhury, D., Walsh, J.J., Friedman, A.K., Juarez, B., Ku, S.M., Koo, J.W., Ferguson, D., Tsai, H.-C., Pomeranz, L., Christoffel, D.J., et al. (2013). Rapid regulation of depression-related behaviours by control of midbrain dopamine neurons. Nature 493, 532–536.

11. Cao, J.-L., Covington, H.E., 3rd, Friedman, A.K., Wilkinson, M.B., Walsh, J.J., Cooper, D.C., Nestler, E.J., and Han, M.-H. (2010). Mesolimbic dopamine neurons in the brain reward circuit mediate susceptibility to social defeat and antidepressant action. J. Neurosci. 30, 16453–16458.

12. Diaz, V., and Lin, D. (2019). Neural circuits for coping with social defeat. Curr. Opin. Neurobiol. 60, 99–107.

13. McLaughlin, J.P., Li, S., Valdez, J., Chavkin, T.A., and Chavkin, C. (2006). Social defeat stress-induced behavioral responses are mediated by the endogenous kappa opioid system. Neuropsychopharmacology 31, 1241–1248.

14. Haynes, S.E., Lacagnina, A., Seong, H.S., Afzal, M., Morel, C., Menigoz, A., Rajan, K., Clem, R.L., Mayberg, H.S., Rannie, D.G., et al. (2022). CRF neurons establish resilience via stress-history dependent BNST modulation. bioRxiv, 2022.08.31.505596. 10.1101/2022.08.31.505596.

15. Larrieu, T., Cherix, A., Duque, A., Rodrigues, J., Lei, H., Gruetter, R., and Sandi, C. (2017). Hierarchical Status Predicts Behavioral Vulnerability and Nucleus Accumbens Metabolic Profile Following Chronic Social Defeat Stress. Curr. Biol. 27, 2202–2210.e4.

16. Duclot, F., and Kabbaj, M. (2013). Individual differences in novelty seeking predict subsequent vulnerability to social defeat through a differential epigenetic regulation of brain-derived neurotrophic factor expression. J. Neurosci. 33, 11048–11060.

17. Sandi, C., and Richter-Levin, G. (2009). From high anxiety trait to depression: a neurocognitive hypothesis. Trends Neurosci. 32, 312–320.

18. Henckens, M.J.A.G., Klumpers, F., Everaerd, D., Kooijman, S.C., van Wingen, G.A., and Fernández, G. (2016). Interindividual differences in stress sensitivity: basal and stress-induced cortisol levels differentially predict neural vigilance processing under stress. Soc. Cogn. Affect. Neurosci. 11, 663–673.

19. Nasca, C., Menard, C., Hodes, G., Bigio, B., Pena, C., Lorsch, Z., Zelli, D., Ferris, A., Kana, V., Purushothaman, I., et al. (2019). Multidimensional Predictors of Susceptibility and Resilience to Social Defeat Stress. Biol. Psychiatry 86, 483–491.

20. Radwan, B., Jansen, G., and Chaudhury, D. (2020). Abnormal Sleep Signals Vulnerability to Chronic Social Defeat Stress. Front. Neurosci. 14, 610655.

21. Cerniauskas, I., Winterer, J., de Jong, J.W., Lukacsovich, D., Yang, H., Khan, F., Peck, J.R., Obayashi, S.K., Lilascharoen, V., Lim, B.K., et al. (2019). Chronic Stress Induces Activity, Synaptic, and Transcriptional Remodeling of the Lateral Habenula Associated with Deficits in Motivated Behaviors. Neuron 104, 899–915.e8.

22. Friedman, A.K., Walsh, J.J., Juarez, B., Ku, S.M., Chaudhury, D., Wang, J., Li, X., Dietz, D.M., Pan, N., Vialou, V.F., et al. (2014). Enhancing depression mechanisms in midbrain dopamine neurons achieves homeostatic resilience. Science 344, 313–319.

23. LeClair, K.B., Chan, K.L., Kaster, M.P., Parise, L.F., Burnett, C.J., and Russo, S.J. (2021). Individual history of winning and hierarchy landscape influence stress susceptibility in mice. Elife 10. 10.7554/eLife.71401.

24. Lemos, J.C., Roth, C.A., Messinger, D.I., Gill, H.K., Phillips, P.E.M., and Chavkin, C. (2012). Repeated stress dysregulates κ-opioid receptor signaling in the dorsal raphe through a p38α MAPK-dependent mechanism. J. Neurosci. 32, 12325–12336.

25. Lemos, J.C., Wanat, M.J., Smith, J.S., Reyes, B.A.S., Hollon, N.G., Van Bockstaele, E.J., Chavkin, C., and Phillips, P.E.M. (2012). Severe stress switches CRF action in the nucleus accumbens from appetitive to aversive. Nature 490, 402–406.

26. Nygard, S.K., Hourguettes, N.J., Sobczak, G.G., Carlezon, W.A., and Bruchas, M.R. (2016). Stress-Induced Reinstatement of Nicotine Preference Requires Dynorphin/Kappa Opioid Activity in the Basolateral Amygdala. J. Neurosci. 36, 9937–9948.

27. Willmore, L., Cameron, C., Yang, J., Witten, I.B., and Falkner, A.L. (2022). Behavioural and dopaminergic signatures of resilience. Nature 611, 124–132.

28. Schultz, W., Dayan, P., and Montague, P.R. (1997). A neural substrate of prediction and reward. Science 275, 1593–1599.

29. Cohen, J.Y., Haesler, S., Vong, L., Lowell, B.B., and Uchida, N. (2012). Neuron-type-specific signals for reward and punishment in the ventral tegmental area. Nature 482, 85–88.

30. Steinberg, E.E., Keiflin, R., Boivin, J.R., Witten, I.B., Deisseroth, K., and Janak, P.H. (2013). A causal link between prediction errors, dopamine neurons and learning. Nat. Neurosci. 16, 966–973.

31. Tsutsui-Kimura, I., Matsumoto, H., Akiti, K., Yamada, M.M., Uchida, N., and Watabe-Uchida, M. (2020). Distinct temporal difference error signals in dopamine axons in three regions of the striatum in a decision-making task. Elife 9. 10.7554/eLife.62390.

32. Witten, I.B., Steinberg, E.E., Lee, S.Y., Davidson, T.J., Zalocusky, K.A., Brodsky, M., Yizhar, O., Cho, S.L., Gong, S., Ramakrishnan, C., et al. (2011). Recombinase-driver rat lines: tools, techniques, and optogenetic application to dopamine-mediated reinforcement. Neuron 72, 721–733.

33. Matsumoto, M., and Hikosaka, O. (2007). Lateral habenula as a source of negative reward signals in dopamine neurons. Nature 447, 1111–1115.

34. Li, H., Pullmann, D., and Jhou, T.C. The entopeduncular nucleus drives lateral habenula responses to negative but not positive or neutral affective stimuli. Preprint, 10.1101/408963 10.1101/408963.

35. Hu, H., Cui, Y., and Yang, Y. (2020). Circuits and functions of the lateral habenula in health and in disease. Nat. Rev. Neurosci. 21, 277–295.

36. Stamatakis, A.M., and Stuber, G.D. (2012). Activation of lateral habenula inputs to the ventral midbrain promotes behavioral avoidance. Nat. Neurosci. 15, 1105–1107.

37. Wang, D., Li, Y., Feng, Q., Guo, Q., Zhou, J., and Luo, M. (2017). Learning shapes the aversion and reward responses of lateral habenula neurons. Elife 6, e23045.

38. Yang, Y., Cui, Y., Sang, K., Dong, Y., Ni, Z., Ma, S., and Hu, H. (2018). Ketamine blocks bursting in the lateral habenula to rapidly relieve depression. Nature 554, 317–322.

39. Wang, D., Li, A., Dong, K., Li, H., Guo, Y., Zhang, X., Cai, M., Li, H., Zhao, G., and Yang, Q. (2021). Lateral hypothalamus orexinergic inputs to lateral habenula modulate maladaptation after social defeat stress. Neurobiol Stress 14, 100298.

40. Fan, Z., Chang, J., Liang, Y., Zhu, H., Zhang, C., Zheng, D., Wang, J., Xu, Y., Li, Q.-J., and Hu, H. (2023). Neural mechanism underlying depressive-like state associated with social status loss. Cell 186, 560–576.e17.

41. Park, H., Rhee, J., Park, K., Han, J.-S., Malinow, R., and Chung, C. (2017). Exposure to Stressors Facilitates Long-Term Synaptic Potentiation in the Lateral Habenula. J. Neurosci. 37, 6021–6030.

42. Li, B., Piriz, J., Mirrione, M., Chung, C., Proulx, C.D., Schulz, D., Henn, F., and Malinow, R. (2011). Synaptic potentiation onto habenula neurons in the learned helplessness model of depression. Nature 470, 535–539.

43. Nuno-Perez, A., Trusel, M., Lalive, A.L., Congiu, M., Gastaldo, D., Tchenio, A., Lecca, S., Soiza-Reilly, M., Bagni, C., and Mameli, M. (2021). Stress undermines reward-guided cognitive performance through synaptic depression in the lateral habenula. Neuron 109, 947–956.e5.

44. Li, K., Zhou, T., Liao, L., Yang, Z., Wong, C., Henn, F., Malinow, R., Yates, J.R., 3rd, and Hu, H. (2013). βCaMKII in lateral habenula mediates core symptoms of depression. Science 341, 1016–1020.

45. Winter, C., Vollmayr, B., Djodari-Irani, A., Klein, J., and Sartorius, A. (2011). Pharmacological inhibition of the lateral habenula improves depressive-like behavior in an animal model of treatment resistant depression. Behav. Brain Res. 216, 463–465.

46. Amat, J., Sparks, P.D., Matus-Amat, P., Griggs, J., Watkins, L.R., and Maier, S.F. (2001). The role of the habenular complex in the elevation of dorsal raphe nucleus serotonin and the changes in the behavioral responses produced by uncontrollable stress. Brain Res. 917, 118–126.

47. Tsankova, N.M., Berton, O., Renthal, W., Kumar, A., Neve, R.L., and Nestler, E.J. (2006). Sustained hippocampal chromatin regulation in a mouse model of depression and antidepressant action. Nat. Neurosci. 9, 519–525.

48. Golden, S.A., Covington, H.E., 3rd, Berton, O., and Russo, S.J. (2011). A standardized protocol for repeated social defeat stress in mice. Nat. Protoc. 6, 1183–1191.

49. McAllister, B.B., Pochakom, A., Fu, S., and Dyck, R.H. (2020). Effects of social defeat stress and fluoxetine treatment on neurogenesis and behavior in mice that lack zinc transporter 3 (ZnT3) and vesicular zinc. Hippocampus 30, 623–637.

50. Morais-Silva, G., Costa-Ferreira, W., Gomes-de-Souza, L., Pavan, J.C., Crestani, C.C., and Marin, M.T. (2019). Cardiovascular outcomes related to social defeat stress: New insights from resilient and susceptible rats. Neurobiol Stress 11, 100181.

51. Murra, D., Hilde, K.L., Fitzpatrick, A., Maras, P.M., Watson, S.J., and Akil, H. (2022). Characterizing the behavioral and neuroendocrine features of susceptibility and resilience to social stress. Neurobiol Stress 17, 100437.

52. Ayash, S., Schmitt, U., and Müller, M.B. (2020). Chronic social defeat-induced social avoidance as a proxy of stress resilience in mice involves conditioned learning. J. Psychiatr. Res. 120, 64–71.

53. Li, L., Durand-de Cuttoli, R., Aubry, A.V., Burnett, C.J., Cathomas, F., Parise, L.F., Chan, K.L., Morel, C., Yuan, C., Shimo, Y., et al. (2022). Social trauma engages lateral septum circuitry to occlude social reward. Nature. 10.1038/s41586-022-05484-5.

54. Hashikawa, Y., Hashikawa, K., Rossi, M.A., Basiri, M.L., Liu, Y., Johnston, N.L., Ahmad, O.R., and Stuber, G.D. (2020). Transcriptional and Spatial Resolution of Cell Types in the Mammalian Habenula. Neuron 106, 743–758.e5.

55. Liu, H., Rastogi, A., Narain, P., Xu, Q., Sabanovic, M., Alhammadi, A.D., Guo, L., Cao, J.-L., Zhang, H., Aqel, H., et al. (2021). Blunted diurnal firing in lateral habenula projections to dorsal raphe nucleus and delayed photoentrainment in stress-susceptible mice. PLoS Biol. 19, e3000709.

56. Pisano, T.J., Hoag, A.T., Dhanerawala, Z.M., Guariglia, S.R., Jung, C., Boele, H.-J., Seagraves, K.M., Verpeut, J.L., and Wang, S.S.-H. (2022). Automated high-throughput mouse transsynaptic viral tracing using iDISCO+ tissue clearing, light-sheet microscopy, and BrainPipe. STAR Protoc 3, 101289.

57. Renier, N., Adams, E.L., Kirst, C., Wu, Z., Azevedo, R., Kohl, J., Autry, A.E., Kadiri, L., Umadevi Venkataraju, K., Zhou, Y., et al. (2016). Mapping of Brain Activity by Automated Volume Analysis of Immediate Early Genes. Cell 165, 1789–1802.

58. Zimmerman, C.A., Pan-Vazquez, A., Wu, B., Keppler, E.F., Guthman, E.M., Fetcho, R.N., Bolkan, S.S., McMannon, B., Lee, J., Hoag, A.T., et al. (2023). A neural mechanism for learning from delayed postingestive feedback. bioRxiv, 2023.10.06.561214. 10.1101/2023.10.06.561214.

59. Wang, Q., Ding, S.-L., Li, Y., Royall, J., Feng, D., Lesnar, P., Graddis, N., Naeemi, M., Facer, B., Ho, A., et al. (2020). The Allen Mouse Brain Common Coordinate Framework: A 3D Reference Atlas. Cell 181, 936–953.e20.

60. Lee, H., and Hikosaka, O. (2022). Lateral habenula neurons signal step-by-step changes of reward prediction. iScience, 105440.

61. Matsumoto, M., and Hikosaka, O. (2009). Representation of negative motivational value in the primate lateral habenula. Nat. Neurosci. 12, 77–84.

62. Hennigan, K., D’Ardenne, K., and McClure, S.M. (2015). Distinct midbrain and habenula pathways are involved in processing aversive events in humans. J. Neurosci. 35, 198–208.

63. Li, Z.-L., Wang, Y., Zou, H.-W., Jing, X.-Y., Liu, Y.-J., and Li, L.-F. (2021). GABA(B) receptors within the lateral habenula modulate stress resilience and vulnerability in mice. Physiol. Behav. 230, 113311.

64. Ma, S., Chen, M., Jiang, Y., Xiang, X., Wang, S., Wu, Z., Li, S., Cui, Y., Wang, J., Zhu, Y., et al. (2023). Sustained antidepressant effect of ketamine through NMDAR trapping in the LHb. Nature 622, 802–809.

65. Wang, H., Li, F., Zheng, X., Meng, L., Chen, M., Hui, Y., Li, Y., Xie, K., Zhang, J., and Guo, G. (2022). Social defeat drives hyperexcitation of the piriform cortex to induce learning and memory impairment but not mood-related disorders in mice. Transl. Psychiatry 12, 380.

66. Lu, J., Gong, X., Yao, X., Guang, Y., Yang, H., Ji, R., He, Y., Zhou, W., Wang, H., Wang, W., et al. (2021). Prolonged chronic social defeat stress promotes less resilience and higher uniformity in depression-like behaviors in adult male mice. Biochem. Biophys. Res. Commun. 553, 107–113.

67. Lu, J., Zhang, Z., Yin, X., Tang, Y., Ji, R., Chen, H., Guang, Y., Gong, X., He, Y., Zhou, W., et al. (2022). An entorhinal-visual cortical circuit regulates depression-like behaviors. Mol. Psychiatry 27, 3807–3820.

68. Cui, Q.-Q., Hu, Z.-L., Hu, Y.-L., Chen, X., Wang, J., Mao, L., Lu, X.-J., Ni, M., Chen, J.-G., and Wang, F. (2020). Hippocampal CD39/ENTPD1 promotes mouse depression-like behavior through hydrolyzing extracellular ATP. EMBO Rep. 21, e47857.

69. Lorsch, Z.S., Hamilton, P.J., Ramakrishnan, A., Parise, E.M., Salery, M., Wright, W.J., Lepack, A.E., Mews, P., Issler, O., McKenzie, A., et al. (2019). Stress resilience is promoted by a Zfp189-driven transcriptional network in prefrontal cortex. Nat. Neurosci. 22, 1413–1423.

70. Amat, J., Paul, E., Watkins, L.R., and Maier, S.F. (2008). Activation of the ventral medial prefrontal cortex during an uncontrollable stressor reproduces both the immediate and long-term protective effects of behavioral control. Neuroscience 154, 1178–1186.

71. Fetcho, R.N., Hall, B.S., Estrin, D.J., Walsh, A.P., Schuette, P.J., Kaminsky, J., Singh, A., Roshgodal, J., Bavley, C.C., Nadkarni, V., et al. (2023). Regulation of social interaction in mice by a frontostriatal circuit modulated by established hierarchical relationships. Nat. Commun. 14, 2487.

72. Bagot, R.C., Parise, E.M., Peña, C.J., Zhang, H.-X., Maze, I., Chaudhury, D., Persaud, B., Cachope, R., Bolaños-Guzmán, C.A., Cheer, J.F., et al. (2015). Ventral hippocampal afferents to the nucleus accumbens regulate susceptibility to depression. Nat. Commun. 6, 7062.

73. Kumar, S., Hultman, R., Hughes, D., Michel, N., Katz, B.M., and Dzirasa, K. (2014). Prefrontal cortex reactivity underlies trait vulnerability to chronic social defeat stress. Nat. Commun. 5, 4537.

74. Li, H.-Y., Zhu, M.-Z., Yuan, X.-R., Guo, Z.-X., Pan, Y.-D., Li, Y.-Q., and Zhu, X.-H. (2023). A thalamic-primary auditory cortex circuit mediates resilience to stress. Cell 186, 1352–1368.e18.

75. Morel, C., Montgomery, S.E., Li, L., Durand-de Cuttoli, R., Teichman, E.M., Juarez, B., Tzavaras, N., Ku, S.M., Flanigan, M.E., Cai, M., et al. (2022). Midbrain projection to the basolateral amygdala encodes anxiety-like but not depression-like behaviors. Nat. Commun. 13, 1532.

76. Heshmati, M., Christoffel, D.J., LeClair, K., Cathomas, F., Golden, S.A., Aleyasin, H., Turecki, G., Friedman, A.K., Han, M.-H., Menard, C., et al. (2020). Depression and Social Defeat Stress Are Associated with Inhibitory Synaptic Changes in the Nucleus Accumbens. J. Neurosci. 40, 6228–6233.

77. Jasnow, A.M., Davis, M., and Huhman, K.L. (2004). Involvement of central amygdalar and bed nucleus of the stria terminalis corticotropin-releasing factor in behavioral responses to social defeat. Behav. Neurosci. 118, 1052–1061.

78. Markham, C.M., Norvelle, A., and Huhman, K.L. (2009). Role of the bed nucleus of the stria terminalis in the acquisition and expression of conditioned defeat in Syrian hamsters. Behav. Brain Res. 198, 69–73.

79. Colyn, L., Venzala, E., Marco, S., Perez-Otaño, I., and Tordera, R.M. (2019). Chronic social defeat stress induces sustained synaptic structural changes in the prefrontal cortex and amygdala. Behav. Brain Res. 373, 112079.

80. Zhuang, L., Gao, W., Chen, Y., Fang, W., Lo, H., Dai, X., Zhang, J., Chen, W., Ye, Q., Chen, X., et al. (2023). LHPP in glutamatergic neurons of the ventral hippocampus mediates depression-like behavior by dephosphorylating CaMKIIα and ERK. Biol. Psychiatry. 10.1016/j.biopsych.2023.08.026.

81. Bruchas, M.R., Land, B.B., Lemos, J.C., and Chavkin, C. (2009). CRF1-R activation of the dynorphin/kappa opioid system in the mouse basolateral amygdala mediates anxiety-like behavior. PLoS One 4, e8528.

82. Jhou, T.C., Fields, H.L., Baxter, M.G., Saper, C.B., and Holland, P.C. (2009). The rostromedial tegmental nucleus (RMTg), a GABAergic afferent to midbrain dopamine neurons, encodes aversive stimuli and inhibits motor responses. Neuron 61, 786–800.

83. Hong, S., Jhou, T.C., Smith, M., Saleem, K.S., and Hikosaka, O. (2011). Negative reward signals from the lateral habenula to dopamine neurons are mediated by rostromedial tegmental nucleus in primates. J. Neurosci. 31, 11457–11471.

84. Post, R.J., Bulkin, D.A., Ebitz, R.B., Lee, V., Han, K., and Warden, M.R. (2022). Tonic activity in lateral habenula neurons acts as a neutral valence brake on reward-seeking behavior. Curr. Biol. 32, 4325–4336.e5.

85. Christoph, G.R., Leonzio, R.J., and Wilcox, K.S. (1986). Stimulation of the lateral habenula inhibits dopamine-containing neurons in the substantia nigra and ventral tegmental area of the rat. J. Neurosci. 6, 613–619.

86. Ji, H., and Shepard, P.D. (2007). Lateral habenula stimulation inhibits rat midbrain dopamine neurons through a GABA(A) receptor-mediated mechanism. J. Neurosci. 27, 6923–6930.

87. van Zessen, R., Phillips, J.L., Budygin, E.A., and Stuber, G.D. (2012). Activation of VTA GABA neurons disrupts reward consumption. Neuron 73, 1184–1194.

88. Pollak Dorocic, I., Fürth, D., Xuan, Y., Johansson, Y., Pozzi, L., Silberberg, G., Carlén, M. and Meletis, K. (2014). A whole-brain atlas of inputs to serotonergic neurons of the dorsal and median raphe nuclei. Neuron 83, 663–678.

89. Ogawa, S.K., Cohen, J.Y., Hwang, D., Uchida, N., and Watabe-Uchida, M. (2014). Organization of monosynaptic inputs to the serotonin and dopamine neuromodulatory systems. Cell Rep. 8, 1105–1118.

90. Weissbourd, B., Ren, J., DeLoach, K.E., Guenthner, C.J., Miyamichi, K., and Luo, L. (2014). Presynaptic partners of dorsal raphe serotonergic and GABAergic neurons. Neuron 83, 645–662.

91. Zhou, L., Liu, M.-Z., Li, Q., Deng, J., Mu, D., and Sun, Y.-G. (2017). Organization of Functional Long-Range Circuits Controlling the Activity of Serotonergic Neurons in the Dorsal Raphe Nucleus. Cell Rep. 18, 3018–3032.

92. Herkenham, M., and Nauta, W.J. (1979). Efferent connections of the habenular nuclei in the rat. J. Comp. Neurol. 187, 19–47.

93. Behzadi, G., Kalén, P., Parvopassu, F., and Wiklund, L. (1990). Afferents to the median raphe nucleus of the rat: retrograde cholera toxin and wheat germ conjugated horseradish peroxidase tracing, and selective D-[3H]aspartate labelling of possible excitatory amino acid inputs. Neuroscience 37, 77–100.

94. Vertes, R.P., Fortin, W.J., and Crane, A.M. (1999). Projections of the median raphe nucleus in the rat. J. Comp. Neurol. 407, 555–582.

95. Challis, C., Boulden, J., Veerakumar, A., Espallergues, J., Vassoler, F.M., Pierce, R.C., Beck, S.G., and Berton, O. (2013). Raphe GABAergic neurons mediate the acquisition of avoidance after social defeat. J. Neurosci. 33, 13978–13988, 13988a.

96. Zheng, Z., Guo, C., Li, M., Yang, L., Liu, P., Zhang, X., Liu, Y., Guo, X., Cao, S., Dong, Y., et al. (2022). Hypothalamus-habenula potentiation encodes chronic stress experience and drives depression onset. Neuron 110, 1400–1415.e6.

97. Tye, K.M., Mirzabekov, J.J., Warden, M.R., Ferenczi, E.A., Tsai, H.-C., Finkelstein, J., Kim, S.-Y., Adhikari, A., Thompson, K.R., Andalman, A.S., et al. (2013). Dopamine neurons modulate neural encoding and expression of depression-related behaviour. Nature 493, 537–541.

98. Bagot, R.C., Cates, H.M., Purushothaman, I., Lorsch, Z.S., Walker, D.M., Wang, J., Huang, X., Schlüter, O.M., Maze, I., Peña, C.J., et al. (2016). Circuit-wide Transcriptional Profiling Reveals Brain Region-Specific Gene Networks Regulating Depression Susceptibility. Neuron 90, 969–983.

99. Schindelin, J., Arganda-Carreras, I., Frise, E., Kaynig, V., Longair, M., Pietzsch, T., Preibisch, S., Rueden, C., Saalfeld, S., Schmid, B., et al. (2012). Fiji: an open-source platform for biological-image analysis. Nat. Methods 9, 676–682.

100. Dombeck, D.A., Khabbaz, A.N., Collman, F., Adelman, T.L., and Tank, D.W. (2007). Imaging large-scale neural activity with cellular resolution in awake, mobile mice. Neuron 56, 43–57.

101. Friard, O., and Gamba, M. (2016). BORIS: a free, versatile open-source event-logging software for video/audio coding and live observations. Methods Ecol. Evol. 7, 1325–1330.

102. Mathis, A., Mamidanna, P., Cury, K.M., Abe, T., Murthy, V.N., Mathis, M.W., and Bethge, M. (2018). DeepLabCut: markerless pose estimation of user-defined body parts with deep learning. Nat. Neurosci. 21, 1281–1289.

103. Berman, G.J., Choi, D.M., Bialek, W., and Shaevitz, J.W. (2014). Mapping the stereotyped behaviour of freely moving fruit flies. J. R. Soc. Interface 11. 10.1098/rsif.2014.0672.

104. Pnevmatikakis, E.A., and Giovannucci, A. (2017). NoRMCorre: An online algorithm for piecewise rigid motion correction of calcium imaging data. J. Neurosci. Methods 291, 83–94.

105. Zhou, P., Resendez, S.L., Rodriguez-Romaguera, J., Jimenez, J.C., Neufeld, S.Q., Giovannucci, A., Friedrich, J., Pnevmatikakis, E.A., Stuber, G.D., Hen, R., et al. (2018). Efficient and accurate extraction of in vivo calcium signals from microendoscopic video data. Elife 7. 10.7554/eLife.28728.

106. Pnevmatikakis, E.A., Soudry, D., Gao, Y., Machado, T.A., Merel, J., Pfau, D., Reardon, T., Mu, Y., Lacefield, C., Yang, W., et al. (2016). Simultaneous Denoising, Deconvolution, and Demixing of Calcium Imaging Data. Neuron 89, 285–299.

107. Harris, K.D. (2021). Nonsense correlations in neuroscience. bioRxiv, 2020.11.29.402719. 10.1101/2020.11.29.402719.

108. Tyson, A.L., Rousseau, C.V., Niedworok, C.J., Keshavarzi, S., Tsitoura, C., Cossell, L., Strom, M., and Margrie, T.W. (2021). A deep learning algorithm for 3D cell detection in whole mouse brain image datasets. PLoS Comput. Biol. 17, e1009074.

