## Supplemental figures & tables for "Heightened lateral habenula activity during stress produces brainwide and behavioral substrates of susceptibility"

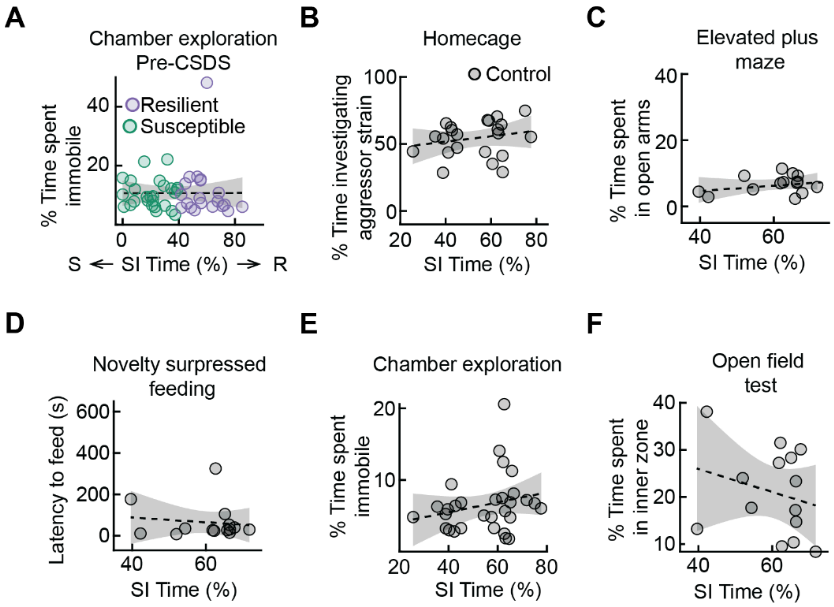


**Supplementary Figure 1. Additional data related to Figure 1. A**. Relationship between time spent immobile during chamber exploration pre-CSDS and SI time (susceptible *N* = 26 mice, resilient *N* = 20 mice). **B**. Relationship between percent of time spent investigating a juvenile from the aggressor strain in the homecage assay and aggressor strain SI time for control mice that did not undergo defeat: *R* = 0.2177, *p* = 0.3305 (*N* = 26 mice). **C**. Relationship between percent of time spent in the open arms of the elevated plus maze and aggressor strain SI time for control mice that did not undergo defeat (*N* = 14 mice). **D**. Relationship between latency to feed in the novelty suppressed feeding assay and aggressor strain SI time for control mice that did not undergo defeat (*N* = 14 mice). **E**. Relationship between time spent immobile during chamber exploration and SI time for control mice that did not undergo defeat (*N* = 30 mice). **F**. Relationship between percent of time spent in the inner zone of the open field test and aggressor strain SI time for control mice that did not undergo defeat (*N* = 14 mice). Shaded areas in **A**-**F** represent 95% confidence interval for linear fit. See Supplementary Table 1 for detailed statistics summary.

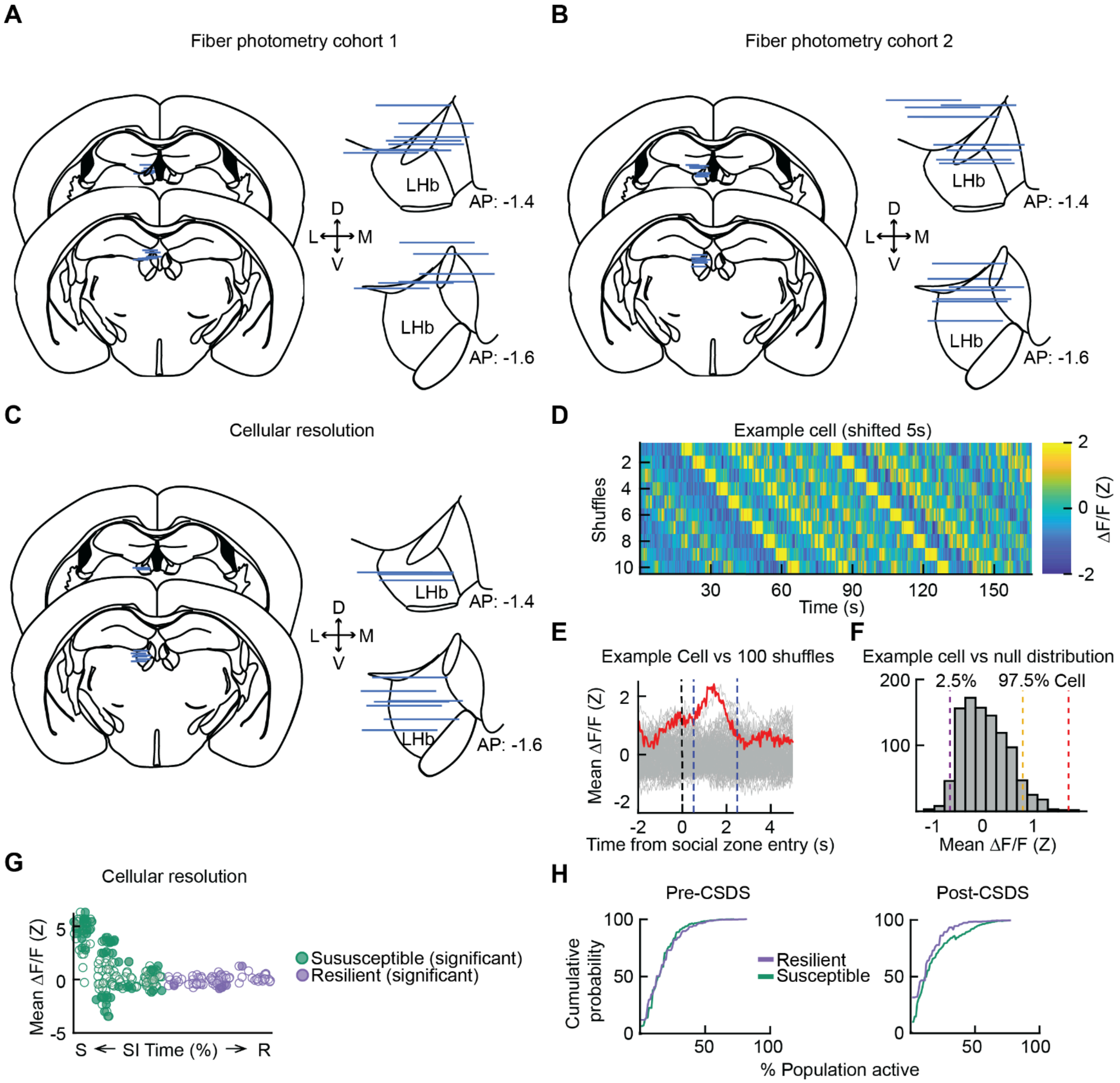


**Supplementary Figure 2. Additional data related to Figure 2. A.** Left: Histology summary of fiber tips for first photometry cohort (all plotted on left hemisphere for visualization). Right: Zoom in on LHb for fiber locations (*N* = 11 mice). **B**. Left: Histology summary of fiber tips for second photometry cohort (all plotted on left hemisphere for visualization). Right: Zoom in on LHb for fiber locations (*N* = 10 mice). **C**. Left: Histology summary of fiber tips for all cellular resolution calcium imaging mice (all plotted on left hemisphere for visualization). Right: Zoom in on LHb for lens locations (*N* = 11 mice). **D**. Fluorescence trace of an example cell shifted 5 s 10 times. **E**. Real (red) and shifted (gray) fluorescence traces of an example cell time-locked to entry into the social zone, averaging across region labeled with blue dotted lines. **F**. Histogram of the null distribution for an example cell with a dashed line at the average activity of the real trace (red) and the 2 extrema (purple and yellow) which indicate significance of $p < 0.05$ for a 2-sided test. **G**. Average neural activity of each cell plotted against avoidance magnitude with significantly activated or inhibited cells filled-in. **H**. Left: inter-cell activity synchrony pre-CSDS. Right: inter-cell activity synchrony post-CSDS (susceptible *N* = 6 mice, resilient *N* = 5 mice).

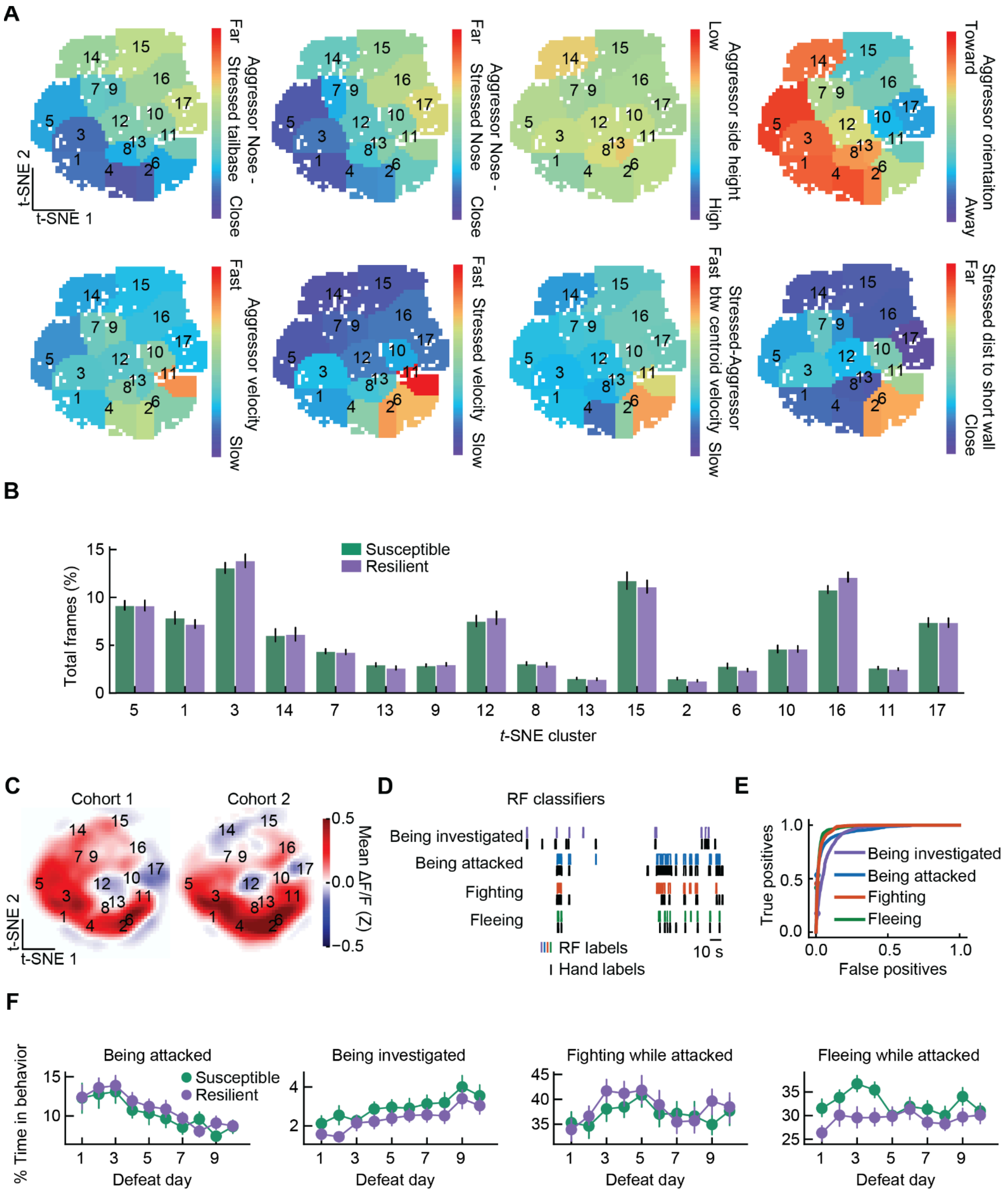


**Supplemental Figure 3. Additional data related to Figure 3. A.** Average raw feature value within each *t*-SNE cluster. **B**. Average percent of total frames in each *t*-SNE cluster for each individual, split by susceptible and resilient groups (mice from fiber photometry experiments: susceptible *N* = 21 mice, resilient *N* = 14 mice). **C**. Mean LHb GCaMP responses across *t*-SNE behavior space in defeated mice plotted separately for each cohort (mice from fiber photometry experiments: cohort 1: AAV5-CaMKII-GCaMP6f; *N* = 11 mice; cohort 2: AAV5-syn-jGCaMP7f; *N* = 10 mice). **D**. Comparison of hand-labeled behavior to performance of binary random forest (RF) classification of behaviors. **E**.

Accuracy of behavior classification. **F**. Amount of time spent in classified behaviors across days, mean±SEM plotted (mice from fiber photometry experiments and unstimulated mice from figure 6: susceptible $N = 34$ mice, resilient $N = 41$ mice). Time being attacked: effect of SI time $Z = -0.216$, $p = 0.829$, effect of day $Z = -5.665$, $p < 0.001$ , interaction, $Z = 0.061$, $p = 0.951$; Time being investigated: effect of SI time $Z = -0.885$, $p = 0.376$, effect of day $Z = 5.146$, $p < 0.001$, interaction, $Z = -0.551$, $p = 0.581$; Time fighting while attacked: effect of SI time $Z = -0.702$, $p = 0.483$, effect of day $Z = 0.294$, $p = 0.769$, interaction, $Z = -0.578$, $p = 0.563$; Time fleeing while attacked: effect of SI time $Z = -1.932$, $p = 0.053$, effect of day $Z = -0.336$, $p = 0.737$, interaction, $Z = 0.755$, $p = 0.450$. See Supplementary Tables 2-9 for more information on GEE statistics.

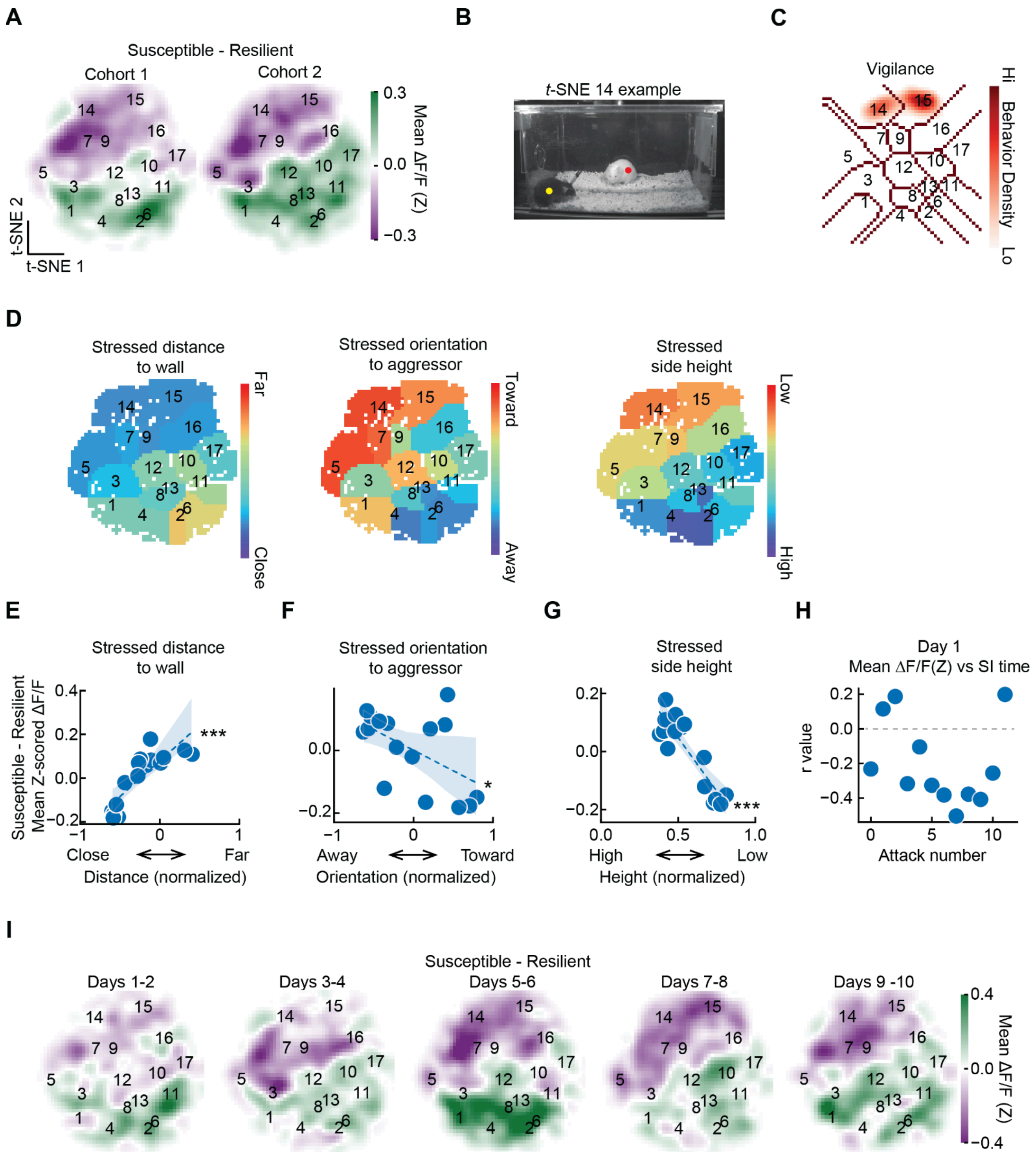


**Supplemental Figure 4. Additional data related to Figure 4. A**. Difference between susceptible and resilient maps of mean LHb GCaMP responses across *t*-SNE behavior space for each cohort (cohort 1: susceptible *N* = 6 mice, resilient *N* = 5 mice; cohort 2: susceptible *N* = 5 mice, resilient *N* = 5 mice). **B**. Example frame from cluster 14. **C**. Density of vigilance behavior annotated with supervised classifiers within *t*-SNE space. **D**. Average raw feature value within *t*-SNE clusters that have a similar pattern to the susceptible-resilient GCaMP activity map. **E**. Difference between susceptible and resilient mean LHb GCaMP of each cluster plotted against stressed mouse distance to closest wall (*N* = 17 clusters). *R* = 0.8230, *p* = 9.01-5. **F**. Difference between susceptible and resilient mean LHb GCaMP of each cluster plotted against stressed mouse orientation to aggressor (*N* = 17 clusters). *R* = 0.521, *p* = 0.0384. **G**.

Difference between susceptible and resilient mean LHb GCaMP of each cluster plotted against stressed mouse side height ($N$ = 17 clusters). $R$ = -0.8912, $p$ = 3.6E-6. **H**. Pearson's r value from correlation of the response to each attack with the SI score plotted as a function of attack number on day 1. **I**. Evolution of susceptible-resilient GCaMP activity mapped onto $t$-SNE behavior space across defeat (susceptible $N$ = 11 mice, resilient $N$ = 10 mice). $p$-values in **E**-**G** are from Pearson's correlations. Shaded areas in **E-G** represent 95% confidence interval for linear fit. *$p \leq 0.05$, **$p \leq 0.01$, ***$p \leq 0.001$. See Supplementary Table 1 for detailed statistics summary.

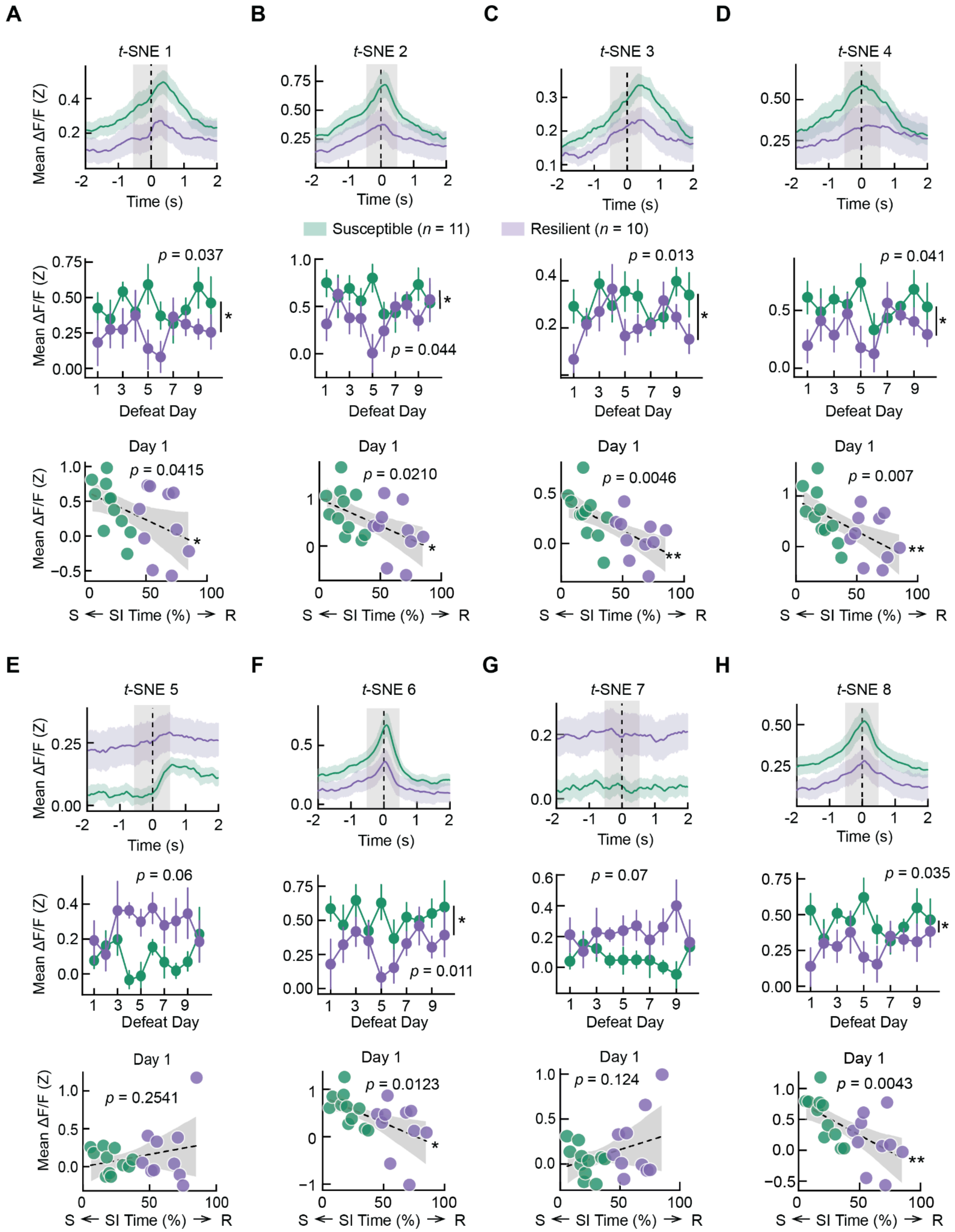


**Supplemental Figure 5. Susceptible and resilient neural activity time-locked to *t*-SNE clusters 1-8, related to Figure 4. A.** Top: LHb responses aligned to onset of *t*-SNE 1 during defeat averaged across individuals in susceptible (*N* = 11 mice) and resilient (*N* = 10 mice) groups (mean ± s.e.m. plotted). Grey region indicates +/-0.5 s surrounding onset of *t*-SNE cluster. Middle: average LHb GCaMP dF/F to onset of *t*-SNE 1 from susceptible and resilient groups across defeat (mean ± s.e.m across mice; averaging across labeled gray region in plot above(+/-0.5 s surrounding onset of *t*-SNE cluster). Onset activity by SI time, day, and their interaction: main effect of SI time, $Z$=-2.091, $p$ = 0.037; main effect of day, $Z$ = 0.470, $p$ = 0.638. Interaction, $Z$ = 0.261, $p$ = 0.794. Bottom: average LHb GCaMP responses to onset of *t*-SNE cluster (labeled gray region in top plot) on day 1 plotted against SI time for each mouse ($N$ = 21 mice): $R$ = -0.4483, $p$ = 0.0415. **B**. Top: Same as **A** for *t*-SNE 2. Middle: onset activity by SI time, day, and their interaction: main effect of SI time, $Z$ = -2.018, $p$ = 0.044; main effect of day, $Z$ = -0.366, $p$ = 0.714. Interaction, $Z$ = 1.044, $p$ = 0.296. Bottom: $R$ = -0.5, $p$ = 0.0210. **C**. Top: Same as **A** for *t*-SNE 3. Middle: onset activity by SI time, day, and their interaction: main effect of SI time, $Z$ = -2.475, $p$ = 0.013; main effect of day, $Z$ = 0.989, $p$ = 0.323. Interaction, $Z$ = -0.163, $p$ = 0.871. Bottom: $R$ = -0.5934, $p$ = 0.0046. **D**. Top: Same as **A** for *t*-SNE 4. Middle: onset activity by SI time, day, and their interaction: main effect of SI time, $Z$ = -2.045, $p$ = 0.041; main effect of day, $Z$ = 0.330, $p$ = 0.741. Interaction, $Z$ = 1.250, $p$ = 0.211. Bottom: $R$ = -0.567, $p$ = 0.0073. **E**. Top: Same as **A** for *t*-SNE 5. Middle: onset activity by SI time, day, and their interaction: main effect of SI time, $Z$ = 1.879, $p$ = 0.060; main effect of day, $Z$ = 0.610, $p$ = 0.542. Interaction, $Z$ = 0.255, $p$ = 0.799. Bottom: $R$ = 0.2605, $p$ = 0.2541. **F**. Top: Same as **A** for *t*-SNE 6. Middle: onset activity by SI time, day, and their interaction: main effect of SI time, $Z$ = -2.549, $p$ = 0.011; main effect of day, $Z$ = 0.527, $p$ = 0.598. Interaction, $Z$ = 0.760, $p$ = 0.447. Bottom: $R$ = -5.36, $p$ = 0.0123. **G**. Same as **A** for *t*-SNE 7. Middle: onset activity by SI time, day, and their interaction: main effect of SI time, $Z$ = 1.810, $p$ = 0.070; main effect of day, $Z$ = 0.238, $p$ = 0.812. Interaction, $Z$ = 1.615, $p$ = 0.106. Bottom: $R$ = 0.3464, $p$ = 0.124. **H**. Top: Same as **A** for *t*-SNE 8. Middle: onset activity by SI time, day, and their interaction: main effect of SI time, $Z$ = -2.113, $p$ = 0.035; main effect of day, $Z$ = 0.834, $p$ = 0.404. Interaction, $Z$ = 1.289, $p$ = 0.197. Bottom: $R$ = -0.596, $p$ = 0.0043. $p$-values in **A-H** (bottom) are from Pearson's correlations. Shaded areas in **A**-**H** (bottom) represent 95% confidence interval for linear fit. *$p \leq 0.05$, **$p \leq 0.01$, ***$p \leq 0.001$. in See Supplementary Tables 18-33 for more information on GEE statistics.

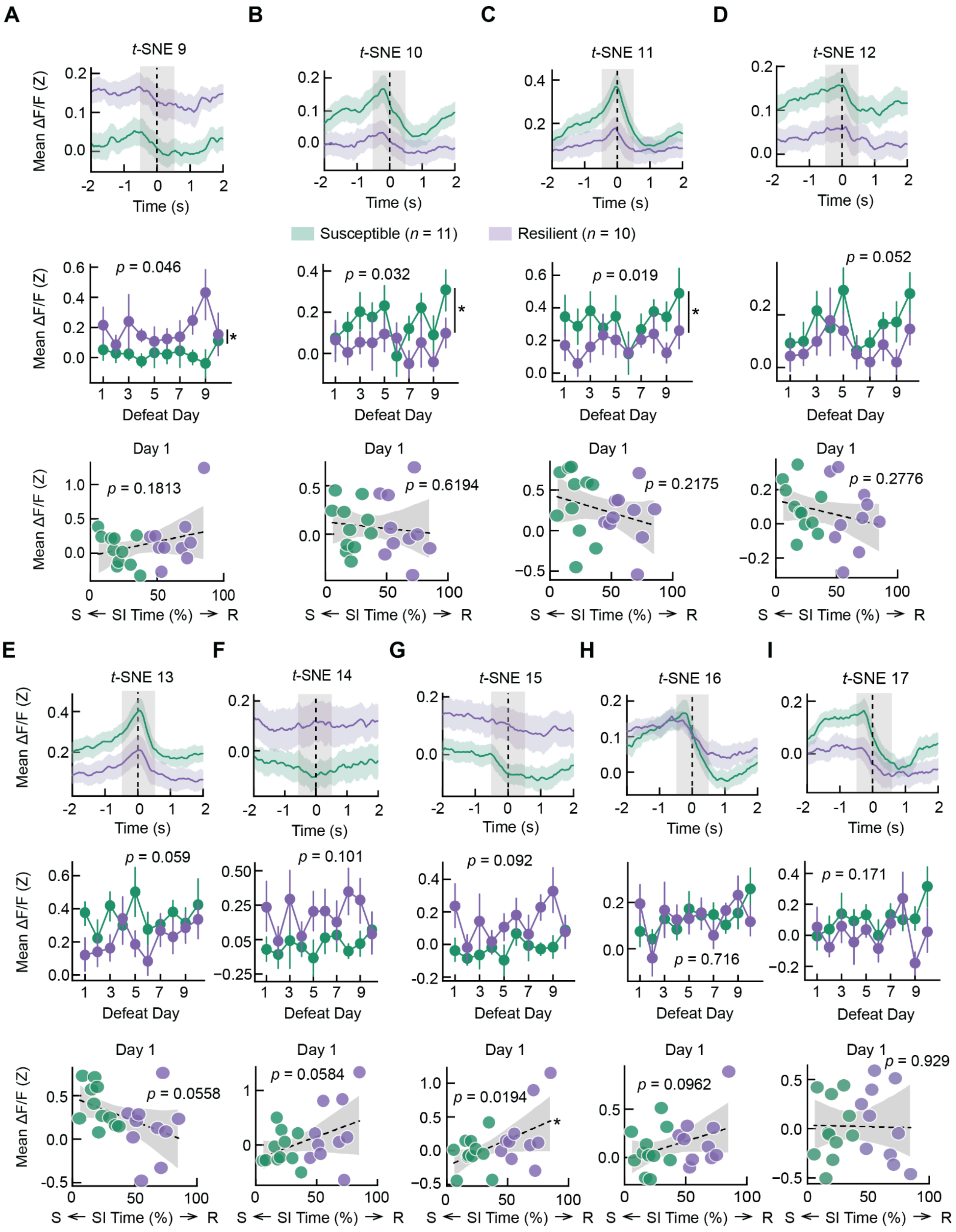


**Supplemental Figure 6. Susceptible and resilient neural activity time-locked to *t*-SNE clusters 9-17, related to Figure 4. A.** Top: LHb responses aligned to onset of *t*-SNE 9 during defeat averaged across individuals in susceptible (*N* = 11 mice) and resilient (*N* = 10 mice) groups (mean ± s.e.m. plotted). Grey region indicates +/-0.5 s surrounding onset of *t*-SNE cluster. Middle: average LHb GCaMP dF/F to onset of *t*-SNE 9 from susceptible and resilient groups across defeat (mean ± s.e.m across mice; averaging across labeled gray region in plot above(+/-0.5 s surrounding onset of *t*-SNE cluster). Onset activity by SI time, day, and their interaction: main effect of SI time, $Z = 1.997$, $p = 0.046$; main effect of day, $Z = 0.903$, $p = 0.366$. Interaction, $Z = 1.025$, $p = 0.305$. Bottom: average LHb GCaMP responses to onset of *t*-SNE cluster (labeled gray region in top plot) on day 1 plotted against SI time for each mouse ($N = 21$ mice): $R = 0.3033$, $p = 0.1813$. **B**. Top: Same as **A** for *t*-SNE 10. Middle: onset activity by SI time, day, and their interaction: main effect of SI time, $Z = -2.139$, $p = 0.032$; main effect of day, $Z = 0.616$, $p = 0.538$. Interaction, $Z = -0.654$, $p = 0.513$. Bottom: $R = -0.115$, $p = 0.6194$. **C**. Top: Same as **A** for *t*-SNE 11. Middle: onset activity by SI time, day, and their interaction: main effect of SI time, $Z = -2.354$, $p = 0.019$; main effect of day, $Z = 1.076$, $p = 0.282$. Interaction, $Z = -0.119$, $p = 0.905$. Bottom: $R = -0.281$, $p = 0.2175$. **D**. Top: Same as **A** for *t*-SNE 12. Middle: onset activity by SI time, day, and their interaction: main effect of SI time, $Z = -1.944$, $p = 0.052$; main effect of day, $Z = 1.079$, $p = 0.281$. Interaction, $Z = -0.382$, $p = 0.702$. Bottom: $R = -0.248$, $p = 0.2776$. **E**. Top: Same as **A** for *t*-SNE 13. Middle: onset activity by SI time, day, and their interaction: main effect of SI time, $Z = -1.888$, $p = 0.059$; main effect of day, $Z = 1.714$, $p = 0.087$. Interaction, $Z = 1.055$, $p = 0.292$. Bottom: $R = -0.423$, $p = 0.0558$. **F**. Top: Same as **A** for *t*-SNE 14. Middle: onset activity by SI time, day, and their interaction: main effect of SI time, $Z = 1.638$, $p = 0.101$; main effect of day, $Z = 1.509$, $p = 0.131$. Interaction, $Z = -0.452$, $p = 0.651$. Bottom: $R = 0.4194$, $p = 0.0584$. **G**. Top: Same as **A** for *t*-SNE 15. Middle: onset activity by SI time, day, and their interaction: main effect of SI time, $Z = 1.684$, $p = 0.092$; main effect of day, $Z = 2.029$, $p = 0.042$. Interaction, $Z = -0.023$, $p = 0.982$. Bottom: $R = 0.5056$, $p = 0.0194$. **H**. Top: Same as **A** for *t*-SNE 16. Middle: onset activity by SI time, day, and their interaction: main effect of SI time, $Z = 0.364$, $p = 0.716$; main effect of day, $Z = 2.119$, $p = 0.034$. Interaction, $Z = -1.121$, $p = 0.262$. Bottom: $R = 0.3726$, $p = 0.0962$. **I**. Top: Same as **A** for *t*-SNE 17. Middle: onset activity by SI time, day, and their interaction: main effect of SI time, $Z = -1.368$, $p = 0.171$; main effect of day, $Z = 1.573$, $p = 0.116$. Interaction, $Z = -2.197$, $p = 0.028$. Bottom: $R = -0.021$, $p = 0.9292$. *p*-values in **A**-**I** (bottom) are from Pearson's correlations. Shaded areas in **A**-**I** (bottom) represent 95% confidence interval for linear fit. $^*p \leq 0.05$, $^{**}p \leq 0.01$, $^{***}p \leq 0.001$. See Supplementary Tables 34-51 for more information on GEE statistics.

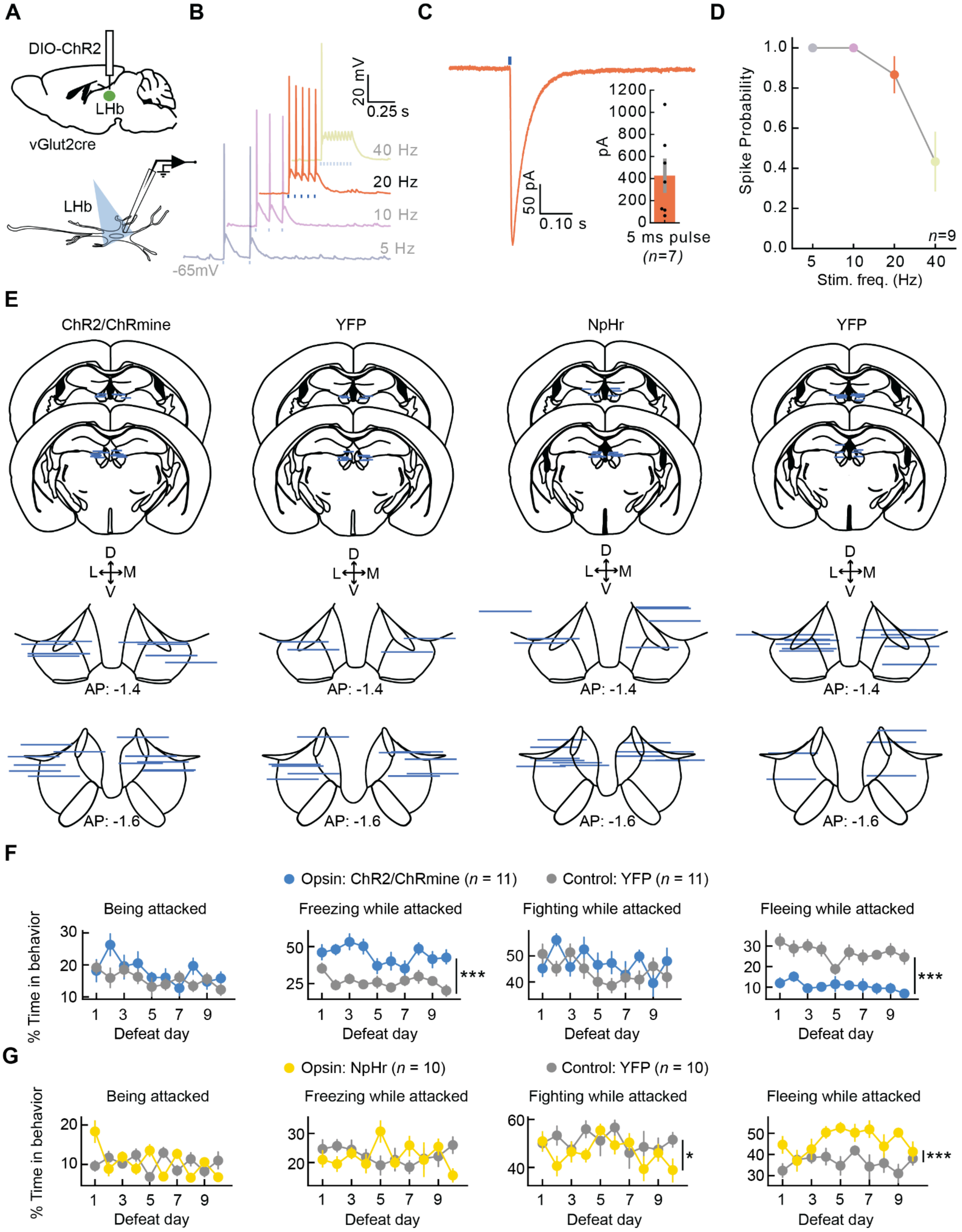


**Supplemental Figure 7. Additional data related to Figure 5. A.** Validation of optogenetic stimulation parameters through patch-clamp electrophysiology. Top: Schematic of virus injection (AAV5-EF1a-DIO-ChR2-eYFP) into LHb. Bottom: Schematic of slice recordings from neurons in LHb (5

ms, 470 nm, 8 $mW/mm^2$ light pulses). **B**. Representative current-clamp traces generated from 5 - 40 Hz optical stimulation. Highlighted trace (20 Hz, orange) is the frequency used in our in vivo experiments. **C**. Representative and average evoked photocurrents (426.8 +/- 140.2 pA). **D**. Average spike fidelity for tested stimulation frequencies (5 - 40Hz). **E**. Histology summary of implant targeting for mice expressing ChR2 or ChRmine and NpHr and respective control mice expressing YFP. **F**. Time spent in classified behaviors across days (mean ± s.e.m. plotted). Time being attacked: effect of opsin (ChR2 or ChRmine) group $Z = -1.831$, $p = 0.0687$, effect of day $Z = -1.897$, $p = 0.058$, interaction, $Z = 0.405$, $p = 0.685$; Freezing while attacked: effect of opsin (ChR2 or ChRmine) group $Z = -4.956$, $p < 0.001$, effect of day $Z = -2.109$, $p = 0.035$, interaction, $Z = -0.177$, $p = 0.859$; Fighting while attacked: effect of opsin (ChR2 or ChRmine) group $Z = -0.399$, $p = 0.690$, effect of day $Z = -0.273$, $p = 0.785$, interaction, $Z = -0.772$, $p = 0.440$; Fleeing while attacked: effect of opsin (ChR2 or ChRmine) group $Z = 8.721$, $p = <0.001$, effect of day $Z = -2.740$, $p = 0.006$, interaction, $Z = 0.111$, $p = 0.912$. **G**. Time spent in classified behaviors across days (mean ± s.e.m. plotted). Time being attacked: effect of opsin (NpHr) group $Z = 0.0357$, $p = 0.721$, effect of day $Z = -0.423$, $p = 0.672$, interaction, $Z = -2.187$, $p = 0.029$; Freezing while attacked: effect of opsin (NpHr) group $Z = -0.235$, $p = 0.814$, effect of day $Z = -0.585$, $p = 0.558$, interaction, $Z = 0.367$, $p = 0.714$; Fighting while attacked: effect of opsin (NpHr) group $Z = -2.545$, $p = 0.011$, effect of day $Z = -0.687$, $p = 0.492$, interaction, $Z = -0.436$, $p = 0.663$; Fleeing while attacked: effect of opsin (NpHr) group $Z = 3.788$, $p = <0.001$, effect of day $Z = -0.213$, $p = 0.831$, interaction, $Z = 0.803$, $p = 0.422$. $p$-values in **F**-**G** are from two-sided GEE. *$p \leq 0.05$, **$p \leq 0.01$, ***$p \leq 0.001$. See Supplementary Tables 52-59 for more information on GEE statistics.

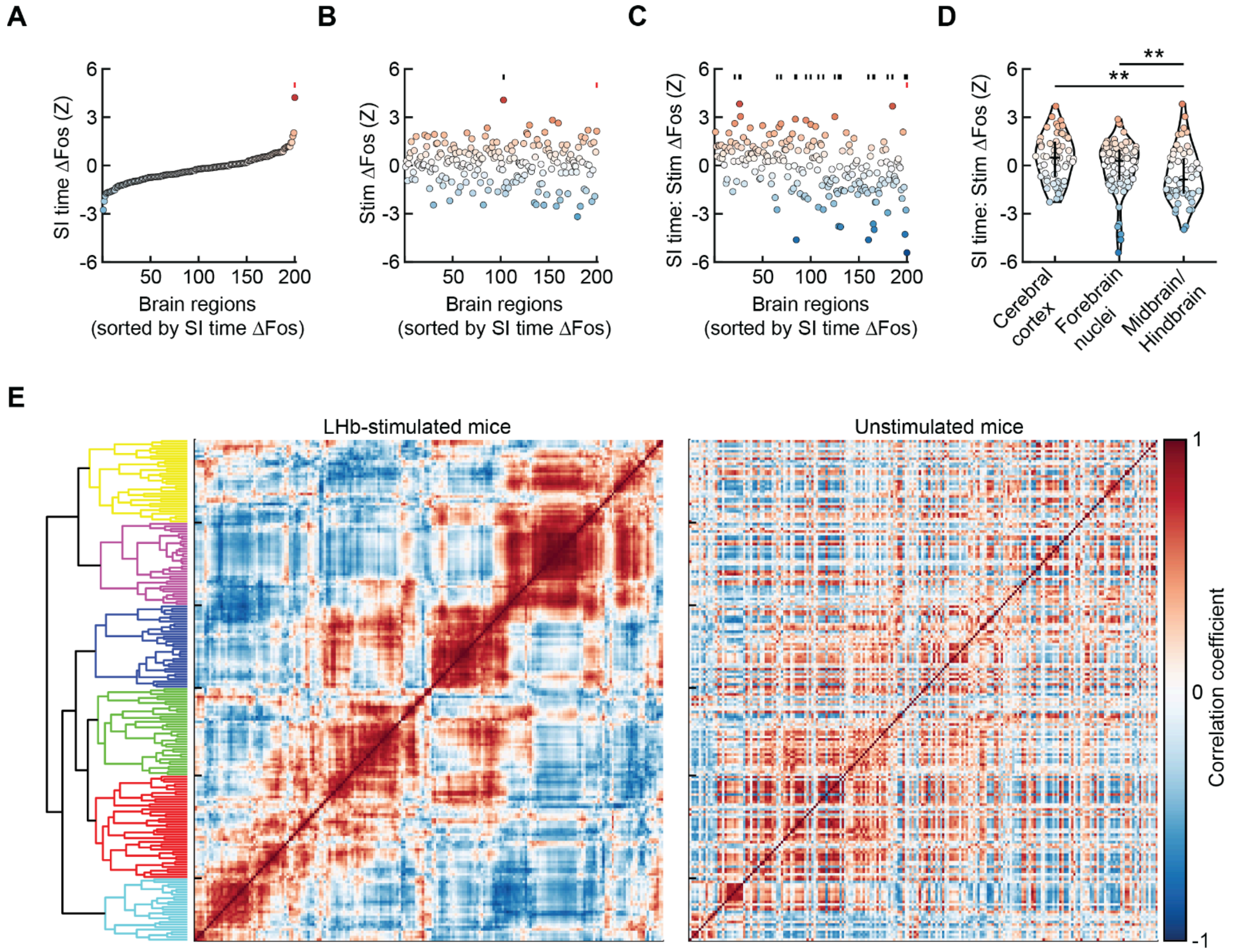


**Supplemental Figure 8. Additional analysis related to Figure 6. A–B.** Summary of coefficient estimates from the GLMM fit to all 54 mice in the Fos dataset, where for each brain region: *Counts* ~ *SI Time* + *Stim* + *SI Time:Stim* + ln(*Total Counts*) + (1+*SI time*|*Cohort*). **A.** Individual brain regions sorted by the *SI Time* coefficient estimate; the *SI Time* coefficient captures the contribution of SI time to $Fos^{+}$ cell counts for the unstimulated mice. Significance is denoted with red ticks across **A**-**C**. **B.** *Stim* coefficient estimates, shown using the sorting from **A**. Significance is highlighted by black ticks. **C.** *SI Time:Stim* [interaction term] coefficient estimates, shown using the sorting from **A**; the *SI Time:Stim* interaction coefficient captures the contribution of SI time to $Fos^{+}$ cell counts for the LHb-stimulated mice. Significance is highlighted by black ticks. **D**. Comparison of distributions of *SI Time:Stim* interaction coefficients (from **C**) across all brain regions in cerebral cortex ($n$ = 61 regions), forebrain nuclei ($n$ = 83 regions), and midbrain/hindbrain ($n$ = 56 regions). **E.** Correlation matrices showing the animal-by-animal pairwise Fos correlation for every pair of brain regions for LHb-stimulated mice (left) and unstimulated control mice (right). Each row/column represents one brain region, and regions are sorted by hierarchical clustering of the correlation matrix of the LHb-stimulated mice. Significance in **A**-**C** is based on GLMM coefficient estimate *z*-tests corrected for 10% false discovery rate. Error bars in **D** represent median ± interquartile. *p*-values in **D** are from Kolmogorov-Smirnov tests with Hochberg-Bonferroni correction for multiple comparisons. See Supplementary Tables 1 and 63-65 for detailed statistics summary. See Supplementary Table 66 for a list of brain regions in each cluster for **E**. *$p \leq 0.05$, **$p \leq 0.01$, ***$p \leq 0.001$.

### Supplemental Tables

**Supplementary Data Table 1**: **Statistics summary**
Note: all t-tests and correlations are 2-sided unless noted otherwise. In "Significance" column: *p<0.05, **p<0.01, ***p<0.001, NS not significant.

| Figure | Description | Sample size | Statistical test | Test Statistic | *p* Value | Significance |
|---|---|---|---|---|---|---|
| Figure 1C | Aggressor strain SI time in control mice vs stressed mice pre-CSDS | 22 mice, 46 mice | t-test | t = 0.7341 | 0.4655 | NS |
| Figure 1C | Aggressor strain SI time in control mice vs stressed mice post-CSDS | 26 mice, 66 mice | t-test | t = 3.5505 | 6.2E-4 | *** |
| Figure 1D | Self strain SI time in susceptible mice pre-CSDS vs post-CSDS | 26 mice, 26 mice | Paired t-test (with Bonferroni correction) | t = -6.1047 | 1.3E-05 | *** |
| Figure 1D | Self strain SI time in resilient mice pre-CSDS vs post-CSDS | 20 mice, 20 mice | Paired t-test (with Bonferroni correction) | t = -2.9147 | 0.0393 | * |
| Figure 1D | Other strain SI time in susceptible mice pre-CSDS vs post-CSDS | 14 mice, 14 mice | Paired t-test (with Bonferroni correction) | t = -3.1374 | 0.0393 | * |
| Figure 1E | Investigation time difference across stress groups to aggressor strain in homecage assay | 68 mice, 3 groups | 1-way ANOVA | $F_{(65,2)} = 5$ | 0.0096 | ** |
| Figure 1EC | Investigation time difference across control and susceptible to aggressor strain in homecage assay | 22 mice, 26 mice | t-test | t = 3.2324 | 0.0023 | ** |
| Figure 1E | Investigation time difference across susceptible and resilient to aggressor strain in homecage assay | 26 mice, 20 mice | t-test | t = -2.1898 | 0.0339 | * |
| Figure 1E | Investigation time difference across control and resilient to aggressor strain in homecage assay | 22 mice, 20 mice | t-test | t = 0.3753 | 0.7095 | NS |

| | | | | | | |
|---|---|---|---|---|---|---|
| Figure 1E | Investigation time difference across stress groups to self strain in homecage assay | 67 mice, 3 groups | 1-way ANOVA | $F_{(64,2)}$ = 3.16 | 0.0491 | * |
| Figure 1EC | Investigation time difference across control and susceptible to self strain in homecage assay | 22 mice, 26 mice | t-test | t = -2.1285 | 0.0387 | * |
| Figure 1E | Investigation time difference across susceptible and resilient to self strain in homecage assay | 26 mice, 20 mice | t-test | t = 2.0548 | 0.0460 | * |
| Figure 1E | Investigation time difference across control and resilient to self strain in homecage assay | 19 mice, 22 mice | t-test | t = 0.1218 | 0.9037 | NS |
| Figure 1F | Time spent investigating aggressor strain juvenile in homecage vs SI time | 46 mice | Pearson's correlation | R = 0.3965 | 0.0064 | ** |
| Figure 1G | Open arm time difference across stress groups | 65 mice, 3 groups | 1-way ANOVA | $F_{(62,2)}$ = 3.99 | 0.0234 | * |
| Figure 1G | Open arm time difference across control vs susceptible | 14 mice, 20 mice | t-test | t = 4.5352 | 7.6E-5 | *** |
| Figure 1G | Open arm time difference across susceptible vs resilient | 20 mice, 31 mice | t-test | t = -2.1630 | 0.038 | * |
| Figure 1G | Open arm time difference across control vs resilient | 14 mice, 31 mice | t-test | t = 0.5717 | 0.5705 | NS |
| Figure 1H | Open arm time vs SI time | 51 mice | Pearson's correlation | R = 0.3100 | 0.0269 | * |
| Figure 1H | Latency to feed difference across stress groups | 64 mice, 3 groups | 1-way ANOVA | $F_{(61,2)}$ = 3.86 | 0.0263 | * |
| Figure 1I | Latency to feed difference across control vs susceptible | 14 mice, 19 mice | t-test | t = -2.7623 | 0.0096 | ** |
| Figure 1I | Latency to feed difference across susceptible vs resilient | 19 mice, 31 mice | t-test | t = 1.6294 | 0.1098 | NS |

| | | | | | | |
|---|---|---|---|---|---|---|
| Figure 1I | Latency to feed difference across control vs resilient | 14 mice, 31 mice | t-test | t = -1.5607 | 0.1259 | NS |
| Figure 1J | Latency to feed vs SI time | 50 mice | Pearson's correlation | R = -0.4584 | 0.0008 | *** |
| Figure 1K | Immobility difference across stress groups | 116 mice, 3 groups | 1-way ANOVA | $F_{(113,2)}$ = 9.72 | 0.0001 | * |
| Figure 1K | Immobility difference across control vs susceptible | 30 mice, 40 mice | t-test | t = -3.7733 | 3.4E-4 | *** |
| Figure 1K | Immobility difference across susceptible vs resilient | 40 mice, 46 mice | t-test | t = -2.5642 | 0.0102 | * |
| Figure 1K | Immobility difference across control vs resilient | 30 mice, 46 mice | t-test | t = -2.7313 | 0.0079 | ** |
| Figure 1L | Immobility vs SI time | 86 mice | Pearson's correlation | R = -0.3119 | 0.0035 | ** |
| Figure 1L | Inner zone time difference across control vs resilient | 65 mice | 1-way ANOVA | $F_{(62,2)}$ = | 0.0036 | ** |
| Figure 1M | Inner zone time difference across control vs susceptible | 14 mice, 20 mice | t-test | t = 3.5396 | 0.0013 | ** |
| Figure 1M | Inner zone time difference across susceptible vs resilient | 20 mice, 31 mice | t-test | t = -1.0845 | 0.2835 | NS |
| Figure 1M | Inner zone time difference across control vs resilient | 14 mice, 31 mice | t-test | t = 2.6244 | 0.0120 | * |
| Figure 1N | Inner zone time vs SI time | 51 mice | Pearson's correlation | R = 0.2008 | 0.1578 | NS |
| Figure 2I | Magnitude of the fluorescence response during the SI test with the aggressor strain vs avoidance level (in mice from fiber photometry experiments) | 20 mice | Pearson's correlation | R = 0.7291 | 0.0003 | *** |
| Figure 2F | Susceptible mice LHb(GCaMP) at social | 10 mice, 10 mice | Paired t-test (with | t = -3.7842 | 0.0389 | * |

| | | | | | | |
|---|---|---|---|---|---|---|
| | zone entry pre-CSDS vs post-CSDS | | Bonferroni correction) | | | |
| Figure 2J | Susceptible mice LHb(GCaMP) at first social zone entry pre-CSDS vs post-CSDS | 10 mice, 10 mice | t-test | t = -2.0530 | 0.0549 | NS |
| Figure 2J | Susceptible mice LHb(GCaMP) at second social zone entry pre-CSDS vs post-CSDS | 10 mice, 8 mice | t-test | t = -2.4735 | 0.0250 | * |
| Figure 2J | Susceptible mice LHb(GCaMP) at third social zone entry pre-CSDS vs post-CSDS | 10 mice, 8 mice | t-test | t = -1.6412 | 0.1203 | NS |
| Figure 2J | Susceptible mice LHb(GCaMP) at fourth social zone entry pre-CSDS vs post-CSDS | 10 mice, 6 mice | t-test | t = -2.4105 | 0.0303 | * |
| Figure 2J | Susceptible mice LHb(GCaMP) at fifth social zone entry pre-CSDS vs post-CSDS | 7 mice, 5 mice | t-test | t = -3.8559 | 0.0032 | ** |
| Figure 2J | Susceptible mice LHb(GCaMP) at sixth social zone entry pre-CSDS vs post-CSDS | 6 mice, 3 mice | t-test | t = -1.5090 | 0.1750 | NS |
| Figure 2Q | Susceptible mice LHb(GCaMP) at social zone entry pre-CSDS vs post-CSDS | 165 neurons from 6 mice, 131 neurons from 4 mice | t-test (with Bonferroni correction) | t = -5.8304 | <0.0001 | *** |
| Figure 2U | Distribution of responses during the SI test | 131 neurons from 4 mice, 75 neurons from 5 mice | Kolmogorov-Smirnov test | ks = 0.4123 | 9.5E-8 | ** |
| Figure 2W | Magnitude of the fluorescence response during the SI test with the aggressor strain vs | 7 mice | Pearson’s correlation | R = -0.7732 | 0.0415 | * |

| | | | | | | |
|---|---|---|---|---|---|---|
| | avoidance level (in significantly activated cells) | | | | | |
| Figure 2X | Magnitude of the fluorescence response during the SI test with the aggressor strain vs avoidance level (in significantly inhibited cells) | 7 mice | Pearson's correlation | R = -0.4015 | 0.3720 | NS |
| Figure 2Y | Pre-CSDS Spontaneous transient rates | 165 neurons from 6 mice, 71 neurons from 5 mice | t-test | t = -0.1427 | 0.8867 | NS |
| Figure 2Z | Post-CSDS Spontaneous transient rates | 182 neurons from 6 mice, 75 neurons from 5 mice | t-test | t = -3.0187 | 0.0028 | *** |
| Figure 3F | LHb(GCaMP) mean of each cluster vs mean distance between mice of each cluster | 17 clusters | Pearson's correlation | R = -0.8215 | 5.3E-5 | *** |
| Figure 4L | LHb(GCaMP) at attack onset on day 1 vs SI time | 21 mice | Pearson's correlation | R = -0.5783 | 0.0060 | ** |
| Figure 4M | LHb(GCaMP) at fighting onset on day 1 vs SI time | 21 mice | Pearson's correlation | R = -0.4806 | 0.0274 | * |
| Figure 4N | LHb(GCaMP) at fleeing onset on day 1 vs SI time | 21 mice | Pearson's correlation | R = -0.5807 | 0.0058 | ** |
| Figure 4O | LHb(GCaMP) at vigilance onset on day 1 vs SI time | 21 mice | Pearson's correlation | R = 0.4954 | 0.0224 | * |
| Figure 5G | SI time difference across opsin (ChR2 or ChRmine) vs YFP | 11 mice, 11 mice | t-test | t = -2.1263 | 0.0461 | * |
| Figure 5H | Open arm time difference across opsin (ChR2 or ChRmine) vs YFP | 11 mice, 11 mice | t-test | t = -2.2934 | 0.0328 | * |

| | | | | | | |
|---|---|---|---|---|---|---|
| Figure 5I | Inner zone time difference across opsin (ChR2 or ChRmine) vs YFP | 11 mice, 11 mice | t-test | t = -2.8231 | 0.0105 | * |
| Figure 5J | Latency to feed difference across opsin (ChR2 or ChRmine) vs YFP | 11 mice, 11 mice | t-test | t = 0.7336 | 0.4717 | NS |
| Figure 5K | SI time difference across opsin (NpHr) vs YFP | 10 mice, 10 mice | t-test | t = 1.1623 | 0.2603 | NS |
| Figure 5L | Open arm time difference across opsin (NpHr) vs YFP | 10 mice, 10 mice | t-test | t = -3.9140 | 0.0010 | ** |
| Figure 5M | Inner zone time difference across opsin (NpHr) vs YFP | 10 mice, 10 mice | t-test | t = 1.4660 | 0.1599 | NS |
| Figure 5N | Latency to feed difference across opsin (NpHr) vs YFP | 10 mice, 10 mice | t-test | t = -0.3913 | 0.7002 | NS |
| Figure 6H | Difference in SI time coefficient distributions (LHb-stimulated mice only), cerebral cortex vs. forebrain nuclei | 61 regions, 83 regions | Kolmogorov-Smirnov test | ks = 0.32 | 0.0025 | ** |
| Figure 6H | Difference in SI time coefficient distributions (LHb-stimulated mice only), cerebral cortex vs. midbrain/hindbrain | 61 regions, 56 regions | Kolmogorov-Smirnov test | ks = 0.40 | 0.00038 | *** |
| Figure 6H | Difference in SI time coefficient distributions (LHb-stimulated mice only), forebrain nuclei vs. midbrain/hindbrain | 83 regions, 56 regions | Kolmogorov-Smirnov test | ks = 0.27 | 0.012 | * |
| Figure 6I | Estimated SI time coefficient (LHb-stimulated mice only) vs. estimated LHb stimulation coefficient (all mice) | 200 regions | Pearson correlation | r = 0.468 | 2.8E-12 | *** |
| Figure 6J | Estimated SI time coefficient (unstimulated mice only) vs. estimated LHb stimulation coefficient (all mice) | 200 regions | Pearson correlation | r = -0.0752 | 0.29 | NS |

| | | | | | | |
|---|---|---|---|---|---|---|
| Supp. Figure 1A | Pre-CSDS immobility vs SI time | 46 mice | Pearson's correlation | R = 0.0058 | 0.9695 | NS |
| Supp. Figure 1B | Time spent investigating aggressor strain juvenile in homecage vs SI time for control mice | 26 mice | Pearson's correlation | R = 0.2177 | 0.3305 | NS |
| Supp. Figure 1C | Open arm time vs SI time for control mice | 14 mice | Pearson's correlation | R = 0.2902 | 0.3143 | NS |
| Supp. Figure 1D | Latency to feed vs SI time for control mice | 14 mice | Pearson's correlation | R = -0.1287 | 0.6611 | NS |
| Supp. Figure 1E | Immobility vs SI time for control mice | 30 mice | Pearson's correlation | R = 0.2322 | 0.2169 | NS |
| Supp. Figure 1F | Inner zone time vs SI time for control mice | 14 mice | Pearson's correlation | R = -0.2542 | 0.3806 | NS |
| Supp. Figure 4F | Difference in LHb(GCaMP) mean of each cluster vs mean distance from closest wall of each cluster | 17 clusters | Pearson's correlation | R = 0.8230 | 9.01-5 | *** |
| Supp. Figure 4G | Difference in LHb(GCaMP) mean of each cluster vs mean stressed mouse orientation to aggressor mouse of each cluster | 17 clusters | Pearson's correlation | R = -0.521 | 0.0384 | * |
| Supp. Figure 4H | Difference in LHb(GCaMP) mean of each cluster vs mean stressed mouse side height of each cluster | 17 clusters | Pearson's correlation | R = -0.8912 | 3.6E-6 | *** |
| Supp. Figure 5A | LHb(GCaMP) at *t*-SNE 1 onset on day 1 vs SI time | 21 mice | Pearson's correlation | R = -0.4483 | 0.04151 | * |
| Supp. Figure 5B | LHb(GCaMP) at *t*-SNE 2 onset on day 1 vs SI time | 21 mice | Pearson's correlation | R = -0.5001 | 0.0210 | * |

| | | | | | | |
|---|---|---|---|---|---|---|
| Supp. Figure 5C | LHb(GCaMP) at *t*-SNE 3 onset on day 1 vs SI time | 21 mice | Pearson's correlation | R = -0.5934 | 0.0046 | ** |
| Supp. Figure 5D | LHb(GCaMP) at *t*-SNE 4 onset on day 1 vs SI time | 21 mice | Pearson's correlation | R = -0.5674 | 0.0073 | ** |
| Supp. Figure 5E | LHb(GCaMP) at *t*-SNE 5 onset on day 1 vs SI time | 21 mice | Pearson's correlation | R = 0.2605 | 0.2541 | NS |
| Supp. Figure 5F | LHb(GCaMP) at *t*-SNE 6 onset on day 1 vs SI time | 21 mice | Pearson's correlation | R = -0.5359 | 0.01228 | * |
| Supp. Figure 5G | LHb(GCaMP) at *t*-SNE 7 onset on day 1 vs SI time | 21 mice | Pearson's correlation | R = 0.3464 | 0.1240 | NS |
| Supp. Figure 5H | LHb(GCaMP) at *t*-SNE 8 onset on day 1 vs SI time | 21 mice | Pearson's correlation | R = -0.5961 | 0.0043 | ** |
| Supp. Figure 6A | LHb(GCaMP) at *t*-SNE 9 onset on day 1 vs SI time | 21 mice | Pearson's correlation | R = 0.3033 | 0.1813 | NS |
| Supp. Figure 6B | LHb(GCaMP) at *t*-SNE 10 onset on day 1 vs SI time | 21 mice | Pearson's correlation | R = -0.1151 | 0.6194 | NS |
| Supp. Figure 6C | LHb(GCaMP) at *t*-SNE 11 onset on day 1 vs SI time | 21 mice | Pearson's correlation | R = -0.2808 | 0.2175 | NS |
| Supp. Figure 6D | LHb(GCaMP) at *t*-SNE 12 onset on day 1 vs SI time | 21 mice | Pearson's correlation | R = -0.2484 | 0.2776 | NS |
| Supp. Figure 6E | LHb(GCaMP) at *t*-SNE 13 onset on day 1 vs SI time | 21 mice | Pearson's correlation | R = -0.4234 | 0.0558 | NS |
| Supp. Figure 6F | LHb(GCaMP) at *t*-SNE 14 onset on day 1 vs SI time | 21 mice | Pearson's correlation | R = 0.4194 | 0.0584 | NS |
| Supp. Figure 6G | LHb(GCaMP) at *t*-SNE 15 onset on day 1 vs SI time | 21 mice | Pearson's correlation | R = 0.5056 | 0.0194 | * |
| Supp. Figure 6H | LHb(GCaMP) at *t*-SNE 16 onset on day 1 vs SI time | 21 mice | Pearson's correlation | R = 0.3726 | 0.0962 | NS |

| | | | | | | |
|---|---|---|---|---|---|---|
| Supp. Figure 6I | LHb(GCaMP) at *t*-SNE 17 onset on day 1 vs SI time | 21 mice | Pearson's correlation | R = -0.0207 | 0.9292 | NS |
| Supp. Figure 8D | Difference in *SI time:Stim* interaction coefficient distributions (LHb-stimulated mice only), cerebral cortex vs. forebrain nuclei | 61 regions, 83 regions | Kolmogorov-Smirnov test | ks = 0.19 | 0.14 | NS |
| Supp. Figure 8D | Difference in *SI time:Stim* interaction coefficient distributions (LHb-stimulated mice only), cerebral cortex vs. midbrain/hindbrain | 61 regions, 56 regions | Kolmogorov-Smirnov test | ks = 0.35 | 0.0037 | ** |
| Supp. Figure 8D | Difference in *SI time:Stim* interaction coefficient distributions (LHb-stimulated mice only), forebrain nuclei vs. midbrain/hindbrain | 83 regions, 56 regions | Kolmogorov-Smirnov test | ks = 0.31 | 0.0051 | ** |

**Supplementary Data Table 2:** 2-sided GEE regression of behavior (**Figure S3F**, first panel): % time mice are being attacked = defeat day + SI time + defeat day*SI time + intercept, grouped by mouse. Number of mice (groups) = 75, minimum samples per group 33, maximum samples per group 42, dependence structure = independence, family = Gaussian. NOTE: defeat day is coded as mean centered

| *Model: % time mice are **being attacked** = defeat day + SI time + defeat day*SI time category + intercept* | $\beta$**±standard error** | **z-stat** | **p-value** | **95% CI [lower, upper]** | |
|---|---|---|---|---|---|
| **Intercept** | 10.670±0.465 | 22.954 | <0.001 | 9.759 | 11.581 |
| **Defeat day** | -0.5956±0.105 | -5.665 | <0.001 | -0.802 | -0.390 |
| **SI time** | -0.0062±0.029 | -0.216 | 0.829 | -0.063 | 0.050 |
| **Defeat day * SI time** | 0.0003±0.005 | 0.061 | 0.951 | -0.010 | 0.011 |

**Supplementary Data Table 3:** 2-sided GEE regression of behavior (**Figure S3F**, first panel): % time mice are being attacked = defeat day + susceptibility category + defeat day*susceptibility category + intercept, grouped by mouse. Number of mice (groups) = 75, minimum samples per group 33, maximum samples per group 42, dependence structure = independence, family = Gaussian. NOTE: defeat day is coded as mean centered

| *Model: % time mice are* ***being attacked*** *= defeat day* + susceptibility *category + defeat day**susceptibility *category + intercept* | $\beta$±standard error | z-stat | p-value | 95% CI [lower, upper] | |
|---|---|---|---|---|---|
| **Intercept** | 10.3152±0.777 | 13.278 | <0.001 | 8.793 | 11.838 |
| **Defeat day** | -0.5771±0.176 | -3.283 | 0.001 | -0.922 | -0.233 |
| **Susceptibility category** | 0.6335±0.957 | 0.662 | 0.508 | -1.243 | 2.510 |
| **Defeat day * Susceptibility category** | -0.0330±0.217 | -0.152 | 0.879 | -0.458 | 0.392 |

**Supplementary Data Table 4:** 2-sided GEE regression of behavior (**Figure S3F**, second panel): % time mice are being investigated = defeat day + SI time + defeat day*SI time + intercept, grouped by mouse. Number of mice (groups) = 75, minimum samples per group 33, maximum samples per group 42, dependence structure = independence, family = Gaussian. NOTE: defeat day is coded as mean centered

| *Model: % time mice are* ***being investigated*** *= defeat day + SI time + defeat day*SI time category + intercept* | $\beta$±standard error | z-stat | p-value | 95% CI [lower, upper] | |
|---|---|---|---|---|---|
| **Intercept** | 2.6339±0.145 | 18.221 | <0.001 | 2.351 | 2.917 |
| **Defeat day** | 0.1776±0.035 | 5.146 | <0.001 | 0.110 | 0.245 |
| **SI time** | -0.0069±0.008 | -0.885 | 0.376 | -0.022 | 0.008 |
| **Defeat day * SI time category** | -0.0012±0.002 | -0.551 | 0.581 | -0.005 | 0.003 |

**Supplementary Data Table 5:** 2-sided GEE regression of behavior (**Figure S3F**, second panel): % time mice are being investigated = defeat day + susceptibility category + defeat day*susceptibility category + intercept, grouped by mouse. Number of mice (groups) = 75, minimum samples per group 33, maximum samples per group 42, dependence structure = independence, family = Gaussian. NOTE: defeat day is coded as mean centered

| *Model: % time mice are* ***being investigated*** *= defeat day +* susceptibility *category + defeat day**susceptibility *category + intercept* | $\beta$±standard error | z-stat | p-value | 95% CI [lower, upper] | |
|---|---|---|---|---|---|
| **Intercept** | 2.9541±0.241 | 12.251 | <0.001 | 2.482 | 3.427 |
| **Defeat day** | 0.1727±0.055 | 3.154 | 0.002 | 0.065 | 0.280 |
| **Susceptibility category** | -0.5719±0.293 | -1.950 | 0.051 | -1.147 | 0.003 |

| | | | | | |
|---|---|---|---|---|---|
| **Defeat day * Susceptibility category** | 0.0088±0.070 | 0.124 | 0.901 | -0.129 | 0.147 |

**Supplementary Data Table 6:** 2-sided GEE regression of behavior (**Figure S3F**, third panel): % time mice are fighting while attacked = defeat day + SI time + defeat day*SI time + intercept, grouped by mouse. Number of mice (groups) = 75, minimum samples per group 33, maximum samples per group 42, dependence structure = independence, family = Gaussian. NOTE: defeat day is coded as mean centered

| *Model: % time mice are **fighting while attacked** = defeat day + SI time + defeat day*SI time category + intercept* | $\beta$**±standard error** | **z-stat** | **p-value** | **95% CI [lower, upper]** | |
|---|---|---|---|---|---|
| **Intercept** | 37.709±1.035 | 36.517 | <0.001 | 35.762 | 39.718 |
| **Defeat day** | 0.0638±0.217 | 0.294 | 0.769 | -0.362 | 0.490 |
| **SI time** | -0.0353±0.050 | -0.702 | 0.483 | -0.134 | 0.063 |
| **Defeat day * SI time** | -0.0075±0.013 | -0.578 | 0.563 | -0.033 | 0.018 |

**Supplementary Data Table 7:** 2-sided GEE regression of behavior (**Figure S3F**, third panel): % time mice are fighting while attacked = defeat day + susceptibility category + defeat day*susceptibility category + intercept, grouped by mouse. Number of mice (groups) = 75, minimum samples per group 33, maximum samples per group 42, dependence structure = independence, family = Gaussian. NOTE: defeat day is coded as mean centered

| *Model: % time mice are **fighting while attacked** = defeat day +* susceptibility *category + defeat day**susceptibility *category + intercept* | $\beta$**±standard error** | **z-stat** | **p-value** | **95% CI [lower, upper]** | |
|---|---|---|---|---|---|
| **Intercept** | 37.0788±1.283 | 28.895 | <0.001 | 34.564 | 39.594 |
| **Defeat day** | 0.0514±0.263 | 0.196 | 0.845 | -0.463 | 0.566 |
| **Susceptibility category** | 1.2704±2.011 | 0.632 | 0.528 | -2.671 | 5.211 |
| **Defeat day * Susceptibility category** | 0.0221±0.421 | 0.052 | 0.958 | -0.804 | 0.848 |

**Supplementary Data Table 8:** 2-sided GEE regression of behavior (**Figure S3F**, fourth panel): % time mice are fleeing while attacked = defeat day + SI time + defeat day*SI time + intercept, grouped by mouse. Number of mice (groups) = 75, minimum samples per group 33, maximum samples per group 42, dependence structure = independence, family = Gaussian. NOTE: defeat day is coded as mean centered

| *Model: % time mice are **fleeing while attacked** = defeat day + SI time + defeat day*SI time category + intercept* | **$\beta$±standard error** | **z-stat** | **p-value** | **95% CI [lower, upper]** | |
|---|---|---|---|---|---|
| **Intercept** | 30.7719±0.731 | 42.123 | <0.001 | 29.340 | 32.204 |
| **Defeat day** | -0.0489±0.146 | -0.336 | 0.737 | -0.334 | 0.236 |
| **SI time** | -0.0741±0.038 | -1.932 | 0.053 | -0.149 | 0.001 |
| **Defeat day * SI time** | 0.0052±0.007 | 0.755 | 0.450 | -0.008 | 0.019 |

**Supplementary Data Table 9:** 2-sided GEE regression of behavior (**Figure S3F**, fourth panel): % time mice are fleeing while attacked = defeat day + susceptibility category + defeat day*susceptibility category + intercept, grouped by mouse. Number of mice (groups) = 75, minimum samples per group 33, maximum samples per group 42, dependence structure = independence, family = Gaussian. NOTE: defeat day is coded as mean centered

| *Model: % time mice are **fleeing while attacked** = defeat day +* susceptibility *category + defeat day**susceptibility *category + intercept* | **$\beta$±standard error** | **z-stat** | **p-value** | **95% CI [lower, upper]** | |
|---|---|---|---|---|---|
| **Intercept** | 32.5861±0.876 | 37.192 | <0.001 | 30.869 | 34.303 |
| **Defeat day** | -0.2890±0.186 | -1.551 | 0.121 | -0.654 | 0.076 |
| **Susceptibility category** | -3.2396±1.397 | -2.319 | 0.020 | -5.977 | -0.502 |
| **Defeat day * Susceptibility category** | 0.4288±0.281 | 1.524 | 0.128 | -0.123 | 0.980 |

**Supplementary Data Table 10:** 2-sided GEE regression of LHb GCaMP (**Figure 4H**): Z-scored LHb GCaMP ΔF/F at attack onset = defeat day + SI time + defeat day*SI time + intercept, grouped by mouse. Number of mice (groups) = 21, minimum samples per group 10, maximum samples per group 11, dependence structure = independence, family = Gaussian. NOTE: defeat day and SI time are coded as mean centered.

| *Model: **attack onset** Z-scored LHb (GCaMP) ΔF/F = defeat day + SI time + defeat day*SI time + intercept* | **$\beta$±standard error** | **z-stat** | **p-value** | **95% CI [lower, upper]** | |
|---|---|---|---|---|---|
| **Intercept** | 0.6356±0.062 | 10.321 | <0.001 | 0.515 | 0.756 |
| **Defeat day** | 0.0068±0.013 | 0.537 | 0.591 | -0.018 | 0.031 |
| **SI time** | -0.0062±0.003 | -2.428 | 0.015 | -0.011 | -0.001 |

| | | | | | |
|---|---|---|---|---|---|
| **Defeat day * SI time** | 0.0011±0.001 | 2.088 | 0.037 | 6.58e-05 | 0.002 |

**Supplementary Data Table 11:** 2-sided GEE regression of LHb GCaMP (**Figure 4H**): Z-scored LHb GCaMP ΔF/F at attack onset = defeat day + susceptibility category + defeat day*susceptibility category + intercept, grouped by mouse. Number of mice (groups) = 21, minimum samples per group 10, maximum samples per group 11, dependence structure = independence, family = Gaussian. NOTE: defeat day is coded as mean centered.

| *Model: **attack onset** Z-scored LHb (GCaMP) ΔF/F = defeat day + susceptibility category + defeat day*susceptibility category + intercept* | $\beta$**±standard error** | **z-stat** | **p-value** | **95% CI [lower, upper]** | |
|---|---|---|---|---|---|
| **Intercept** | 0.7753±0.095 | 8.202 | <0.001 | 0.590 | 0.961 |
| **Defeat day** | -0.0158±0.023 | -0.681 | 0.496 | -0.061 | 0.030 |
| **Susceptibility category** | -0.2872±0.122 | -2.355 | 0.019 | -0.526 | -0.048 |
| **Defeat day * Susceptibility category** | 0.0509±0.026 | 1.933 | 0.053 | -0.001 | 0.102 |

**Supplementary Data Table 12:** 2-sided GEE regression of LHb GCaMP (**Figure 4I**): Z-scored LHb GCaMP ΔF/F at fighting onset = defeat day + SI time + defeat day*SI time + intercept, grouped by mouse. Number of mice (groups) = 21, minimum samples per group 10, maximum samples per group 11, dependence structure = independence, family = Gaussian. NOTE: defeat day is coded as mean centered.

| *Model: **fighting onset** Z-scored LHb (GCaMP) ΔF/F = defeat day + SI time + defeat day*SI time + intercept* | $\beta$**±standard error** | **z-stat** | **p-value** | **95% CI [lower, upper]** | |
|---|---|---|---|---|---|
| **Intercept** | 0.7431±0.086 | 8.664 | <0.001 | 0.575 | 0.911 |
| **Defeat day** | -0.0060±0.014 | -0.431 | 0.666 | -0.033 | 0.021 |
| **SI time** | -0.0072±0.003 | -2.089 | 0.037 | -0.014 | -0.000 |
| **Defeat day * SI time** | 0.0004±0.001 | 0.808 | 0.419 | -0.001 | 0.001 |

**Supplementary Data Table 13** 2-sided GEE regression of LHb GCaMP(**Figure 4I**): Z-scored LHb GCaMP ΔF/F at fighting onset = defeat day + susceptibility category + defeat day*susceptibility category + intercept, grouped by mouse. Number of mice (groups) = 21, minimum samples per group 10, maximum samples per group 11, dependence structure = independence, family = Gaussian. NOTE: defeat day is coded as mean centered.

| *Model:* ***fighting onset*** *Z-scored LHb (GCaMP) ΔF/F = defeat day + susceptibility category + defeat day*susceptibility category + intercept* | $\beta$±standard error | z-stat | p-value | 95% CI [lower, upper] | |
|---|---|---|---|---|---|
| **Intercept** | 0.9179±0.126 | 7.297 | <0.001 | 0.671 | 1.164 |
| **Defeat day** | -0.0131±0.025 | -0.532 | 0.594 | -0.061 | 0.035 |
| **Susceptibility category** | -0.3589±0.167 | -2.143 | 0.032 | -0.687 | -0.031 |
| **Defeat day * Susceptibility category** | 0.0163±0.027 | 0.602 | 0.547 | -0.037 | 0.069 |

**Supplementary Data Table 14:** 2-sided GEE regression of LHb GCaMP (**Figure 4J**): Z-scored LHb GCaMP ΔF/F at fleeing onset = defeat day + SI time + defeat day*SI time + intercept, grouped by mouse. Number of mice (groups) = 21, minimum samples per group 10, maximum samples per group 11, dependence structure = independence, family = Gaussian. NOTE: defeat day is coded as mean centered.

| *Model:* ***fleeing onset*** *Z-scored LHb (GCaMP) ΔF/F = defeat day + SI time + defeat day*SI time + intercept* | $\beta$±standard error | z-stat | p-value | 95% CI [lower, upper] | |
|---|---|---|---|---|---|
| **Intercept** | 0.3331±0.038 | 8.657 | <0.001 | 0.258 | 0.409 |
| **Defeat day** | -0.0051±0.010 | -0.539 | 0.590 | -0.024 | 0.014 |
| **SI time** | -0.0032±0.002 | -1.607 | 0.108 | -0.007 | 0.001 |
| **Defeat day * SI time** | 0.0008±0.000 | 2.520 | 0.012 | 0.000 | 0.001 |

**Supplementary DataTable 15:** 2-sided GEE regression of LHb GCaMP (**Figure 4J**): Z-scored LHb GCaMP ΔF/F at fleeing onset = defeat day + susceptibility category + defeat day*susceptibility category + intercept, grouped by mouse. Number of mice (groups) = 21, minimum samples per group 10, maximum samples per group 11, dependence structure = independence, family = Gaussian. NOTE: defeat day is coded as mean centered.

| *Model:* ***fleeing onset*** *Z-scored LHb (GCaMP) ΔF/F = defeat day + susceptibility category + defeat day*susceptibility category + intercept* | $\beta$±standard error | z-stat | p-value | 95% CI [lower, upper] | |
|---|---|---|---|---|---|
| **Intercept** | 0.3839±0.052 | 7.361 | <0.001 | 0.282 | 0.486 |
| **Defeat day** | -0.0181±0.015 | -1.183 | 0.237 | -0.048 | 0. 012 |
| **Susceptibility category** | -0.1035±0.081 | -1.283 | 0.199 | -0.262 | 0.055 |

| | | | | | |
|---|---|---|---|---|---|
| **Defeat day * Susceptibility category** | 0.0277±0.020 | 1.411 | 0.158 | -0.011 | 0.066 |

**Supplementary Data Table 16:** 2-sided GEE regression of LHb GCaMP (**Figure 4K**): Z-scored LHb GCaMP ΔF/F at vigilance onset = defeat day + SI time + defeat day*SI time + intercept, grouped by mouse. Number of mice (groups) = 21, minimum samples per group 10, maximum samples per group 11, dependence structure = independence, family = Gaussian. NOTE: defeat day is coded as mean centered.

| *Model:* ***vigilance onset*** *Z-scored LHb (GCaMP) ΔF/F = defeat day + SI time + defeat day*SI time + intercept* | $\beta$**±standard error** | **z-stat** | **p-value** | **95% CI [lower, upper]** | |
|---|---|---|---|---|---|
| **Intercept** | -0.1200±0.025 | -4.782 | <0.001 | -0.169 | -0.071 |
| **Defeat day** | 0.0009±0.005 | 0.188 | 0.851 | -0.009 | 0.010 |
| **SI time** | 0.0023±0.001 | 2.185 | 0.029 | 0.000 | 0.004 |
| **Defeat day * SI time** | 4.642e-05±0.000 | 0.207 | 0.836 | -0.000 | 0.000 |

**Supplementary DataTable 17:** 2-sided GEE regression of LHb GCaMP (**Figure 4K**): Z-scored LHb GCaMP ΔF/F at vigilance onset = defeat day + susceptibility category + defeat day*susceptibility category + intercept, grouped by mouse. Number of mice (groups) = 21, minimum samples per group 10, maximum samples per group 11, dependence structure = independence, family = Gaussian. NOTE: defeat day is coded as mean centered.

| *Model:* ***vigilance onset*** *Z-scored LHb (GCaMP) ΔF/F = defeat day + susceptibility category + defeat day*susceptibility category + intercept* | $\beta$**±standard error** | **z-stat** | **p-value** | **95% CI [lower, upper]** | |
|---|---|---|---|---|---|
| **Intercept** | -0.1733±0.039 | -4.471 | <0.001 | -0.249 | -0.097 |
| **Defeat day** | 0.0028±0.006 | 0.494 | 0.621 | -0.008 | 0.014 |
| **Susceptibility category** | 0.1094±0.050 | 2.206 | 0.027 | 0.012 | 0.207 |
| **Defeat day * Susceptibility category** | -0.0044±0.010 | -0.441 | 0.659 | -0.024 | 0.015 |

**Supplementary Data Table 18:** 2-sided GEE regression of LHb GCaMP (**Figure S5A**): Z-scored LHb GCaMP ΔF/F at *t*-SNE 1 onset = defeat day + SI time + defeat day*SI time + intercept, grouped by mouse. Number of mice (groups) = 21, minimum samples per group 10, maximum samples per group 11, dependence structure = independence, family = Gaussian. NOTE: defeat day is coded as mean centered.

| *Model: **t-SNE 1 onset** Z-scored LHb (GCaMP) ΔF/F = defeat day + SI time + defeat day*SI time + intercept* | **$\beta$±standard error** | **z-stat** | **p-value** | **95% CI [lower, upper]** | |
|---|---|---|---|---|---|
| **Intercept** | 0.3496±0.052 | 6.733 | <0.001 | 0.248 | 0.451 |
| **Defeat day** | 0.0039±0.008 | 0.470 | 0.638 | -0.012 | 0.020 |
| **SI time** | -0.0055±0.003 | -2.091 | 0.037 | -0.011 | -0.000 |
| **Defeat day * SI time** | 9.822e-05±0.000 | 0.261 | 0.794 | -0.001 | 0.001 |

**Supplementary DataTable 19:** 2-sided GEE regression of LHb GCaMP (**Figure S5A**): Z-scored LHb GCaMP ΔF/F at *t*-SNE 1 onset = defeat day + susceptibility category + defeat day*susceptibility category + intercept, grouped by mouse. Number of mice (groups) = 21, minimum samples per group 10, maximum samples per group 11, dependence structure = independence, family = Gaussian. NOTE: defeat day is coded as mean centered.

| *Model: **t-SNE 1 onset** Z-scored LHb (GCaMP) ΔF/F = defeat day + susceptibility category + defeat day*susceptibility category + intercept* | **$\beta$±standard error** | **z-stat** | **p-value** | **95% CI [lower, upper]** | |
|---|---|---|---|---|---|
| **Intercept** | 0.4447±0.052 | 6.733 | <0.001 | 0.248 | 0.451 |
| **Defeat day** | 0.0048±0.013 | 0.355 | 0.722 | -0.022 | 0.031 |
| **Susceptibility category** | -0.1928±0.110 | -1.753 | 0.080 | -0.408 | 0.023 |
| **Defeat day * Susceptibility category** | -0.0010±0.016 | -0.059 | 0.953 | -0.033 | 0.031 |

**Supplementary Data Table 20:** 2-sided GEE regression of LHb GCaMP (**Figure S5B**): Z-scored LHb GCaMP ΔF/F at *t*-SNE 2 onset = defeat day + SI time + defeat day*SI time + intercept, grouped by mouse. Number of mice (groups) = 21, minimum samples per group 10, maximum samples per group 11, dependence structure = independence, family = Gaussian. NOTE: defeat day is coded as mean centered.

| *Model: **t-SNE 2 onset** Z-scored LHb (GCaMP) ΔF/F = defeat day + SI time + defeat day*SI time + intercept* | **$\beta$±standard error** | **z-stat** | **p-value** | **95% CI [lower, upper]** | |
|---|---|---|---|---|---|
| **Intercept** | 0.5030±0.068 | 7.415 | <0.001 | 0.370 | 0.636 |
| **Defeat day** | -0.0037±0.010 | -0.366 | 0.714 | -0.023 | 0.016 |
| **SI time** | -0.0067±0.003 | -2.018 | 0.044 | -0.013 | -0.000 |
| **Defeat day * SI time** | 0.0005±0.000 | 1.044 | 0.296 | -0.000 | 0.001 |

**Supplementary DataTable 21:** 2-sided GEE regression of LHb GCaMP (**Figure S5B**): Z-scored LHb GCaMP ΔF/F at *t*-SNE 2 onset = defeat day + susceptibility category + defeat day*susceptibility category + intercept, grouped by mouse. Number of mice (groups) = 21, minimum samples per group 10, maximum samples per group 11, dependence structure = independence, family = Gaussian. NOTE: defeat day is coded as mean centered.

| *Model:* ***t-SNE 2 onset*** *Z-scored LHb (GCaMP) ΔF/F = defeat day + susceptibility category + defeat day*susceptibility category + intercept* | **$\beta$±standard error** | **z-stat** | **p-value** | **95% CI [lower, upper]** | |
|---|---|---|---|---|---|
| **Intercept** | 0.6114±0.096 | 6.394 | <0.001 | 0.424 | 0.799 |
| **Defeat day** | -0.0141±0.014 | -0.980 | 0.327 | -0.042 | 0.014 |
| **Susceptibility category** | -0.2197±0.144 | -1.522 | 0.128 | -0.503 | 0.063 |
| **Defeat day * Susceptibility category** | 0.0228±0.020 | 1.159 | 0.247 | -0.016 | 0.061 |

**Supplementary Data Table 22:** 2-sided GEE regression of LHb GCaMP (**Figure S5C**): Z-scored LHb GCaMP ΔF/F at *t*-SNE 3 onset = defeat day + SI time + defeat day*SI time + intercept, grouped by mouse. Number of mice (groups) = 21, minimum samples per group 10, maximum samples per group 11, dependence structure = independence, family = Gaussian. NOTE: defeat day is coded as mean centered.

| *Model:* ***t-SNE 3 onset*** *Z-scored LHb (GCaMP) ΔF/F = defeat day + SI time + defeat day*SI time + intercept* | **$\beta$±standard error** | **z-stat** | **p-value** | **95% CI [lower, upper]** | |
|---|---|---|---|---|---|
| **Intercept** | 0.2649±0.024 | 10.867 | <0.001 | 0.217 | 0.313 |
| **Defeat day** | 0.0040±0.004 | 0.989 | 0.323 | -0.004 | 0.012 |
| **SI time** | -0.0026±0.001 | -2.475 | 0.013 | -0.005 | 0.012 |
| **Defeat day * SI time** | -3.565e-05±0.0 | -0.163 | 0.871 | -0.000 | 0.000 |

**Supplementary DataTable 23:** 2-sided GEE regression of LHb GCaMP (**Figure S5C**): Z-scored LHb GCaMP ΔF/F at *t*-SNE 3 onset = defeat day + susceptibility category + defeat day*susceptibility category + intercept, grouped by mouse. Number of mice (groups) = 21, minimum samples per group 10, maximum samples per group 11, dependence structure = independence, family = Gaussian. NOTE: defeat day is coded as mean centered.

| *Model: **t-SNE 3 onset** Z-scored LHb (GCaMP) ΔF/F = defeat day + susceptibility category + defeat day*susceptibility category + intercept* | **$\beta$±standard error** | **z-stat** | **p-value** | **95% CI [lower, upper]** | |
|---|---|---|---|---|---|
| **Intercept** | 0.3092±0.041 | 7.458 | <0.001 | 0.228 | 0.390 |
| **Defeat day** | 0.0038±0.005 | 0.750 | 0.453 | -0.006 | 0.014 |
| **Susceptibility category** | -0.0898±0.051 | -1.744 | 0.081 | -0.191 | 0.011 |
| **Defeat day * Susceptibility category** | 0.0007±0.008 | 0.086 | 0.932 | -0.015 | 0.016 |

**Supplementary Data Table 24:** 2-sided GEE regression of LHb GCaMP (**Figure S5D**): Z-scored LHb GCaMP ΔF/F at *t*-SNE 4 onset = defeat day + SI time + defeat day*SI time + intercept, grouped by mouse. Number of mice (groups) = 21, minimum samples per group 10, maximum samples per group 11, dependence structure = independence, family = Gaussian. NOTE: defeat day is coded as mean centered.

| *Model: **t-SNE 4 onset** Z-scored LHb (GCaMP) ΔF/F = defeat day + SI time + defeat day*SI time + intercept* | **$\beta$±standard error** | **z-stat** | **p-value** | **95% CI [lower, upper]** | |
|---|---|---|---|---|---|
| **Intercept** | 0.4460±0.063 | 7.042 | <0.001 | 0.322 | 0.570 |
| **Defeat day** | 0.0029±0.009 | 0.330 | 0.741 | -0.014 | 0.020 |
| **SI time** | -0.0061±0.003 | -2.045 | 0.041 | -0.012 | -0.000 |
| **Defeat day * SI time** | 0.0004±0.000 | 1.250 | 0.211 | -0.000 | 0.001 |

**Supplementary DataTable 25:** 2-sided GEE regression of LHb GCaMP (**Figure S5D**): Z-scored LHb GCaMP ΔF/F at *t*-SNE 4 onset = defeat day + susceptibility category + defeat day*susceptibility category + intercept, grouped by mouse. Number of mice (groups) = 21, minimum samples per group 10, maximum samples per group 11, dependence structure = independence, family = Gaussian. NOTE: defeat day is coded as mean centered.

| *Model: **t-SNE 4 onset** Z-scored LHb (GCaMP) ΔF/F = defeat day + susceptibility category + defeat day*susceptibility category + intercept* | **$\beta$±standard error** | **z-stat** | **p-value** | **95% CI [lower, upper]** | |
|---|---|---|---|---|---|
| **Intercept** | 0.5532±0.091 | 6.053 | <0.001 | 0.374 | 0.732 |
| **Defeat day** | -0.0028±0.015 | -0.184 | 0.854 | -0.032 | 0.027 |
| **Susceptibility category** | -0.2178±0.133 | -1.635 | 0.102 | -0.479 | 0.043 |

| | | | | | |
|---|---|---|---|---|---|
| **Defeat day * Susceptibility category** | 0.0127±0.017 | 0.740 | 0.469 | -0.021 | 0.046 |

**Supplementary Data Table 26:** 2-sided GEE regression of LHb GCaMP (**Figure S5E**): Z-scored LHb GCaMP ΔF/F at *t*-SNE 5 onset = defeat day + SI time + defeat day*SI time + intercept, grouped by mouse. Number of mice (groups) = 21, minimum samples per group 10, maximum samples per group 11, dependence structure = independence, family = Gaussian. NOTE: defeat day is coded as mean centered.

| *Model: **t-SNE 5 onset** Z-scored LHb (GCaMP) ΔF/F = defeat day + SI time + defeat day*SI time + intercept* | $\beta$**±standard error** | **z-stat** | **p-value** | **95% CI [lower, upper]** | |
|---|---|---|---|---|---|
| **Intercept** | 0.1853±0.038 | 4.896 | <0.001 | 0.111 | 0.259 |
| **Defeat day** | 0.0042±0.007 | 0.610 | 0.542 | -0.009 | 0.018 |
| **SI time** | 0.0046±0.002 | 1.879 | 0.060 | -0.000 | 0.009 |
| **Defeat day * SI time** | 7.316e05±0.0 | 0.255 | 0.799 | -0.000 | 0.001 |

**Supplementary DataTable 27:** 2-sided GEE regression of LHb GCaMP (**Figure S5E**): Z-scored LHb GCaMP ΔF/F at *t*-SNE 5 onset = defeat day + susceptibility category + defeat day*susceptibility category + intercept, grouped by mouse. Number of mice (groups) = 21, minimum samples per group 10, maximum samples per group 11, dependence structure = independence, family = Gaussian. NOTE: defeat day is coded as mean centered.

| *Model: **t-SNE 5 onset** Z-scored LHb (GCaMP) ΔF/F = defeat day + susceptibility category + defeat day*susceptibility category + intercept* | $\beta$**±standard error** | **z-stat** | **p-value** | **95% CI [lower, upper]** | |
|---|---|---|---|---|---|
| **Intercept** | 0.0930±0.039 | 2.360 | 0.018 | 0.016 | 0.170 |
| **Defeat day** | 0.0010±0.008 | 0.127 | 0.899 | -0.014 | 0.016 |
| **Susceptibility category** | 0.1877±0.080 | 2.339 | 0.019 | 0.030 | 0.345 |
| **Defeat day * Susceptibility category** | 0.0061±0.014 | 0.427 | 0.669 | -0.022 | 0.034 |

**Supplementary Data Table 28**: 2-sided GEE regression of LHb GCaMP (**Figure S5F**): Z-scored LHb GCaMP ΔF/F at *t*-SNE 6 onset = defeat day + SI time + defeat day*SI time + intercept, grouped by mouse. Number of mice (groups) = 21, minimum samples per group 10, maximum samples per group 11, dependence structure = independence, family = Gaussian. NOTE: defeat day is coded as mean centered.

| *Model: **t-SNE 6 onset** Z-scored LHb (GCaMP) ΔF/F = defeat day + SI time + defeat day*SI time + intercept* | $\beta$±standard error | z-stat | p-value | 95% CI [lower, upper] | |
|---|---|---|---|---|---|
| **Intercept** | 0.4166±0.051 | 8.157 | <0.001 | 0.317 | 0.517 |
| **Defeat day** | 0.0049±0.009 | 0.527 | 0.598 | -0.013 | 0.023 |
| **SI time** | -0.0063±0.002 | -2.549 | 0.011 | -0.011 | -0.001 |
| **Defeat day * SI time** | 0.0003±0.000 | 0.760 | 0.447 | -0.000 | 0.001 |

**Supplementary DataTable 29**: 2-sided GEE regression of LHb GCaMP (**Figure S5F**): Z-scored LHb GCaMP ΔF/F at *t*-SNE 6 onset = defeat day + susceptibility category + defeat day*susceptibility category + intercept, grouped by mouse. Number of mice (groups) = 21, minimum samples per group 10, maximum samples per group 11, dependence structure = independence, family = Gaussian. NOTE: defeat day is coded as mean centered.

| *Model: **t-SNE 6 onset** Z-scored LHb (GCaMP) ΔF/F = defeat day + susceptibility category + defeat day*susceptibility category + intercept* | $\beta$±standard error | z-stat | p-value | 95% CI [lower, upper] | |
|---|---|---|---|---|---|
| **Intercept** | 0.5310±0.072 | 7.338 | <0.001 | 0.389 | 0.673 |
| **Defeat day** | -0.0003±0.015 | -0.022 | 0.983 | -0.029 | 0.029 |
| **Susceptibility category** | -0.2324±0.110 | -2.107 | 0.035 | -0.449 | -0.016 |
| **Defeat day * Susceptibility category** | 0.0118±0.018 | 0.650 | 0.516 | -0.024 | 0.048 |

**Supplementary Data Table 30**: 2-sided GEE regression of LHb GCaMP (**Figure S5G**): Z-scored LHb GCaMP ΔF/F at *t*-SNE 7 onset = defeat day + SI time + defeat day*SI time + intercept, grouped by mouse. Number of mice (groups) = 21, minimum samples per group 10, maximum samples per group 11, dependence structure = independence, family = Gaussian. NOTE: defeat day is coded as mean centered.

| *Model: **t-SNE 7 onset** Z-scored LHb (GCaMP) ΔF/F = defeat day + SI time + defeat day*SI time + intercept* | $\beta$±standard error | z-stat | p-value | 95% CI [lower, upper] | |
|---|---|---|---|---|---|
| **Intercept** | 0.1421±0.037 | 3.886 | <0.001 | 0.070 | 0.214 |
| **Defeat day** | 0.0017±0.007 | 0.238 | 0.812 | -0.012 | 0.016 |
| **SI time** | 0.0041±0.002 | 1.810 | 0.070 | -0.000 | 0.009 |

| | | | | | |
|---|---|---|---|---|---|
| **Defeat day * SI time** | 0.0005±0.000 | 1.615 | 0.106 | -0.000 | 0.001 |

**Supplementary DataTable 31**: 2-sided GEE regression of LHb GCaMP (**Figure S5G**): Z-scored LHb GCaMP ΔF/F at *t*-SNE 7 onset = defeat day + susceptibility category + defeat day*susceptibility category + intercept, grouped by mouse. Number of mice (groups) = 21, minimum samples per group 10, maximum samples per group 11, dependence structure = independence, family = Gaussian. NOTE: defeat day is coded as mean centered.

| *Model: **t-SNE 7 onset** Z-scored LHb (GCaMP) ΔF/F = defeat day + susceptibility category + defeat day*susceptibility category + intercept* | **$\beta$±standard error** | **z-stat** | **p-value** | **95% CI [lower, upper]** | |
|---|---|---|---|---|---|
| **Intercept** | 0.0592±0.036 | 1.652 | 0.098 | -0.011 | 0.129 |
| **Defeat day** | -0.0071±0.006 | -1.096 | 0.273 | -0.020 | 0.006 |
| **Susceptibility category** | 0.1682±0.076 | 2.200 | 0.028 | 0.018 | 0.318 |
| **Defeat day * Susceptibility category** | 0.0179±0.015 | 1.184 | 0.236 | -0.012 | 0.047 |

**Supplementary Data Table 32:** 2-sided GEE regression of LHb GCaMP (**Figure S5H**): Z-scored LHb GCaMP ΔF/F at *t*-SNE 8 onset = defeat day + SI time + defeat day*SI time + intercept, grouped by mouse. Number of mice (groups) = 21, minimum samples per group 10, maximum samples per group 11, dependence structure = independence, family = Gaussian. NOTE: defeat day is coded as mean centered.

| *Model: **t-SNE 8 onset** Z-scored LHb (GCaMP) ΔF/F = defeat day + SI time + defeat day*SI time + intercept* | **$\beta$±standard error** | **z-stat** | **p-value** | **95% CI [lower, upper]** | |
|---|---|---|---|---|---|
| **Intercept** | 0.3719±0.046 | 8.002 | <0.001 | 0.281 | 0.463 |
| **Defeat day** | 0.0058±0.007 | 0.834 | 0.404 | -0.008 | 0.019 |
| **SI time** | -0.0050±0.002 | -2.113 | 0.035 | -0.010 | -0.000 |
| **Defeat day * SI time** | 0.0004±0.000 | 1.289 | 0.197 | -0.000 | 0.001 |

**Supplementary DataTable 33:** 2-sided GEE regression of LHb GCaMP (**Figure S5H**): Z-scored LHb GCaMP ΔF/F at *t*-SNE 8 onset = defeat day + susceptibility category + defeat day*susceptibility category + intercept, grouped by mouse. Number of mice (groups) = 21, minimum samples per group 10, maximum samples per group 11, dependence structure = independence, family = Gaussian. NOTE: defeat day is coded as mean centered.

| *Model: **t-SNE 8 onset** Z-scored LHb (GCaMP) ΔF/F = defeat day + susceptibility category + defeat day*susceptibility category + intercept* | **$\beta$±standard error** | **z-stat** | **p-value** | **95% CI [lower, upper]** | |
|---|---|---|---|---|---|
| **Intercept** | 0.4588±0.067 | 6.812 | 0.000 | 0.327 | 0.591 |
| **Defeat day** | -0.0013±0.011 | -0.125 | 0.900 | -0.022 | 0.019 |
| **Susceptibility category** | -0.1766±0.099 | -1.784 | 0.074 | -0.371 | 0.017 |
| **Defeat day * Susceptibility category** | 0.0156±0.014 | 1.121 | 0.262 | -0.012 | 0.043 |

**Supplementary Data Table 34:** 2-sided GEE regression of LHb GCaMP (**Figure S6A**): Z-scored LHb GCaMP ΔF/F at *t*-SNE 9 onset = defeat day + SI time + defeat day*SI time + intercept, grouped by mouse. Number of mice (groups) = 21, minimum samples per group 10, maximum samples per group 11, dependence structure = independence, family = Gaussian. NOTE: defeat day is coded as mean centered.

| *Model: **t-SNE 9 onset** Z-scored LHb (GCaMP) ΔF/F = defeat day + SI time + defeat day*SI time + intercept* | **$\beta$±standard error** | **z-stat** | **p-value** | **95% CI [lower, upper]** | |
|---|---|---|---|---|---|
| **Intercept** | 0.1070±0.030 | 3.566 | <0.001 | 0.048 | 0.166 |
| **Defeat day** | 0.0062±0.007 | 0.903 | 0.366 | -0.007 | 0.020 |
| **SI time** | 0.0037±0.002 | 1.997 | 0.046 | 6.85e-05 | 0.007 |
| **Defeat day * SI time** | 0.0003±0.000 | 1.025 | 0.305 | -0.000 | 0.001 |

**Supplementary DataTable 35:** 2-sided GEE regression of LHb GCaMP (**Figure S6A**): Z-scored LHb GCaMP ΔF/F at *t*-SNE 9 onset = defeat day + susceptibility category + defeat day*susceptibility category + intercept, grouped by mouse. Number of mice (groups) = 21, minimum samples per group 10, maximum samples per group 11, dependence structure = independence, family = Gaussian. NOTE: defeat day is coded as mean centered.

| *Model: **t-SNE 9 onset** Z-scored LHb (GCaMP) ΔF/F = defeat day + susceptibility category + defeat day*susceptibility category + intercept* | **$\beta$±standard error** | **z-stat** | **p-value** | **95% CI [lower, upper]** | |
|---|---|---|---|---|---|
| **Intercept** | 0.0263±0.031 | 0.846 | 0.398 | -0.035 | 0.087 |
| **Defeat day** | 0.0007±0.007 | 0.094 | 0.925 | -0.014 | 0.015 |
| **Susceptibility category** | 0.1647±0.061 | 2.681 | 0.007 | 0.044 | 0.285 |

| | | | | | |
|---|---|---|---|---|---|
| **Defeat day * Susceptibility category** | 0.0110±0.014 | 0.781 | 0.435 | -0.017 | 0.039 |

**Supplementary Data Table 36:** 2-sided GEE regression of LHb GCaMP (**Figure S6B**): Z-scored LHb GCaMP ΔF/F at *t*-SNE 10 onset = defeat day + SI time + defeat day*SI time + intercept, grouped by mouse. Number of mice (groups) = 21, minimum samples per group 10, maximum samples per group 11, dependence structure = independence, family = Gaussian. NOTE: defeat day is coded as mean centered.

| *Model: **t-SNE 10 onset** Z-scored LHb (GCaMP) ΔF/F = defeat day + SI time + defeat day*SI time + intercept* | **$\beta$±standard error** | **z-stat** | **p-value** | **95% CI [lower, upper]** | |
|---|---|---|---|---|---|
| **Intercept** | 0.0998±0.027 | 3.656 | <0.001 | 0.046 | 0.153 |
| **Defeat day** | 0.0038±0.006 | 0.616 | 0.538 | -0.008 | 0.016 |
| **SI time** | -0.0027±0.001 | -2.139 | 0.032 | -0.005 | -0.000 |
| **Defeat day * SI time** | -0.0002±0.000 | -0.654 | 0.513 | -0.001 | 0.000 |

**Supplementary DataTable 37:** 2-sided GEE regression of LHb GCaMP (**Figure S6B**): Z-scored LHb GCaMP ΔF/F at *t*-SNE 10 onset = defeat day + susceptibility category + defeat day*susceptibility category + intercept, grouped by mouse. Number of mice (groups) = 21, minimum samples per group 10, maximum samples per group 11, dependence structure = independence, family = Gaussian. NOTE: defeat day is coded as mean centered.

| *Model: **t-SNE 10 onset** Z-scored LHb (GCaMP) ΔF/F = defeat day + susceptibility category + defeat day*susceptibility category + intercept* | **$\beta$±standard error** | **z-stat** | **p-value** | **95% CI [lower, upper]** | |
|---|---|---|---|---|---|
| **Intercept** | 0.1568±0.043 | 3.652 | <0.001 | 0.073 | 0.241 |
| **Defeat day** | 0.0094±0.009 | 1.022 | 0.307 | -0.009 | 0.027 |
| **Susceptibility category** | -0.1160±0.056 | -2.067 | 0.039 | -0.226 | -0.006 |
| **Defeat day * Susceptibility category** | -0.0112±0.012 | -0.933 | 0.351 | -0.035 | 0.012 |

**Supplementary Data Table 38:** 2-sided GEE regression of LHb GCaMP (**Figure S6C**): Z-scored LHb GCaMP ΔF/F at *t*-SNE 11 onset = defeat day + SI time + defeat day*SI time + intercept, grouped by mouse. Number of mice (groups) = 21, minimum samples per group 10, maximum samples per group 11, dependence structure = independence, family = Gaussian. NOTE: defeat day is coded as mean centered.

| *Model: **t-SNE 11 onset** Z-scored LHb (GCaMP) ΔF/F = defeat day + SI time + defeat day*SI time + intercept* | **$\beta$±standard error** | **z-stat** | **p-value** | **95% CI [lower, upper]** | |
|---|---|---|---|---|---|
| **Intercept** | 0.2530±0.037 | 6.769 | <0.001 | 0.180 | 0.326 |
| **Defeat day** | 0.0083±0.008 | 1.076 | 0.282 | -0.007 | 0.028 |
| **SI time** | -0.0042±0.002 | -2.354 | 0.019 | -0.008 | -0.001 |
| **Defeat day * SI time** | -3.888e-05±0.0 | -0.119 | 0.905 | -0.001 | 0.001 |

**Supplementary DataTable 39:** 2-sided GEE regression of LHb GCaMP (**Figure S6C**): Z-scored LHb GCaMP ΔF/F at *t*-SNE 11 onset = defeat day + susceptibility category + defeat day*susceptibility category + intercept, grouped by mouse. Number of mice (groups) = 21, minimum samples per group 10, maximum samples per group 11, dependence structure = independence, family = Gaussian. NOTE: defeat day is coded as mean centered.

| *Model: **t-SNE 11 onset** Z-scored LHb (GCaMP) ΔF/F = defeat day + susceptibility category + defeat day*susceptibility category + intercept* | **$\beta$±standard error** | **z-stat** | **p-value** | **95% CI [lower, upper]** | |
|---|---|---|---|---|---|
| **Intercept** | 0.3275±0.053 | 6.174 | <0.001 | 0.224 | 0.432 |
| **Defeat day** | 0.0083±0.014 | 0.592 | 0.554 | -0.019 | 0.036 |
| **Susceptibility category** | -0.1512±0.081 | -1.877 | 0.061 | -0.309 | 0.007 |
| **Defeat day * Susceptibility category** | 0.0006±0.015 | 0.042 | 0.967 | -0.029 | 0.030 |

**Supplementary Data Table 40:** 2-sided GEE regression of LHb GCaMP (**Figure S6D**): Z-scored LHb GCaMP ΔF/F at *t*-SNE 12 onset = defeat day + SI time + defeat day*SI time + intercept, grouped by mouse. Number of mice (groups) = 21, minimum samples per group 10, maximum samples per group 11, dependence structure = independence, family = Gaussian. NOTE: defeat day is coded as mean centered.

| *Model: **t-SNE 12 onset** Z-scored LHb (GCaMP) ΔF/F = defeat day + SI time + defeat day*SI time + intercept* | **$\beta$±standard error** | **z-stat** | **p-value** | **95% CI [lower, upper]** | |
|---|---|---|---|---|---|
| **Intercept** | 0.1242±0.022 | 5.541 | <0.001 | 0.080 | 0.168 |
| **Defeat day** | 0.0052±0.005 | 1.079 | 0.281 | -0.004 | 0.015 |
| **SI time** | -0.0020±0.001 | -1.944 | 0.052 | -0.004 | 1.7e-05 |
| **Defeat day * SI time** | -9.071e-05±0.0 | -0.382 | 0.702 | -0.001 | 0.000 |

**Supplementary DataTable 41:** 2-sided GEE regression of LHb GCaMP (**Figure S6D**): Z-scored LHb GCaMP ΔF/F at *t*-SNE 12 onset = defeat day + susceptibility category + defeat day*susceptibility category + intercept, grouped by mouse. Number of mice (groups) = 21, minimum samples per group 10, maximum samples per group 11, dependence structure = independence, family = Gaussian. NOTE: defeat day is coded as mean centered.

| *Model: **t-SNE 12 onset** Z-scored LHb (GCaMP) ΔF/F = defeat day + susceptibility category + defeat day*susceptibility category + intercept* | **$\beta$±standard error** | **z-stat** | **p-value** | **95% CI [lower, upper]** | |
|---|---|---|---|---|---|
| **Intercept** | 0.1637±0.035 | 4.688 | <0.001 | 0.095 | 0.232 |
| **Defeat day** | 0.0095±0.007 | 1.313 | 0.189 | -0.005 | 0.024 |
| **Susceptibility category** | -0.0802±0.046 | -1.733 | 0.083 | -0.171 | 0.011 |
| **Defeat day * Susceptibility category** | -0.0087±0.009 | -0.916 | 0.360 | -0.027 | 0.010 |

**Supplementary Data Table 42:** 2-sided GEE regression of LHb GCaMP (**Figure S6E**): Z-scored LHb GCaMP ΔF/F at *t*-SNE 13 onset = defeat day + SI time + defeat day*SI time + intercept, grouped by mouse. Number of mice (groups) = 21, minimum samples per group 10, maximum samples per group 11, dependence structure = independence, family = Gaussian. NOTE: defeat day is coded as mean centered.

| *Model: **t-SNE 13 onset** Z-scored LHb (GCaMP) ΔF/F = defeat day + SI time + defeat day*SI time + intercept* | **$\beta$±standard error** | **z-stat** | **p-value** | **95% CI [lower, upper]** | |
|---|---|---|---|---|---|
| **Intercept** | 0.2858±0.038 | 7.451 | <0.001 | 0.211 | 0.361 |
| **Defeat day** | 0.0106±0.006 | 1.714 | 0.087 | -0.002 | 0.023 |
| **SI time** | -0.0038±0.002 | -1.888 | 0.059 | -0.008 | 0.000 |
| **Defeat day * SI time** | 0.0003±0.000 | 1.055 | 0.292 | -0.000 | 0.001 |

**Supplementary DataTable 43:** 2-sided GEE regression of LHb GCaMP (**Figure S6E**): Z-scored LHb GCaMP ΔF/F at *t*-SNE 13 onset = defeat day + susceptibility category + defeat day*susceptibility category + intercept, grouped by mouse. Number of mice (groups) = 21, minimum samples per group 10, maximum samples per group 11, dependence structure = independence, family = Gaussian. NOTE: defeat day is coded as mean centered.

| *Model:* ***t-SNE 13 onset*** *Z-scored LHb (GCaMP) ΔF/F = defeat day + susceptibility category + defeat day*susceptibility category + intercept* | **$\beta$±standard error** | **z-stat** | **p-value** | **95% CI [lower, upper]** | |
|---|---|---|---|---|---|
| **Intercept** | 0.3534±0.055 | 6.437 | <0.001 | 0.246 | 0.461 |
| **Defeat day** | 0.0042±0.009 | 0.457 | 0.648 | -0.014 | 0.022 |
| **Susceptibility category** | -0.1374±0.081 | -1.698 | 0.090 | -0.296 | 0.021 |
| **Defeat day * Susceptibility category** | 0.0138±0.012 | 1.128 | 0.259 | -0.010 | 0.038 |

**Supplementary Data Table 44:** 2-sided GEE regression of LHb GCaMP (**Figure S6F**): Z-scored LHb GCaMP ΔF/F at *t*-SNE 14 onset = defeat day + SI time + defeat day*SI time + intercept, grouped by mouse. Number of mice (groups) = 21, minimum samples per group 10, maximum samples per group 11, dependence structure = independence, family = Gaussian. NOTE: defeat day is coded as mean centered.

| *Model:* ***t-SNE 14 onset*** *Z-scored LHb (GCaMP) ΔF/F = defeat day + SI time + defeat day*SI time + intercept* | **$\beta$±standard error** | **z-stat** | **p-value** | **95% CI [lower, upper]** | |
|---|---|---|---|---|---|
| **Intercept** | 0.0647±0.051 | 1.266 | 0.206 | -0.035 | 0.165 |
| **Defeat day** | 0.0086±0.006 | 1.509 | 0.131 | -0.003 | 0.020 |
| **SI time** | 0.0049±0.003 | 1.638 | 0.101 | -0.001 | 0.011 |
| **Defeat day * SI time** | -8.11e-05±0.0 | -0.452 | 0.651 | -0.000 | 0.000 |

**Supplementary DataTable 45:** 2-sided GEE regression of LHb GCaMP (**Figure S6F**): Z-scored LHb GCaMP ΔF/F at *t*-SNE 14 onset = defeat day + susceptibility category + defeat day*susceptibility category + intercept, grouped by mouse. Number of mice (groups) = 21, minimum samples per group 10, maximum samples per group 11, dependence structure = independence, family = Gaussian. NOTE: defeat day is coded as mean centered.

| *Model:* ***t-SNE 14 onset*** *Z-scored LHb (GCaMP) ΔF/F = defeat day + susceptibility category + defeat day*susceptibility category + intercept* | **$\beta$±standard error** | **z-stat** | **p-value** | **95% CI [lower, upper]** | |
|---|---|---|---|---|---|
| **Intercept** | -0.0378±0.056 | -0.671 | 0.502 | -0.148 | 0.073 |
| **Defeat day** | 0.0109±0.008 | 1.366 | 0.172 | -0.005 | 0.027 |
| **Susceptibility category** | 0.2092±0.106 | 1.978 | 0.048 | 0.002 | 0.417 |

| | | | | | |
|---|---|---|---|---|---|
| **Defeat day * Susceptibility category** | -0.0056±0.011 | -0.492 | 0.623 | -0.028 | 0.017 |

**Supplementary Data Table 46:** 2-sided GEE regression of LHb GCaMP (**Figure S6G**): Z-scored LHb GCaMP ΔF/F at *t*-SNE 15 onset = defeat day + SI time + defeat day*SI time + intercept, grouped by mouse. Number of mice (groups) = 21, minimum samples per group 10, maximum samples per group 11, dependence structure = independence, family = Gaussian. NOTE: defeat day is coded as mean centered.

| *Model: **t-SNE 15 onset** Z-scored LHb (GCaMP) ΔF/F = defeat day + SI time + defeat day*SI time + intercept* | **$\beta$±standard error** | **z-stat** | **p-value** | **95% CI [lower, upper]** | |
|---|---|---|---|---|---|
| **Intercept** | 0.0561±0.035 | 1.626 | 0.104 | -0.012 | 0.124 |
| **Defeat day** | 0.0112±0.006 | 2.029 | 0.042 | 0.000 | 0.022 |
| **SI time** | 0.0038±0.002 | 1.684 | 0.092 | -0.001 | 0.008 |
| **Defeat day * SI time** | -4.742e-06±0.0 | -0.023 | 0.982 | -0.000 | 0.000 |

**Supplementary DataTable 47:** 2-sided GEE regression of LHb GCaMP (**Figure S76G**): Z-scored LHb GCaMP ΔF/F at *t*-SNE 15 onset = defeat day + susceptibility category + defeat day*susceptibility category + intercept, grouped by mouse. Number of mice (groups) = 21, minimum samples per group 10, maximum samples per group 11, dependence structure = independence, family = Gaussian. NOTE: defeat day is coded as mean centered.

| *Model: **t-SNE 15 onset** Z-scored LHb (GCaMP) ΔF/F = defeat day + susceptibility category + defeat day*susceptibility category + intercept* | **$\beta$±standard error** | **z-stat** | **p-value** | **95% CI [lower, upper]** | |
|---|---|---|---|---|---|
| **Intercept** | -0.0212±0.033 | -0.636 | 0.525 | -0.086 | 0.044 |
| **Defeat day** | 0.0119±0.007 | 1.600 | 0.110 | -0.003 | 0.026 |
| **Susceptibility category** | 0.1576±0.072 | 2.201 | 0.028 | 0.017 | 0.298 |
| **Defeat day * Susceptibility category** | -0.0020±0.011 | -0.179 | 0.858 | -0.024 | 0.020 |

**Supplementary Data Table 48:** 2-sided GEE regression of LHb GCaMP (**Figure S6H**): Z-scored LHb GCaMP ΔF/F at *t*-SNE 16 onset = defeat day + SI time + defeat day*SI time + intercept, grouped by mouse. Number of mice (groups) = 21, minimum samples per group 10, maximum samples per group 11, dependence structure = independence, family = Gaussian. NOTE: defeat day is coded as mean centered.

| *Model:* ***t-SNE 16 onset*** *Z-scored LHb (GCaMP) ΔF/F = defeat day + SI time + defeat day*SI time + intercept* | *β*±standard error | z-stat | p-value | 95% CI [lower, upper] | |
|---|---|---|---|---|---|
| **Intercept** | 0.1314±0.022 | 5.942 | <0.001 | 0.088 | 0.175 |
| **Defeat day** | 0.0109±0.005 | 2.119 | 0.034 | 0.001 | 0.021 |
| **SI time** | 0.0005±0.001 | 0.364 | 0.716 | -0.002 | 0.003 |
| **Defeat day * SI time** | -0.0002±0.000 | -1.121 | 0.262 | -0.001 | 0.000 |

**Supplementary DataTable 49:** 2-sided GEE regression of LHb GCaMP (**Figure S6H**): Z-scored LHb GCaMP ΔF/F at *t*-SNE 16 onset = defeat day + susceptibility category + defeat day*susceptibility category + intercept, grouped by mouse. Number of mice (groups) = 21, minimum samples per group 10, maximum samples per group 11, dependence structure = independence, family = Gaussian. NOTE: defeat day is coded as mean centered.

| *Model:* ***t-SNE 16 onset*** *Z-scored LHb (GCaMP) ΔF/F = defeat day + susceptibility category + defeat day*susceptibility category + intercept* | *β*±standard error | z-stat | p-value | 95% CI [lower, upper] | |
|---|---|---|---|---|---|
| **Intercept** | 0.1308±0.027 | 4.934 | <0.001 | 0.079 | 0.183 |
| **Defeat day** | 0.0148±0.006 | 2.303 | 0.021 | 0.002 | 0.027 |
| **Susceptibility category** | 0.0008±0.044 | 0.017 | 0.986 | -0.086 | 0.088 |
| **Defeat day * Susceptibility category** | -0.0083±0.010 | -0.790 | 0.430 | -0.029 | 0.012 |

**Supplementary Data Table 50:** 2-sided GEE regression of LHb GCaMP (**Figure S6I**): Z-scored LHb GCaMP ΔF/F at *t*-SNE 17 onset = defeat day + SI time + defeat day*SI time + intercept, grouped by mouse. Number of mice (groups) = 21, minimum samples per group 10, maximum samples per group 11, dependence structure = independence, family = Gaussian. NOTE: defeat day is coded as mean centered.

| *Model:* ***t-SNE 17 onset*** *Z-scored LHb (GCaMP) ΔF/F = defeat day + SI time + defeat day*SI time + intercept* | *β*±standard error | z-stat | p-value | 95% CI [lower, upper] | |
|---|---|---|---|---|---|
| **Intercept** | 0.0557±0.036 | 1.545 | 0.122 | -0.015 | 0.126 |
| **Defeat day** | 0.0094±0.006 | 1.573 | 0.116 | -0.002 | 0.021 |
| **SI time** | -0.0021±0.002 | -1.368 | 0.171 | -0.005 | 0.001 |
| **Defeat day * SI time** | -0.0004±0.000 | -2.197 | 0.028 | -0.001 | -4.3e-05 |

**Supplementary DataTable 51:** 2-sided GEE regression of LHb GCaMP (**Figure S6I**): Z-scored LHb GCaMP ΔF/F at *t*-SNE 17 onset = defeat day + susceptibility category + defeat day*susceptibility category + intercept, grouped by mouse. Number of mice (groups) = 21, minimum samples per group 10, maximum samples per group 11, dependence structure = independence, family = Gaussian. NOTE: defeat day is coded as mean centered.

| *Model: **t-SNE 17 onset** Z-scored LHb (GCaMP) ΔF/F = defeat day + susceptibility category + defeat day*susceptibility category + intercept* | **$\beta$±standard error** | **z-stat** | **p-value** | **95% CI [lower, upper]** | |
|---|---|---|---|---|---|
| **Intercept** | 0.1067±0.043 | 2.488 | 0.013 | 0.023 | 0.191 |
| **Defeat day** | 0.0189±0.009 | 2.209 | 0.027 | 0.002 | 0.036 |
| **Susceptibility category** | -0.1039±0.073 | -1.431 | 0.153 | -0.246 | 0.038 |
| **Defeat day * Susceptibility category** | -0.0195±0.012 | -1.652 | 0.098 | -0.043 | 0.004 |

**Supplementary Data Table 52:** 2-sided GEE regression of behavior (**Figure S7F**, first panel): % time mice are being attacked = defeat day + opsin (ChR2/ChRmine vs YFP) category + defeat day*opsin (ChR2ChRmine/YFP) category + intercept, grouped by mouse. Number of mice (groups) = 22, minimum samples per group 11, maximum samples per group 11, dependence structure = independence, family = Gaussian. NOTE: defeat day is coded as mean centered

| *Model: % time mice are **being attacked** = defeat day + opsin + defeat day*opsin category + intercept* | **$\beta$±standard error** | **z-stat** | **p-value** | **95% CI [lower, upper]** | |
|---|---|---|---|---|---|
| **Intercept** | 12.3172±0.583 | 21.121 | <0.001 | 11.174 | 13.460 |
| **Defeat day** | -0.6116±0.322 | -1.897 | 0.058 | -1.243 | 0.020 |
| **Opsin category** | -1.7172±0.938 | -1.831 | 0.067 | -3.556 | 0.121 |
| **Defeat day * Opsin category** | 0.1413±0.349 | 0.405 | 0.685 | -0.157 | 0.272 |

**Supplementary Data Table 53:** 2-sided GEE regression of behavior (**Figure S7F**, second panel): % time mice are freezing during attack = defeat day + opsin (ChR2 or ChRmine) category + defeat day*opsin (ChR2 or ChRmine) category + intercept, grouped by mouse. Number of mice (groups) = 22, minimum samples per group 11, maximum samples per group 11, dependence structure = independence, family = Gaussian. NOTE: defeat day is coded as mean centered

| *Model: % time mice are* ***freezing during attack****= defeat day + opsin category + defeat day*opsin category + intercept* | **$\beta$±standard error** | **z-stat** | **p-value** | **95% CI [lower, upper]** | |
|---|---|---|---|---|---|
| **Intercept** | 43.7667±1.883 | 25.898 | <0.001 | 45.076 | 52.457 |
| **Defeat day** | -1.1064±0.524 | -2.109 | 0.035 | -2.134 | -0.078 |
| **Opsin category** | -12.6126±2.545 | -4.956 | <0.000 | -17.601 | -7.625 |
| **Defeat day * Opsin category** | -0.1156±0.652 | -0.177 | 0.859 | -1.394 | 1.162 |

**Supplementary Data Table 54:** 2-sided GEE regression of behavior (**Figure S7F**, third panel): % time mice are fighting during attack = defeat day + opsin (ChR2 or ChRmine) category + defeat day*opsin (ChR2/ChRmine vs YFP) category + intercept, grouped by mouse. Number of mice (groups) = 22, minimum samples per group 11, maximum samples per group 11, dependence structure = independence, family = Gaussian. NOTE: defeat day is coded as mean centered

| *Model: % time mice are* ***fighting during attack*** *= defeat day + opsin category + defeat day*opsin category + intercept* | **$\beta$±standard error** | **z-stat** | **p-value** | **95% CI [lower, upper]** | |
|---|---|---|---|---|---|
| **Intercept** | 50.1998±1.932 | 25.983 | <0.001 | 46.413 | 53.987 |
| **Defeat day** | -0.1333±0.488 | -0.273 | 0.785 | -1.090 | 0.823 |
| **Opsin category** | -1.1085±2.775 | -0.399 | 0.690 | -6.548 | -4.331 |
| **Defeat day * Opsin category** | -0.4933±0.639 | -0.772 | 0.440 | -1.745 | 0.759 |

**Supplementary Data Table 55:** 2-sided GEE regression of behavior (**Figure S7F**, fourth panel): % time mice are fleeing during attack = defeat day + opsin (ChR2 or ChRmine) category + defeat day*opsin (ChR2 or ChRmine) category + intercept, grouped by mouse. Number of mice (groups) = 22, minimum samples per group 11, maximum samples per group 11, dependence structure = independence, family = Gaussian. NOTE: defeat day is coded as mean centered

| *Model: % time mice are* ***fleeing during attack*** *= defeat day + opsin (ChR2 or ChRmine) category + defeat day*opsin category + intercept* | **$\beta$±standard error** | **z-stat** | **p-value** | **95% CI [lower, upper]** | |
|---|---|---|---|---|---|
| **Intercept** | 11.4885±1.582 | 7.263 | <0.001 | 8.388 | 14.589 |
| **Defeat day** | -0.6205±0.226 | -2.740 | 0.006 | -1.064 | -0.177 |
| **Opsin category** | 16.3787±1.878 | 8.721 | <0.001 | 12.698 | 20.060 |

| | | | | | |
|---|---|---|---|---|---|
| **Defeat day * Opsin category** | 0.0676±0.610 | 0.111 | 0.912 | -1.128 | 1.263 |

**Supplementary Data Table 56:** 2-sided GEE regression of behavior (**Figure S7G**, first panel): % time mice are being attacked = defeat day + opsin (NpHr) category + defeat day*opsin (NpHr) category + intercept, grouped by mouse. Number of mice (groups) = 20, minimum samples per group 10, maximum samples per group 10, dependence structure = independence, family = Gaussian. NOTE: defeat day is coded as mean centered

| *Model: % time mice are **being attacked** = defeat day + opsin (NpHr) category + defeat day*opsin category + intercept* | **$\beta$±standard error** | **z-stat** | **p-value** | **95% CI [lower, upper]** | |
|---|---|---|---|---|---|
| **Intercept** | 10.2633±0.499 | 20.555 | <0.001 | 9.285 | 11.242 |
| **Defeat day** | -0.0721±0.170 | -0.423 | 0.672 | -0.406 | 0.262 |
| **Opsin category** | 0.2470±0.691 | 0.357 | 0.721 | -1.107 | 1.601 |
| **Defeat day * Opsin category** | -0.6185±0.283 | -2.187 | 0.029 | -1.173 | -0.064 |

**Supplementary Data Table 57:** 2-sided GEE regression of behavior (**Figure S7G**, second panel): % time mice are freezing during attack = defeat day + opsin (NpHr) category + defeat day*opsin (NpHr) category + intercept, grouped by mouse. Number of mice (groups) = 20, minimum samples per group 10, maximum samples per group 10, dependence structure = independence, family = Gaussian. NOTE: defeat day is coded as mean centered

| *Model: % time mice are **freezing during attack**= defeat day + opsin (NpHr) category + defeat day*opsin category + intercept* | **$\beta$±standard error** | **z-stat** | **p-value** | **95% CI [lower, upper]** | |
|---|---|---|---|---|---|
| **Intercept** | 22.4674±0.776 | 28.948 | <0.001 | 20.946 | 23.989 |
| **Defeat day** | -0.2212±0.378 | -0.585 | 0.558 | -0.962 | 0.520 |
| **Opsin category** | -0.2843±1.211 | -0.235 | 0.814 | -2.659 | 2.090 |
| **Defeat day * Opsin category** | 0.1604±0.437 | 0.367 | 0.714 | -0.696 | 1.017 |

**Supplementary Data Table 58:** 2-sided GEE regression of behavior (**Figure S7G**, third panel): % time mice are fighting during attack = defeat day + opsin (NpHr) category + defeat day*opsin (NpHr) category + intercept, grouped by mouse. Number of mice (groups) = 20, minimum samples per group 10, maximum samples per group 10, dependence structure = independence, family = Gaussian. NOTE: defeat day is coded as mean centered

| *Model: % time mice are **fighting during attack** = defeat day + opsin (NpHr) category + defeat day*opsin category + intercept* | **$\beta$±standard error** | **z-stat** | **p-value** | **95% CI [lower, upper]** | |
|---|---|---|---|---|---|
| **Intercept** | 50.8145±1.226 | 41.456 | <0.001 | 48.412 | 53.217 |
| **Defeat day** | -0.2843±0.414 | -0.687 | 0.492 | -1.096 | 0.527 |
| **Opsin category** | -4.5477±1.787 | -2.545 | 0.011 | -8.050 | -1.045 |
| **Defeat day * Opsin category** | -0.3011±0.690 | -0.436 | 0.663 | -1.653 | 1.051 |

**Supplementary Data Table 59**: 2-sided GEE regression of behavior (**Figure S7G**, fourth panel): % time mice are fleeing during attack = defeat day + opsin (NpHr) category + defeat day*opsin (NpHr) category + intercept, grouped by mouse. Number of mice (groups) = 20, minimum samples per group 10, maximum samples per group 10, dependence structure = independence, family = Gaussian. NOTE: defeat day is coded as mean centered

| *Model: % time mice are **fleeing during attack** = defeat day + opsin (NpHr) category + defeat day*opsin category + intercept* | **$\beta$±standard error** | **z-stat** | **p-value** | **95% CI [lower, upper]** | |
|---|---|---|---|---|---|
| **Intercept** | 36.2985±1.665 | 21.799 | <0.001 | 33.035 | 39.562 |
| **Defeat day** | -0.0859±0.403 | 3.788 | 0.831 | -0.875 | 0.704 |
| **Opsin category** | 10.1383±2.676 | 3.788 | <0.001 | 4.893 | 15.384 |
| **Defeat day * Opsin category** | 0.5224±0.651 | 0.803 | 0.422 | -0.753 | 1.798 |

**Supplementary Data Table 60:** GLM coefficient estimates of SI time contribution to $Fos^+$ cell counts in LHb-stimulated mice (**Figure 6E**). The total number of $Fos^+$ cells in each brain region was modeled as ***Counts ~ SI Time* + ln(*Total Counts*)** using a negative binomial link function (*N* = 10 mice). The critical *p*-value for significance while permitting a 10% FDR was 0.0371; 60 regions were significant.

| **Brain region name** | **SI time (Stim) coefficient estimate** | **SI time (Stim) coefficient std error** | **SI time (Stim) coefficient Z-statistic** | **SI time (Stim) coefficient P-value** |
|---|---|---|---|---|
| Frontal pole | -0.7314016 | 0.1849516 | -3.954556 | 7.667685E-05 |
| Primary motor area | 0.04163166 | 0.09308766 | 0.4472307 | 0.6547085 |
| Secondary motor area | 0.1555208 | 0.05654003 | 2.750633 | 0.00594803 |
| Primary somatosensory area, nose | 0.1739313 | 0.1633481 | 1.064789 | 0.2869714 |

| | | | | |
|---|---|---|---|---|
| Primary somatosensory area, barrel field | 0.2933146 | 0.1174378 | 2.497617 | 0.01250311 |
| Primary somatosensory area, lower limb | 0.1552762 | 0.1416929 | 1.095864 | 0.2731382 |
| Primary somatosensory area, mouth | -0.003168969 | 0.1861798 | -0.01702102 | 0.9864198 |
| Primary somatosensory area, upper limb | 0.08774347 | 0.1454138 | 0.6034052 | 0.5462391 |
| Primary somatosensory area, trunk | 0.1093963 | 0.09586774 | 1.141117 | 0.2538214 |
| Supplemental somatosensory area | 0.08872401 | 0.09056348 | 0.9796886 | 0.3272398 |
| Gustatory areas | -0.1694817 | 0.09013567 | -1.880295 | 0.06006789 |
| Visceral area | -0.03226184 | 0.09941656 | -0.3245117 | 0.7455506 |
| Dorsal auditory area | 0.1837334 | 0.1631325 | 1.126284 | 0.2600455 |
| Primary auditory area | -0.1018297 | 0.1245446 | -0.8176162 | 0.4135764 |
| Posterior auditory area | -0.03900642 | 0.3279325 | -0.1189465 | 0.9053177 |
| Ventral auditory area | -0.1349892 | 0.1301455 | -1.037218 | 0.2996345 |
| Anterolateral visual area | 0.6343466 | 0.2238639 | 2.833626 | 0.004602318 |
| Anteromedial visual area | 0.6566582 | 0.172218 | 3.812949 | 0.0001373186 |
| Lateral visual area | 0.7053728 | 0.2127889 | 3.314895 | 0.0009167755 |
| Primary visual area | 0.8620199 | 0.2476434 | 3.480892 | 0.000499747 |
| Posterolateral visual area | 0.3478742 | 0.08880988 | 3.917066 | 8.963309E-05 |
| Posteromedial visual area | 0.7418979 | 0.2214379 | 3.350365 | 0.0008070505 |
| Anterior cingulate area, dorsal part | 0.2382019 | 0.06744824 | 3.531625 | 0.0004130145 |
| Anterior cingulate area, ventral part | 0.3870767 | 0.101386 | 3.817853 | 0.000134618 |
| Prelimbic area | 0.002944094 | 0.1486476 | 0.01980587 | 0.9841982 |
| Infralimbic area | 0.02678873 | 0.09042469 | 0.2962545 | 0.7670357 |
| Orbital area, lateral part | 0.2886696 | 0.09369222 | 3.081042 | 0.002062779 |
| Orbital area, medial part | -0.2335652 | 0.1108752 | -2.106559 | 0.03515578 |
| Orbital area, ventrolateral part | 0.3141479 | 0.09798285 | 3.206152 | 0.001345228 |
| Agranular insular area, dorsal part | 0.1045462 | 0.06434629 | 1.624744 | 0.1042172 |
| Agranular insular area, posterior part | 0.2698059 | 0.08724678 | 3.092445 | 0.001985149 |
| Agranular insular area, ventral part | 0.0118411 | 0.07542433 | 0.1569931 | 0.8752503 |
| Retrosplenial area, lateral agranular part | 0.3265871 | 0.08658281 | 3.771962 | 0.0001619689 |
| Retrosplenial area, dorsal part | -0.1287764 | 0.06803493 | -1.892799 | 0.05838464 |
| Retrosplenial area, ventral part | -0.01986674 | 0.09742396 | -0.2039205 | 0.8384156 |
| Rostrolateral visual area | 0.4705157 | 0.1680059 | 2.800591 | 0.005100914 |
| Temporal association areas | -0.1244013 | 0.05688354 | -2.186947 | 0.0287464 |
| Perirhinal area | 0.0549359 | 0.08582782 | 0.640071 | 0.5221264 |
| Ectorhinal area | -0.1468615 | 0.09480798 | -1.549041 | 0.1213718 |
| Anterior olfactory nucleus | 0.2312797 | 0.08652238 | 2.673062 | 0.007516239 |
| Taenia tecta, dorsal part | 0.04222891 | 0.08560573 | 0.4932954 | 0.6218039 |
| Taenia tecta, ventral part | -0.5283536 | 0.08714838 | -6.06269 | 1.338636E-09 |
| Dorsal peduncular area | 0.1024163 | 0.1157033 | 0.8851632 | 0.3760686 |
| Piriform area | 0.3463955 | 0.1238715 | 2.796411 | 0.005167368 |
| Nucleus of the lateral olfactory tract | 0.5010421 | 0.4623802 | 1.083615 | 0.2785355 |
| Cortical amygdalar area, anterior part | 0.5264915 | 0.2789621 | 1.887323 | 0.05911693 |

| | | | | |
|---|---|---|---|---|
| Cortical amygdalar area, posterior part | 0.4102002 | 0.3063754 | 1.338881 | 0.1806094 |
| Piriform,amygdalar area | -0.03153026 | 0.1496777 | -0.2106543 | 0.833157 |
| Postpiriform transition area | 0.201428 | 0.221475 | 0.9094841 | 0.3630947 |
| Field c a1 | -0.03712589 | 0.09824932 | -0.3778743 | 0.705524 |
| Field c a2 | -0.2401251 | 0.1453749 | -1.651765 | 0.09858249 |
| Field c a3 | -0.02804974 | 0.1192571 | -0.235204 | 0.8140504 |
| Dentate gyrus | -0.2609662 | 0.179082 | -1.457244 | 0.1450491 |
| Induseum griseum | -0.5149479 | 0.4143951 | -1.24265 | 0.213997 |
| Entorhinal area, lateral part | 0.04921013 | 0.08151402 | 0.6037015 | 0.5460421 |
| Entorhinal area, medial part | 0.1822872 | 0.07023926 | 2.595233 | 0.009452694 |
| Parasubiculum | -0.3307718 | 0.2207432 | -1.498446 | 0.1340174 |
| Postsubiculum | 0.08865971 | 0.0917386 | 0.9664384 | 0.3338248 |
| Presubiculum | -0.05097407 | 0.06265621 | -0.8135517 | 0.4159018 |
| Subiculum | -0.2392685 | 0.1017158 | -2.352323 | 0.01865654 |
| Prosubiculum | 0.1064967 | 0.05515821 | 1.930749 | 0.05351405 |
| Claustrum | 0.1625206 | 0.1193206 | 1.362049 | 0.1731823 |
| Endopiriform nucleus, dorsal part | 0.4618091 | 0.172034 | 2.684407 | 0.007265867 |
| Endopiriform nucleus, ventral part | 0.2465357 | 0.1338811 | 1.841452 | 0.06555529 |
| Lateral amygdalar nucleus | -0.263296 | 0.08220707 | -3.202839 | 0.0013608 |
| Basolateral amygdalar nucleus, anterior part | 0.1695389 | 0.08099015 | 2.093327 | 0.03631998 |
| Basolateral amygdalar nucleus, posterior part | 0.009952845 | 0.09477041 | 0.1050206 | 0.9163595 |
| Basolateral amygdalar nucleus, ventral part | 0.4040623 | 0.199038 | 2.030076 | 0.0423488 |
| Basomedial amygdalar nucleus, anterior part | 0.4084054 | 0.331475 | 1.232085 | 0.2179172 |
| Basomedial amygdalar nucleus, posterior part | 0.1975485 | 0.2079677 | 0.9498999 | 0.3421631 |
| Posterior amygdalar nucleus | -0.003826692 | 0.3382347 | -0.01131372 | 0.9909732 |
| Caudoputamen | -0.1618055 | 0.09884296 | -1.636995 | 0.1016315 |
| Nucleus accumbens | 0.2504399 | 0.1404632 | 1.782957 | 0.07459337 |
| Fundus of striatum | -0.05841888 | 0.1443635 | -0.4046652 | 0.6857236 |
| Olfactory tubercle | -0.2113748 | 0.0780571 | -2.707951 | 0.006770007 |
| Lateral septal nucleus | 0.3688252 | 0.1956787 | 1.884851 | 0.05945 |
| Septofimbrial nucleus | 0.1653383 | 0.2965934 | 0.5574579 | 0.5772146 |
| Anterior amygdalar area | -0.2301262 | 0.306297 | -0.7513173 | 0.4524617 |
| Central amygdalar nucleus, capsular part | -0.3617941 | 0.1381454 | -2.618936 | 0.008820451 |
| Central amygdalar nucleus, lateral part | -0.08125791 | 0.1727744 | -0.4703122 | 0.638132 |
| Central amygdalar nucleus, medial part | 0.2951745 | 0.2584186 | 1.142234 | 0.2533569 |
| Intercalated amygdalar nucleus | 0.08395031 | 0.2868701 | 0.2926423 | 0.7697956 |
| Medial amygdalar nucleus | 0.2413205 | 0.4207641 | 0.5735292 | 0.5662864 |
| Globus pallidus, external segment | -0.2684994 | 0.1773076 | -1.514314 | 0.1299463 |
| Globus pallidus, internal segment | 0.3331863 | 0.2647918 | 1.258295 | 0.208285 |
| Substantia innominata | -0.1234805 | 0.188975 | -0.6534223 | 0.5134841 |
| Magnocellular nucleus | -0.5858663 | 0.2271826 | -2.578834 | 0.009913448 |
| Medial septal nucleus | -0.053557 | 0.1093459 | -0.4897945 | 0.6242793 |

| | | | | |
|---|---|---|---|---|
| Diagonal band nucleus | -0.2894237 | 0.09260241 | -3.125444 | 0.001775368 |
| Triangular nucleus of septum | -0.2000238 | 0.2949106 | -0.6782524 | 0.4976117 |
| Bed nuclei of the stria terminalis | 0.09172938 | 0.1893776 | 0.4843729 | 0.6281212 |
| Ventral anterior,lateral complex of the thalamus | 0.7189982 | 0.6031576 | 1.192057 | 0.2332389 |
| Ventral medial nucleus of the thalamus | -0.03125853 | 0.2737155 | -0.1142008 | 0.9090786 |
| Ventral posterior complex of the thalamus | -0.01730245 | 0.1953722 | -0.08856148 | 0.9294304 |
| Subparafascicular nucleus | 0.2526613 | 0.2880439 | 0.8771628 | 0.3803982 |
| Subparafascicular area | 0.07468876 | 0.5691948 | 0.1312183 | 0.8956026 |
| Medial geniculate complex | -0.6386775 | 0.1606624 | -3.975278 | 7.029721E-05 |
| Lateral geniculate complex | -0.3271515 | 0.0616582 | -5.305888 | 1.121259E-07 |
| Lateral posterior nucleus of the thalamus | -0.2818305 | 0.1223991 | -2.302554 | 0.02130394 |
| Posterior complex of the thalamus | -0.687739 | 0.1308011 | -5.257898 | 1.457115E-07 |
| Posterior limiting nucleus of the thalamus | -0.3971762 | 0.09588208 | -4.14234 | 3.437802E-05 |
| Suprageniculate nucleus | -0.591656 | 0.1464017 | -4.041318 | 5.315154E-05 |
| Anteroventral nucleus of thalamus | -0.01644025 | 0.3157823 | -0.05206196 | 0.9584793 |
| Anteromedial nucleus | 0.7094845 | 0.4114808 | 1.724223 | 0.08466765 |
| Anterodorsal nucleus | -0.3792642 | 0.1897503 | -1.998754 | 0.04563501 |
| Interanterodorsal nucleus of the thalamus | 0.9536466 | 0.3870404 | 2.463946 | 0.01374169 |
| Lateral dorsal nucleus of thalamus | -0.1590042 | 0.1378022 | -1.153858 | 0.2485582 |
| Intermediodorsal nucleus of the thalamus | -0.07109112 | 0.3637729 | -0.1954272 | 0.8450585 |
| Mediodorsal nucleus of thalamus | -0.02925827 | 0.1744685 | -0.1676994 | 0.8668198 |
| Submedial nucleus of the thalamus | -1.274216 | 0.4920157 | -2.589787 | 0.009603522 |
| Perireunensis nucleus | -0.1117313 | 0.3050968 | -0.366216 | 0.7142039 |
| Paraventricular nucleus of the thalamus | -0.1897655 | 0.1825975 | -1.039256 | 0.2986857 |
| Parataenial nucleus | -0.004615776 | 0.1795629 | -0.02570562 | 0.9794921 |
| Nucleus of reuniens | -0.2315966 | 0.2758274 | -0.8396434 | 0.4011083 |
| Central medial nucleus of the thalamus | 0.4061229 | 0.3795686 | 1.069959 | 0.2846377 |
| Paracentral nucleus | -0.651202 | 0.299325 | -2.175569 | 0.02958754 |
| Central lateral nucleus of the thalamus | 0.04285389 | 0.1741498 | 0.2460749 | 0.8056242 |
| Parafascicular nucleus | -0.12695 | 0.1534304 | -0.827411 | 0.4080042 |
| Reticular nucleus of the thalamus | -0.2722539 | 0.184447 | -1.476055 | 0.1399291 |
| Medial habenula | -0.9738178 | 0.1580583 | -6.161129 | 7.222799E-10 |
| Lateral habenula | -0.2497861 | 0.2279821 | -1.095639 | 0.2732368 |
| Paraventricular hypothalamic nucleus | 0.3274925 | 0.2603035 | 1.258118 | 0.2083492 |
| Periventricular hypothalamic nucleus | -0.357596 | 0.3900434 | -0.9168109 | 0.3592417 |
| Arcuate hypothalamic nucleus | -0.4339995 | 0.576894 | -0.7523037 | 0.4518684 |
| Anterodorsal preoptic nucleus | 0.0884651 | 0.2308219 | 0.3832613 | 0.701526 |
| Anteroventral periventricular nucleus | -0.06808806 | 0.2948822 | -0.2308992 | 0.8173931 |
| Dorsomedial hypothalamic nucleus | -0.2885431 | 0.4366378 | -0.6608294 | 0.5087217 |
| Medial preoptic area | 0.07462574 | 0.1813418 | 0.4115198 | 0.6806914 |
| Subparaventricular zone | 0.2884794 | 0.5601746 | 0.5149812 | 0.6065662 |
| Anterior hypothalamic nucleus | 0.1542997 | 0.3432005 | 0.4495905 | 0.6530057 |

| | | | | |
|---|---|---|---|---|
| Supramammillary nucleus | -0.2029339 | 0.6067278 | -0.3344727 | 0.7380229 |
| Tuberomammillary nucleus | 0.4585401 | 0.507093 | 0.9042525 | 0.3658615 |
| Medial preoptic nucleus | -0.2770029 | 0.423436 | -0.654179 | 0.5129965 |
| Dorsal premammillary nucleus | -0.3149158 | 0.6236127 | -0.5049861 | 0.6135686 |
| Ventral premammillary nucleus | 0.1237997 | 0.7424729 | 0.1667397 | 0.8675749 |
| Ventromedial hypothalamic nucleus | 0.07014335 | 0.5456631 | 0.128547 | 0.8977161 |
| Posterior hypothalamic nucleus | -0.119619 | 0.4546849 | -0.263081 | 0.7924881 |
| Lateral hypothalamic area | 0.07519841 | 0.2198425 | 0.3420558 | 0.7323089 |
| Lateral preoptic area | 0.08505081 | 0.1540873 | 0.5519652 | 0.5809722 |
| Parasubthalamic nucleus | 0.2688046 | 0.4242063 | 0.6336649 | 0.5262995 |
| Retrochiasmatic area | 0.3402017 | 0.5642826 | 0.6028924 | 0.5465802 |
| Subthalamic nucleus | 0.1497166 | 0.2877702 | 0.5202644 | 0.6028793 |
| Tuberal nucleus | 0.3094244 | 0.4305652 | 0.7186469 | 0.4723585 |
| Zona incerta | 0.04947893 | 0.1237397 | 0.3998632 | 0.6892573 |
| Superior colliculus, dorsal part | -0.06146608 | 0.1076395 | -0.5710366 | 0.5679748 |
| Inferior colliculus, central part | -0.4139421 | 0.1458797 | -2.837558 | 0.004546014 |
| Inferior colliculus, dorsal part | -0.4393368 | 0.1213334 | -3.620904 | 0.0002935751 |
| Inferior colliculus, external part | -0.3223625 | 0.1295522 | -2.488282 | 0.01283618 |
| Substantia nigra, reticular part | -0.4276571 | 0.08645127 | -4.946799 | 7.544366E-07 |
| Ventral tegmental area | 0.02059024 | 0.4833655 | 0.04259765 | 0.9660223 |
| Midbrain reticular nucleus | -0.159509 | 0.1203988 | -1.324839 | 0.1852246 |
| Superior colliculus, ventral part | -0.2097347 | 0.103546 | -2.025522 | 0.04281375 |
| Periaqueductal gray | -0.105694 | 0.09600211 | -1.100955 | 0.2709163 |
| Anterior pretectal nucleus | -0.2243703 | 0.1958359 | -1.145706 | 0.251917 |
| Nucleus of the optic tract | -0.4385215 | 0.1817514 | -2.412755 | 0.01583244 |
| Nucleus of the posterior commissure | -0.214548 | 0.2897952 | -0.7403434 | 0.4590916 |
| Posterior pretectal nucleus | -0.3123867 | 0.1088459 | -2.869989 | 0.004104861 |
| Cuneiform nucleus | -0.05762814 | 0.08824413 | -0.6530535 | 0.5137218 |
| Red nucleus | 0.01148355 | 0.2410147 | 0.04764669 | 0.9619978 |
| Substantia nigra, compact part | 0.07573934 | 0.2282799 | 0.3317828 | 0.7400533 |
| Pedunculopontine nucleus | -0.2829347 | 0.07449492 | -3.79804 | 0.000145845 |
| Dorsal nucleus raphe | -0.2313467 | 0.2557011 | -0.9047544 | 0.3655955 |
| Nucleus of the lateral lemniscus | 0.02620023 | 0.1455483 | 0.1800105 | 0.8571443 |
| Principal sensory nucleus of the trigeminal | 0.755922 | 0.2141155 | 3.53044 | 0.0004148688 |
| Parabrachial nucleus | -0.4492131 | 0.121283 | -3.703843 | 0.0002123577 |
| Superior olivary complex | -0.1565898 | 0.1507921 | -1.038449 | 0.2990613 |
| Dorsal tegmental nucleus | -0.8018735 | 0.2532351 | -3.166518 | 0.001542757 |
| Pontine central gray | -0.7485026 | 0.1465017 | -5.109175 | 3.235684E-07 |
| Pontine gray | 0.7944909 | 0.38337 | 2.072387 | 0.03822937 |
| Supratrigeminal nucleus | -0.3950062 | 0.1556073 | -2.538482 | 0.01113347 |
| Tegmental reticular nucleus | 0.2543683 | 0.4805386 | 0.5293399 | 0.5965697 |
| Motor nucleus of trigeminal | 0.1256878 | 0.14708 | 0.8545541 | 0.3927981 |

| | | | | |
|---|---|---|---|---|
| Superior central nucleus raphe | 0.3142269 | 0.3040467 | 1.033483 | 0.3013781 |
| Laterodorsal tegmental nucleus | -0.5445012 | 0.1984766 | -2.743403 | 0.006080606 |
| Nucleus incertus | -0.3351602 | 0.1907434 | -1.757126 | 0.0788963 |
| Pontine reticular nucleus | 0.03717075 | 0.1130418 | 0.3288231 | 0.7422894 |
| Dorsal cochlear nucleus | -0.1814855 | 0.1413664 | -1.283795 | 0.1992138 |
| Ventral cochlear nucleus | -0.3561585 | 0.2265167 | -1.572327 | 0.1158747 |
| Cuneate nucleus | 0.1379057 | 0.3303435 | 0.4174614 | 0.6763409 |
| External cuneate nucleus | -0.1333319 | 0.174199 | -0.7653996 | 0.4440337 |
| Nucleus of the trapezoid body | -0.02324342 | 0.346784 | -0.06702565 | 0.9465613 |
| Nucleus of the solitary tract | -0.8231558 | 0.2081215 | -3.95517 | 7.648038E-05 |
| Spinal nucleus of the trigeminal | -0.2839096 | 0.1361867 | -2.08471 | 0.03709565 |
| Facial motor nucleus | 0.2392683 | 0.2172025 | 1.101591 | 0.2706394 |
| Dorsal motor nucleus of the vagus nerve | -0.7546433 | 0.2195987 | -3.436465 | 0.0005893582 |
| Gigantocellular reticular nucleus | 0.4516057 | 0.1702198 | 2.653074 | 0.007976233 |
| Inferior olivary complex | 0.07347595 | 0.2635633 | 0.2787791 | 0.7804143 |
| Intermediate reticular nucleus | -0.3049118 | 0.1743223 | -1.749127 | 0.08026907 |
| Lateral reticular nucleus | 0.5570417 | 0.25824 | 2.15707 | 0.03100022 |
| Magnocellular reticular nucleus | 1.123009 | 0.6447175 | 1.741861 | 0.08153272 |
| Medullary reticular nucleus | -0.3416906 | 0.1960752 | -1.742651 | 0.08139459 |
| Parvicellular reticular nucleus | -0.2797066 | 0.1534423 | -1.822879 | 0.06832176 |
| Paragigantocellular reticular nucleus | 0.02005862 | 0.1186495 | 0.1690578 | 0.8657512 |
| Nucleus prepositus | 0.1887702 | 0.1598303 | 1.181067 | 0.2375762 |
| Lateral vestibular nucleus | -0.06541776 | 0.257114 | -0.2544309 | 0.7991627 |
| Medial vestibular nucleus | -0.4056525 | 0.07463156 | -5.435402 | 5.46731E-08 |
| Spinal vestibular nucleus | -0.114425 | 0.1154098 | -0.9914672 | 0.3214575 |
| Superior vestibular nucleus | -0.1221085 | 0.2781986 | -0.4389256 | 0.6607155 |
| Hypoglossal nucleus | -0.459039 | 0.1826233 | -2.513585 | 0.0119511 |
| Nucleus raphe magnus | 0.2526663 | 0.4194547 | 0.6023685 | 0.5469289 |

**Supplementary Data Table 61:** GLMM coefficient estimates of SI time contribution to Fos$^{+}$ cell counts in unstimulated control mice (**Figure 6E**). The total number of Fos$^{+}$ cells in each brain region was modeled as ***Counts ~ SI Time* + ln(*Total Counts*) + (*1+SI Time*|*Cohort*)** using a negative binomial link function (*N* = 44 mice). The critical *p*-value for significance while permitting a 10% FDR was 0.000068; 1 region was significant.

| **Brain region name** | **SI time (NoStim) coefficient estimate** | **SI time (NoStim) coefficient std error** | **SI time (NoStim) coefficient Z-statistic** | **SI time (NoStim) coefficient P-value** |
|---|---|---|---|---|
| Frontal pole | -0.1244752 | 0.2070876 | -0.6010749 | 0.5477901 |
| Primary motor area | 0.03187053 | 0.0850538 | 0.3747102 | 0.707876 |
| Secondary motor area | 0.006213008 | 0.04968765 | 0.1250413 | 0.9004908 |

| | | | | |
|---|---|---|---|---|
| Primary somatosensory area, nose | -0.1219617 | 0.102128 | -1.194205 | 0.2323979 |
| Primary somatosensory area, barrel field | 0.01697556 | 0.09205788 | 0.1844009 | 0.853699 |
| Primary somatosensory area, lower limb | 0.1107461 | 0.1209749 | 0.9154467 | 0.3599572 |
| Primary somatosensory area, mouth | -0.1141899 | 0.1215792 | -0.9392227 | 0.3476164 |
| Primary somatosensory area, upper limb | 0.01778997 | 0.09557645 | 0.1861335 | 0.8523401 |
| Primary somatosensory area, trunk | 0.03059643 | 0.07019274 | 0.4358917 | 0.6629153 |
| Supplemental somatosensory area | -0.0322329 | 0.07818083 | -0.4122865 | 0.6801294 |
| Gustatory areas | 0.02235067 | 0.1233906 | 0.1811376 | 0.8562596 |
| Visceral area | -0.09042612 | 0.08649857 | -1.045406 | 0.2958354 |
| Dorsal auditory area | 0.004810851 | 0.1083562 | 0.0443985 | 0.9645868 |
| Primary auditory area | 0.01416738 | 0.1300086 | 0.1089726 | 0.9132242 |
| Posterior auditory area | -0.1433355 | 0.2773161 | -0.5168668 | 0.6052491 |
| Ventral auditory area | -0.036433 | 0.09338198 | -0.3901502 | 0.6964255 |
| Anterolateral visual area | -0.05749006 | 0.221683 | -0.2593346 | 0.7953771 |
| Anteromedial visual area | -0.04041426 | 0.1257587 | -0.3213636 | 0.7479349 |
| Lateral visual area | -0.0401709 | 0.2467249 | -0.1628166 | 0.8706628 |
| Primary visual area | -0.02554216 | 0.1822249 | -0.1401683 | 0.888527 |
| Posterolateral visual area | -0.06485103 | 0.2312065 | -0.2804896 | 0.7791019 |
| Posteromedial visual area | -0.07899774 | 0.141031 | -0.5601446 | 0.5753808 |
| Anterior cingulate area, dorsal part | -0.01151697 | 0.04599909 | -0.2503739 | 0.8022982 |
| Anterior cingulate area, ventral part | 0.03874381 | 0.04237275 | 0.9143566 | 0.3605295 |
| Prelimbic area | 0.009497338 | 0.07445022 | 0.1275663 | 0.8984922 |
| Infralimbic area | -0.04221049 | 0.07770423 | -0.5432199 | 0.5869784 |
| Orbital area, lateral part | -0.01395659 | 0.07087148 | -0.1969282 | 0.8438838 |
| Orbital area, medial part | 0.007382164 | 0.105216 | 0.07016202 | 0.9440647 |
| Orbital area, ventrolateral part | -0.06707003 | 0.09459961 | -0.7089884 | 0.4783317 |
| Agranular insular area, dorsal part | 0.07643315 | 0.1005513 | 0.7601411 | 0.4471702 |
| Agranular insular area, posterior part | -0.05742976 | 0.09493345 | -0.6049476 | 0.5452138 |
| Agranular insular area, ventral part | 0.05861033 | 0.1037669 | 0.5648268 | 0.5721916 |
| Retrosplenial area, lateral agranular part | -0.0447323 | 0.1178547 | -0.3795545 | 0.7042761 |
| Retrosplenial area, dorsal part | 0.02324616 | 0.1165858 | 0.199391 | 0.8419569 |
| Retrosplenial area, ventral part | 0.0475773 | 0.05381408 | 0.8841051 | 0.3766395 |
| Rostrolateral visual area | -0.02220282 | 0.1376004 | -0.1613572 | 0.8718121 |
| Temporal association areas | -0.0211156 | 0.1446013 | -0.1460264 | 0.8839006 |
| Perirhinal area | -0.1960756 | 0.1601596 | -1.224251 | 0.2208574 |
| Ectorhinal area | -0.1185687 | 0.1150942 | -1.030189 | 0.3029215 |
| Anterior olfactory nucleus | -0.02846436 | 0.1654752 | -0.1720158 | 0.8634251 |
| Taenia tecta, dorsal part | -0.006718019 | 0.06165789 | -0.1089564 | 0.9132371 |
| Taenia tecta, ventral part | -0.04903991 | 0.2190231 | -0.223903 | 0.8228328 |
| Dorsal peduncular area | -0.08993022 | 0.08503945 | -1.057512 | 0.2902781 |
| Piriform area | -0.05072274 | 0.1373249 | -0.3693629 | 0.7118572 |
| Nucleus of the lateral olfactory tract | -0.5964235 | 0.5195186 | -1.148031 | 0.2509557 |

| | | | | |
|---|---|---|---|---|
| Cortical amygdalar area, anterior part | -1.034304 | 0.8728108 | -1.185027 | 0.2360068 |
| Cortical amygdalar area, posterior part | -0.2738256 | 0.4129487 | -0.6630983 | 0.5072676 |
| Piriform,amygdalar area | -0.3792913 | 0.4180772 | -0.9072278 | 0.3642863 |
| Postpiriform transition area | -0.3633383 | 0.2373998 | -1.530491 | 0.1258953 |
| Field c a1 | 0.05705185 | 0.0459914 | 1.24049 | 0.2147944 |
| Field c a2 | -0.03171736 | 0.0717703 | -0.4419288 | 0.6585407 |
| Field c a3 | 0.008740511 | 0.06273482 | 0.1393247 | 0.8891936 |
| Dentate gyrus | 0.004960057 | 0.07044779 | 0.07040757 | 0.9438693 |
| Induseum griseum | -0.06895816 | 0.1057855 | -0.6518676 | 0.5144866 |
| Entorhinal area, lateral part | -0.07016447 | 0.2328053 | -0.3013869 | 0.7631195 |
| Entorhinal area, medial part | -0.02382059 | 0.08625518 | -0.2761642 | 0.7824219 |
| Parasubiculum | -0.002766915 | 0.08663841 | -0.03193636 | 0.9745228 |
| Postsubiculum | 0.06678519 | 0.06161419 | 1.083925 | 0.2783978 |
| Presubiculum | 0.08288441 | 0.06061536 | 1.367383 | 0.1715053 |
| Subiculum | -0.0470562 | 0.06897867 | -0.6821848 | 0.4951221 |
| Prosubiculum | -0.01057175 | 0.05412897 | -0.1953066 | 0.8451529 |
| Claustrum | 0.04410515 | 0.05381203 | 0.8196151 | 0.4124355 |
| Endopiriform nucleus, dorsal part | 0.08595679 | 0.1343619 | 0.6397409 | 0.5223411 |
| Endopiriform nucleus, ventral part | -0.4706728 | 0.1908692 | -2.465945 | 0.01366523 |
| Lateral amygdalar nucleus | 0.006022568 | 0.1060278 | 0.0568018 | 0.9547031 |
| Basolateral amygdalar nucleus, anterior part | -0.07986055 | 0.1591652 | -0.5017464 | 0.6158459 |
| Basolateral amygdalar nucleus, posterior part | -0.1813625 | 0.2128445 | -0.8520896 | 0.3941644 |
| Basolateral amygdalar nucleus, ventral part | -0.815103 | 0.8001528 | -1.018684 | 0.3083529 |
| Basomedial amygdalar nucleus, anterior part | -0.5041722 | 0.40067 | -1.258323 | 0.208275 |
| Basomedial amygdalar nucleus, posterior part | -0.3054298 | 0.1999838 | -1.527273 | 0.1266932 |
| Posterior amygdalar nucleus | -0.0302394 | 0.2273958 | -0.1329813 | 0.8942082 |
| Caudoputamen | 0.05868319 | 0.07011798 | 0.8369208 | 0.4026371 |
| Nucleus accumbens | 0.04271247 | 0.09669703 | 0.4417144 | 0.6586959 |
| Fundus of striatum | -0.02337132 | 0.1060585 | -0.2203626 | 0.8255888 |
| Olfactory tubercle | -0.1334093 | 0.2802409 | -0.4760522 | 0.6340372 |
| Lateral septal nucleus | -0.01417287 | 0.07222929 | -0.1962206 | 0.8444375 |
| Septofimbrial nucleus | -0.06522049 | 0.0598274 | -1.090144 | 0.2756497 |
| Anterior amygdalar area | -0.4249364 | 0.1458051 | -2.914413 | 0.00356358 |
| Central amygdalar nucleus, capsular part | -0.1316331 | 0.1315279 | -1.0008 | 0.3169235 |
| Central amygdalar nucleus, lateral part | -0.1733695 | 0.1129924 | -1.534347 | 0.1249443 |
| Central amygdalar nucleus, medial part | -0.1403542 | 0.1338931 | -1.048256 | 0.2945208 |
| Intercalated amygdalar nucleus | -0.2171916 | 0.2027443 | -1.071259 | 0.2840531 |
| Medial amygdalar nucleus | -0.130111 | 0.1734786 | -0.7500115 | 0.4532478 |
| Globus pallidus, external segment | -0.009954849 | 0.09719895 | -0.1024172 | 0.9184255 |
| Globus pallidus, internal segment | -0.06736114 | 0.1444259 | -0.4664061 | 0.6409248 |
| Substantia innominata | 0.006613689 | 0.09228287 | 0.07166757 | 0.9428665 |
| Magnocellular nucleus | -0.1579383 | 0.127245 | -1.241215 | 0.2145264 |

| | | | | |
|---|---|---|---|---|
| Medial septal nucleus | -0.1342138 | 0.07305477 | -1.837166 | 0.06618537 |
| Diagonal band nucleus | -0.1440691 | 0.1088528 | -1.323522 | 0.1856618 |
| Triangular nucleus of septum | 0.02207067 | 0.06609697 | 0.3339134 | 0.7384449 |
| Bed nuclei of the stria terminalis | -0.04458337 | 0.07231628 | -0.6165052 | 0.5375611 |
| Ventral anterior,lateral complex of the thalamus | -0.03250431 | 0.09408716 | -0.3454702 | 0.7297409 |
| Ventral medial nucleus of the thalamus | -0.04301005 | 0.08166001 | -0.5266966 | 0.5984043 |
| Ventral posterior complex of the thalamus | 0.0427066 | 0.06868881 | 0.6217404 | 0.5341126 |
| Subparafascicular nucleus | -0.05652446 | 0.07458612 | -0.7578415 | 0.4485459 |
| Subparafascicular area | -0.09364922 | 0.140408 | -0.666979 | 0.5047855 |
| Medial geniculate complex | 0.06644385 | 0.06855351 | 0.9692261 | 0.3324324 |
| Lateral geniculate complex | 0.01888782 | 0.04640382 | 0.4070316 | 0.6839848 |
| Lateral posterior nucleus of the thalamus | 0.05898301 | 0.06174132 | 0.9553248 | 0.3394135 |
| Posterior complex of the thalamus | 0.2421485 | 0.1362463 | 1.777284 | 0.07552144 |
| Posterior limiting nucleus of the thalamus | -0.1000042 | 0.11951 | -0.836785 | 0.4027134 |
| Suprageniculate nucleus | -0.07668891 | 0.08046049 | -0.9531251 | 0.3405267 |
| Anteroventral nucleus of thalamus | -0.004098541 | 0.0974158 | -0.04207265 | 0.9664408 |
| Anteromedial nucleus | -0.1659288 | 0.1164348 | -1.425079 | 0.1541343 |
| Anterodorsal nucleus | -0.1714044 | 0.1049446 | -1.633284 | 0.1024093 |
| Interanterodorsal nucleus of the thalamus | -0.04998603 | 0.1039893 | -0.4806845 | 0.6307407 |
| Lateral dorsal nucleus of thalamus | 0.09467539 | 0.1194365 | 0.7926837 | 0.4279621 |
| Intermediodorsal nucleus of the thalamus | -0.2147501 | 0.1144995 | -1.875554 | 0.06071654 |
| Mediodorsal nucleus of thalamus | -0.08001558 | 0.06739608 | -1.187244 | 0.2351315 |
| Submedial nucleus of the thalamus | -0.00274229 | 0.1129606 | -0.02427651 | 0.980632 |
| Perireunensis nucleus | -0.07694016 | 0.1055337 | -0.7290576 | 0.4659664 |
| Paraventricular nucleus of the thalamus | -0.03417365 | 0.06399778 | -0.5339818 | 0.5933541 |
| Parataenial nucleus | -0.07444211 | 0.06667198 | -1.116543 | 0.26419 |
| Nucleus of reuniens | -0.07105897 | 0.07595245 | -0.9355718 | 0.3494937 |
| Central medial nucleus of the thalamus | -0.06987667 | 0.1030239 | -0.6782566 | 0.497609 |
| Paracentral nucleus | -0.02032759 | 0.1146871 | -0.177244 | 0.8593168 |
| Central lateral nucleus of the thalamus | 0.08014783 | 0.09479059 | 0.8455252 | 0.3978177 |
| Parafascicular nucleus | 0.03781085 | 0.1239704 | 0.3049991 | 0.7603669 |
| Reticular nucleus of the thalamus | -0.1133894 | 0.09337151 | -1.214389 | 0.2245991 |
| Medial habenula | 0.5142396 | 0.1291072 | 3.983042 | 6.803877E-05 |
| Lateral habenula | 0.03461826 | 0.09246247 | 0.3744033 | 0.7081043 |
| Paraventricular hypothalamic nucleus | -0.02014541 | 0.11002708 | -0.183095 | 0.8547235 |
| Periventricular hypothalamic nucleus | 0.07804844 | 0.1397465 | 0.5585003 | 0.5765028 |
| Arcuate hypothalamic nucleus | -0.473564 | 0.6318556 | -0.7494814 | 0.4535671 |
| Anterodorsal preoptic nucleus | -0.08999211 | 0.09710568 | -0.9267441 | 0.3540594 |
| Anteroventral periventricular nucleus | -0.1258881 | 0.1556587 | -0.8087445 | 0.4186621 |
| Dorsomedial hypothalamic nucleus | -0.006132622 | 0.1100541 | -0.05572372 | 0.9555619 |
| Medial preoptic area | -0.112921 | 0.09231301 | -1.22324 | 0.221239 |
| Subparaventricular zone | -0.113467 | 0.1209568 | -0.9380782 | 0.3482042 |

| | | | | |
|---|---|---|---|---|
| Anterior hypothalamic nucleus | -0.06867683 | 0.1136775 | -0.6041374 | 0.5457523 |
| Supramammillary nucleus | -0.0865908 | 0.1638546 | -0.5284612 | 0.5971793 |
| Tuberomammillary nucleus | -0.2289766 | 0.3577237 | -0.6400936 | 0.5221118 |
| Medial preoptic nucleus | -0.07412443 | 0.09617689 | -0.7707094 | 0.4408792 |
| Dorsal premammillary nucleus | -0.08034717 | 0.1842738 | -0.4360207 | 0.6628217 |
| Ventral premammillary nucleus | 0.0470095 | 0.4341849 | 0.1082707 | 0.913781 |
| Ventromedial hypothalamic nucleus | -0.1151733 | 0.2170411 | -0.5306522 | 0.5956598 |
| Posterior hypothalamic nucleus | -0.08743613 | 0.1194955 | -0.7317107 | 0.4643452 |
| Lateral hypothalamic area | -0.025912 | 0.1298872 | -0.1994962 | 0.8418747 |
| Lateral preoptic area | -0.1414283 | 0.09497951 | -1.48904 | 0.1364768 |
| Parasubthalamic nucleus | -0.0226905 | 0.1535515 | -0.1477713 | 0.8825233 |
| Retrochiasmatic area | -0.4026865 | 0.6309765 | -0.6381958 | 0.5233462 |
| Subthalamic nucleus | -0.0857626 | 0.1348378 | -0.6360429 | 0.5247484 |
| Tuberal nucleus | -0.04375665 | 0.3376577 | -0.1295888 | 0.8968918 |
| Zona incerta | -0.03918807 | 0.071961 | -0.5445738 | 0.5860467 |
| Superior colliculus, dorsal part | 0.06460688 | 0.08345984 | 0.7741074 | 0.4388673 |
| Inferior colliculus, central part | 0.07671999 | 0.09017196 | 0.8508188 | 0.39487 |
| Inferior colliculus, dorsal part | 0.009863801 | 0.1347811 | 0.07318385 | 0.9416598 |
| Inferior colliculus, external part | 0.1195571 | 0.07719087 | 1.54885 | 0.1214177 |
| Substantia nigra, reticular part | -0.0411861 | 0.1188127 | -0.3466472 | 0.7288564 |
| Ventral tegmental area | -0.374182 | 0.3278732 | -1.14124 | 0.2537701 |
| Midbrain reticular nucleus | -0.06506472 | 0.1006753 | -0.6462832 | 0.518096 |
| Superior colliculus, ventral part | 0.001278192 | 0.06042271 | 0.02115417 | 0.9831227 |
| Periaqueductal gray | 0.008944951 | 0.07455235 | 0.1199821 | 0.9044973 |
| Anterior pretectal nucleus | 0.03531751 | 0.0796308 | 0.4435157 | 0.6573928 |
| Nucleus of the optic tract | -0.001803136 | 0.08450648 | -0.02133726 | 0.9829766 |
| Nucleus of the posterior commissure | 0.1254446 | 0.0897448 | 1.397793 | 0.1621753 |
| Posterior pretectal nucleus | 0.002127863 | 0.07532883 | 0.02824766 | 0.9774646 |
| Cuneiform nucleus | 0.005843659 | 0.09452633 | 0.06182044 | 0.9507058 |
| Red nucleus | -0.1754132 | 0.1046026 | -1.676949 | 0.09355247 |
| Substantia nigra, compact part | -0.1238306 | 0.1573336 | -0.7870578 | 0.431248 |
| Pedunculopontine nucleus | -0.06988815 | 0.1156845 | -0.6041271 | 0.5457591 |
| Dorsal nucleus raphe | -0.005675236 | 0.1228326 | -0.04620301 | 0.9631484 |
| Nucleus of the lateral lemniscus | 0.05119508 | 0.1028265 | 0.4978782 | 0.6185699 |
| Principal sensory nucleus of the trigeminal | 0.05275123 | 0.1416626 | 0.3723723 | 0.7096157 |
| Parabrachial nucleus | -0.07095405 | 0.1162609 | -0.6103 | 0.5416631 |
| Superior olivary complex | -0.2970388 | 0.1823838 | -1.628647 | 0.1033878 |
| Dorsal tegmental nucleus | -0.02346846 | 0.1745765 | -0.1344308 | 0.8930619 |
| Pontine central gray | -0.006636479 | 0.139774 | -0.04748007 | 0.9621306 |
| Pontine gray | -0.07550478 | 0.1630392 | -0.4631082 | 0.6432868 |
| Supratrigeminal nucleus | -0.04206037 | 0.1256335 | -0.3347862 | 0.7377864 |
| Tegmental reticular nucleus | -0.7681638 | 0.6876721 | -1.117049 | 0.2639732 |

| | | | | |
|---|---|---|---|---|
| Motor nucleus of trigeminal | -0.08232653 | 0.1658703 | -0.4963307 | 0.6196611 |
| Superior central nucleus raphe | -0.6374828 | 0.5742885 | -1.110039 | 0.2669821 |
| Laterodorsal tegmental nucleus | -0.06353252 | 0.1599744 | -0.3971418 | 0.6912629 |
| Nucleus incertus | 0.0197115 | 0.11397266 | 0.1729494 | 0.8626912 |
| Pontine reticular nucleus | -0.2202817 | 0.2349622 | -0.9375198 | 0.3484912 |
| Dorsal cochlear nucleus | 0.05782994 | 0.114564 | 0.5047829 | 0.6137113 |
| Ventral cochlear nucleus | 0.1658639 | 0.139262 | 1.19102 | 0.2336455 |
| Cuneate nucleus | 0.2773747 | 0.2413588 | 1.149222 | 0.2504646 |
| External cuneate nucleus | 0.001148166 | 0.2029795 | 0.005656559 | 0.9954867 |
| Nucleus of the trapezoid body | -0.4960915 | 0.4587105 | -1.081492 | 0.2794785 |
| Nucleus of the solitary tract | 0.02958777 | 0.1638275 | 0.1806033 | 0.856679 |
| Spinal nucleus of the trigeminal | -0.008767952 | 0.1655691 | -0.05295645 | 0.9577666 |
| Facial motor nucleus | -0.05334312 | 0.169067 | -0.3155147 | 0.7523709 |
| Dorsal motor nucleus of the vagus nerve | 0.1481606 | 0.2915792 | 0.5081315 | 0.6113612 |
| Gigantocellular reticular nucleus | 0.1069361 | 0.1546492 | 0.6914756 | 0.4892667 |
| Inferior olivary complex | 0.1735732 | 0.5340075 | 0.3250389 | 0.7451516 |
| Intermediate reticular nucleus | 0.03933921 | 0.1594241 | 0.2467583 | 0.8050953 |
| Lateral reticular nucleus | 0.3113005 | 0.3034911 | 1.025732 | 0.3050179 |
| Magnocellular reticular nucleus | 0.01150378 | 0.2978956 | 0.03861682 | 0.9691959 |
| Medullary reticular nucleus | 0.07775043 | 0.1674628 | 0.4642848 | 0.6424437 |
| Parvicellular reticular nucleus | 0.04765496 | 0.1208028 | 0.3944854 | 0.6932227 |
| Paragigantocellular reticular nucleus | 0.1189127 | 0.1576135 | 0.7544575 | 0.4505745 |
| Nucleus prepositus | 0.1826923 | 0.1641877 | 1.112704 | 0.2658357 |
| Lateral vestibular nucleus | -0.05145628 | 0.1333934 | -0.3857484 | 0.699683 |
| Medial vestibular nucleus | 0.08411719 | 0.1222016 | 0.6883479 | 0.4912337 |
| Spinal vestibular nucleus | -0.007403631 | 0.09990784 | -0.07410461 | 0.9409271 |
| Superior vestibular nucleus | -0.2026147 | 0.216005 | -0.9380094 | 0.3482396 |
| Hypoglossal nucleus | 0.380394 | 0.1950168 | 1.950571 | 0.05110815 |
| Nucleus raphe magnus | 0.160579 | 0.3169754 | 0.5065977 | 0.6124371 |

**Supplementary Data Table 62:** GLMM coefficient estimates of LHb stimulation contribution to Fos$^+$ cell counts across all mice (**Figure 6E**). The total number of Fos$^+$ cells in each brain region was modeled as ***Counts ~ NoStim* + ln(*Total Counts*) + (*1*|*Cohort*)** using a negative binomial link function ($N$ = 54 mice). The critical $p$-value for significance while permitting a 10% FDR was 0.0108; 24 regions were significant.

| Brain region name | SI time (NoStim–Stim) coefficient estimate | SI time (NoStim–Stim) coefficient std error | SI time (NoStim–Stim) coefficient Z-statistic | SI time (NoStim–Stim) coefficient P-value |
|---|---|---|---|---|
| Frontal pole | -1.448516 | 0.5927378 | -2.443773 | 0.01453459 |
| Primary motor area | 0.5884432 | 0.2120615 | 2.77487 | 0.005522378 |

| | | | | |
|---|---|---|---|---|
| Secondary motor area | 0.3268285 | 0.10068718 | 3.245979 | 0.001170475 |
| Primary somatosensory area, nose | 0.4876159 | 0.3381211 | 1.442134 | 0.1492646 |
| Primary somatosensory area, barrel field | 0.54962 | 0.1711741 | 3.210883 | 0.00132328 |
| Primary somatosensory area, lower limb | 0.731869 | 0.2071353 | 3.53329 | 0.0004104227 |
| Primary somatosensory area, mouth | 0.3750864 | 0.40709 | 0.9213845 | 0.3568497 |
| Primary somatosensory area, upper limb | 0.4643062 | 0.2352882 | 1.973351 | 0.04845557 |
| Primary somatosensory area, trunk | 0.4005701 | 0.15181094 | 2.638612 | 0.008324626 |
| Supplemental somatosensory area | 0.3613312 | 0.2046063 | 1.765983 | 0.07739868 |
| Gustatory areas | -0.3626419 | 0.2373255 | -1.528036 | 0.1265036 |
| Visceral area | -0.006613192 | 0.264708 | -0.02498297 | 0.9800685 |
| Dorsal auditory area | 0.3815158 | 0.2464848 | 1.547827 | 0.121664 |
| Primary auditory area | 0.3513487 | 0.3513649 | 0.9999539 | 0.3173328 |
| Posterior auditory area | -0.1428659 | 0.530661 | -0.2692226 | 0.7877584 |
| Ventral auditory area | -0.1495439 | 0.27824 | -0.5374638 | 0.5909473 |
| Anterolateral visual area | -0.1985893 | 0.4540335 | -0.437389 | 0.6618293 |
| Anteromedial visual area | -0.07642186 | 0.3440248 | -0.2221405 | 0.8242045 |
| Lateral visual area | -0.4269711 | 0.4883651 | -0.8742867 | 0.3819621 |
| Primary visual area | -0.460163 | 0.4446626 | -1.034859 | 0.3007348 |
| Posterolateral visual area | 0.006602699 | 0.459901 | 0.01435678 | 0.9885453 |
| Posteromedial visual area | -0.3021566 | 0.3894748 | -0.7758051 | 0.4378641 |
| Anterior cingulate area, dorsal part | 0.1575336 | 0.10021997 | 1.571878 | 0.1159789 |
| Anterior cingulate area, ventral part | 0.1447553 | 0.15905091 | 0.9101195 | 0.3627595 |
| Prelimbic area | 0.08524969 | 0.16659292 | 0.5117246 | 0.6088438 |
| Infralimbic area | 0.250842 | 0.15582037 | 1.609815 | 0.1074382 |
| Orbital area, lateral part | 0.4429749 | 0.2052411 | 2.158315 | 0.03090338 |
| Orbital area, medial part | -0.1987482 | 0.251356 | -0.7907042 | 0.4291166 |
| Orbital area, ventrolateral part | 0.02260728 | 0.18084147 | 0.1250116 | 0.9005144 |
| Agranular insular area, dorsal part | 0.1759461 | 0.2965886 | 0.5932328 | 0.5530253 |
| Agranular insular area, posterior part | 0.2263496 | 0.2828577 | 0.8002241 | 0.423581 |
| Agranular insular area, ventral part | 0.08590713 | 0.2863841 | 0.2999717 | 0.7641987 |
| Retrosplenial area, lateral agranular part | -0.09550695 | 0.264971 | -0.360443 | 0.7185159 |
| Retrosplenial area, dorsal part | -0.2712026 | 0.2592133 | -1.046252 | 0.2954445 |
| Retrosplenial area, ventral part | 0.1766387 | 0.11572602 | 1.526352 | 0.1269222 |
| Rostrolateral visual area | -0.1519449 | 0.3582295 | -0.4241551 | 0.6714527 |
| Temporal association areas | -0.289173 | 0.2632088 | -1.098645 | 0.271923 |
| Perirhinal area | -0.1222689 | 0.44401 | -0.2753742 | 0.7830288 |
| Ectorhinal area | -0.08238387 | 0.3365775 | -0.2447694 | 0.806635 |
| Anterior olfactory nucleus | 0.4473582 | 0.3338847 | 1.339858 | 0.1802914 |
| Taenia tecta, dorsal part | 0.2062441 | 0.12559753 | 1.642103 | 0.1005686 |
| Taenia tecta, ventral part | 0.09854577 | 0.4687827 | 0.2102163 | 0.8334989 |
| Dorsal peduncular area | 0.4325683 | 0.1876764 | 2.304862 | 0.02117429 |
| Piriform area | 0.1804987 | 0.4150149 | 0.434921 | 0.6636197 |

| | | | | |
|---|---|---|---|---|
| Nucleus of the lateral olfactory tract | -0.3965505 | 0.9236607 | -0.4293249 | 0.6676868 |
| Cortical amygdalar area, anterior part | -0.3992196 | 0.9634507 | -0.4143643 | 0.6786073 |
| Cortical amygdalar area, posterior part | -0.1557786 | 0.7549912 | -0.2063317 | 0.8365318 |
| Piriform,amygdalar area | 0.01590863 | 0.7184263 | 0.02214372 | 0.9823333 |
| Postpiriform transition area | -0.1707377 | 0.6646866 | -0.2568695 | 0.7972795 |
| Field c a1 | 0.2179599 | 0.13225912 | 1.647976 | 0.09935758 |
| Field c a2 | 0.4073569 | 0.2230294 | 1.826472 | 0.06777922 |
| Field c a3 | 0.2870473 | 0.1690533 | 1.697969 | 0.08951367 |
| Dentate gyrus | -0.07291174 | 0.1971033 | -0.3699164 | 0.7114448 |
| Induseum griseum | -0.5192171 | 0.3322468 | -1.562745 | 0.1181126 |
| Entorhinal area, lateral part | -0.031642 | 0.5121754 | -0.06177962 | 0.9507383 |
| Entorhinal area, medial part | -0.0753039 | 0.244374 | -0.3081502 | 0.757968 |
| Parasubiculum | -0.6632297 | 0.20869911 | -3.177923 | 0.001483343 |
| Postsubiculum | 0.05011429 | 0.1907654 | 0.2627012 | 0.7927809 |
| Presubiculum | -0.2557405 | 0.1790161 | -1.428589 | 0.1531224 |
| Subiculum | -0.1993997 | 0.1837327 | -1.085271 | 0.2778018 |
| Prosubiculum | 0.02988401 | 0.11485533 | 0.2601883 | 0.7947186 |
| Claustrum | 0.03077765 | 0.17169102 | 0.1792618 | 0.8577321 |
| Endopiriform nucleus, dorsal part | 0.1199075 | 0.3330066 | 0.3600755 | 0.7187907 |
| Endopiriform nucleus, ventral part | 0.01722877 | 0.5717711 | 0.03013228 | 0.9759616 |
| Lateral amygdalar nucleus | -0.5229927 | 0.2694656 | -1.940852 | 0.05227629 |
| Basolateral amygdalar nucleus, anterior part | -0.2928538 | 0.4123356 | -0.7102316 | 0.4775605 |
| Basolateral amygdalar nucleus, posterior part | -0.2978764 | 0.5144031 | -0.5790718 | 0.5625407 |
| Basolateral amygdalar nucleus, ventral part | -0.009075707 | 0.8163896 | -0.01111688 | 0.9911302 |
| Basomedial amygdalar nucleus, anterior part | -0.2564467 | 0.8042535 | -0.3188631 | 0.7498303 |
| Basomedial amygdalar nucleus, posterior part | -0.3008758 | 0.5526251 | -0.5444484 | 0.586133 |
| Posterior amygdalar nucleus | -0.4919111 | 0.5235572 | -0.9395557 | 0.3474455 |
| Caudoputamen | -0.07061924 | 0.2356578 | -0.2996686 | 0.7644299 |
| Nucleus accumbens | 0.2580504 | 0.2898658 | 0.890241 | 0.3733365 |
| Fundus of striatum | 0.03524639 | 0.2315589 | 0.1522135 | 0.8790185 |
| Olfactory tubercle | -0.01708012 | 0.5542317 | -0.03081765 | 0.975415 |
| Lateral septal nucleus | 0.02855094 | 0.23348 | 0.1222843 | 0.9026739 |
| Septofimbrial nucleus | -0.4630892 | 0.4527207 | -1.022903 | 0.3063538 |
| Anterior amygdalar area | -0.6770354 | 0.5110169 | -1.324879 | 0.1852114 |
| Central amygdalar nucleus, capsular part | -0.7960965 | 0.2629731 | -3.027292 | 0.002467552 |
| Central amygdalar nucleus, lateral part | 0.1262841 | 0.2528909 | 0.499362 | 0.6175244 |
| Central amygdalar nucleus, medial part | -0.3431147 | 0.2743697 | -1.250556 | 0.2110965 |
| Intercalated amygdalar nucleus | -0.1944765 | 0.4621923 | -0.4207696 | 0.6739233 |
| Medial amygdalar nucleus | -0.6202674 | 0.5183793 | -1.196551 | 0.2314815 |
| Globus pallidus, external segment | 0.1582658 | 0.3810345 | 0.4153581 | 0.6778798 |
| Globus pallidus, internal segment | 0.7701132 | 0.6544696 | 1.176698 | 0.2393159 |
| Substantia innominata | -0.1891231 | 0.20483407 | -0.9232991 | 0.3558514 |

| | | | | |
|---|---|---|---|---|
| Magnocellular nucleus | -0.5767355 | 0.4408082 | -1.308359 | 0.1907514 |
| Medial septal nucleus | 0.1561568 | 0.2356335 | 0.6627103 | 0.5075161 |
| Diagonal band nucleus | -0.3212498 | 0.3186444 | -1.008177 | 0.3133697 |
| Triangular nucleus of septum | -1.011283 | 0.4292159 | -2.356117 | 0.01846709 |
| Bed nuclei of the stria terminalis | 0.1804478 | 0.229172 | 0.7873902 | 0.4310535 |
| Ventral anterior,lateral complex of the thalamus | -1.796687 | 0.4667599 | -3.849273 | 0.0001184687 |
| Ventral medial nucleus of the thalamus | 0.238154 | 0.3275081 | 0.7271697 | 0.467122 |
| Ventral posterior complex of the thalamus | -0.3397661 | 0.2549696 | -1.332575 | 0.1826712 |
| Subparafascicular nucleus | -0.4547136 | 0.3389766 | -1.341431 | 0.1797806 |
| Subparafascicular area | 0.2089501 | 0.7739845 | 0.2699668 | 0.7871858 |
| Medial geniculate complex | -0.9249893 | 0.339288 | -2.726266 | 0.006405531 |
| Lateral geniculate complex | -0.4974655 | 0.11364579 | -4.377334 | 1.2014E-05 |
| Lateral posterior nucleus of the thalamus | -0.8229428 | 0.2260546 | -3.64046 | 0.0002721517 |
| Posterior complex of the thalamus | -0.1578372 | 0.3177987 | -0.4966577 | 0.6194305 |
| Posterior limiting nucleus of the thalamus | -0.6067726 | 0.2380935 | -2.548463 | 0.01081986 |
| Suprageniculate nucleus | -0.8277045 | 0.2878887 | -2.875085 | 0.00403919 |
| Anteroventral nucleus of thalamus | -1.149953 | 0.4437757 | -2.591293 | 0.009561608 |
| Anteromedial nucleus | -0.696325 | 0.4732021 | -1.471517 | 0.1411513 |
| Anterodorsal nucleus | -0.9582275 | 0.3605164 | -2.65793 | 0.007862211 |
| Interanterodorsal nucleus of the thalamus | -0.4529004 | 0.3480007 | -1.301435 | 0.1931094 |
| Lateral dorsal nucleus of thalamus | -0.6115842 | 0.3670608 | -1.666166 | 0.09568037 |
| Intermediodorsal nucleus of the thalamus | -0.3871287 | 0.4093915 | -0.9456198 | 0.3443425 |
| Mediodorsal nucleus of thalamus | -0.3709288 | 0.2436461 | -1.522408 | 0.1279068 |
| Submedial nucleus of the thalamus | -0.7623623 | 0.8390614 | -0.9085894 | 0.3635669 |
| Perireunensis nucleus | 0.2283013 | 0.2898008 | 0.7877869 | 0.4308214 |
| Paraventricular nucleus of the thalamus | -0.6828314 | 0.2283991 | -2.989641 | 0.002793051 |
| Parataenial nucleus | -0.4803399 | 0.234972 | -2.044243 | 0.04092951 |
| Nucleus of reuniens | 0.3040992 | 0.1929893 | 1.57573 | 0.1150879 |
| Central medial nucleus of the thalamus | -0.1175455 | 0.3623718 | -0.3243783 | 0.7456517 |
| Paracentral nucleus | 0.3091355 | 0.4499995 | 0.6869685 | 0.4921026 |
| Central lateral nucleus of the thalamus | -0.5235853 | 0.3067146 | -1.707076 | 0.08780789 |
| Parafascicular nucleus | -0.1487215 | 0.3543676 | -0.4196813 | 0.6747183 |
| Reticular nucleus of the thalamus | -0.6171223 | 0.3378166 | -1.826797 | 0.06773035 |
| Medial habenula | -1.786815 | 0.3718796 | -4.804822 | 1.548892E-06 |
| Lateral habenula | -0.3033435 | 0.3717012 | -0.8160949 | 0.4144458 |
| Paraventricular hypothalamic nucleus | 0.0150156 | 0.2347123 | 0.06397449 | 0.9489905 |
| Periventricular hypothalamic nucleus | -0.3962473 | 0.4072614 | -0.9729556 | 0.3305753 |
| Arcuate hypothalamic nucleus | -1.239951 | 0.8605458 | -1.440889 | 0.1496159 |
| Anterodorsal preoptic nucleus | -0.1717768 | 0.263822 | -0.6511088 | 0.5149762 |
| Anteroventral periventricular nucleus | -0.07990803 | 0.4388064 | -0.1821031 | 0.8555018 |
| Dorsomedial hypothalamic nucleus | -0.2380205 | 0.2763872 | -0.8611849 | 0.3891362 |
| Medial preoptic area | 0.06357373 | 0.20158629 | 0.3153673 | 0.7524828 |

| | | | | |
|---|---|---|---|---|
| Subparaventricular zone | -0.02093185 | 0.3729992 | -0.05611768 | 0.9552481 |
| Anterior hypothalamic nucleus | 0.06469772 | 0.2455263 | 0.2635063 | 0.7921604 |
| Supramammillary nucleus | -1.015551 | 0.7842204 | -1.294982 | 0.1953265 |
| Tuberomammillary nucleus | -1.123202 | 0.9351017 | -1.201155 | 0.2296913 |
| Medial preoptic nucleus | -0.1736042 | 0.2471188 | -0.7025132 | 0.4823592 |
| Dorsal premammillary nucleus | -1.266924 | 0.707424 | -1.790898 | 0.07330969 |
| Ventral premammillary nucleus | -1.048835 | 1.1345989 | -0.9244108 | 0.3552725 |
| Ventromedial hypothalamic nucleus | -0.2573631 | 0.6793395 | -0.3788432 | 0.7048043 |
| Posterior hypothalamic nucleus | -0.348847 | 0.2577819 | -1.353264 | 0.1759712 |
| Lateral hypothalamic area | -0.05146403 | 0.2575134 | -0.1998499 | 0.841598 |
| Lateral preoptic area | 0.2402169 | 0.21107186 | 1.138081 | 0.2550866 |
| Parasubthalamic nucleus | -0.06657257 | 0.3384523 | -0.196697 | 0.8440646 |
| Retrochiasmatic area | -0.05938953 | 0.8013911 | -0.07410805 | 0.9409244 |
| Subthalamic nucleus | 0.5362049 | 0.3834688 | 1.398301 | 0.1620226 |
| Tuberal nucleus | -0.2361503 | 0.8062487 | -0.2929001 | 0.7695985 |
| Zona incerta | 0.3772034 | 0.2517821 | 1.498134 | 0.1340984 |
| Superior colliculus, dorsal part | -0.2790528 | 0.2393125 | -1.16606 | 0.2435901 |
| Inferior colliculus, central part | 0.4100654 | 0.2410228 | 1.701355 | 0.08887631 |
| Inferior colliculus, dorsal part | 0.2546776 | 0.2825131 | 0.9014716 | 0.3673376 |
| Inferior colliculus, external part | 0.2187117 | 0.2665116 | 0.8206461 | 0.4118479 |
| Substantia nigra, reticular part | -0.5498089 | 0.3369903 | -1.631527 | 0.1027791 |
| Ventral tegmental area | 0.588877 | 0.3101824 | 1.898486 | 0.05763207 |
| Midbrain reticular nucleus | 0.1365546 | 0.12904016 | 1.058233 | 0.2899492 |
| Superior colliculus, ventral part | -0.0714606 | 0.1586441 | -0.4504461 | 0.6523888 |
| Periaqueductal gray | -0.3337478 | 0.1728062 | -1.931342 | 0.05344079 |
| Anterior pretectal nucleus | -0.3540254 | 0.3248668 | -1.089756 | 0.2758207 |
| Nucleus of the optic tract | -0.7440978 | 0.2776777 | -2.679718 | 0.007368424 |
| Nucleus of the posterior commissure | -0.6244987 | 0.4094112 | -1.525358 | 0.1271698 |
| Posterior pretectal nucleus | -0.6344413 | 0.2166377 | -2.928582 | 0.003405122 |
| Cuneiform nucleus | -0.3730072 | 0.2066465 | -1.80505 | 0.07106686 |
| Red nucleus | 0.1575498 | 0.3648276 | 0.4318474 | 0.6658524 |
| Substantia nigra, compact part | -0.00205516 | 0.3507504 | -0.005859323 | 0.995325 |
| Pedunculopontine nucleus | -0.2414875 | 0.255859 | -0.9438301 | 0.3452565 |
| Dorsal nucleus raphe | 0.06181184 | 0.3524876 | 0.1753589 | 0.8607976 |
| Nucleus of the lateral lemniscus | -0.005216064 | 0.7053569 | -0.007394929 | 0.9940998 |
| Principal sensory nucleus of the trigeminal | 0.1829322 | 0.3532823 | 0.5178075 | 0.6045926 |
| Parabrachial nucleus | -0.9509668 | 0.2847278 | -3.339915 | 0.0008380393 |
| Superior olivary complex | -0.00714207 | 0.5646187 | -0.01264937 | 0.9899075 |
| Dorsal tegmental nucleus | -0.7757344 | 0.4731047 | -1.639667 | 0.1010743 |
| Pontine central gray | -1.074062 | 0.2914699 | -3.684985 | 0.0002287165 |
| Pontine gray | -0.03197004 | 0.5961329 | -0.05362904 | 0.9572307 |
| Supratrigeminal nucleus | -0.5702325 | 0.3260015 | -1.749172 | 0.08026137 |

| | | | | |
|---|---|---|---|---|
| Tegmental reticular nucleus | 1.09061 | 0.6907327 | 1.578918 | 0.1143549 |
| Motor nucleus of trigeminal | 0.1284899 | 0.3722911 | 0.3451328 | 0.7299946 |
| Superior central nucleus raphe | 1.211382 | 0.5231266 | 2.315658 | 0.02057696 |
| Laterodorsal tegmental nucleus | -0.2904853 | 0.3957969 | -0.7339251 | 0.4629944 |
| Nucleus incertus | -0.613166 | 0.23329689 | -2.628265 | 0.008582164 |
| Pontine reticular nucleus | -0.05391102 | 0.4316807 | -0.1248863 | 0.9006135 |
| Dorsal cochlear nucleus | 0.4205356 | 0.2871624 | 1.464452 | 0.1430705 |
| Ventral cochlear nucleus | 0.4787787 | 0.3102012 | 1.543446 | 0.1227227 |
| Cuneate nucleus | 0.5882448 | 0.616343 | 0.9544115 | 0.3398754 |
| External cuneate nucleus | 0.1861427 | 0.5253176 | 0.3543432 | 0.7230817 |
| Nucleus of the trapezoid body | -0.7060439 | 0.7899295 | -0.8938062 | 0.3714256 |
| Nucleus of the solitary tract | -1.453742 | 0.5345729 | -2.719447 | 0.006539122 |
| Spinal nucleus of the trigeminal | -0.2727857 | 0.3624425 | -0.7526316 | 0.4516713 |
| Facial motor nucleus | -0.662725 | 0.4843427 | -1.368298 | 0.1712189 |
| Dorsal motor nucleus of the vagus nerve | -1.764594 | 0.5411714 | -3.260694 | 0.0011114 |
| Gigantocellular reticular nucleus | 0.1495062 | 0.4591995 | 0.32558 | 0.7447421 |
| Inferior olivary complex | -1.134499 | 1.0196761 | -1.112607 | 0.265877 |
| Intermediate reticular nucleus | -0.4056733 | 0.4605896 | -0.8807695 | 0.3784426 |
| Lateral reticular nucleus | -1.087946 | 0.5462209 | -1.99177 | 0.04639633 |
| Magnocellular reticular nucleus | -0.7524659 | 0.7125782 | -1.055977 | 0.2909789 |
| Medullary reticular nucleus | -0.9030645 | 0.4813991 | -1.875916 | 0.06066675 |
| Parvicellular reticular nucleus | -0.2965332 | 0.3895286 | -0.7612617 | 0.4465008 |
| Paragigantocellular reticular nucleus | -0.1365971 | 0.3793242 | -0.3601065 | 0.7187675 |
| Nucleus prepositus | 0.07890425 | 0.2964712 | 0.2661447 | 0.7901278 |
| Lateral vestibular nucleus | 0.3009039 | 0.3581353 | 0.840196 | 0.4007985 |
| Medial vestibular nucleus | -0.4214147 | 0.2589051 | -1.62768 | 0.1035928 |
| Spinal vestibular nucleus | 0.0647102 | 0.2871863 | 0.2253248 | 0.8217266 |
| Superior vestibular nucleus | 0.3536197 | 0.4136274 | 0.8549233 | 0.3925936 |
| Hypoglossal nucleus | -0.9129928 | 0.5897548 | -1.548089 | 0.1216009 |
| Nucleus raphe magnus | -0.2438884 | 0.6388284 | -0.3817745 | 0.7026286 |

**Supplementary Data Table 63:** GLMM coefficient estimates of ***SI Time*** contribution to Fos$^+$ cell counts in unstimulated control mice (**Figure S8A**). The total number of Fos$^+$ cells in each brain region was modeled as ***Counts ~ SI Time + Stim + SI Time:Stim* + ln(*Total Counts*) + (*1+SI Time*|*Cohort*)** using a negative binomial link function (*N* = 44 mice). The critical *p*-value for significance while permitting a 10% FDR was 0.000025; 1 region was significant.

| Brain region name | SI time coefficient estimate | SI time coefficient std error | SI time coefficient Z-statistic | SI time coefficient P-value |
|---|---|---|---|---|
| Frontal pole | -0.1264267 | 0.1954137 | -0.6469698 | 0.5176515 |
| Primary motor area | 0.030783744 | 0.0817474 | 0.37657152 | 0.7064921 |

| | | | | |
|---|---|---|---|---|
| Secondary motor area | 0.00621545 | 0.047332 | 0.131316 | 0.89552531 |
| Primary somatosensory area, nose | -0.1006953 | 0.1133659 | -0.8882333 | 0.3744152 |
| Primary somatosensory area, barrel field | 0.006530494 | 0.0873 | 0.0748052 | 0.9403697 |
| Primary somatosensory area, lower limb | 0.0938509 | 0.1157641 | 0.81070828 | 0.4175332 |
| Primary somatosensory area, mouth | -0.1139833 | 0.1174932 | -0.9701263 | 0.3319835 |
| Primary somatosensory area, upper limb | 0.01455021 | 0.09612965 | 0.1513603 | 0.8796915 |
| Primary somatosensory area, trunk | 0.0305854 | 0.06758642 | 0.4525376 | 0.65088171 |
| Supplemental somatosensory area | -0.03098379 | 0.0778573 | -0.3979561 | 0.6906625 |
| Gustatory areas | 0.02236063 | 0.115418 | 0.1937361 | 0.8463825 |
| Visceral area | -0.08925373 | 0.08578594 | -1.0404237 | 0.2981431 |
| Dorsal auditory area | 0.01020095 | 0.1069429 | 0.09538692 | 0.9240075 |
| Primary auditory area | 0.01364062 | 0.1233951 | 0.1105443 | 0.9119777 |
| Posterior auditory area | -0.14349007 | 0.2746675 | -0.5224138 | 0.6013823 |
| Ventral auditory area | -0.006157011 | 0.1092788 | -0.05634224 | 0.9550692 |
| Anterolateral visual area | -0.05292371 | 0.2159978 | -0.2450197 | 0.8064412 |
| Anteromedial visual area | -0.03936384 | 0.1212109 | -0.3247548 | 0.7453666 |
| Lateral visual area | -0.0378537 | 0.2368907 | -0.159794 | 0.8730434 |
| Primary visual area | -0.02202972 | 0.1763243 | -0.1249387 | 0.9005721 |
| Posterolateral visual area | -0.1984164749 | 0.1488327 | -1.3331513872 | 0.1824821 |
| Posteromedial visual area | -0.0771167 | 0.1354888 | -0.5691739 | 0.5692381 |
| Anterior cingulate area, dorsal part | -0.002062887 | 0.04042858 | -0.05102546 | 0.959305235 |
| Anterior cingulate area, ventral part | 0.03722007 | 0.04547977 | 0.8183873 | 0.4131360466 |
| Prelimbic area | 0.009532914 | 0.08016829 | 0.11891128 | 0.9053456 |
| Infralimbic area | -0.04287446 | 0.0761928 | -0.5627102 | 0.57363225 |
| Orbital area, lateral part | -0.01270432 | 0.06861798 | -0.1851456 | 0.8531148 |
| Orbital area, medial part | 0.0106742 | 0.1030591 | 0.10357354 | 0.9175078 |
| Orbital area, ventrolateral part | -0.06719145 | 0.09090004 | -0.7391796 | 0.45979797 |
| Agranular insular area, dorsal part | 0.06751079 | 0.08711006 | 0.7750056 | 0.4383363 |
| Agranular insular area, posterior part | -0.05817762 | 0.08869604 | -0.6559213 | 0.5118748 |
| Agranular insular area, ventral part | 0.03149859 | 0.1065779 | 0.2955453 | 0.7675774 |
| Retrosplenial area, lateral agranular part | -0.04269047 | 0.112633 | -0.3790227 | 0.704671 |
| Retrosplenial area, dorsal part | 0.02305168 | 0.1092254 | 0.2110469 | 0.8328507 |
| Retrosplenial area, ventral part | 0.04755993 | 0.05662699 | 0.8398809 | 0.4009751 |
| Rostrolateral visual area | -0.05949862 | 0.1033374 | -0.5757705 | 0.5647703 |
| Temporal association areas | -0.02196668 | 0.1382091 | -0.158938 | 0.8737177 |
| Perirhinal area | -0.19923484 | 0.1444613 | -1.3791575 | 0.1678462 |
| Ectorhinal area | -0.011661002 | 0.1578813 | -0.07385932 | 0.9411223 |
| Anterior olfactory nucleus | -0.02854369 | 0.155897 | -0.1830932 | 0.8547249 |
| Taenia tecta, dorsal part | -0.01236022 | 0.06292675 | -0.1964223 | 0.8442796 |
| Taenia tecta, ventral part | -0.05150716 | 0.205766 | -0.2503191 | 0.8023406 |
| Dorsal peduncular area | -0.09181247 | 0.08841588 | -1.038416 | 0.2990764 |
| Piriform area | 0.008610064 | 0.1632057 | 0.05275589 | 0.9579264 |

| | | | | |
|---|---|---|---|---|
| Nucleus of the lateral olfactory tract | -0.6161204 | 0.5143028 | -1.1979722 | 0.2309278 |
| Cortical amygdalar area, anterior part | -1.1050967 | 0.8745885 | -1.263562 | 0.2063873 |
| Cortical amygdalar area, posterior part | -0.2809676 | 0.3931279 | -0.7146977 | 0.4747958 |
| Piriform,amygdalar area | -0.39986133 | 0.4029429 | -0.99235233 | 0.3210257 |
| Postpiriform transition area | -0.37000981 | 0.213594 | -1.7323047 | 0.08321933 |
| Field c a1 | 0.05647456 | 0.05415365 | 1.0428579 | 0.2970141 |
| Field c a2 | -0.03236074 | 0.07331365 | -0.4414013 | 0.6589225 |
| Field c a3 | 0.02379144 | 0.05531103 | 0.4301391 | 0.6670945 |
| Dentate gyrus | 0.03617501 | 0.06739426 | 0.536767 | 0.5914286 |
| Induseum griseum | -0.06573308 | 0.1364009 | -0.481911 | 0.6298692 |
| Entorhinal area, lateral part | -0.11565822 | 0.1633876 | -0.70787639 | 0.479022 |
| Entorhinal area, medial part | -0.05305423 | 0.09924029 | -0.5346037 | 0.5929239 |
| Parasubiculum | -0.002891138 | 0.09851037 | -0.02934857 | 0.97658659 |
| Postsubiculum | 0.06897404 | 0.06346763 | 1.08675925 | 0.2771432 |
| Presubiculum | 0.08303751 | 0.05780987 | 1.43639 | 0.1508915 |
| Subiculum | -0.05059324 | 0.07360104 | -0.68739843 | 0.4918317 |
| Prosubiculum | -0.01050149 | 0.05040592 | -0.2083383 | 0.8349648 |
| Claustrum | 0.04316053 | 0.06748869 | 0.6395224 | 0.5224831 |
| Endopiriform nucleus, dorsal part | 0.08879711 | 0.1295925 | 0.6852024 | 0.4932162 |
| Endopiriform nucleus, ventral part | -0.47453375 | 0.1719336 | -2.75998247 | 0.005780446 |
| Lateral amygdalar nucleus | 0.005610335 | 0.09932867 | 0.05648253 | 0.9549574 |
| Basolateral amygdalar nucleus, anterior part | -0.08517842 | 0.1567809 | -0.5432958 | 0.5869261 |
| Basolateral amygdalar nucleus, posterior part | -0.1862663 | 0.1919807 | -0.9702347 | 0.3319295 |
| Basolateral amygdalar nucleus, ventral part | -0.8369194 | 0.7838769 | -1.067667 | 0.2856708 |
| Basomedial amygdalar nucleus, anterior part | -0.5130189 | 0.3849316 | -1.3327532 | 0.1826128 |
| Basomedial amygdalar nucleus, posterior part | -0.3091427 | 0.1833453 | -1.6861232 | 0.09177207 |
| Posterior amygdalar nucleus | -0.1194613 | 0.1661523 | -0.7189868 | 0.472149 |
| Caudoputamen | 0.061779601 | 0.08897638 | 0.69433709 | 0.4874708 |
| Nucleus accumbens | 0.04244781 | 0.09314477 | 0.4557186 | 0.6485923 |
| Fundus of striatum | -0.023439523 | 0.10077879 | -0.2325839 | 0.8160845 |
| Olfactory tubercle | -0.1372783 | 0.2610811 | -0.5258071 | 0.5990222 |
| Lateral septal nucleus | -0.01475418 | 0.06801403 | -0.2169285 | 0.828264 |
| Septofimbrial nucleus | -0.07341144 | 0.1006598 | -0.7293022 | 0.4658168 |
| Anterior amygdalar area | -0.31779907 | 0.1989126 | -1.59768234 | 0.1101137 |
| Central amygdalar nucleus, capsular part | -0.1314241 | 0.1268297 | -1.036225 | 0.30009719 |
| Central amygdalar nucleus, lateral part | -0.1759051 | 0.1070823 | -1.6427091 | 0.1004431 |
| Central amygdalar nucleus, medial part | -0.1332277 | 0.1364389 | -0.9764644 | 0.3288344 |
| Intercalated amygdalar nucleus | -0.3241171 | 0.1459698 | -2.2204401 | 0.0263889 |
| Medial amygdalar nucleus | -0.1275957 | 0.1737279 | -0.7344572 | 0.4626701 |
| Globus pallidus, external segment | -0.01048457 | 0.0995155 | -0.1053561 | 0.9160932 |
| Globus pallidus, internal segment | -0.06736136 | 0.139619 | -0.4824655 | 0.6294753 |
| Substantia innominata | 0.006605879 | 0.09274836 | 0.07122367 | 0.9432197 |

| | | | | |
|---|---|---|---|---|
| Magnocellular nucleus | -0.157502247 | 0.1273346 | -1.2369166 | 0.216118 |
| Medial septal nucleus | -0.13432896 | 0.07131442 | -1.8836156 | 0.05961698 |
| Diagonal band nucleus | -0.14523949 | 0.09574078 | -1.5170076 | 0.1292648 |
| Triangular nucleus of septum | 0.0154493 | 0.1088643 | 0.1419134 | 0.8871484 |
| Bed nuclei of the stria terminalis | -0.07511383 | 0.08995078 | -0.8350548 | 0.4036869 |
| Ventral anterior,lateral complex of the thalamus | -0.02738486 | 0.1374078 | -0.1992963 | 0.842031 |
| Ventral medial nucleus of the thalamus | -0.08655617 | 0.1557822 | -0.5556229 | 0.5784687 |
| Ventral posterior complex of the thalamus | 0.04247172 | 0.07751441 | 0.5479203 | 0.5837466 |
| Subparafascicular nucleus | -0.05324835 | 0.09385869 | -0.5673247 | 0.5704936 |
| Subparafascicular area | -0.09568648 | 0.1668172 | -0.5736009 | 0.5662379 |
| Medial geniculate complex | 0.04703251 | 0.09792973 | 0.480268 | 0.6310369 |
| Lateral geniculate complex | 0.01872897 | 0.04363248 | 0.4292439 | 0.6677458 |
| Lateral posterior nucleus of the thalamus | 0.02528143 | 0.08531631 | 0.2963258 | 0.7669813 |
| Posterior complex of the thalamus | 0.2479171 | 0.1364589 | 1.816789 | 0.06924937 |
| Posterior limiting nucleus of the thalamus | -0.1019839 | 0.1202638 | -0.8480017 | 0.3964371 |
| Suprageniculate nucleus | -0.0279923 | 0.06786634 | -0.4124622 | 0.6800007 |
| Anteroventral nucleus of thalamus | -0.003285927 | 0.1381521 | -0.02378486 | 0.9810242 |
| Anteromedial nucleus | -0.1539657 | 0.1267462 | -1.214756 | 0.2244591 |
| Anterodorsal nucleus | -0.1705883 | 0.1107625 | -1.540127 | 0.1235294 |
| Interanterodorsal nucleus of the thalamus | -0.04856269 | 0.1137043 | -0.4270965 | 0.669309 |
| Lateral dorsal nucleus of thalamus | 0.09442304 | 0.1139739 | 0.8284622 | 0.4074088 |
| Intermediodorsal nucleus of the thalamus | -0.2122318 | 0.1272525 | -1.6678001 | 0.09535543 |
| Mediodorsal nucleus of thalamus | -0.07842985 | 0.07983645 | -0.9823815 | 0.3259119 |
| Submedial nucleus of the thalamus | -0.003370518 | 0.1427868 | -0.02360524 | 0.9811675 |
| Perireunensis nucleus | -0.08282489 | 0.1114208 | -0.7433525 | 0.4572683 |
| Paraventricular nucleus of the thalamus | -0.03150503 | 0.07868054 | -0.4004171 | 0.6888493 |
| Parataenial nucleus | -0.07166078 | 0.08113141 | -0.8832681 | 0.3770914 |
| Nucleus of reuniens | -0.0710729 | 0.09133681 | -0.7781408 | 0.43648599 |
| Central medial nucleus of the thalamus | -0.06586492 | 0.1270575 | -0.5183869 | 0.6041883 |
| Paracentral nucleus | -0.02102195 | 0.1181864 | -0.1778711 | 0.8588242 |
| Central lateral nucleus of the thalamus | 0.08047318 | 0.09904601 | 0.8124828 | 0.4165147 |
| Parafascicular nucleus | 0.03782199 | 0.118414 | 0.3194048 | 0.7494196 |
| Reticular nucleus of the thalamus | -0.1132467 | 0.09493788 | -1.1928506 | 0.2329279 |
| Medial habenula | 0.5142014 | 0.1218878 | 4.218644 | 2.457759E-05 |
| Lateral habenula | 0.03558852 | 0.09855755 | 0.3610938 | 0.7180294 |
| Paraventricular hypothalamic nucleus | -0.01973027 | 0.11521264 | -0.1712509 | 0.8640265 |
| Periventricular hypothalamic nucleus | 0.08217766 | 0.1466325 | 0.5604328 | 0.5751843 |
| Arcuate hypothalamic nucleus | -0.4742477 | 0.6367102 | -0.7448407 | 0.456368 |
| Anterodorsal preoptic nucleus | -0.09261268 | 0.09865965 | -0.9387088 | 0.3478803 |
| Anteroventral periventricular nucleus | -0.1517142 | 0.1332633 | -1.1384543 | 0.2549308 |
| Dorsomedial hypothalamic nucleus | -0.00609135 | 0.1297512 | -0.0469464 | 0.962556 |
| Medial preoptic area | -0.11297299 | 0.0903439 | -1.2504773 | 0.2111252 |

| | | | | |
|---|---|---|---|---|
| Subparaventricular zone | -0.1085096 | 0.1389729 | -0.7807968 | 0.434922 |
| Anterior hypothalamic nucleus | -0.068674547 | 0.1170565 | -0.58667874 | 0.5574195 |
| Supramammillary nucleus | -0.0711657 | 0.1877829 | -0.3789786 | 0.7047038 |
| Tuberomammillary nucleus | -0.4093644 | 0.2466494 | -1.659701 | 0.09697455 |
| Medial preoptic nucleus | -0.07411133 | 0.1086408 | -0.682168748 | 0.4951323 |
| Dorsal premammillary nucleus | -0.07145146 | 0.1976116 | -0.3615752 | 0.7176695 |
| Ventral premammillary nucleus | 0.06011673 | 0.4069731 | 0.1477167 | 0.8825663 |
| Ventromedial hypothalamic nucleus | -0.1163118 | 0.2092843 | -0.5557597 | 0.5783751 |
| Posterior hypothalamic nucleus | -0.04763004 | 0.1247626 | -0.3817653 | 0.7026355 |
| Lateral hypothalamic area | -0.02120213 | 0.1236522 | -0.1714659 | 0.8638575 |
| Lateral preoptic area | -0.1414789 | 0.09138293 | -1.5481982 | 0.1215746 |
| Parasubthalamic nucleus | -0.02259563 | 0.1571881 | -0.143749 | 0.8856987 |
| Retrochiasmatic area | -0.41678496 | 0.6300028 | -0.6615605 | 0.5082529 |
| Subthalamic nucleus | -0.08064712 | 0.1348986 | -0.5978351 | 0.5499499 |
| Tuberal nucleus | -0.1895058 | 0.2332058 | -0.8126119 | 0.4164406 |
| Zona incerta | -0.08347846 | 0.119953 | -0.6959266 | 0.4864748 |
| Superior colliculus, dorsal part | 0.06760481 | 0.08446078 | 0.8004285 | 0.4234626 |
| Inferior colliculus, central part | 0.07324502 | 0.09368692 | 0.7818062 | 0.4343285 |
| Inferior colliculus, dorsal part | 0.00866651 | 0.1314439 | 0.06593315 | 0.947431 |
| Inferior colliculus, external part | 0.05909524 | 0.1072623 | 0.5509414 | 0.5816739 |
| Substantia nigra, reticular part | -0.04062963 | 0.1157111 | -0.3511298 | 0.72549096 |
| Ventral tegmental area | -0.10061886 | 0.1426521 | -0.7053443 | 0.480596 |
| Midbrain reticular nucleus | -0.007795432 | 0.07251467 | -0.1075015 | 0.9143912 |
| Superior colliculus, ventral part | -0.0008393465 | 0.06474092 | -0.0129647 | 0.989656 |
| Periaqueductal gray | 0.005999178 | 0.07738623 | 0.07752256 | 0.9382078 |
| Anterior pretectal nucleus | 0.03395264 | 0.09054907 | 0.374964 | 0.7076873 |
| Nucleus of the optic tract | -0.005428492 | 0.1018012 | -0.05332442 | 0.9574734 |
| Nucleus of the posterior commissure | 0.1238715 | 0.1051018 | 1.178586 | 0.2385632 |
| Posterior pretectal nucleus | -0.002998761 | 0.07950271 | -0.03771897 | 0.96991175 |
| Cuneiform nucleus | 0.05909191 | 0.0670748 | 0.8809852 | 0.37832583 |
| Red nucleus | -0.1751559 | 0.1093433 | -1.6018889 | 0.1091802 |
| Substantia nigra, compact part | -0.12677315 | 0.1520086 | -0.8339866 | 0.4042885 |
| Pedunculopontine nucleus | -0.008596527 | 0.08285333 | -0.103756 | 0.917363 |
| Dorsal nucleus raphe | -0.01480811 | 0.1389906 | -0.1065404 | 0.9151536 |
| Nucleus of the lateral lemniscus | -0.1650233 | 0.2349852 | -0.7022709 | 0.4825102 |
| Principal sensory nucleus of the trigeminal | 0.13751063 | 0.09669903 | 1.4220477 | 0.1550124 |
| Parabrachial nucleus | -0.07221681 | 0.1219731 | -0.5920715 | 0.5538027 |
| Superior olivary complex | -0.24167176 | 0.1964354 | -1.23028604 | 0.21859 |
| Dorsal tegmental nucleus | -0.02640416 | 0.1814597 | -0.1455098 | 0.8843084 |
| Pontine central gray | -0.003831885 | 0.1345471 | -0.02847988 | 0.977279413 |
| Pontine gray | -0.07560482 | 0.1650892 | -0.4579633 | 0.6469788 |
| Supratrigeminal nucleus | -0.04403334 | 0.1316026 | -0.3345933 | 0.7379319 |

| | | | | |
|---|---|---|---|---|
| Tegmental reticular nucleus | -0.7043041 | 0.6585238 | -1.0695196 | 0.2848356 |
| Motor nucleus of trigeminal | -0.02734259 | 0.1313643 | -0.2081433 | 0.8351171 |
| Superior central nucleus raphe | -0.6198934 | 0.5673013 | -1.092706 | 0.2745229 |
| Laterodorsal tegmental nucleus | -0.06368266 | 0.1672526 | -0.3807574 | 0.7033833 |
| Nucleus incertus | 0.0193313 | 0.1207085 | 0.1601487 | 0.87276397 |
| Pontine reticular nucleus | -0.2194759 | 0.2284048 | -0.9609077 | 0.3365986 |
| Dorsal cochlear nucleus | 0.01304403 | 0.1387095 | 0.09403846 | 0.9250786 |
| Ventral cochlear nucleus | 0.1659023 | 0.1406008 | 1.179952 | 0.23801919 |
| Cuneate nucleus | 0.27182643 | 0.2310285 | 1.1765928 | 0.2393581 |
| External cuneate nucleus | 0.08727849 | 0.1421712 | 0.6138972 | 0.5392833 |
| Nucleus of the trapezoid body | -0.4989683 | 0.4377726 | -1.139789 | 0.2543743 |
| Nucleus of the solitary tract | 0.07599482 | 0.1461172 | 0.5200948 | 0.6029975 |
| Spinal nucleus of the trigeminal | -0.01205235 | 0.1596045 | -0.07551382 | 0.9398059 |
| Facial motor nucleus | -0.05283062 | 0.1629856 | -0.3241428 | 0.7458299 |
| Dorsal motor nucleus of the vagus nerve | 0.1477514 | 0.2807947 | 0.5261902 | 0.59875609 |
| Gigantocellular reticular nucleus | 0.10817392 | 0.1459695 | 0.74107214 | 0.4586497 |
| Inferior olivary complex | -0.110099 | 0.6159553 | -0.178745 | 0.8581379 |
| Intermediate reticular nucleus | 0.0797491 | 0.1436429 | 0.5551898 | 0.5787648 |
| Lateral reticular nucleus | 0.3111742 | 0.2784899 | 1.1173625 | 0.2638394 |
| Magnocellular reticular nucleus | 0.02510036 | 0.2666601 | 0.09412869 | 0.9250069 |
| Medullary reticular nucleus | 0.07631515 | 0.1628739 | 0.4685534 | 0.6393889 |
| Parvicellular reticular nucleus | 0.04708099 | 0.122602 | 0.3840149 | 0.7009674 |
| Paragigantocellular reticular nucleus | 0.1185748 | 0.1480001 | 0.8011806 | 0.4230271 |
| Nucleus prepositus | 0.18262742 | 0.1559823 | 1.17082139 | 0.2416706 |
| Lateral vestibular nucleus | -0.006885964 | 0.1118596 | -0.06155902 | 0.950914 |
| Medial vestibular nucleus | 0.04056485 | 0.1396542 | 0.2904664 | 0.7714595 |
| Spinal vestibular nucleus | -0.049489211 | 0.1134345 | -0.43628017 | 0.6626335 |
| Superior vestibular nucleus | -0.2035824 | 0.2203693 | -0.9238239 | 0.355578 |
| Hypoglossal nucleus | 0.3797446 | 0.1870445 | 2.030236 | 0.04233253 |
| Nucleus raphe magnus | 0.18528409 | 0.3101586 | 0.5973851 | 0.5502503 |

**Supplementary Data Table 64:** GLMM coefficient estimates of ***Stim*** contribution to Fos$^{+}$ cell counts across all mice (**Figure S8A**). The total number of Fos$^{+}$ cells in each brain region was modeled as ***Counts ~ SI Time + Stim + SI Time:Stim* + ln(*Total Counts*) + (*1+SI Time*|*Cohort*)** using a negative binomial link function (*N* = 54 mice). The critical *p*-value for significance while permitting a 10% FDR was 0.000046; 1 region was significant.

| Brain region name | Stim coefficient estimate | Stim coefficient std error | Stim coefficient Z-statistic | Stim coefficient P-value |
|---|---|---|---|---|
| Frontal pole | 0.4426291 | 0.5430733 | 0.8150449 | 0.4150466 |
| Primary motor area | -0.562061357 | 0.2221522 | -2.5300731 | 0.01140388 |

| | | | | |
|---|---|---|---|---|
| Secondary motor area | -0.25948675 | 0.10519181 | -2.466796 | 0.0136328 |
| Primary somatosensory area, nose | -0.436114 | 0.3481784 | -1.252559 | 0.2103662 |
| Primary somatosensory area, barrel field | -0.44453024 | 0.18254361 | -2.4352002 | 0.01488355 |
| Primary somatosensory area, lower limb | -0.58617995 | 0.2308095 | -2.5396703 | 0.0110957 |
| Primary somatosensory area, mouth | -0.3823107 | 0.4215946 | -0.9068206 | 0.3645017 |
| Primary somatosensory area, upper limb | -0.446434 | 0.24532152 | -1.8197915 | 0.06879077 |
| Primary somatosensory area, trunk | -0.34871731 | 0.16212925 | -2.1508599 | 0.03148726 |
| Supplemental somatosensory area | -0.32525663 | 0.2119438 | -1.5346365 | 0.1248732 |
| Gustatory areas | 0.24961667 | 0.2487453 | 1.0035032 | 0.3156182 |
| Visceral area | -0.05327407 | 0.26544994 | -0.2006935 | 0.8409383 |
| Dorsal auditory area | -0.3018395 | 0.2619007 | -1.15249619 | 0.2491172 |
| Primary auditory area | -0.41114943 | 0.3619692 | -1.1358686 | 0.2560115 |
| Posterior auditory area | 0.07894997 | 0.5257554 | 0.1501648 | 0.8806346 |
| Ventral auditory area | 0.055650927 | 0.2754182 | 0.20205972 | 0.83987 |
| Anterolateral visual area | 0.37827867 | 0.4432742 | 0.8533739 | 0.393452 |
| Anteromedial visual area | 0.19429227 | 0.3575782 | 0.543356 | 0.5868847 |
| Lateral visual area | 0.5920809 | 0.4789456 | 1.236218 | 0.2163777 |
| Primary visual area | 0.57423398 | 0.400159 | 1.4350147 | 0.1512829 |
| Posterolateral visual area | 0.0001938621 | 0.469639 | 0.0004127897 | 0.9996706 |
| Posteromedial visual area | 0.4581661 | 0.4186095 | 1.0944954 | 0.2737378 |
| Anterior cingulate area, dorsal part | -0.060503461 | 0.10121129 | -0.59779358 | 0.549977678 |
| Anterior cingulate area, ventral part | 0.03055002 | 0.14849034 | 0.2057374 | 0.8369960306 |
| Prelimbic area | -0.08209342 | 0.17782168 | -0.46166149 | 0.6443241 |
| Infralimbic area | -0.25933672 | 0.15073828 | -1.7204437 | 0.08535182 |
| Orbital area, lateral part | -0.35601615 | 0.20342289 | -1.7501283 | 0.08009619 |
| Orbital area, medial part | 0.02087956 | 0.2602235 | 0.08023701 | 0.9360488 |
| Orbital area, ventrolateral part | 0.06805414 | 0.18416278 | 0.3695326 | 0.71173081 |
| Agranular insular area, dorsal part | -0.12197678 | 0.29770096 | -0.4097292 | 0.6820046 |
| Agranular insular area, posterior part | -0.15117727 | 0.28501776 | -0.5304135 | 0.5958253 |
| Agranular insular area, ventral part | -0.06490639 | 0.2979395 | -0.2178509 | 0.8275453 |
| Retrosplenial area, lateral agranular part | 0.18730522 | 0.2531769 | 0.7398196 | 0.4594095 |
| Retrosplenial area, dorsal part | 0.19291135 | 0.2575398 | 0.7490545 | 0.4538243 |
| Retrosplenial area, ventral part | -0.18079521 | 0.12294482 | -1.4705395 | 0.1414157 |
| Rostrolateral visual area | 0.24783881 | 0.3593444 | 0.6896971 | 0.4903847 |
| Temporal association areas | 0.22297219 | 0.2460902 | 0.9060587 | 0.3649048 |
| Perirhinal area | 0.07736879 | 0.4504886 | 0.1717442 | 0.8636386 |
| Ectorhinal area | 0.006318135 | 0.3301864 | 0.01913505 | 0.9847334 |
| Anterior olfactory nucleus | -0.3413431 | 0.3261311 | -1.0466439 | 0.2952639 |
| Taenia tecta, dorsal part | -0.18649304 | 0.1370198 | -1.3610664 | 0.1734927 |
| Taenia tecta, ventral part | -0.69286611 | 0.4245279 | -1.6320862 | 0.1026613 |
| Dorsal peduncular area | -0.43622765 | 0.17748469 | -2.457833 | 0.01397783 |
| Piriform area | -0.048679028 | 0.4221246 | -0.11531909 | 0.9081922 |

| | | | | |
|---|---|---|---|---|
| Nucleus of the lateral olfactory tract | 0.5402154 | 0.7865159 | 0.6868461 | 0.4921797 |
| Cortical amygdalar area, anterior part | 0.3723464 | 0.6813558 | 0.5464787 | 0.584737 |
| Cortical amygdalar area, posterior part | 0.2455821 | 0.7307785 | 0.3360554 | 0.7368291 |
| Piriform,amygdalar area | -0.1221503 | 0.6013901 | -0.20311324 | 0.8390465 |
| Postpiriform transition area | 0.07199079 | 0.7007295 | 0.1027369 | 0.9181718 |
| Field c a1 | -0.22209135 | 0.1413913 | -1.5707569 | 0.1162391 |
| Field c a2 | -0.5480861 | 0.22336599 | -2.4537581 | 0.0141372 |
| Field c a3 | -0.2956507 | 0.17634613 | -1.6765364 | 0.09363318 |
| Dentate gyrus | -0.09137129 | 0.19981532 | -0.4572787 | 0.6474707 |
| Induseum griseum | 0.13761116 | 0.3436637 | 0.4004239 | 0.6888443 |
| Entorhinal area, lateral part | 0.01404744 | 0.5319298 | 0.02640845 | 0.9789316 |
| Entorhinal area, medial part | 0.12661377 | 0.25313259 | 0.5001876 | 0.616943 |
| Parasubiculum | 0.419661902 | 0.21685244 | 1.93524174 | 0.05296064 |
| Postsubiculum | 0.01341043 | 0.195652 | 0.06854228 | 0.945354 |
| Presubiculum | 0.26238668 | 0.18202804 | 1.441463 | 0.1494539 |
| Subiculum | -0.01622799 | 0.17426451 | -0.09312275 | 0.925806 |
| Prosubiculum | 0.01493867 | 0.12175768 | 0.1226918 | 0.9023512 |
| Claustrum | 0.0560559 | 0.17589334 | 0.3186926 | 0.7499597 |
| Endopiriform nucleus, dorsal part | 0.06167727 | 0.3460003 | 0.1782578 | 0.8585205 |
| Endopiriform nucleus, ventral part | -0.04838818 | 0.5711218 | -0.08472479 | 0.9324802 |
| Lateral amygdalar nucleus | 0.335299097 | 0.27139305 | 1.23547414 | 0.2166541 |
| Basolateral amygdalar nucleus, anterior part | 0.34672299 | 0.4281728 | 0.8097735 | 0.4180704 |
| Basolateral amygdalar nucleus, posterior part | 0.2803123 | 0.5415219 | 0.517638 | 0.6047109 |
| Basolateral amygdalar nucleus, ventral part | 0.1168356 | 0.5741749 | 0.2034843 | 0.8387565 |
| Basomedial amygdalar nucleus, anterior part | 0.3379701 | 0.7435735 | 0.4545214 | 0.6494536 |
| Basomedial amygdalar nucleus, posterior part | 0.2901147 | 0.5867703 | 0.4944264 | 0.6210051 |
| Posterior amygdalar nucleus | 0.4364702 | 0.5795388 | 0.7531338 | 0.4513695 |
| Caudoputamen | -0.005307495 | 0.24014421 | -0.02210128 | 0.9823672 |
| Nucleus accumbens | -0.13952137 | 0.30180358 | -0.462292 | 0.643872 |
| Fundus of striatum | -0.051036743 | 0.25047862 | -0.20375689 | 0.8385435 |
| Olfactory tubercle | -0.1466278 | 0.5285142 | -0.2774339 | 0.7814469 |
| Lateral septal nucleus | 0.11275619 | 0.23274353 | 0.4844654 | 0.6280556 |
| Septofimbrial nucleus | 0.49286304 | 0.4663487 | 1.0568551 | 0.2905777 |
| Anterior amygdalar area | 0.45223323 | 0.4881314 | 0.92645801 | 0.354208 |
| Central amygdalar nucleus, capsular part | 0.495502 | 0.2564506 | 1.932154 | 0.05334055 |
| Central amygdalar nucleus, lateral part | -0.198396 | 0.2336298 | -0.8491897 | 0.3957757 |
| Central amygdalar nucleus, medial part | 0.4459695 | 0.321397 | 1.3875971 | 0.1652598 |
| Intercalated amygdalar nucleus | 0.1352833 | 0.4802978 | 0.2816655 | 0.7782 |
| Medial amygdalar nucleus | 0.7233412 | 0.5688429 | 1.2716011 | 0.2035149 |
| Globus pallidus, external segment | -0.28168958 | 0.3806719 | -0.7399799 | 0.4593122 |
| Globus pallidus, internal segment | -0.68252688 | 0.6385985 | -1.0687887 | 0.2851649 |
| Substantia innominata | 0.118209077 | 0.22557457 | 0.52403548 | 0.6002539 |

| | | | | |
|---|---|---|---|---|
| Magnocellular nucleus | 0.008276661 | 0.4332723 | 0.01910268 | 0.9847592 |
| Medial septal nucleus | -0.22176809 | 0.2338871 | -0.9481844 | 0.3430356 |
| Diagonal band nucleus | 0.03837666 | 0.31757056 | 0.1208445 | 0.9038142 |
| Triangular nucleus of septum | 0.8865247 | 0.4291009 | 2.0660053 | 0.03882798 |
| Bed nuclei of the stria terminalis | -0.18626934 | 0.23377357 | -0.7967939 | 0.4255708 |
| Ventral anterior,lateral complex of the thalamus | 1.96923013 | 0.4836437 | 4.071655 | 4.668029E-05 |
| Ventral medial nucleus of the thalamus | -0.17089499 | 0.2874277 | -0.5945668 | 0.5521331 |
| Ventral posterior complex of the thalamus | 0.3371586 | 0.26278575 | 1.2830171 | 0.1994861 |
| Subparafascicular nucleus | 0.5657772 | 0.34030754 | 1.6625467 | 0.0964032 |
| Subparafascicular area | -0.10885182 | 0.9184069 | -0.1185224 | 0.9056537 |
| Medial geniculate complex | 0.21009096 | 0.26162321 | 0.8030288 | 0.4219581 |
| Lateral geniculate complex | 0.23596808 | 0.08941141 | 2.6391273 | 0.008311975 |
| Lateral posterior nucleus of the thalamus | 0.62199265 | 0.22041714 | 2.8218888 | 0.004774173 |
| Posterior complex of the thalamus | -0.5365185 | 0.2794646 | -1.919809 | 0.05488209 |
| Posterior limiting nucleus of the thalamus | 0.274261 | 0.196333 | 1.3969172 | 0.1624385 |
| Suprageniculate nucleus | 0.2368909 | 0.22300095 | 1.0622865 | 0.2881056 |
| Anteroventral nucleus of thalamus | 1.139887926 | 0.4541916 | 2.50970744 | 0.01208312 |
| Anteromedial nucleus | 0.8582153 | 0.4525711 | 1.89631 | 0.05791902 |
| Anterodorsal nucleus | 0.6220877 | 0.3445938 | 1.805278 | 0.07103114 |
| Interanterodorsal nucleus of the thalamus | 0.64056984 | 0.3509632 | 1.8251766 | 0.06797438 |
| Lateral dorsal nucleus of thalamus | 0.54464776 | 0.3705738 | 1.4697417 | 0.1416317 |
| Intermediodorsal nucleus of the thalamus | 0.3301968 | 0.416118 | 0.7935172 | 0.4274766 |
| Mediodorsal nucleus of thalamus | 0.33900704 | 0.24913894 | 1.3607148 | 0.1736038 |
| Submedial nucleus of the thalamus | -0.188399069 | 0.6568479 | -0.28682298 | 0.7742479 |
| Perireunensis nucleus | -0.31867198 | 0.3111094 | -1.0243083 | 0.3056897 |
| Paraventricular nucleus of the thalamus | 0.54554364 | 0.2336956 | 2.3344198 | 0.01957375 |
| Parataenial nucleus | 0.45927485 | 0.24285101 | 1.8911795 | 0.05860038 |
| Nucleus of reuniens | -0.458216 | 0.20877554 | -2.1947783 | 0.02817951 |
| Central medial nucleus of the thalamus | 0.23802536 | 0.3750431 | 0.6346614 | 0.5256493 |
| Paracentral nucleus | -0.63011053 | 0.4517867 | -1.3947081 | 0.1631039 |
| Central lateral nucleus of the thalamus | 0.56414108 | 0.31522684 | 1.7896353 | 0.07351256 |
| Parafascicular nucleus | 0.10831662 | 0.363577 | 0.2979194 | 0.7657647 |
| Reticular nucleus of the thalamus | 0.3949921 | 0.33399538 | 1.1826274 | 0.2369569 |
| Medial habenula | 0.7012075 | 0.328508 | 2.134522 | 0.03280011 |
| Lateral habenula | 0.16946198 | 0.37907801 | 0.4470372 | 0.6548482 |
| Paraventricular hypothalamic nucleus | 0.10961495 | 0.25874153 | 0.4236465 | 0.6718236 |
| Periventricular hypothalamic nucleus | 0.19785625 | 0.4432933 | 0.4463326 | 0.655357 |
| Arcuate hypothalamic nucleus | 0.7369641 | 0.7451613 | 0.9889994 | 0.3226634 |
| Anterodorsal preoptic nucleus | 0.19731017 | 0.28101002 | 0.7021464 | 0.4825879 |
| Anteroventral periventricular nucleus | -0.0237025 | 0.4683611 | -0.05060731 | 0.9596384 |
| Dorsomedial hypothalamic nucleus | 0.05262437 | 0.313262 | 0.1679883 | 0.8665925 |
| Medial preoptic area | -0.03255195 | 0.22526086 | -0.1445078 | 0.8850995 |

| | | | | |
|---|---|---|---|---|
| Subparaventricular zone | 0.1087871 | 0.4148113 | 0.2622568 | 0.7931235 |
| Anterior hypothalamic nucleus | 0.002064889 | 0.2925826 | 0.007057459 | 0.994369 |
| Supramammillary nucleus | 0.9422979 | 0.8276457 | 1.1385281 | 0.2549 |
| Tuberomammillary nucleus | 1.4477174 | 0.9756055 | 1.483917 | 0.137831 |
| Medial preoptic nucleus | -0.00187075 | 0.2929893 | -0.006385046 | 0.9949055 |
| Dorsal premammillary nucleus | 1.09484647 | 0.7587378 | 1.442984 | 0.149025 |
| Ventral premammillary nucleus | 1.20571849 | 1.2575126 | 0.9588123 | 0.3376533 |
| Ventromedial hypothalamic nucleus | 0.2479778 | 0.78062 | 0.3176678 | 0.750737 |
| Posterior hypothalamic nucleus | 0.27335769 | 0.2942727 | 0.9289265 | 0.3529272 |
| Lateral hypothalamic area | 0.08569068 | 0.2878094 | 0.2977342 | 0.7659061 |
| Lateral preoptic area | -0.2021547 | 0.22814252 | -0.8860896 | 0.3755693 |
| Parasubthalamic nucleus | 0.1820977 | 0.3932613 | 0.463045 | 0.6433321 |
| Retrochiasmatic area | 0.07383277 | 0.6742551 | 0.1095027 | 0.9128038 |
| Subthalamic nucleus | -0.44240514 | 0.4231402 | -1.0455285 | 0.2957788 |
| Tuberal nucleus | 0.3745652 | 0.8723087 | 0.4293952 | 0.6676356 |
| Zona incerta | -0.29330238 | 0.2217709 | -1.3225468 | 0.1859861 |
| Superior colliculus, dorsal part | 0.28118852 | 0.24740611 | 1.1365464 | 0.255728 |
| Inferior colliculus, central part | -0.68676476 | 0.21581033 | -3.1822608 | 0.001461302 |
| Inferior colliculus, dorsal part | -0.57219787 | 0.2466159 | -2.32019865 | 0.02033013 |
| Inferior colliculus, external part | -0.42597827 | 0.242301 | -1.7580542 | 0.07873829 |
| Substantia nigra, reticular part | 0.1482141 | 0.3296724 | 0.4495799 | 0.65301338 |
| Ventral tegmental area | -0.59961096 | 0.358054 | -1.6746383 | 0.09400521 |
| Midbrain reticular nucleus | -0.234842565 | 0.22846467 | -1.0279163 | 0.3039892 |
| Superior colliculus, ventral part | -0.0554211724 | 0.15001558 | -0.3694361 | 0.7118027 |
| Periaqueductal gray | 0.26940309 | 0.16665009 | 1.61657929 | 0.1059691 |
| Anterior pretectal nucleus | 0.22649691 | 0.32261823 | 0.7020586 | 0.4826426 |
| Nucleus of the optic tract | 0.327748734 | 0.2401819 | 1.36458559 | 0.1723833 |
| Nucleus of the posterior commissure | 0.5497035 | 0.4080829 | 1.347039 | 0.1779678 |
| Posterior pretectal nucleus | 0.399007465 | 0.19654999 | 2.03005589 | 0.04235086 |
| Cuneiform nucleus | 0.36609612 | 0.2124738 | 1.7230181 | 0.08488525 |
| Red nucleus | -0.1553841 | 0.3676771 | -0.4226103 | 0.6725796 |
| Substantia nigra, compact part | 0.04747939 | 0.3637794 | 0.130517 | 0.8961574 |
| Pedunculopontine nucleus | 0.040643461 | 0.25079678 | 0.1620573 | 0.8712607 |
| Dorsal nucleus raphe | -0.23327387 | 0.3642338 | -0.6404509 | 0.5218795 |
| Nucleus of the lateral lemniscus | 0.2579272 | 0.3091448 | 0.8343249 | 0.4040979 |
| Principal sensory nucleus of the trigeminal | -0.09397853 | 0.33132671 | -0.2836431 | 0.7766839 |
| Parabrachial nucleus | 0.44801913 | 0.2211763 | 2.0256199 | 0.04280375 |
| Superior olivary complex | 0.03493938 | 0.5787314 | 0.06037236 | 0.9518591 |
| Dorsal tegmental nucleus | -0.46669062 | 0.4498909 | -1.0373417 | 0.2995767 |
| Pontine central gray | -0.014898005 | 0.2454065 | -0.06070747 | 0.9515921812 |
| Pontine gray | 0.26952529 | 0.6200075 | 0.4347129 | 0.6637708 |
| Supratrigeminal nucleus | 0.23077277 | 0.2894625 | 0.7972459 | 0.4253082 |

| | | | | |
|---|---|---|---|---|
| Tegmental reticular nucleus | -0.20267 | 0.5230711 | -0.3874617 | 0.6984145 |
| Motor nucleus of trigeminal | -0.10205343 | 0.3784696 | -0.2696476 | 0.7874313 |
| Superior central nucleus raphe | -0.5783528 | 0.401464 | -1.440609 | 0.1496951 |
| Laterodorsal tegmental nucleus | -0.35227129 | 0.3423152 | -1.0290844 | 0.30344 |
| Nucleus incertus | 0.3341309 | 0.2390728 | 1.3976113 | 0.16222982 |
| Pontine reticular nucleus | 0.217975 | 0.2714528 | 0.8029941 | 0.4219782 |
| Dorsal cochlear nucleus | -0.47046998 | 0.2871189 | -1.63858924 | 0.1012988 |
| Ventral cochlear nucleus | -0.7009772 | 0.3131295 | -2.238617 | 0.02518082 |
| Cuneate nucleus | -0.38500298 | 0.6760075 | -0.5695247 | 0.5690001 |
| External cuneate nucleus | -0.21998643 | 0.5267441 | -0.4176344 | 0.6762145 |
| Nucleus of the trapezoid body | 1.3525976 | 0.7527702 | 1.796827 | 0.07236313 |
| Nucleus of the solitary tract | -0.01573293 | 0.4634865 | -0.03394475 | 0.9729212 |
| Spinal nucleus of the trigeminal | 0.03855514 | 0.3374901 | 0.11424081 | 0.9090469 |
| Facial motor nucleus | 0.71516978 | 0.4944177 | 1.4464891 | 0.1480401 |
| Dorsal motor nucleus of the vagus nerve | 0.5327978 | 0.5217076 | 1.0212576 | 0.30713241 |
| Gigantocellular reticular nucleus | -0.02703528 | 0.4603919 | -0.05872232 | 0.9531733 |
| Inferior olivary complex | 1.0971142 | 1.0526595 | 1.0422309 | 0.2973047 |
| Intermediate reticular nucleus | 0.154744 | 0.4584569 | 0.3375323 | 0.7357157 |
| Lateral reticular nucleus | 1.255255 | 0.5711381 | 2.1978135 | 0.0279624 |
| Magnocellular reticular nucleus | 0.96734788 | 0.765289 | 1.26402947 | 0.2062195 |
| Medullary reticular nucleus | 0.61991997 | 0.4878333 | 1.2707619 | 0.2038134 |
| Parvicellular reticular nucleus | 0.07187669 | 0.3849772 | 0.1867037 | 0.8518929 |
| Paragigantocellular reticular nucleus | 0.1696412 | 0.3955927 | 0.4288279 | 0.6680485 |
| Nucleus prepositus | 0.006954042 | 0.3069464 | 0.02265556 | 0.981925 |
| Lateral vestibular nucleus | -0.323179944 | 0.3650094 | -0.8854017 | 0.37594 |
| Medial vestibular nucleus | 0.16306432 | 0.2695229 | 0.605011 | 0.5451717 |
| Spinal vestibular nucleus | -0.170000723 | 0.2833208 | -0.60002909 | 0.5484868 |
| Superior vestibular nucleus | -0.2650012 | 0.3988982 | -0.6643329 | 0.5064773 |
| Hypoglossal nucleus | 0.4770088 | 0.5698165 | 0.837127 | 0.4025212 |
| Nucleus raphe magnus | 0.4690844 | 0.6910528 | 0.6787968 | 0.4972666 |

**Supplementary Data Table 65:** GLMM coefficient estimates of ***SI Time:Stim*** interaction contribution to Fos$^+$ cell counts, which captures the contribution of SI time in LHb-stimulated mice (**Figure S8A**). The total number of Fos$^+$ cells in each brain region was modeled as ***Counts ~ SI Time + Stim + SI Time:Stim* + ln(*Total Counts*) + (*1+SI Time*|*Cohort*)** using a negative binomial link function (*N* = 10 mice). The critical *p*-value for significance while permitting a 10% FDR was 0.0108; 22 regions were significant.

| Brain region name | SI:Stim coefficient estimate | SI:Stim coefficient std error | SI:Stim coefficient Z-statistic | SI:Stim coefficient P-value |
|---|---|---|---|---|
| Frontal pole | -0.6003859 | 0.3009757 | -1.9947988 | 0.04606483 |

| | | | | |
|---|---|---|---|---|
| Primary motor area | -0.006325648 | 0.1537393 | -0.04114528 | 0.9671801 |
| Secondary motor area | 0.15015587 | 0.08919164 | 1.68352 | 0.09227455 |
| Primary somatosensory area, nose | 0.2649597 | 0.2228576 | 1.1889194 | 0.2344714 |
| Primary somatosensory area, barrel field | 0.290537979 | 0.14257287 | 2.0378209 | 0.04156784 |
| Primary somatosensory area, lower limb | 0.01066322 | 0.2051755 | 0.05197123 | 0.9585516 |
| Primary somatosensory area, mouth | 0.1118432 | 0.2551192 | 0.4383959 | 0.6610993 |
| Primary somatosensory area, upper limb | 0.04911854 | 0.16087797 | 0.3053155 | 0.7601258 |
| Primary somatosensory area, trunk | 0.07918124 | 0.13684763 | 0.5786088 | 0.56285315 |
| Supplemental somatosensory area | 0.1226987 | 0.1320488 | 0.9291922 | 0.3527895 |
| Gustatory areas | -0.19389357 | 0.1970494 | -0.9839847 | 0.325123 |
| Visceral area | 0.05632929 | 0.13607387 | 0.4139611 | 0.6789026 |
| Dorsal auditory area | 0.17361618 | 0.2021536 | 0.85883295 | 0.3904327 |
| Primary auditory area | -0.1160242 | 0.2069751 | -0.5605708 | 0.5750902 |
| Posterior auditory area | -0.03144794 | 0.462263 | -0.0680304 | 0.9457614 |
| Ventral auditory area | -0.12796475 | 0.1633322 | -0.78346291 | 0.4333553 |
| Anterolateral visual area | 0.49809825 | 0.3988772 | 1.248751 | 0.2117562 |
| Anteromedial visual area | 0.70654381 | 0.2514671 | 2.8096867 | 0.004958975 |
| Lateral visual area | 0.5535882 | 0.4360039 | 1.269686 | 0.2041964 |
| Primary visual area | 0.72684243 | 0.3874366 | 1.8760296 | 0.06065122 |
| Posterolateral visual area | 0.5518385338 | 0.2739941 | 2.0140527636 | 0.04400399 |
| Posteromedial visual area | 0.8341306 | 0.3217293 | 2.5926471 | 0.009524045 |
| Anterior cingulate area, dorsal part | 0.24043406 | 0.07934036 | 3.03041306 | 0.002442195 |
| Anterior cingulate area, ventral part | 0.34575449 | 0.09376646 | 3.6874003 | 0.0002265568 |
| Prelimbic area | -0.005850432 | 0.14236333 | -0.04109508 | 0.9672201 |
| Infralimbic area | 0.0557542 | 0.11749643 | 0.4745183 | 0.63513038 |
| Orbital area, lateral part | 0.30148997 | 0.12436934 | 2.4241502 | 0.01534426 |
| Orbital area, medial part | -0.24463001 | 0.1665128 | -1.46913663 | 0.1417957 |
| Orbital area, ventrolateral part | 0.37986432 | 0.15337992 | 2.4766235 | 0.01326317 |
| Agranular insular area, dorsal part | 0.03588516 | 0.15679888 | 0.2288611 | 0.8189769 |
| Agranular insular area, posterior part | 0.32961124 | 0.1660731 | 1.9847359 | 0.04717387 |
| Agranular insular area, ventral part | -0.03672896 | 0.2079391 | -0.1766332 | 0.8597965 |
| Retrosplenial area, lateral agranular part | 0.31941271 | 0.1849716 | 1.7268207 | 0.08419987 |
| Retrosplenial area, dorsal part | -0.18819543 | 0.1715879 | -1.0967873 | 0.2727344 |
| Retrosplenial area, ventral part | -0.06721344 | 0.09956192 | -0.6750919 | 0.4996174 |
| Rostrolateral visual area | 0.5336878 | 0.2234108 | 2.3888185 | 0.01690265 |
| Temporal association areas | -0.220171 | 0.1777927 | -1.238358 | 0.2155833 |
| Perirhinal area | 0.23069863 | 0.2451773 | 0.9409461 | 0.3467325 |
| Ectorhinal area | -0.23405455 | 0.2400005 | -0.97522516 | 0.3294486 |
| Anterior olfactory nucleus | 0.14478798 | 0.2505297 | 0.5779274 | 0.5633132 |
| Taenia tecta, dorsal part | 0.05620632 | 0.11682563 | 0.4811129 | 0.6304363 |
| Taenia tecta, ventral part | -0.59329952 | 0.2864105 | -2.0715003 | 0.03831206 |
| Dorsal peduncular area | 0.21050272 | 0.12116686 | 1.737296 | 0.08233491 |

| | | | | |
|---|---|---|---|---|
| Piriform area | 0.278691581 | 0.3080372 | 0.90473342 | 0.3656066 |
| Nucleus of the lateral olfactory tract | 0.886327 | 0.6488628 | 1.3659697 | 0.1719485 |
| Cortical amygdalar area, anterior part | 0.8502402 | 0.5773224 | 1.4727303 | 0.1408238 |
| Cortical amygdalar area, posterior part | 0.4057969 | 0.6626842 | 0.6123535 | 0.5403039 |
| Piriform,amygdalar area | 0.01884123 | 0.4302467 | 0.04379168 | 0.9650705 |
| Postpiriform transition area | 0.59958531 | 0.4774075 | 1.2559193 | 0.2091452 |
| Field c a1 | -0.09475606 | 0.09763444 | -0.9705188 | 0.331788 |
| Field c a2 | -0.21420328 | 0.14949414 | -1.4328541 | 0.1518996 |
| Field c a3 | -0.05380282 | 0.10996855 | -0.4892565 | 0.6246601 |
| Dentate gyrus | -0.29810495 | 0.13123695 | -2.2715017 | 0.02311662 |
| Induseum griseum | -0.44996353 | 0.2812582 | -1.5998237 | 0.1096377 |
| Entorhinal area, lateral part | 0.16545112 | 0.3006639 | 0.55028592 | 0.5821233 |
| Entorhinal area, medial part | 0.2533183 | 0.16942219 | 1.4951897 | 0.134865 |
| Parasubiculum | -0.325632502 | 0.18599293 | -1.75077893 | 0.079984 |
| Postsubiculum | 0.02664849 | 0.1134865 | 0.23481639 | 0.8143512 |
| Presubiculum | -0.13600974 | 0.10044329 | -1.354095 | 0.1757061 |
| Subiculum | -0.13919541 | 0.12194856 | -1.14142733 | 0.2536921 |
| Prosubiculum | 0.12362497 | 0.10602964 | 1.1659473 | 0.2436357 |
| Claustrum | 0.12369958 | 0.13107282 | 0.943747 | 0.3452989 |
| Endopiriform nucleus, dorsal part | 0.38223517 | 0.2610431 | 1.464261 | 0.1431226 |
| Endopiriform nucleus, ventral part | 0.79305623 | 0.3810809 | 2.08107053 | 0.03742745 |
| Lateral amygdalar nucleus | -0.269920371 | 0.17095476 | -1.57889946 | 0.1143591 |
| Basolateral amygdalar nucleus, anterior part | 0.26338469 | 0.272194 | 0.9676359 | 0.3332263 |
| Basolateral amygdalar nucleus, posterior part | 0.2672877 | 0.3675385 | 0.7272373 | 0.4670806 |
| Basolateral amygdalar nucleus, ventral part | 0.6332527 | 0.572505 | 1.1061085 | 0.2686796 |
| Basomedial amygdalar nucleus, anterior part | 0.7062184 | 0.6453008 | 1.0944019 | 0.2737787 |
| Basomedial amygdalar nucleus, posterior part | 0.5412841 | 0.4406588 | 1.2283519 | 0.2193149 |
| Posterior amygdalar nucleus | 0.1203852 | 0.4662955 | 0.2581737 | 0.7962729 |
| Caudoputamen | -0.22944468 | 0.20710306 | -1.10787681 | 0.267915 |
| Nucleus accumbens | 0.21065498 | 0.19134206 | 1.100934 | 0.2709254 |
| Fundus of striatum | -0.005585185 | 0.2394606 | -0.02332403 | 0.9813918 |
| Olfactory tubercle | -0.2633965 | 0.3767224 | -0.6991793 | 0.48444 |
| Lateral septal nucleus | 0.37526781 | 0.14718641 | 2.5496092 | 0.01078437 |
| Septofimbrial nucleus | 0.23551854 | 0.185563 | 1.2692104 | 0.204366 |
| Anterior amygdalar area | 0.01145345 | 0.3581238 | 0.03198181 | 0.9744866 |
| Central amygdalar nucleus, capsular part | -0.2338823 | 0.2052933 | -1.13926 | 0.25459486 |
| Central amygdalar nucleus, lateral part | 0.0945056 | 0.21494287 | 0.4396778 | 0.6601705 |
| Central amygdalar nucleus, medial part | 0.4294781 | 0.2952903 | 1.4544264 | 0.1458281 |
| Intercalated amygdalar nucleus | 0.4254669 | 0.3596884 | 1.1828762 | 0.2368582 |
| Medial amygdalar nucleus | 0.3691845 | 0.4616104 | 0.7997752 | 0.423841 |
| Globus pallidus, external segment | -0.24966655 | 0.1877288 | -1.3299322 | 0.1835406 |
| Globus pallidus, internal segment | 0.41580692 | 0.3534556 | 1.1764049 | 0.2394331 |

| | | | | |
|---|---|---|---|---|
| Substantia innominata | -0.13027515 | 0.20684413 | -0.62982281 | 0.5288105 |
| Magnocellular nucleus | -0.428506416 | 0.2644215 | -1.62054325 | 0.1051156 |
| Medial septal nucleus | 0.08092318 | 0.14230831 | 0.5686469 | 0.56959577 |
| Diagonal band nucleus | -0.15032542 | 0.17439662 | -0.8619744 | 0.3887016 |
| Triangular nucleus of septum | -0.2281759 | 0.1977693 | -1.1537477 | 0.2486036 |
| Bed nuclei of the stria terminalis | 0.18862533 | 0.18734455 | 1.0068365 | 0.3140133 |
| Ventral anterior,lateral complex of the thalamus | 0.57077686 | 0.3764818 | 1.5160809 | 0.1294989 |
| Ventral medial nucleus of the thalamus | 0.289409 | 0.295045 | 0.9808977 | 0.3266432 |
| Ventral posterior complex of the thalamus | -0.07223306 | 0.15900819 | -0.4542725 | 0.6496327 |
| Subparafascicular nucleus | 0.28175208 | 0.204197 | 1.3798052 | 0.1676466 |
| Subparafascicular area | 0.13901204 | 0.4063285 | 0.3421174 | 0.7322625 |
| Medial geniculate complex | -0.66309317 | 0.18269415 | -3.6295259 | 0.0002839422 |
| Lateral geniculate complex | -0.34550338 | 0.07480991 | -4.6184171 | 3.866783E-06 |
| Lateral posterior nucleus of the thalamus | -0.26057861 | 0.16824127 | -1.5488388 | 0.1214205 |
| Posterior complex of the thalamus | -0.9428083 | 0.2205599 | -4.274613 | 1.914696E-05 |
| Posterior limiting nucleus of the thalamus | -0.0730044 | 0.1884145 | -0.387467 | 0.6984105 |
| Suprageniculate nucleus | -0.5607864 | 0.12145996 | -4.6170476 | 3.89238E-06 |
| Anteroventral nucleus of thalamus | -0.015089664 | 0.2340918 | -0.06446045 | 0.9486036 |
| Anteromedial nucleus | 0.8453603 | 0.3236753 | 2.611754 | 0.009007904 |
| Anterodorsal nucleus | -0.2138259 | 0.1929811 | -1.108015 | 0.2678553 |
| Interanterodorsal nucleus of the thalamus | 0.9075771 | 0.3156371 | 2.8753814 | 0.004035397 |
| Lateral dorsal nucleus of thalamus | -0.25263417 | 0.2089296 | -1.2091832 | 0.2265925 |
| Intermediodorsal nucleus of the thalamus | 0.1262187 | 0.2792182 | 0.4520432 | 0.6512379 |
| Mediodorsal nucleus of thalamus | 0.04729071 | 0.14454396 | 0.3271718 | 0.743538 |
| Submedial nucleus of the thalamus | -1.270480169 | 0.3334069 | -3.81059898 | 0.0001386305 |
| Perireunensis nucleus | -0.04428719 | 0.2753777 | -0.1608234 | 0.8722325 |
| Paraventricular nucleus of the thalamus | -0.15887048 | 0.14386119 | -1.1043318 | 0.2694492 |
| Parataenial nucleus | 0.06593954 | 0.14781592 | 0.4460923 | 0.6555306 |
| Nucleus of reuniens | -0.1642305 | 0.19678417 | -0.8345717 | 0.40395888 |
| Central medial nucleus of the thalamus | 0.44009017 | 0.2715193 | 1.6208428 | 0.1050513 |
| Paracentral nucleus | -0.66229299 | 0.2552813 | -2.594365 | 0.009476582 |
| Central lateral nucleus of the thalamus | -0.03732998 | 0.18273196 | -0.2042882 | 0.8381283 |
| Parafascicular nucleus | -0.14712581 | 0.2286622 | -0.64342 | 0.5199516 |
| Reticular nucleus of the thalamus | -0.1621609 | 0.16312768 | -0.9940734 | 0.3201871 |
| Medial habenula | -1.4901657 | 0.2751422 | -5.415984 | 6.095263E-08 |
| Lateral habenula | -0.2880095 | 0.21185299 | -1.3594781 | 0.1739951 |
| Paraventricular hypothalamic nucleus | 0.34239866 | 0.2499379 | 1.369935 | 0.1707072 |
| Periventricular hypothalamic nucleus | -0.43990545 | 0.3694142 | -1.1908191 | 0.2337246 |
| Arcuate hypothalamic nucleus | -0.5137431 | 0.6504901 | -0.7897785 | 0.4296571 |
| Anterodorsal preoptic nucleus | 0.17701998 | 0.23817957 | 0.7432207 | 0.4573481 |
| Anteroventral periventricular nucleus | 0.094201 | 0.3532045 | 0.26670387 | 0.7896972 |
| Dorsomedial hypothalamic nucleus | -0.28261215 | 0.3117986 | -0.9063934 | 0.3647277 |

| | | | | |
|---|---|---|---|---|
| Medial preoptic area | 0.1910699 | 0.22066062 | 0.8658994 | 0.3865453 |
| Subparaventricular zone | 0.3463767 | 0.3744121 | 0.9251216 | 0.3549027 |
| Anterior hypothalamic nucleus | 0.216579834 | 0.322889 | 0.670756347 | 0.5023758 |
| Supramammillary nucleus | -0.1332464 | 0.4822863 | -0.2762807 | 0.7823325 |
| Tuberomammillary nucleus | 0.9623214 | 0.7581502 | 1.269302 | 0.2043335 |
| Medial preoptic nucleus | -0.20390001 | 0.3205789 | -0.636036834 | 0.5247524 |
| Dorsal premammillary nucleus | -0.24365621 | 0.535407 | -0.455086 | 0.6490474 |
| Ventral premammillary nucleus | 0.16194271 | 1.0482246 | 0.1544924 | 0.8772215 |
| Ventromedial hypothalamic nucleus | 0.2055728 | 0.7212893 | 0.2850074 | 0.7756385 |
| Posterior hypothalamic nucleus | -0.07309003 | 0.2936 | -0.2489442 | 0.8034039 |
| Lateral hypothalamic area | 0.09715235 | 0.2786411 | 0.3486648 | 0.727341 |
| Lateral preoptic area | 0.240148 | 0.21322605 | 1.1262602 | 0.2600554 |
| Parasubthalamic nucleus | 0.28498716 | 0.4096517 | 0.6956816 | 0.4866283 |
| Retrochiasmatic area | 0.19213348 | 0.6727916 | 0.2855765 | 0.7752025 |
| Subthalamic nucleus | 0.2403192 | 0.3337832 | 0.7199858 | 0.4715337 |
| Tuberal nucleus | 0.5331735 | 0.6773921 | 0.7870972 | 0.4312249 |
| Zona incerta | 0.32618465 | 0.2347843 | 1.3892952 | 0.164743 |
| Superior colliculus, dorsal part | -0.18425668 | 0.20724965 | -0.8890567 | 0.3739726 |
| Inferior colliculus, central part | -0.46244521 | 0.15770658 | -2.9323139 | 0.003364465 |
| Inferior colliculus, dorsal part | -0.33941438 | 0.2142335 | -1.58431991 | 0.1131209 |
| Inferior colliculus, external part | -0.30316686 | 0.1945784 | -1.5580703 | 0.1192166 |
| Substantia nigra, reticular part | -0.38388666 | 0.1963656 | -1.9549589 | 0.05058792 |
| Ventral tegmental area | 0.09577804 | 0.37263 | 0.2570325 | 0.79715365 |
| Midbrain reticular nucleus | -0.149718552 | 0.17832566 | -0.8395794 | 0.4011443 |
| Superior colliculus, ventral part | -0.1765986719 | 0.11656508 | -1.5150221 | 0.1297668 |
| Periaqueductal gray | -0.030385513 | 0.14565423 | -0.20861402 | 0.8347496 |
| Anterior pretectal nucleus | -0.25850301 | 0.16804901 | -1.5382596 | 0.1239852 |
| Nucleus of the optic tract | -0.383303852 | 0.1616537 | -2.37114163 | 0.01773323 |
| Nucleus of the posterior commissure | -0.3430891 | 0.2262653 | -1.516313 | 0.1294402 |
| Posterior pretectal nucleus | -0.240554146 | 0.1508627 | -1.59452369 | 0.11081879 |
| Cuneiform nucleus | -0.11027139 | 0.1302412 | -0.8466708 | 0.39717862 |
| Red nucleus | 0.1717384 | 0.2328684 | 0.7374911 | 0.4608238 |
| Substantia nigra, compact part | 0.28877213 | 0.3436566 | 0.8402927 | 0.4007443 |
| Pedunculopontine nucleus | -0.275237772 | 0.1435577 | -1.9172623 | 0.05520462 |
| Dorsal nucleus raphe | -0.21628596 | 0.2317819 | -0.9331443 | 0.3507455 |
| Nucleus of the lateral lemniscus | 0.6403633 | 0.3326262 | 1.9251741 | 0.05420758 |
| Principal sensory nucleus of the trigeminal | 0.60017706 | 0.28919202 | 2.0753583 | 0.03795334 |
| Parabrachial nucleus | -0.36033156 | 0.1312303 | -2.7457957 | 0.006036434 |
| Superior olivary complex | 0.10474143 | 0.4023605 | 0.26031735 | 0.794619 |
| Dorsal tegmental nucleus | -0.77704247 | 0.2842224 | -2.7339243 | 0.006258442 |
| Pontine central gray | -0.75282534 | 0.1990634 | -3.7818369 | 0.0001556754 |
| Pontine gray | 0.88957808 | 0.451628 | 1.9697141 | 0.04887114 |

| | | | | |
|---|---|---|---|---|
| Supratrigeminal nucleus | -0.33034561 | 0.1757641 | -1.8794833 | 0.06017853 |
| Tegmental reticular nucleus | 2.380056 | 0.7820305 | 3.0434313 | 0.002338968 |
| Motor nucleus of trigeminal | 0.11458372 | 0.2384628 | 0.4805098 | 0.630865 |
| Superior central nucleus raphe | 2.2464117 | 0.58774 | 3.822118 | 0.0001323103 |
| Laterodorsal tegmental nucleus | -0.41127494 | 0.2341183 | -1.756697 | 0.07896947 |
| Nucleus incertus | -0.3565122 | 0.1977092 | -1.8032156 | 0.07135436 |
| Pontine reticular nucleus | 0.7215137 | 0.3092427 | 2.3331631 | 0.01963959 |
| Dorsal cochlear nucleus | -0.18035708 | 0.2429265 | -0.7424349 | 0.4578239 |
| Ventral cochlear nucleus | -0.5243363 | 0.2551736 | -2.054822 | 0.03989616 |
| Cuneate nucleus | -0.07929914 | 0.5531547 | -0.143358 | 0.8860075 |
| External cuneate nucleus | -0.24536741 | 0.2769971 | -0.8858124 | 0.3757186 |
| Nucleus of the trapezoid body | 0.9518255 | 0.790624 | 1.203891 | 0.2286315 |
| Nucleus of the solitary tract | -0.89927847 | 0.2258871 | -3.98109649 | 6.859809E-05 |
| Spinal nucleus of the trigeminal | -0.260157 | 0.2299203 | -1.1315097 | 0.2578406 |
| Facial motor nucleus | 0.28028045 | 0.3031412 | 0.9245872 | 0.3551807 |
| Dorsal motor nucleus of the vagus nerve | -0.9022889 | 0.3960274 | -2.2783497 | 0.02270575 |
| Gigantocellular reticular nucleus | 0.26926387 | 0.2689686 | 1.00109795 | 0.3167795 |
| Inferior olivary complex | -0.2625265 | 0.9369996 | -0.2801778 | 0.7793411 |
| Intermediate reticular nucleus | -0.3843454 | 0.2206435 | -1.7419295 | 0.08152078 |
| Lateral reticular nucleus | 0.197913 | 0.4872879 | 0.4061521 | 0.6846308 |
| Magnocellular reticular nucleus | 1.3810685 | 1.0719709 | 1.28834513 | 0.1976258 |
| Medullary reticular nucleus | -0.4189387 | 0.2782779 | -1.5054686 | 0.1322036 |
| Parvicellular reticular nucleus | -0.32636472 | 0.1900912 | -1.716885 | 0.08600019 |
| Paragigantocellular reticular nucleus | -0.1285616 | 0.238582 | -0.5388573 | 0.5899853 |
| Nucleus prepositus | -0.014806067 | 0.2625373 | -0.05639604 | 0.9550263 |
| Lateral vestibular nucleus | -0.063305467 | 0.267392 | -0.23675153 | 0.8128496 |
| Medial vestibular nucleus | -0.3718009 | 0.2499002 | -1.4877977 | 0.1368043 |
| Spinal vestibular nucleus | -0.003738046 | 0.2070898 | -0.01805036 | 0.9855987 |
| Superior vestibular nucleus | 0.2505237 | 0.3546777 | 0.7063418 | 0.4799756 |
| Hypoglossal nucleus | -0.8408187 | 0.3035061 | -2.770352 | 0.005599568 |
| Nucleus raphe magnus | 0.09270091 | 0.593278 | 0.156252 | 0.8758344 |

**Supplementary Data Table 66**: Brain regions and corresponding clusters from **Figure S8E**, listed in the same order as the rows of the correlation matrices.

| | **Yellow cluster** | | **Purple cluster** | | **Dark blue cluster** | | **Green cluster** | | **Red cluster** | | **Light blue cluster** |
|---|---|---|---|---|---|---|---|---|---|---|---|
| | Entorhinal area, medial part | | Subiculum | | Paracentral nucleus | | Red nucleus | | Posterior amygdalar nucleus | | Pontine gray |
| | Gigantocellular reticular nucleus | | Spinal nucleus of the trigeminal | | Caudoputamen | | Zona incerta | | Bed nuclei of the stria terminalis | | Orbital area, lateral part |

| | | | | | |
|---|---|---|---|---|---|
| Principal sensory nucleus of the trigeminal | Gustatory areas | Paraventricular nucleus of the thalamus | Posterior auditory area | Basomedial amygdalar nucleus, posterior part | Rostrolateral visual area |
| Nucleus prepositus | Lateral amygdalar nucleus | Reticular nucleus of the thalamus | Primary somatosensory area, trunk | Intercalated amygdalar nucleus | Primary visual area |
| Motor nucleus of trigeminal | Posterior complex of the thalamus | Field CA2 | Primary somatosensory area, nose | Medial amygdalar nucleus | Posteromedial visual area |
| Inferior olivary complex | Medial geniculate complex | Dentate gyrus | Superior central nucleus raphe | Ventral premammillary nucleus | Retrosplenial area, lateral agranular part |
| Paragigantocellular reticular nucleus | Suprageniculate nucleus | Periaqueductal gray | Primary somatosensory area, mouth | Cortical amygdalar area, posterior part | Anteromedial visual area |
| Perirhinal area | Frontal pole | Ventral tegmental area | Submedial nucleus of the thalamus | Lateral septal nucleus | Lateral visual area |
| Retrosplenial area, ventral part | Taenia tecta, ventral part | Parasubthalamic nucleus | Parasubiculum | Endopiriform nucleus, ventral part | Anterolateral visual area |
| Superior colliculus, dorsal part | Pontine central gray | Subthalamic nucleus | Magnocellular nucleus | Basolateral amygdalar nucleus, ventral part | Anterior olfactory nucleus |
| Presubiculum | Laterodorsal tegmental nucleus | Field CA3 | Inferior colliculus, central part | Retrochiasmatic area | Orbital area, ventrolateral part |
| Visceral area | Dorsal tegmental nucleus | Substantia nigra, compact part | Superior colliculus, ventral part | Globus pallidus, internal segment | Posterolateral visual area |
| Ventral auditory area | Supratrigeminal nucleus | Nucleus raphe magnus | Midbrain reticular nucleus | Prosubiculum | Basolateral amygdalar nucleus, anterior part |
| Primary auditory area | Parabrachial nucleus | Pontine reticular nucleus | Nucleus of the lateral lemniscus | Cuneiform nucleus | Anterior cingulate area, ventral part |
| External cuneate nucleus | Nucleus of the solitary tract | Taenia tecta, dorsal part | Parafascicular nucleus | Superior olivary complex | Anterior cingulate area, dorsal part |
| Orbital area, medial part | Dorsal motor nucleus of the vagus nerve | Subparafascicular nucleus | Perireunensis nucleus | Spinal vestibular nucleus | Secondary motor area |
| Pedunculopontine nucleus | Hypoglossal nucleus | Ventral posterior complex of the thalamus | Subparafascicular area | Lateral habenula | Primary somatosensory area, barrel field |
| Nucleus of the optic tract | Parvicellular reticular nucleus | Parataenial nucleus | Supramammillary nucleus | Piriform,amygdalar area | Dorsal auditory area |

| | | | | | |
|---|---|---|---|---|---|
| Posterior pretectal nucleus | Intermediate reticular nucleus | Septofimbrial nucleus | Intermediodorsal nucleus of the thalamus | Tuberomammillary nucleus | Postsubiculum |
| Lateral posterior nucleus of the thalamus | Temporal association areas | Ventral anterior,lateral complex of the thalamus | Posterior hypothalamic nucleus | Tuberal nucleus | Supplemental somatosensory area |
| Anterior pretectal nucleus | Retrosplenial area, dorsal part | Anteromedial nucleus | Arcuate hypothalamic nucleus | Ventromedial hypothalamic nucleus | Prelimbic area |
| Nucleus of the posterior commissure | Central amygdalar nucleus, capsular part | Central medial nucleus of the thalamus | Induseum griseum | Anterior hypothalamic nucleus | Primary somatosensory area, upper lim |
| Central lateral nucleus of the thalamus | Medullary reticular nucleus | Interanterodorsal nucleus of the thalamus | Ventral medial nucleus of the thalamus | Lateral preoptic area | Primary somatosensory area, lower lim |
| Lateral vestibular nucleus | Olfactory tubercle | Lateral reticular nucleus | Nucleus of reuniens | Basolateral amygdalar nucleus, posterior part | Primary motor area |
| Superior vestibular nucleus | Diagonal band nucleus | Magnocellular reticular nucleus | Periventricular hypothalamic nucleus | Subparaventricular zone | Cuneate nucleus |
| Dorsal cochlear nucleus | Inferior colliculus, external part | Postpiriform transition area | Dorsomedial hypothalamic nucleus | Paraventricular hypothalamic nucleus | |
| Ectorhinal area | Globus pallidus, external segment | Field CA1 | Anterior amygdalar area | Agranular insular area, ventral part | |
| Medial septal nucleus | Inferior colliculus, dorsal part | Anteroventral nucleus of thalamus | Substantia innominata | Central amygdalar nucleus, lateral part | |
| Dorsal nucleus raphe | Medial vestibular nucleus | Triangular nucleus of septum | Dorsal premammillary nucleus | Infralimbic area | |
| Lateral dorsal nucleus of thalamus | Medial habenula | Mediodorsal nucleus of thalamus | Anteroventral periventricular nucleus | Tegmental reticular nucleus | |
| Anterodorsal nucleus | Posterior limiting nucleus of the thalamus | Facial motor nucleus | Lateral hypothalamic area | Central amygdalar nucleus, medial part | |
| Nucleus incertus | Substantia nigra, reticular part | Dorsal peduncular area | Medial preoptic area | Basomedial amygdalar nucleus, anterior part | |
| Ventral cochlear nucleus | Lateral geniculate complex | Entorhinal area, lateral part | Medial preoptic nucleus | Nucleus of the lateral olfactory tract | |
| | | | Fundus of striatum | Cortical amygdalar area, anterior part | |

| | |
|---|---|
| Anterodorsal preoptic nucleus | Endopiriform nucleus, dorsal part |
| | Claustrum |
